# A general, highly efficient and facile synthesis of biocompatible rhodamine dyes and probes for live-cell multicolor nanoscopy

**DOI:** 10.1101/2022.06.13.495683

**Authors:** Jonas Bucevičius, Rūta Gerasimaitė, Kamila A. Kiszka, Shalini Pradhan, Georgij Kostiuk, Tanja Koenen, Gražvydas Lukinavičius

## Abstract

The development of live-cell fluorescence nanoscopy is powered by the availability of suitable fluorescent probes. Rhodamines are among the best fluorophores for labeling intracellular structures. Isomeric tuning is a powerful method for optimizing biocompatibility of the rhodamine-containing probes without affecting their spectral properties. However, the efficient synthesis pathway for rhodamine 4-isomers is still lacking. Herein, we present a facile protecting-group-free 4-carboxyrhodamines’ synthesis based on nucleophilic addition of lithium dicarboxybenzenide to the corresponding xanthone. This approach drastically reduces the number of synthesis steps and expands the achievable structural diversity, increases overall yields and permits a cheap gram-scale synthesis of the dyes. We prepared a wide range of symmetric and asymmetric 4-carboxyrhodamines covering the whole visible spectrum and targeted them to multiple structures in living cells – microtubules, DNA, actin, mitochondria, lysosomes, Halo-tagged and SNAP-tagged proteins. The enhanced permeability fluorescent probes operate at submicromolar concentrations allowing high contrast STED and confocal microscopy of living cells and tissues.

## INTRODUCTION

Developments in fluorescence microscopy often rely on fluorescent dyes and probes which are highly photostable, bright and cover a broad spectral range^1^. Multiple super-resolution imaging techniques have become routine experimental tools and are now driving a fundamental knowledge of the molecular biology^2,3^. The transition from standard imaging of fixed samples towards living specimen in super-resolution microscopy is empowered by the discovery of compatible and cell-permeable fluorescent dyes^4–7^, with silicon-rhodamine being the most recognized^8^. In general, rhodamine class dyes and probes are the most often used fluorescent dyes for the labeling of biological targets in living samples^9–12^. This excellent property is stemming from the rhodamines’ dynamic equilibrium between fluorescent zwitterionic (polar) and colorless non-fluorescent (non-polar) spirolactone species. The equilibrium is environmentally sensitive and influenced by pH, ion concentration, enzyme activity, local microenvironment polarity or light^13–15^. The non-polar spirolactone species are responsible for the passive cell membrane permeability. Multiple groups have observed that the target binding is shifting the equilibrium towards fluorescent zwitterionic state^6,7,16,17^. This results in so called fluorogenicity which allows imaging of living samples without washing off the unbound dye. The development of such dyes has recently attracted a lot of attention and multiple efforts were focused on the generation of “tuned” dyes for self-labeling tags (Halo-tag, SNAP-tag etc.) which form a covalent bond with the fluorophore^6,7,18–20^. This process is irreversible and even low cell-membrane permeability could be sufficient for the target labeling. On the contrary, for the reversible binding probes, a certain concentration has to be reached and maintained inside the cell in order to label the target efficiently. Moreover, the reversibly binding probes are susceptible to efflux pumps, which tend to reduce the probe concentration inside the cell through time^21^. From the user’s perspective, the fluorogenic probes which do not require genetic manipulations are the most appealing. However, the design of such tools is undoubtedly challenging. Most often, screening study for the optimal dye-linker-ligand combination is required. Multiple fails are usually due to the low cell membrane permeability, off-targeting, low signal to noise ratio, drastically altered lipophilicity or decreased solubility.

Recently, we have discovered that rhodamines’ 4-isomer based fluorescent probes, in comparison to isomer-5 or -6 derived probes, demonstrate increased cell permeability and reduced susceptibility to efflux pumps^22^. In isomer-4 probes, the spirolactone forming carboxyl group is in close proximity to the amide group (or formerly before conjugation - carboxyl) resulting in the phenomenon, which we named neighboring group effect (NGE). However, the synthesis of such fluorophores required multiple steps and relied on a protection of 3-bromophtalic acid by formation of bis *tert*-butyl ester. *Tert*-butyl ester mainly creates a steric hindrance around carbonyl group and does not mask the partial positive charge resulting in successful lithium-halogen exchange only at low temperature (−116 °C). At such low temperature, dialkylaminoxanthones turned out to be unreactive, thus siloxy xanthones had to be employed, followed by 5 additional synthetic steps^22^. The multistep synthetic path resulted in relatively low total yield of 13-21% and limited structural diversity (Figure 1). Recently, Butkevich A. N. proposed an alternative synthetic strategy relying on nucleophilic addition of bis-aryllanthanum species to 3-bromophtalic anhydride, followed by palladium catalyzed carbonylative hydroxylation resulting in a total 24-45% yield^23^ (Figure 1). While this method did provide 4-carboxy rhodamines in less steps and in higher total yield, we found that many starting compounds, especially unsymmetrical dibromides, are challenging or even not possible to obtain.

**Figure 1.**
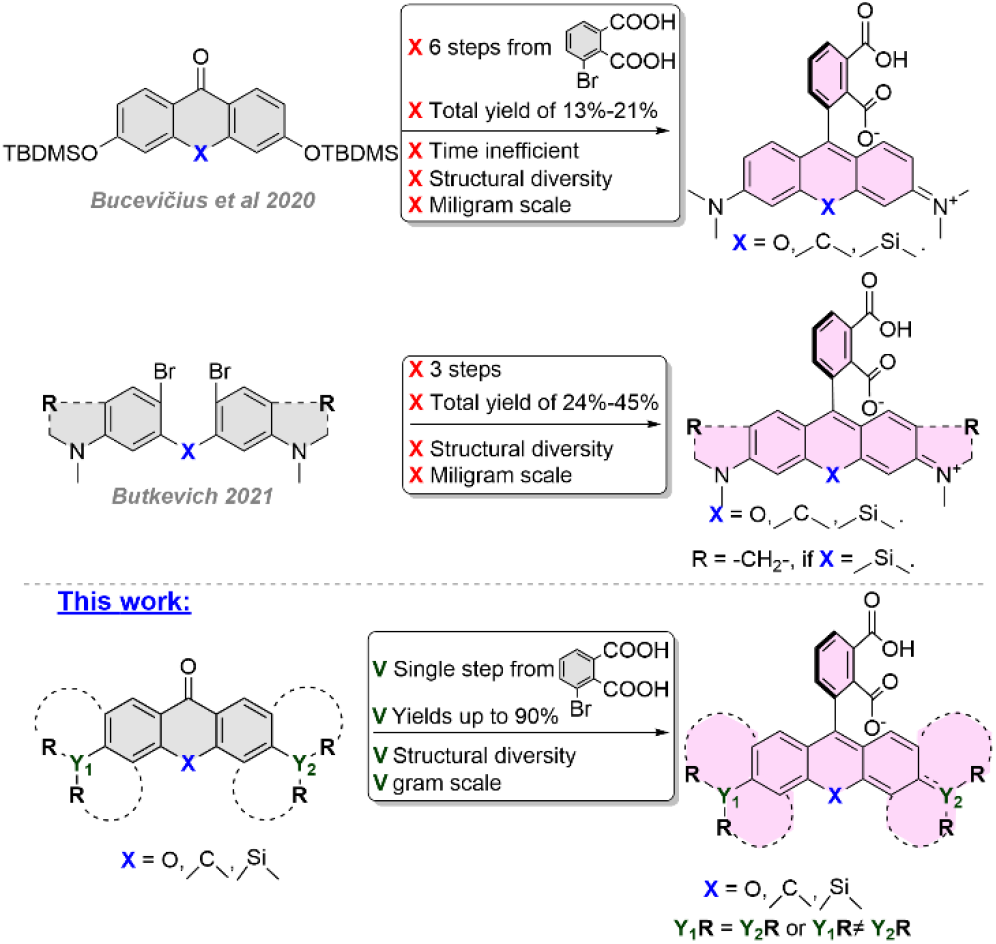
Synthetic approaches to 4-carboxy rhodamines and their analogs.

We aimed to further simplify the synthesis of rhodamine 4-isomers by eliminating the need of protecting groups. As a result, herein we report facile, efficient, protecting-group-free and scalable synthesis of structurally diverse 4-carboxyrhodamines. This allows to systematically examine performance of fluorescent probes derived from symmetrical and unsymmetrical fluorophores in living specimens. We exploit the dyes’ structural diversity for selection of highly biocompatible probes and apply them for multi-color confocal microscopy and STED nanoscopy imaging of living cells and tissues. High cell-membrane permeability of the newly synthesized dyes allows target staining at low nanomolar concentrations with excellent specificity, whereas spectral tuning reduces the need of high STED laser power. Our results clearly demonstrate the high potential of rhodamines’ 4-isomers for the development of biocompatible fluorescent probes.

## RESULTS AND DISCUSSION

### Synthesis of 4-carboxyrhodamine dyes

*In situ* formed carboxylic acid salts may be applied to suppress carbonyl group reactivity towards nucleophiles^24^, but the polar carboxylate salts tend to have low solubility in ethereal solvents, which are typically used in lithium halogen exchange reactions. We have prepared and investigated the solubility of 3-bromophthalic acid salts in anhydrous tetrahydrofuran (THF) and diethyl ether (Et_2_O). Neither disodium nor dilithium 3-bromophthalates were soluble. Monosodium salt showed little solubility, while monolithium salt was sufficiently soluble in THF at ambient temperature. Lithium hydrogen phthalate stands out from other phthalate salts with one of the shortest intramolecular hydrogen bonds observed (reported O-O distance 2.385Å) and a planar phthalate ion structure^25^, which both are strong indications of significantly reduced acidity, especially in organic solvents. This encouraged us to perform lithium-halogen exchange with lithium hydrogen 3-bromophthalate (Table 1, **II**) under standard conditions followed by quenching with excess of I2 and LC/MS analysis. Upon addition of 1 eq of *n*-BuLi no significant dilithium salt precipitation was observed and the solution turned light yellow, indicating some formation of ArLi species. LC/MS analysis revealed that the obtained mixture consisted of unreacted starting bromide, phthalic acid and a trace of 3-iodophthalic acid (Table 1, entry 1). Increasing *n*-BuLi to 2 eq. resulted in 64% conversion to 3-iodophthalic acid (Table 1, entry 2). After first successful attempts, we switched the starting material from lithium hydrogen 3-bromophthalate (Table 1, **II**) to 3-bromophthalic acid (Table 1, **I**) as the monolithium salt can be formed *in situ* by neutralization of 3-bromophthalic acid with additional equivalent of *n*-BuLi. Thus, addition of 2 eq of *n*-BuLi resulted in only trace formation of 3-iodophthalate (Table 1, entry 3), but 55-57% conversion to 3-iodophthalate was achieved once 2.5 or 3 eq of *n*-BuLi were used (Table 1, entry 5-6). This indicates that >2.5 eq are optimal BuLi amount for efficient formation of ArLi species. Inspired by these results, we decided to further examine nucleophilic addition of the formed ArLi species to the silaxanthone **K2** (Table 2), which should afford **4-SiR-COOH** (**2**) fluorophore in a single step without any protecting groups or any need of functional group interconversions. First, we used stoichiometric ratio of reactants: 1 eq of 3-bromophthalic acid and 3 eq of *n*-BuLi followed by an addition of 1 eq of ketone **(K2**). The obtained mixture was analyzed with LC/MS indicating 62% conversion of ketone, but two products were formed: target compound **2** and the *n*-BuLi addition product **1** with a ratio of 6:4 (Table 2, entry 1). Formation of **1** indicated that unreacted *n*-BuLi was still present in the reaction mixture. Reduction of *n*-BuLi amount to 2 eq resulted in decreased conversion of **K2** to 5% and a trace formation of side product **1** (Table 2, entry 2). Usage of 2.5 eq resulted in increased conversion to 26% and retained selectivity ratio (Table 2, entry 3), indicating optimal *n*-BuLi quantity. Abatement of ketone to 0.5 eq and eventually to 0.33 eq led to optimal conditions with 95% conversion of **K2** and with a 98:2 selectivity ratio resulting in an excellent 90% yield of **4-SiRCOOH** (**2**) after flash column purification (Table 2, entry 4-5).

**Table 1.** Optimization of Li-halogen exchange reaction conditions of 3-bromophthalic acid

| Entry | Starting | n-BuLi eq. | A (%)* | B (%)* | C (%)* |
| --- | --- | --- | --- | --- | --- |
| 1 | II | 1.0 | 69 | trace | 30 |
| 2 | II | 2.0 | 3 | 64 | 33 |
| 3 | I | 2.0 | 55 | trace | 44 |
| 4 | I | 2.2 | 33 | 7 | 60 |
| 5 | I | 2.5 | 4 | 55 | 41 |
| 6 | I | 3.0 | 1 | 57 | 32 |

\*Conversion was determined by LC/MS analysis

**Table 2.**
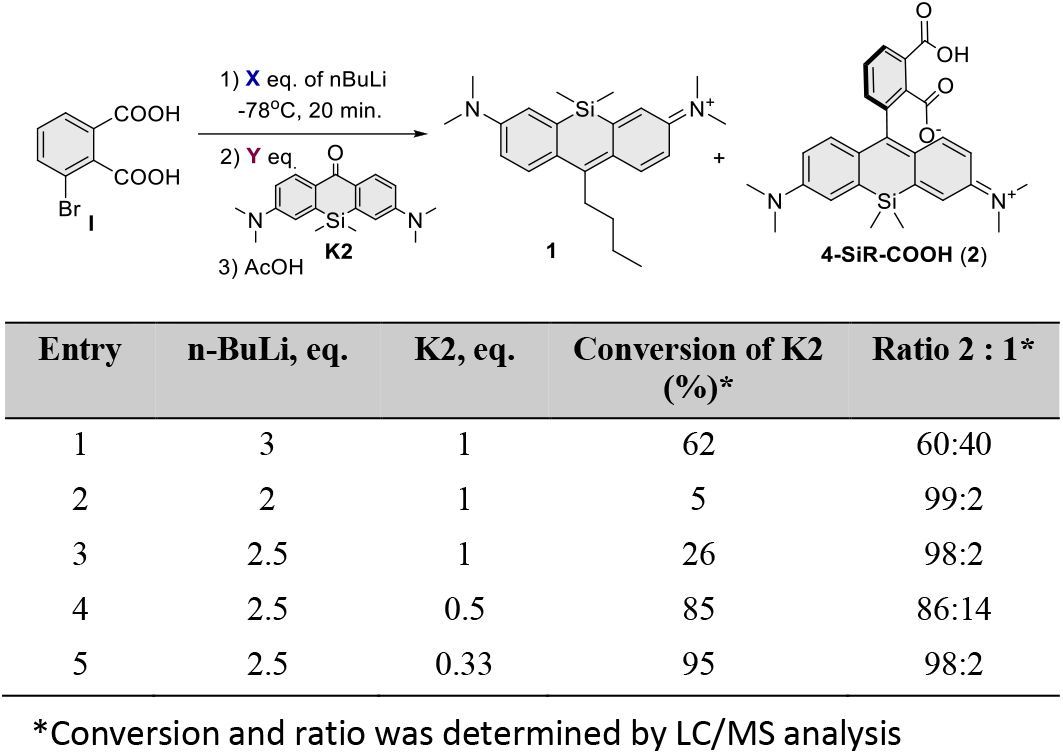
Optimization of nucleophilic addition reaction conditions to silaxanthone.

Relying on the data obtained in optimization of the experimental conditions, we propose the following reaction course (Scheme 1, A). In STEP A lithium 3-bromophthalate (**II**) is formed after addition of 1 eq *n*-BuLi to 3-bromophthalic acid (**I**). In STEP B, due to significantly reduced acidity of **II**, reaction can follow thermodynamic or kinetic paths forming either dilithium salt **III** or ArLi species **IV** respectively. In our favor, thermodynamic product was not dominant even at −78°C and reaction mainly followed a kinetic pathway yielding **IV**. Once formed, ArLi species **IV** are metastable and may self-quench by intramolecular proton transfer forming dilithium phthalate **V** or the proton can be neutralized by additional *n*-BuLi and form highly reactive trianionic ArLi species **VI**. These reaction paths are competing, which results in non-stoichiometric requirement of *n*-BuLi. We did not observe significant degradation of **VI** in THF at −78°C at least for 40 min, and 20 min of aging is sufficient. Due to multiple reaction paths, **VI** forms from starting compound **I** non-stoichiometrically and reduced ratio of electrophilic reactive partner (xanthone) has to be used (Scheme 1, A).

In order to demonstrate the scope of the developed synthesis method, we focused on preparation of a series of structurally diverse xanthones **K2**-**K21** (Figure S1). Xanthones **K2-K4** were synthesized as previously published^8,26,27^. Xanthones with a carbon bridging atom (**K6, K7** and **K10-K12**) were synthesized by Lewis acid mediated condensation of substituted benzyl alcohols and substituted α-methylstyrenes, followed by Brønsted acid initiated cyclization and finally oxidation with KMnO_4_ (Scheme S1). The symmetrical silaxanthones (**K8-K9**) were obtained by condensation of two brominated aniline type units with formaldehyde followed by dilithiation and substitution with Me2SiCl2 to introduce a bridging fragment and lastly temperature controlled oxidation with KMnO_4_ (Scheme S2, a). For unsymmetrical silaxanthones (**K13-K16**), first step was performed by BF3-OEt2 mediated condensation^28^ of bromo substituted benzylic alcohols with the 3-bromoaniline type counter partner and further followed by the same synthetic steps as for symmetrical ketones (Scheme S2, b). Xanthones with methoxy and allyl amino substituents (**K17**-**K21**) were obtained by alkylation of hydroxyxanthones or by nucleophilic aromatic substitution of 3,6-bistriflate xanthones with corresponding amines (Scheme S3).

**Scheme 1.**
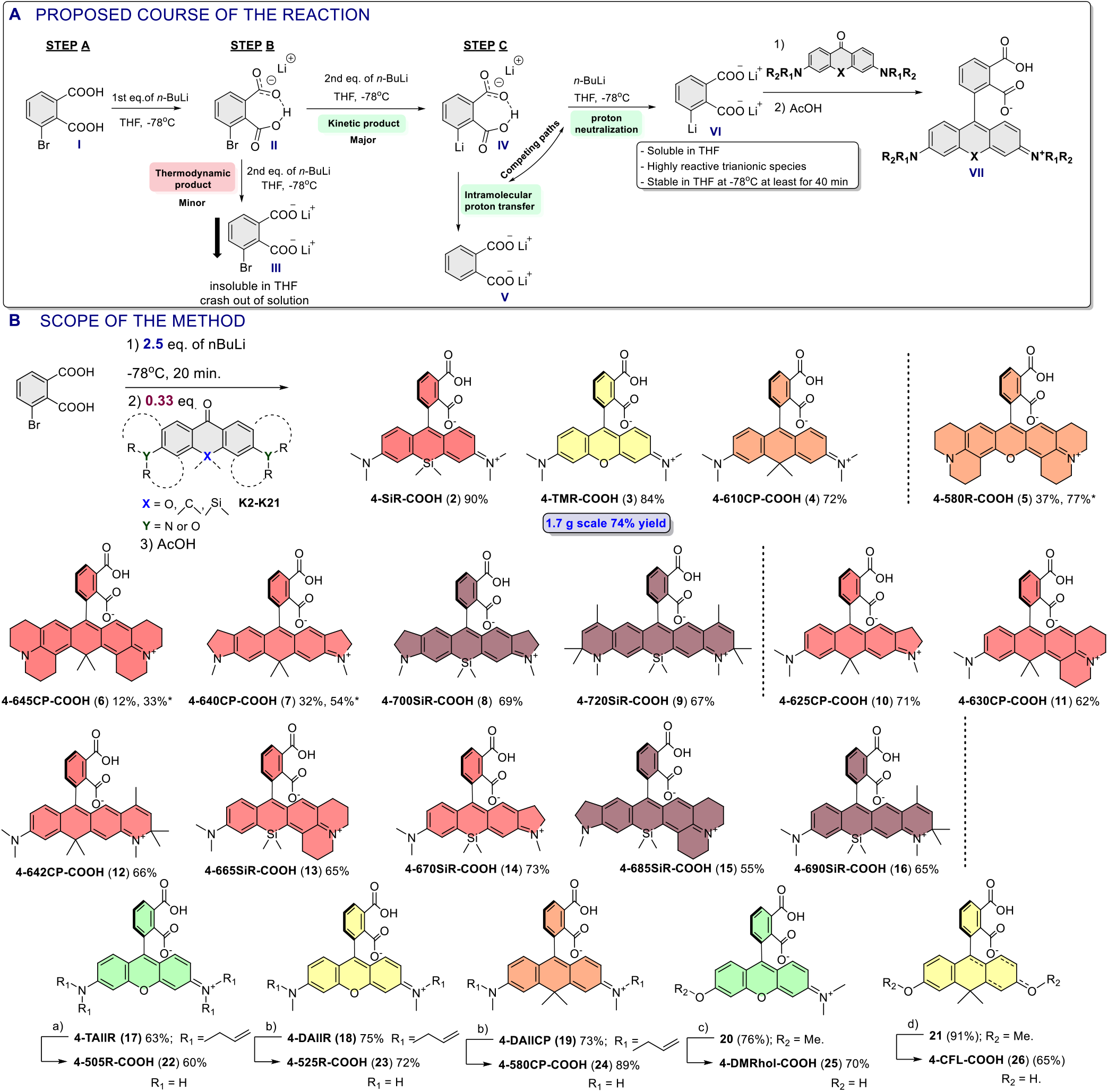
Reaction course and scope. A. Proposed course of the lithium halogen exchange reaction of 3-bromophthalic acid B. The scope of the described 4-carboxyrhodamines’ synthetic method a) 14 mol% Pd(PPh_3_)_4_, 7 eq NDMBA, MeCN. 45°C, 10h b) 7 mol%, Pd(PPh_3_)_4_, 2.5-3 eq NDMBA, MeCN, 40 °C, overnight c) 40 eq BBr_3_, 1,2-DCE, 55 °C, 24h d) 40 eq BBr_3_, 1,2-DCE, 55 °C, 72h.. *Marks the yields when 0.167 eq (instead of 0.33 eq) of ketone reactant was used.

With the key intermediates **K2-K21** in hand, we explored the scope of the developed 4-carboxyrhodamine synthesis method (Scheme 1, B). Classical oxygen, carbon or silicon bridged rhodamines (**2**-**4**) were obtained in significantly fewer synthetic steps and 72-90% yields, which is 3-7 times higher compared to previously published synthetic procedures^22,23^ (Scheme 1, B). In addition, the synthetic method was found to be compatible with ketones possessing amino groups fused to the aromatic ring (Figure S1, **K5-K19**), providing compounds **5-9** in 12-77% yields. Reduced yields for **5**-**7** can be attributed to the reduced solubility of highly rigid symmetrical ketones **K5**-**K7.** They tend to crash out of the mixture resulting in heterogeneous conditions thus prolonged reaction times which are incompatible with the stability of ArLi species **VI**. Increase of 3-bromophthalic acid to 6 eq (abatement of ketone to 0.167 eq) recovered reaction yields (Scheme 1, B, yields marked with *). Unsymmetrical ketones **K10**-**K16** showed a good solubility in THF resulting in usual high 55-71% yields for compounds **10**-**16**. In addition, rhodamines possessing primary (**22**) or secondary amino (**23**-**24**) substituents were obtained via palladium catalyzed allyl group cleavage of the compounds with allyl amino substituents **17-19.** Similarly, rhodol (**25**) or carbofluorescein (**26**) were obtained from methoxy protected intermediates **20**-**21** after BBr3 mediated demethylation (Scheme 1, B).

Finally, we demonstrated scalability of the developed synthetic method by performing a gram-scale synthesis of the **4-TMR-COOH (3)** dye. We obtained 1.7g of product from a single batch in 74% yield after flash column purification without any scrutinized scale-up optimizations. In summary, the described facile, highly efficient and scalable synthetic method provides access to previously inaccessible structurally diverse 4-carboxyrhodamines and their analogs with wide range of photophysical properties.

### Synthesis of fluorescent probes

Rhodamine dyes with a pendant carboxyl group can be attached to the small molecule ligands/inhibitors to provide fluorescent probes with high specificity in cellular staining^9^. We demonstrate the advantages of increased cell-permeability of isomer-4 dyes by conjugating them via peptide coupling to the range of targeting moieties (Scheme 2) and obtaining a library of fluorescent probes targeting SNAP-tag (**27**-**46, 105-109**), β-tubulin (**47**-**67**), Halo-tag (**68**-**79**), mitochondria (**80**-**86**), DNA (**87**-**92**), lysosome (**93**-**99**) and actin (**100**-**104**).

**Scheme 2.**
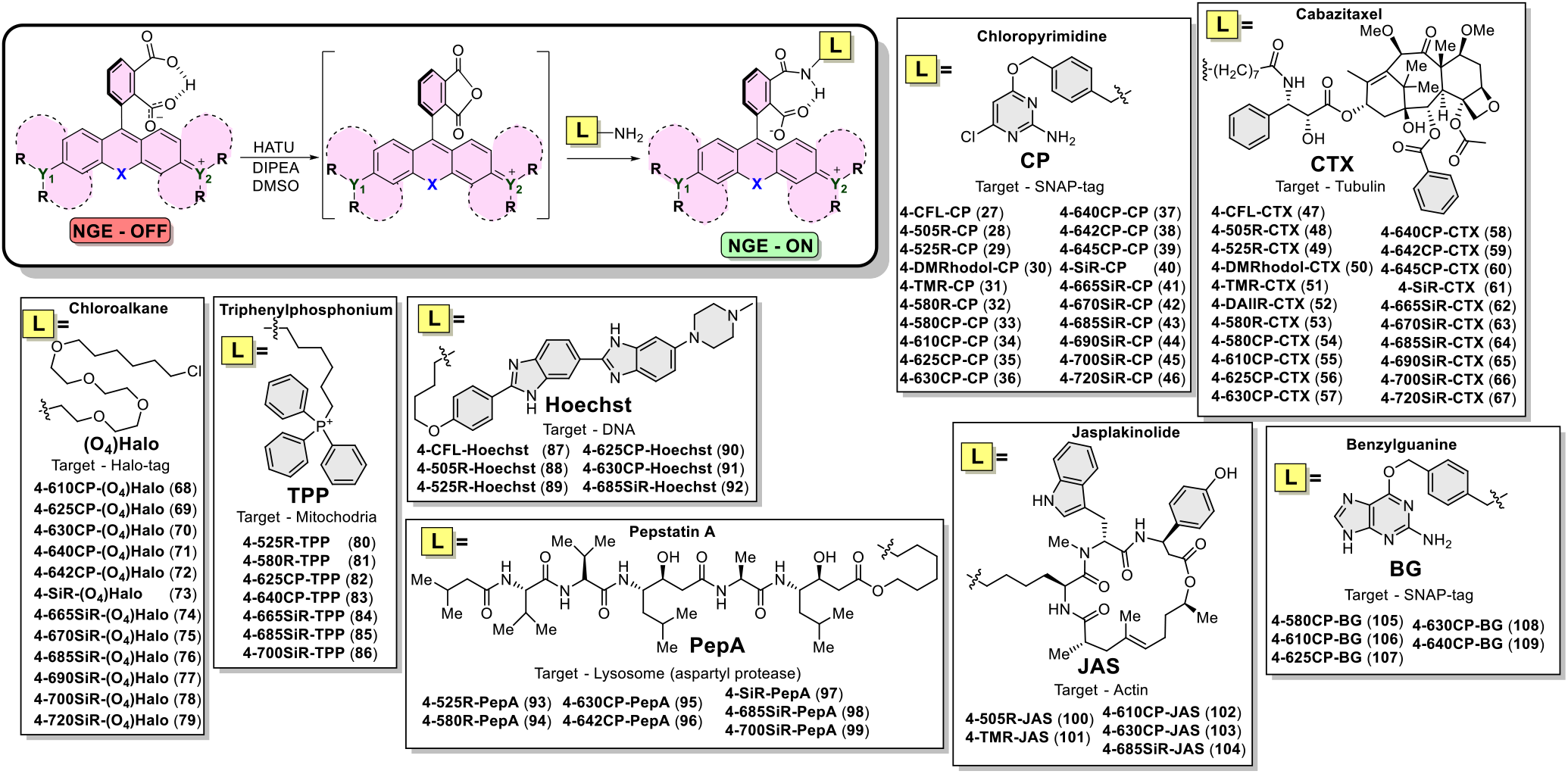
Synthesis of a library of STED imaging compatible fluorescent probes targeting different intracellular targets.

### Characterization of the fluorescent dyes and probes in vitro

The synthesized dyes were characterized by means of absorption and fluorescence spectroscopy, fluorescence quantum yield and lifetime measurements in phosphate-buffered saline (PBS) (Table 3). Additionally, photophysical properties were assessed with 0.1% sodium dodecyl sulfate additive to characterize the dyes avoiding any possible dye aggregation effects (Table S1). The absorption maxima of obtained dyes are in the range from 500 nm to 720 nm and corresponding emission maxima in the range from 522 nm to 757 nm. This coverage is achieved via introduction of substituted amino groups fused to the aromatic ring (indoline, julolidine, dihydroquinoline), variation of amino group substitution pattern (tertiary, secondary, primary), alternation of bridging atom (O, C, Si) and desymmetrization of rhodamine core (**10-16**) (Table 3 and Table S1).

**Table 3.**
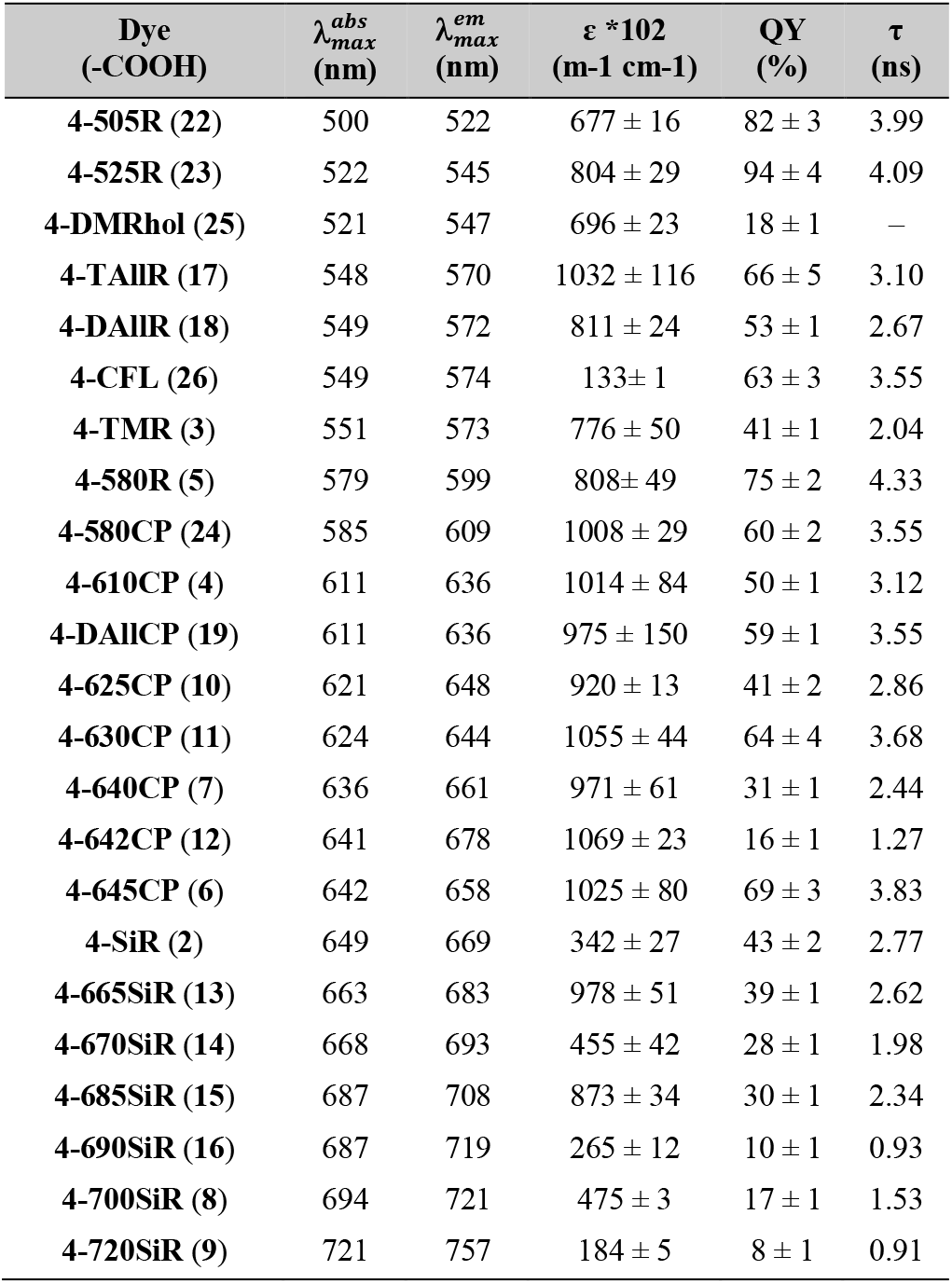
Photophysical properties of the synthesized 4-carboxyrhodamines in PBS at pH =7.4.

To characterize the ability of the synthesized fluorophores and probes to switch between spirolactone and zwitterion states, we measured the absorbance in water – 1,4-dioxane mixtures with known dielectric constants^29^. We fitted the absorbance change to the dose response equation and obtained ^dye^D_50_ value, which indicates the dielectric constant at which absorbance is halved (Figure 2a, S2 and Table S2).

**Figure 2.**
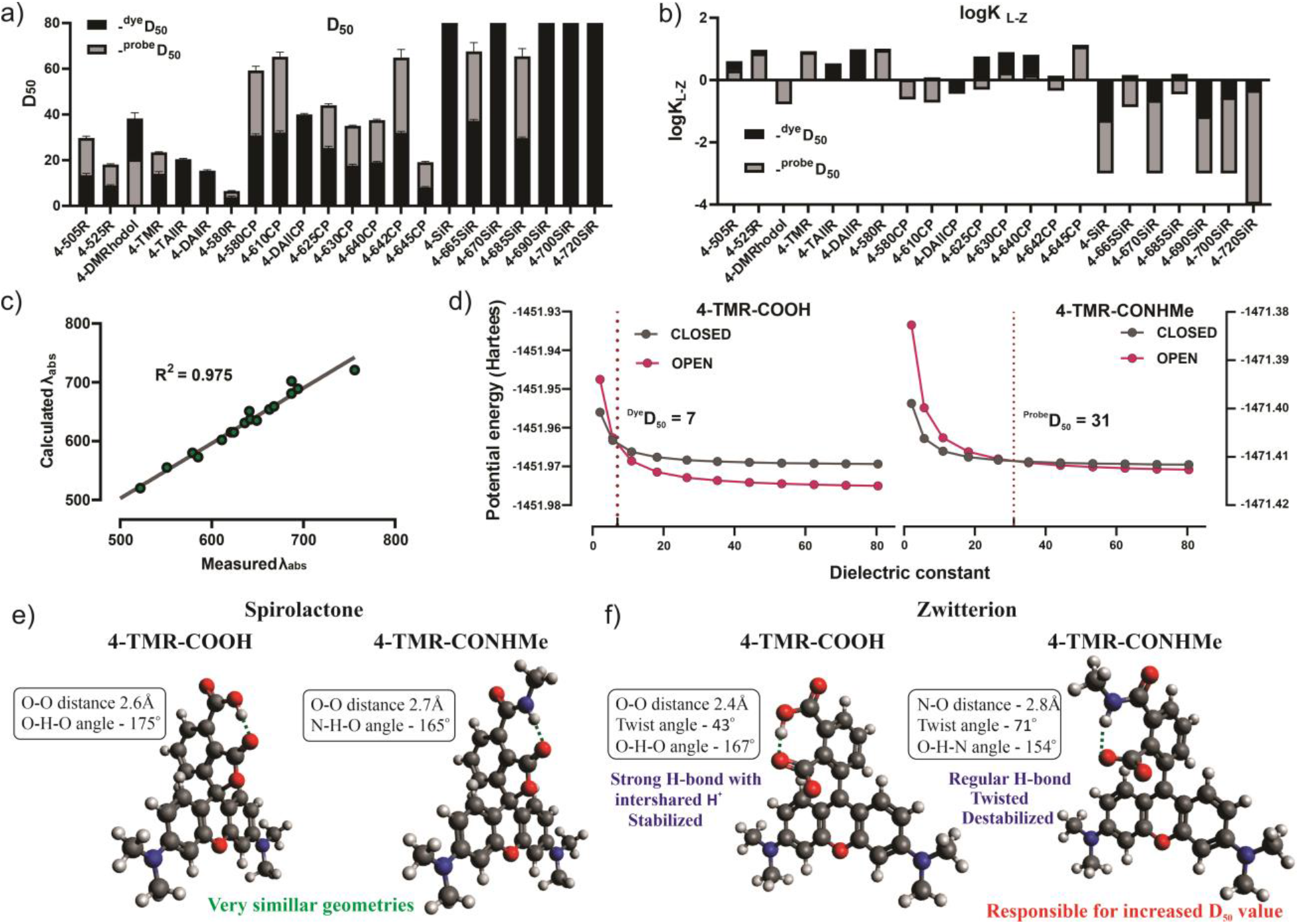
*In vitro* and *in silico* characterization of the dyes. a) ^dye^D_50_ and ^probe^D_50_ and b) ^dye^logK_L-Z_ and ^probe^logK_L-Z_ values represented in stacked bar chart c) Correlation between TDDFT calculated and experimentally measured absorption maxima of the dyes in water d) Plots of DFT calculated potential energies of zwitterion and spirolactone states against simulated dielectric constant, intersection point of the curves represent a simulated D_50_ value e) DFT B3LYP 6-311++G(d,p) IEFPCM optimized geometries of 4-TMR-COOH and 4-TMR-CONHMe spirolactone and f) zwitterion forms in water.

Fluorescent substrates of the self-labeling tags offer a lot of flexibility for the designing of imaging experiments. 4-(benzyloxy)-2-chloropyrimidine (CP) ligand is structurally simple and commercially available SNAP-tag ligand (Scheme 2). Thus, we synthesized a full range of substrates via peptide coupling (**27**-**45**) and used this series to compare the ^dye^D_50_ and the ^probe^D_50_ values. In all cases, ^probe^D_50_ was significantly increased compared to the corresponding ^dye^D_50_, indicating that for all tested dyes the conversion of the carboxyl group to amide induces higher spirolactone content due to NGE^22^. We found that D_50_ values are mainly affected by the bridging atom O < C < Si and, to a lower extent, by the side substituents. In respect to side substituents D_50_ increased in the line: julolidine < indoline < dihydroquinoline ≈ (dimethyl)amino. Interestingly, spirolactone form was dominating in water for majority of the silarhodamines resulting in D_50_ values above or close to 80 (Figure 2a and Table S2), except the ones possessing julolidine fragment (**4-665SiR** and **4-685SiR**).

For this reason we decided to measure alternative ^dye^K_L-Z_ and ^probe^K_L-Z_ values, which can also be used to characterize dye’s propensity to exist in spirolactone or in zwitterion form^6^ (Figure 2b and Table S2). Surprisingly, we did not observe any distinct correlation between D_50_ and K_L-Z_ values. Probably because both concepts have no direct physical link to the true equilibrium constants and are just empirical methods, which provide a numerical value for basic comparison of rhodamine dye’s environmental sensitivity. We noticed that concept of K_L-Z_ proved to be more suitable to assess dyes which are highly closed in water environment, but fail to give comparative values for the dyes which mainly exist in zwitterion form already in 1:1 water-dioxane mixtures (ε = 35). Vice versa, D_50_ concept proved to be more suitable to assess rhodamine dyes, which preferentially exist as zwitterions in water or the dyes which show little sensitivity to the change of surrounding media.

**4-CFL** dye belongs to the fluorescein dyes class and did not demonstrate distinct response to water content in 1,4-dioxane. It is known that fluoresceins are highly sensitive to pH. We measured a pH dependence on absorption of **4-CFL-COOH** dye and found that the dye partially exist in open-fluorescent form at physiological pH range and is suitable for live cell microscopy (Figure S3).

### Characterization of the fluorescent dyes and probes in silico

We performed density functional theory (DFT) calculations, using the Gaussian 09 program package^30^, to estimate the influence of substituents on rhodamines’ core to absorption properties and the impact to the energies of frontier molecular orbitals (FMO’s). The geometries of studied compounds were optimized using B3LYP exchange–correlation hybrid functional together with 6**-** 311++G(d,p) basis set and water was implemented with IEFPCM solvation model^31^. We calculated energies and generated isodensity surface plots of the frontier molecular orbitals. It was found that side substituents and bridging atom has minimal influence to LUMO energies. Whereas systematic increase in HOMO energies was evident in line with increasing donor strength of the substituents and bridging atom, which resulted in steadily decreasing HOMO-LUMO gap and a gradual red shift in absorption (Figure S4). The absorption maxima wavelengths in water (IEFPCM) were calculated from the vertical excitations obtained by TDDFT calculations^32^. It is known that TDDFT gives a systematic error in calculation of vertical excitation energy values for the dyes with methyne fragment ^33^. After applying systematic empirical correction factor of 0.4 eV, the calculated absorption values were in very good agreement with the experimentally measured values for all studied dyes (Figure 2c, Table S3). The obtained calculation data gave us insight to include unsymmetrical rhodamines into the study.

Previously we have compared DFT calculated free Gibb’s energies of isomer 4/5/6-rhodamine dyes and probes upon cyclisation of zwitterion to spirolactone in different surrounding environments^22^. But we had overlooked to study the key reason - why spirolactone-zwitterion equilibrium of 4-isomer rhodamines is drastically affected upon carboxyl group (^dye^D_50_) interconversion to amide (^probe^D_50_). In order to get better insight of how NGE works, we performed potential energy calculations along the increasing dielectric constant for zwitterion and spirolactone forms of **4-TMR-COOH** and of a hypothetical model compound **4-TMR-CONHMe**. The latter compound served as a model for probe with a truncated linker and ligand. The obtained energy values of both forms were plotted against dielectric constant (Figure 2d). The intersection point of two curves corresponds to the equal potential energies, hence equal distribution of spirolactone and zwitterion species, and can be regarded as the simulated D_50_ value. The simulated ^dye^D_50_ value was 7 and ^probe^D_50_ - 31, which is in a rather good agreement with the experimentally obtained values - 14 and 23 respectively and an indication that DFT is able to predict NGE behavior. Based on the data obtained from DFT calculations, we hypothesize that the spirolactone form is stabilized by the intramolecular hydrogen bond (Figure 2e) in both the dye (**4-TMR-COOH)** and the probe (**4-TMR-CONHMe)** on a considerably similar extent. Hence, the major difference could be attributed to the (de)stabilizing effects of zwitterion form. Taking into consideration the unprecedented short intramolecular hydrogen bond length in **4-TMR-COOH** (Figure 2f) it can be considered highly stabilizing as the proton can be shared by both carboxyl groups, which is also supported with previous observations based on x-ray data of lithium hydrogen phthalate^25^. Whereas in the case of **4-TMR-CONHMe**, amide hydrogen is not labile and cannot be intershared with carboxylate group due to large basicity difference of conjugate bases. This results in the formation of a standard length (strength) hydrogen bond and more twisted carboxylate group (Figure 2f), which destabilizes zwitterion form and results increased D_50_ value of the probe. To sum up, an increase in ^probe^D_50_ value (in respect to ^dye^D_50_) is resulting from the loss of the shared proton stabilizing effect between amide and carboxylate groups in zwitterion form of the probe.

### Substrates for self-labeling tags

SNAP-tag is a modified human O6-alkylguanine-DNA alkyltransferase (hAGT) accepting CP and O6-Benzylguanine (BG) derivatives^34,35^. To comprehensively cover the variety of self-labeling tag substrates, additionally to the synthesized CP probes **27**-**46**, we have conjugated the selected fluorophores to benzylguanine (BG) to yield compounds **105-109**. A recent study reported that more hydrophobic CP substrates show 4-14-fold slower reaction kinetics compared to similar BG substrates^36^ and suggesting that BG substrates might perform better in respect of labeling efficiency. In addition, we have also synthesized a series of chloroalkane-PEG4 derivatives **68-79** ((O_4_)Halo) that react with one of the most frequently used self-labeling tag – Halo-tag. It is based on engineered bacterial dehalogenase (DhaA from Rhodococcus sp.), which is accepting halogenated alkanes as substrates^37^. Further, we examined the reactivity of all fluorescent substrates by incubating them with two times excess of respective tag protein and detected if protein labeling took place on denaturing polyacrylamide gel (SDS-PAGE). Most of the CP-based substrates showed a successful reaction with SNAP-tag protein, though **4-SiR-CP**, **4-665SiR-CP**, **4-670SiR-CP** substrates showed incomplete reaction and **4-690SiR-CP**, **4-720SiR-CP** showed no reaction (Figure S5). All of the BG-based substrates reacted with SNAP-tag protein (Figure S6). Similarly, in case of Halo-tag substrates, except for **4-720SiR-(O_4_)Halo** and **4-690SiR-(O_4_)Halo** probes, which showed no or incomplete reaction (Figure S7).

We also determined the fluorogenicity – increase in fluorescence and absorbance upon reacting with the target protein. All of the SNAP-tag interacting substrates demonstrated absorbance and fluorescence increase upon reacting with SNAP-tag protein, except **4-CFL-CP**. In the CP series the **4-685SiR-CP** and **4-700SiR-CP** showed largest, more than 14 times increase in absorbance and more than 80 times increase in fluorescence signal (Figure S8 and S9). In the BG series **4-630CP-BG** demonstrated 8 times increase in absorption and 6 times increase in fluorescence signal (Figure S10). We noticed that CP-based SNAP-tag substrates demonstrated higher fluorogenicity *in vitro* compared to corresponding BG series. **4-685SiR-(O_4_)Halo** and **4-700SiR-(O_4_)Halo** showed highest (~20 times) increase in fluorescence signal in Halo-tag substrate series (Figure S11). In general, fluorescence increase after reaction with target protein of the studied self-labeling substrates was in line with increasing D_50_ value, but the extent of fluorescence increase is highly depended on the attached targeting moiety (CP, BG or (O_4_)Halo).

To test the performance of the self-labeling tag substrates *in cellulo,* we used engineered U-2 OS cells expressing histone-H3-SNAP (Figure S12), Nup96-SNAP ^38^ or vimentin-Halo ^39,40^. We observed less efficient staining of SNAP-tagged histone H3 (Figure S13) by CP probes compared to BG derivatives^36^. We identified **4-625CP-BG** and **4-630CP-BG** as the best performing substrates. High staining efficiency allowed acquisition of high-quality STED microscopy images (Figure S14). All Halo-tag substrates, except **4-690SiR-(O_4_)Halo** and **4-720SiR-(O_4_)Halo**, efficiently labeled vimentin-Halo-tag (Figure S15). The comparative row of staining intensities from the dimmest to the brightest is: **690SiR**, **720SiR** = no staining <<< **4-SiR** < **665SiR** ≤ **670SiR** < **700SiR** ≤ **685SiR** < **642CP** < **625CP** ~ **630CP** ~ **640CP-(O_4_)Halo** (**68**-**79**).

### Tubulin fluorescent probes

Our previous study showed that the incorporation of rhodamines’ 4-isomers enhances cell-membrane permeability of fluorescent probes^22^. We were interested to see whether this conclusion is a general rule valid for a wide variety of fluorophores. Use of the self-labeling tags for this experiment is suboptimal as irreversible reaction with the target makes the staining efficiency less dependent on substrate permeability. Thus, we synthesized a complete series of cabazitaxel (CTX) derivatives targeting tubulin.

We found that almost all probes, except **4-580R-CTX**, were able to stain microtubules in living cells at nanomolar concentrations (Figure 3a). Indeed, the majority compounds (>75%) displayed EC_50_ <100 nM indicating good permeability of the probes (Figure 3b and S16). The best performing tubulin probes were **4-625CP-CTX** and **4-630CP-CTX**, which provided excellent staining at concentration of 10 nM within 1 h (Figure 3c and S17). We observed 20 and 12 signal-to-background ratio in the acquired confocal and 3D STED images of microtubules stained with 10 and 100 nM **4-625CP-CTX** or **4-630CP-CTX**, respectively. Only two-fold signal intensity difference was observed after 10-fold increase in probe concentration (10 nM vs 100 nM, Figure S17). This observation is in excellent agreement with low nanomolar EC_50_ values (Figure 3b, Table S4) and demonstrates labeling saturation. It should be noted that for long term staining, lower concentrations are beneficial because of reduced potential cytotoxicity effects. Furthermore, high efflux pump activity in Vero and U-2 OS cell lines is not hampering microtubule staining with **4-625CP-CTX** and **4-630CP-CTX** probes (Figure S18 and S19). Microtubules are well-defined tubular structures with a diameter of ~20 nm in mammalian cells. We exploited tubulin probes for evaluation of the STED microscopy performance of the new series of dyes (Figure S20). We found that depletion efficiency (smaller I_sat_) is increasing with the shift of emission spectrum maximum towards 775 nm STED laser (Figure 3d, Table S4). Such observations are in agreement with stimulated emission theory^41^. Fluorophores with smaller I_sat_ can operate at lower STED laser powers, thus provide increased resolution and reduce potential phototoxicity. The smallest apparent FWHM of microtubule equal to 21 ± 9 nm was obtained when living cells were stained with **4-665SiR-CTX** probe (Figure 3e and S20, Table S4). This value is close to 16 ± 5 nm obtained using MINFLUX method in fixed cells^42^. Slightly further red shifted **4-670SiR-CTX** probe already demonstrated a slightly worse resolution due to the occurring re-excitation by 775nm STED laser (Figure S20). The 3D STED imaging experiments demonstrated excellent performance of **4-625CP-CTX** and **4-630CP-CTX** probes. The measured apparent FWHM was in the range of 100 - 150 nm along all axes (Table S5, Figure S17). Despite low fluorogenicity (based on low D_50_ value), most of the probes show ratio above 10 of fluorescence signal increase on microtubule compared to cytosol (Table S4 and Figure S20). This observation highlights the highly specific staining of microtubules and high cell-permeability of the probes.

**Figure 3.**
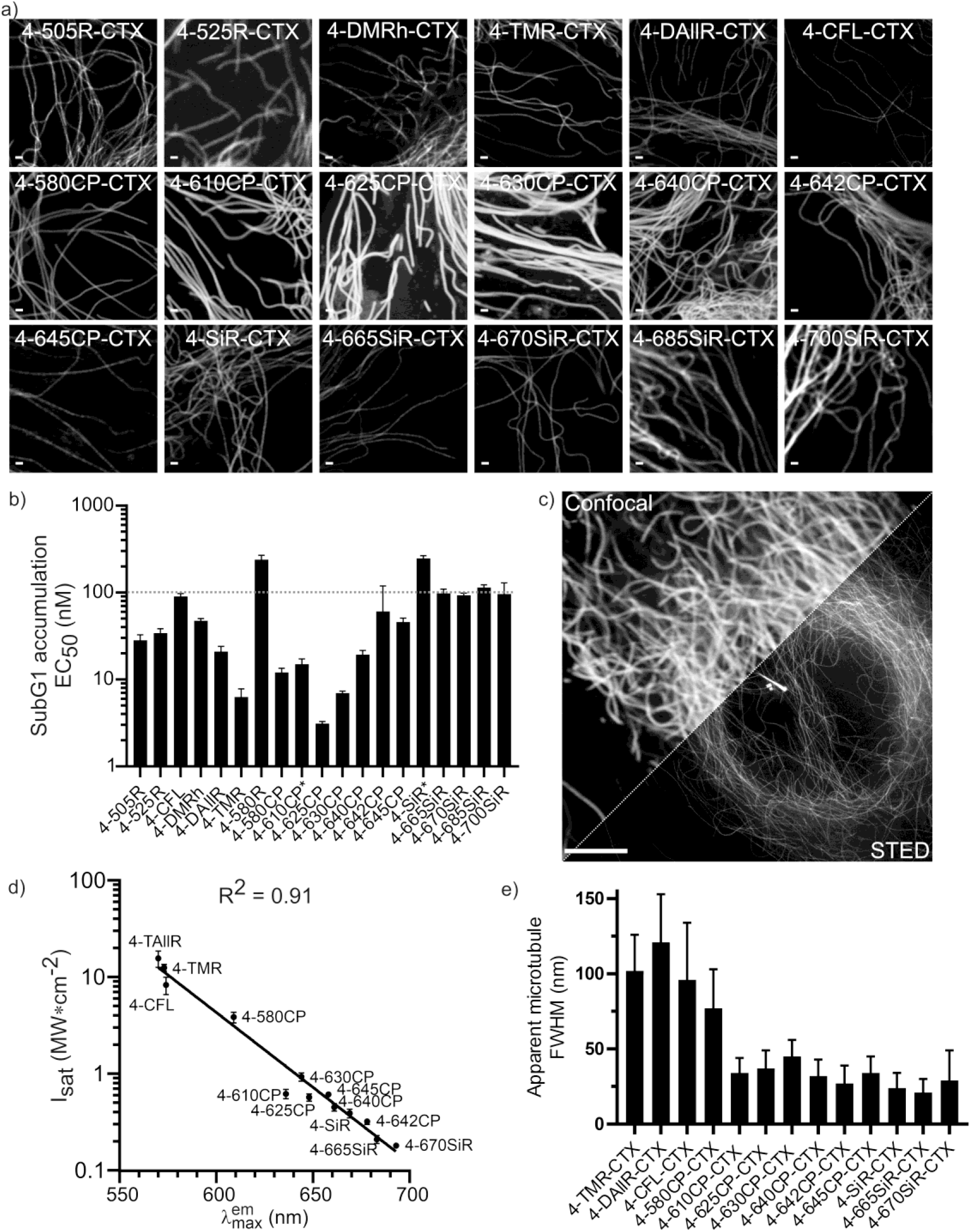
Properties of fluorescent tubulin probes. a) Confocal images of microtubules in living human fibroblasts under no-wash conditions. b) Toxicity of tubulin probes for Hela cells after 24 h incubation. * - EC_50_ of **4-610CP-CTX** and **4-SiR-CTX** was measured in ref. ^22^ c) Confocal and STED microscopy image of human fibroblasts stained with 10nM **4-625CP-CTX** probe. Note, visible in the center centrosome and primary cilium. Scale bar 5 μm. d) measured apparent microtubule FWHM in STED mode with 775 nm depletion laser, data represents mean ± SD (N = 30) e) Plot showing correlation of simulated emission efficiency (I_sat_) and emission maximum of the fluorophore

### Other fluorescent probes

A wide spectral range of synthesized fluorophores opens up the opportunities for multicolor imaging of many cell structures. We used the following targeting moieties: Hoechst – for DNA, jasplakinolide (JAS) for actin, triphenylphosphonium (TPP) for mitochondria and pepstatin A (PepA) for – lysosomes (Scheme 2).

#### Actin probes

Our and several other groups have been reporting fluorescent probes targeting actin which are produced by connecting jasplakinolide derivative^43,44^ via a long aliphatic linker to fluorescent dyes^45^. We hypothesized that increased permeability of 4-isomers would eliminate necessity of a long linker. Accordingly, we synthesized a series of probes by direct attachment of fluorescent dye to jasplakinolide derivative (Scheme 2). Indeed, no major differences of fluorescence signal intensity was observed in fibroblasts stained with a range of **4-610CP-JAS** and **4-610CP-C6-JAS** concentrations (Figure S21). Inspired by this result, we have synthesized actin probes spanning full visible spectrum: **4-505R-JAS**, **4-TMR-JAS**, **4-630CP-JAS** and **4-685SiR-JAS**. Confocal imaging of living fibroblasts confirmed good performance of staining with all these probes (Figure S22a). Periodical actin ring pattern was clearly visible in the STED microscopy images of cultured mouse primary neurons stained with **4-610CP-JAS** or **4-630CP-JAS** (Figure 4c and S22b).

**Figure 4.**
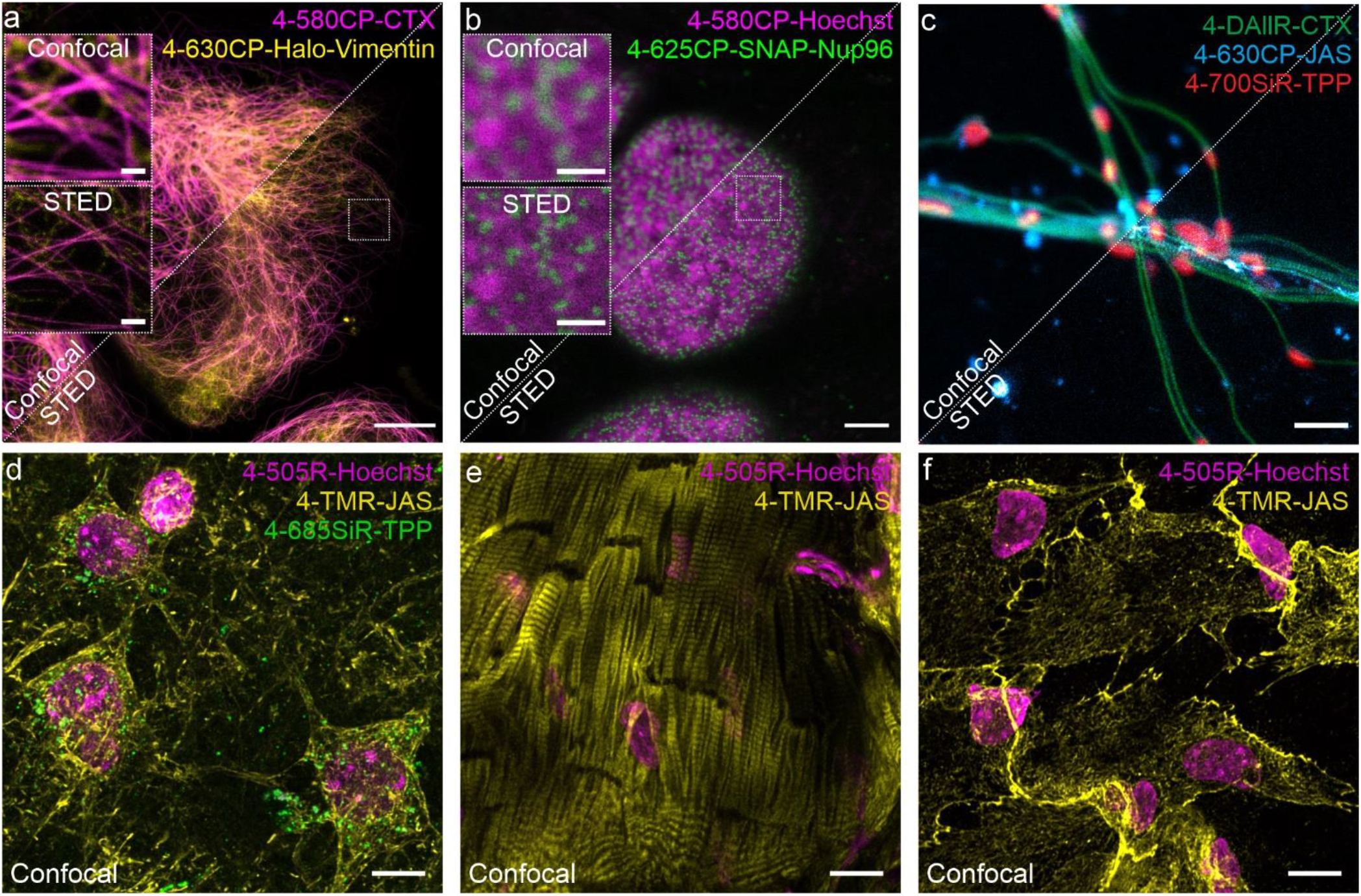
Confocal and STED images of the living cells and tissues stained with the cell membrane permeable probes. a. Confocal and STED images of U-2 OS cells expressing Vimentin-Halo stained with 100 nM **4-630CP-(O_4_)Halo** and 100 nM **4-580CP-CTX**. Images acquired on Abberior Expert Line microscope. Scale bars: large - 10 μm, inserts – 1 μm. b. Confocal and STED images of U-2 OS cells expressing Nup96-SNAP stained with 100 nM **4-625CP-BG** and 100 nM **4-580CP-Hoechst**. Images acquired on Abberior Expert Line microscope. Scale bars: large - 3 μm, inserts – 1 μm. c. Confocal and STED images of isolated mouse living neurons stained with 10 nM **4-DAllR-CTX**, 10 nM **4-630CP-JAS** and 10 nM **4-700SiR-TPP** probes. Images acquired on Abberior Facility Line microscope. Scale bar 1 μm. Confocal images of d. brain, e. heart, and f. liver tissue sections. Samples stained with 250 nM **4-505R-Hoechst** (nuclei-labelling probe, magenta) and 1 μM **4-TMR-JAS** (actin-labelling probe, yellow) and 1.5 μM **4-685SiR-TPP** (mitochondria-labelling probe, green). Images show max intensity projections of z-stacks. Scale bar: 10 μm. Images were deconvoluted with SVI Huygens software.

#### DNA probes

We used a previously reported design of DNA labelling probes using Hoechst linked via aliphatic C4-linker to the fluorescent dyes: **4-505R-**, **4-525R-**, **4-CFL-**, **4-625CP-**, **4-630CP**- and **4-685SiR-** (**87**-**92**). All probes stained DNA in the nucleus of living fibroblasts except **4-685SiR-Hoechst** which showed high background in the cytosol (Figure S23).

#### Mitochondria probes

Matrix of the mitochondria has a pH ~ 8, which is higher than the one of cytosol (7.0-7.4) and membrane potential permitting targeting by positively charged triphenylphosphonium (TPP)^46^. We have synthesized probes by linking TPP via C6-linker to the fluorescent dyes: **4-525R-**, **4-625CP-**, **4-640CP-**, **4-665SiR-**, **4-685SiR-** and **4-700SiR** (**80**-**86**). All probes stained mitochondria except **4-525R-TPP**, which demonstrated some off-targeting to endosomes and high background staining (Figure S24).

#### Lysosome probes

Contrary to mitochondria, lysosomes are acidic organelles (pH ~5) harboring multiple proteases^47^. We synthesized lysosomal fluorescent probes by coupling fluorophores (**4-525R-**, **4-580R-**, **4-630CP-**, **4-642CP-**, **4-SiR-**, **4-685SiR-** and **4-700SiR-**) via aliphatic linker to pepstatin A (**93**-**99**), a known inhibitor of cathepsin D protease. All studied probes featured selective lysosomal staining and the most intensive staining of lysosomal vesicles was obtained with **4-642CP-PepA**, **4-SiR-PepA** and **4-685SiR-PepA** (Figure S25).

In summary, most of the newly generated probes targeting different cell structures and self-labeling tags efficiently stained the living cells. High success rate (>80%) confirms the advantage of incorporation of isomer-4 rhodamines into fluorescent probe design.

### Multicolor imaging of live cells and tissues

Having in hand a set of fluorescent probes spanning the spectrum from 500 to 700 nm, we designed multicolor imaging experiments of living cells and tissue (Figure 4). First, we combined **4-625CP-BG** and **4-630CP-BG** with **4-580CP-Hoechst** for staining U-2 OS cells expressing Nup96-SNAP. The obtained STED images clearly show that the Nup96-SNAP signal localizes to dark areas in 4-580CP channel confirming that the nuclear pore complexes protrude into the heterochromatin layer creating chromatin exclusion zones (Figure 4b).

Small molecule fluorescent probes are a particularly powerful tool for imaging primary cells and tissues ^48^. To demonstrate this, we stained isolated mouse neurons using three probes targeting tubulin (**4-DAllR-CTX**), actin (**4-630CP-JAS**) and mitochondria (**4-700SiR-TPP**). The three color confocal and STED image clearly shows mitochondria (red) embedded into cytoskeleton of axon/dendrite (Figure 4c).

Another advantage of small molecule fluorescent probes is a good penetration of the tissues. We demonstrate this by staining living isolated mouse brain, heart and liver tissue sections with fluorescent probes targeting DNA, actin and mitochondria (Figure 4d-f).

Finally, the full palette of fluorescent dyes allowed us to design four color time-lapse live-cell experiments (Videos S1-S4 and Table S6). To this end, we stained human fibroblasts and HUVEC cells with **4-505R-Hoechst**, **4-DAllR-CTX**, **4-642CP-PepA** and **4-700SiR-TPP**. The obtained time-lapse movies show dynamics of DNA, tubulin, lysosomes and mitochondria with minimal bleaching (Video S1-S2). An alternative scheme employed **4-505R-CTX**, **4-CFL-Hoechst**, **4-624CP-PepA** and **4-700SiR-TPP** (Video S3). The third 4-color scheme consists of **4-505R-Hoechst**, **4-DAllR- CTX**, **4-625CP-(O_4_)Halo** and **4-700SiR-TPP** and was devised to observe the dynamics of mitochondria in the network of vimentin and tubulin in U-2 OS cells expressing Vimentin-Halo fusion (Video S4). In all cases, we took advantage of all four detectors on a commercial Abberior STED Facility line microscope and acquired data using 2 + 2 simultaneous acquisition scheme. This allowed relatively fast imaging (3s per frame) of large fields of view (50 x 50 μm) with minimal or no crosstalk between the channels.

In summary, we have devised a general, highly efficient, facile and scalable strategy to synthesize 4-carboxyrhodamine dyes without the need of protecting groups. We used the obtained dyes to generate a series of fluorescent probes targeting microtubules, actin, mitochondria, lysosomes, DNA, Halo and SNAP tagged proteins. The isomer-4 rhodamines’ probes, due to the NGE phenomenon, demonstrate a combination of excellent cell membrane permeability and specificity. The generality of such biocompatibility enhancement across a large structural variety of fluorescent probes allows fine-tuning to ideally meet the technical and biological requirements of confocal and STED imaging of living cells and tissues. A wide selection of probes operating at different wavelengths allows multicolor imaging of living specimen and tissue. We expect that the demonstrated high biocompatibility together with scalable facile synthesis of rhodamines’ 4-isomers will serve as a general tool-box for scientists to improve the performance of existing fluorescent probes and will accelerate the creation of numerous new.

## Author Contributions

JB and GL conceived and planned the study. JB, GK, RG, TK, KK, SP and GL performed the experiments. JB, KK, SP, GK, RG and GL performed the data analysis. JB and GL wrote the initial draft; all authors contributed to the final version of the manuscript.

## ACKNOWLEDGMENT

The authors are grateful to Dr. Vladimir Belov, Jan Seikowski, Jens Schimpfhauser and Jürgen Bienert for NMR measurements of numerous samples and providing Boc-JAS actin targeting moiety. They also acknowledge Dr. Peter Lenart and Dr. Antonio Politi (Live-cell imaging facility) for the possibility to perform live-cell spinning disk confocal microscopy. JB is grateful to the Max Planck Society for a Nobel Laureate Fellowship. GK is grateful to EMBO for Long-Term Fellowship (ALTF 135-2019). S.P. was supported by the Ph.D. program “Genome Science” - International Max Planck Research School. The authors acknowledge Jaydev Jethwa for critical reading of the manuscript.

## REFERENCES

1 Grimm, J. B. & Lavis, L. D. Caveat fluorophore: an insiders’ guide to small-molecule fluorescent labels. Nat Methods 19, 149–158, doi:10.1038/s41592-021-01338-6 (2022).

2 Hell, S. W. Nanoscopy with Focused Light (Nobel Lecture). Angew Chem Int Ed Engl 54, 8054–8066, doi:10.1002/anie.201504181 (2015).

3 Sahl, S. J., Hell, S. W. & Jakobs, S. Fluorescence nanoscopy in cell biology. Nat Rev Mol Cell Biol 18, 685–701, doi:10.1038/nrm.2017.71 (2017).

4 ßutkevich, A. N., Lukinavičius, G., D’Este, E. & Hell, S. W. Cell-Permeant Large Stokes Shift Dyes for Transfection-Free Multicolor Nanoscopy. J Am Chem Soc 139, 12378–12381, doi:10.1021/jacs.7b06412 (2017).

5 Butkevich, A. N. et al. Hydroxylated Fluorescent Dyes for Live-Cell Labeling: Synthesis, Spectra and Super-Resolution STED. Chemistry 23, 12114–12119, doi:10.1002/chem.201701216 (2017).

6 Grimm, J. B. et al. A general method to fine-tune fluorophores for live-cell and in vivo imaging. Nat Methods 14, 987–994, doi:10.1038/nmeth.4403 (2017).

7 Wang, L. et al. A general strategy to develop cell permeable and fluorogenic probes for multicolour nanoscopy. Nat Chem 12, 165–172, doi:10.1038/s41557-019-0371-1 (2020).

8 Lukinavičius, G. et al. A near-infrared fluorophore for live-cell super-resolution microscopy of cellular proteins. Nat Chem 5, 132–139, doi:10.1038/nchem.1546 (2013).

9 Wang, L., Frei, M. S., Salim, A. & Johnsson, K. Small-Molecule Fluorescent Probes for Live-Cell Super-Resolution Microscopy. J Am Chem Soc 141, 2770–2781, doi:10.1021/jacs.8b11134 (2019).

10 ßucevičius, J., Keller-Findeisen, J., Gilat, T., Hell, S. W. & Lukinavičius, G. Rhodamine-Hoechst positional isomers for highly efficient staining of heterochromatin. Chem Sci 10, 1962–1970, doi:10.1039/c8sc05082a (2019).

11 Lukinavičius, G. et al. Fluorogenic Probes for Multicolor Imaging in Living Cells. J Am Chem Soc 138, 9365–9368, doi:10.1021/jacs.6b04782 (2016).

12 Zheng, Q. et al. Rational Design of Fluorogenic and Spontaneously Blinking Labels for Super-Resolution Imaging. ACS Cent Sci 5, 1602–1613, doi:10.1021/acscentsci.9b00676 (2019).

13 Beija, M., Afonso, C. A. & Martinho, J. M. Synthesis and applications of Rhodamine derivatives as fluorescent probes. Chem Soc Rev 38, 2410–2433, doi:10.1039/b901612k (2009).

14 Chen, X., Pradhan, T., Wang, F., Kim, J. S. & Yoon, J. Fluorescent chemosensors based on spiroring-opening of xanthenes and related derivatives. Chem Rev 112, 1910–1956, doi:10.1021/cr200201z (2012).

15 Zheng, H., Zhan, X. Q., Bian, Q. N. & Zhang, X. J. Advances in modifying fluorescein and rhodamine fluorophores as fluorescent chemosensors. Chem Commun (Camb) 49, 429–447, doi:10.1039/c2cc35997a (2013).

16 Lukinavičius, G. et al. Fluorescent dyes and probes for super-resolution microscopy of microtubules and tracheoles in living cells and tissues. Chem Sci 9, 3324–3334, doi:10.1039/c7sc05334g (2018).

17 Grimm, F., Nizamov, S. & Belov, V. N. Green-Emitting Rhodamine Dyes for Vital Labeling of Cell Organelles Using STED Super-Resolution Microscopy. Chembiochem 20, 2248–2254, doi:10.1002/cbic.201900177 (2019).

18 Lardon, N. et al. Systematic Tuning of Rhodamine Spirocyclization for Super-resolution Microscopy. J Am Chem Soc 143, 14592–14600, doi:10.1021/jacs.1c05004 (2021).

19 Grimm, J. B. et al. A general method to optimize and functionalize red-shifted rhodamine dyes. Nat Methods 17, 815–821, doi:10.1038/s41592-020-0909-6 (2020).

20 Grimm, J. B. et al. A general method to improve fluorophores for live-cell and single-molecule microscopy. Nat Methods 12, 244–250, doi:10.1038/nmeth.3256 (2015).

21 Gerasimaitė, R. et al. Efflux pump insensitive rhodamine-jasplakinolide conjugates for G-and Factin imaging in living cells. Org Biomol Chem 18, 2929–2937, doi:10.1039/d0ob00369g (2020).

22 Bucevičius, J., Kostiuk, G., Gerasimaitė, R., Gilat, T. & Lukinavičius, G. Enhancing the biocompatibility of rhodamine fluorescent probes by a neighbouring group effect. Chem Sci 11, 7313–7323, doi:10.1039/d0sc02154g (2020).

23 Butkevich, A. N. Modular Synthetic Approach to Silicon-Rhodamine Homologues and Analogues via Bis-aryllanthanum Reagents. Org Lett 23, 2604–2609, doi:10.1021/acs.orglett.1c00512 (2021).

24 Gannon, M. K., 2nd & Detty, M. R. Generation of 3-and 5-lithiothiophene-2-carboxylates via metal-halogen exchange and their addition reactions to chalcogenoxanthones. J Org Chem 72, 2647–2650, doi:10.1021/jo062370x (2007).

25 Gonschorek, W. & Kuppers, H. The crystal structure of lithium hydrogen phthalate dihydrate, containing a very short hydrogen bond. Acta Crystallographica Section B 31, 1068–1072, doi:doi:10.1107/S0567740875004517 (1975).

26 Bachman, J. L., Escamilla, P. R., Boley, A. J., Pavlich, C. I. & Anslyn, E. V. Improved Xanthone Synthesis, Stepwise Chemical Redox Cycling. Org Lett 21, 206–209, doi:10.1021/acs.orglett.8b03661 (2019).

27 Pastierik, T., Sebej, P., Medalova, J., Stacko, P. & Klan, P. Near-infrared fluorescent 9-phenylethynylpyronin analogues for bioimaging. J Org Chem 79, 3374–3382, doi:10.1021/jo500140y (2014).

28 Hanaoka, K. et al. Synthesis of unsymmetrical Si-rhodamine fluorophores and application to a far-red to near-infrared fluorescence probe for hypoxia. Chem Commun (Camb) 54, 6939–6942, doi:10.1039/c8cc02451k (2018).

29 Åkerlöf, G. & Short, O. A. The Dielectric Constant of Dioxane—Water Mixtures between 0 and 80°. Journal of the American Chemical Society 58, 1241–1243, doi:10.1021/ja01298a044 (1936).

30 Gaussian 09 Rev. D.01 (Wallingford, CT, 2009).

31 Tomasi, J., Mennucci, B. & Cammi, R. Quantum Mechanical Continuum Solvation Models. Chemical Reviews 105, 2999–3094, doi:10.1021/cr9904009 (2005).

32 Adamo, C. & Jacquemin, D. The calculations of excited-state properties with Time-Dependent Density Functional Theory. Chem Soc Rev 42, 845–856, doi:10.1039/c2cs35394f (2013).

33 Zhou, P. Why the lowest electronic excitations of rhodamines are overestimated by time-dependent density functional theory. International Journal of Quantum Chemistry 118, e25780, doi:https://doi.org/10.1002/qua.25780 (2018).

34 Keppler, A., Arrivoli, C., Sironi, L. & Ellenberg, J. Fluorophores for live cell imaging of AGT fusion proteins across the visible spectrum. Biotechniques 41, 167–170, 172, 174-165, doi:10.2144/000112216 (2006).

35 Keppler, A. et al. A general method for the covalent labeling of fusion proteins with small molecules in vivo. Nat Biotechnol 21, 86–89, doi:10.1038/nbt765 (2003).

36 Wilhelm, J. et al. Kinetic and Structural Characterization of the Self-Labeling Protein Tags HaloTag7, SNAP-tag, and CLIP-tag. Biochemistry 60, 2560–2575, doi:10.1021/acs.biochem.1c00258 (2021).

37 Los, G. V. et al. HaloTag: a novel protein labeling technology for cell imaging and protein analysis. ACS Chem Biol 3, 373–382, doi:10.1021/cb800025k (2008).

38 Thevathasan, J. V. et al. Nuclear pores as versatile reference standards for quantitative superresolution microscopy. Nat Methods 16, 1045–1053, doi:10.1038/s41592-019-0574-9 (2019).

39 Butkevich, A. N. et al. Two-Color 810 nm STED Nanoscopy of Living Cells with Endogenous SNAP-Tagged Fusion Proteins. ACS Chem Biol 13, 475–480, doi:10.1021/acschembio.7b00616 (2018).

40 Ratz, M., Testa, I., Hell, S. W. & Jakobs, S. CRISPR/Cas9-mediated endogenous protein tagging for RESOLFT super-resolution microscopy of living human cells. Sci Rep 5, 9592, doi:10.1038/srep09592 (2015).

41 Vicidomini, G., Moneron, G., Eggeling, C., Rittweger, E. & Hell, S. W. STED with wavelengths closer to the emission maximum. Opt Express 20, 5225–5236, doi:10.1364/OE.20.005225 (2012).

42 Gerasimaitė, R. T. et al. Blinking Fluorescent Probes for Tubulin Nanoscopy in Living and Fixed Cells. ACS Chem Biol 16, 2130–2136, doi:10.1021/acschembio.1c00538 (2021).

43 Milroy, L. G. et al. Selective chemical imaging of static actin in live cells. J Am Chem Soc 134, 8480–8486, doi:10.1021/ja211708z (2012).

44 Tannert, R. et al. Synthesis and structure-activity correlation of natural-product inspired cyclodepsipeptides stabilizing F-actin. J Am Chem Soc 132, 3063–3077, doi:10.1021/ja9095126 (2010).

45 Lukinavičius, G. et al. Fluorogenic probes for live-cell imaging of the cytoskeleton. Nat Methods 11, 731–733, doi:10.1038/nmeth.2972 (2014).

46 Zielonka, J. et al. Mitochondria-Targeted Triphenylphosphonium-Based Compounds: Syntheses, Mechanisms of Action, and Therapeutic and Diagnostic Applications. Chem Rev 117, 10043–10120, doi:10.1021/acs.chemrev.7b00042 (2017).

47 Boya, P. & Kroemer, G. Lysosomal membrane permeabilization in cell death. Oncogene 27, 6434–6451, doi:10.1038/onc.2008.310 (2008).

48 Guo, Z., Park, S., Yoon, J. & Shin, I. Recent progress in the development of near-infrared fluorescent probes for bioimaging applications. Chem Soc Rev 43, 16–29, doi:10.1039/c3cs60271k (2014).

