## Supplementary information for "A general, highly efficient and facile synthesis of biocompatible rhodamine dyes and probes for live-cell multicolor nanoscopy"

### Supplementary Schemes

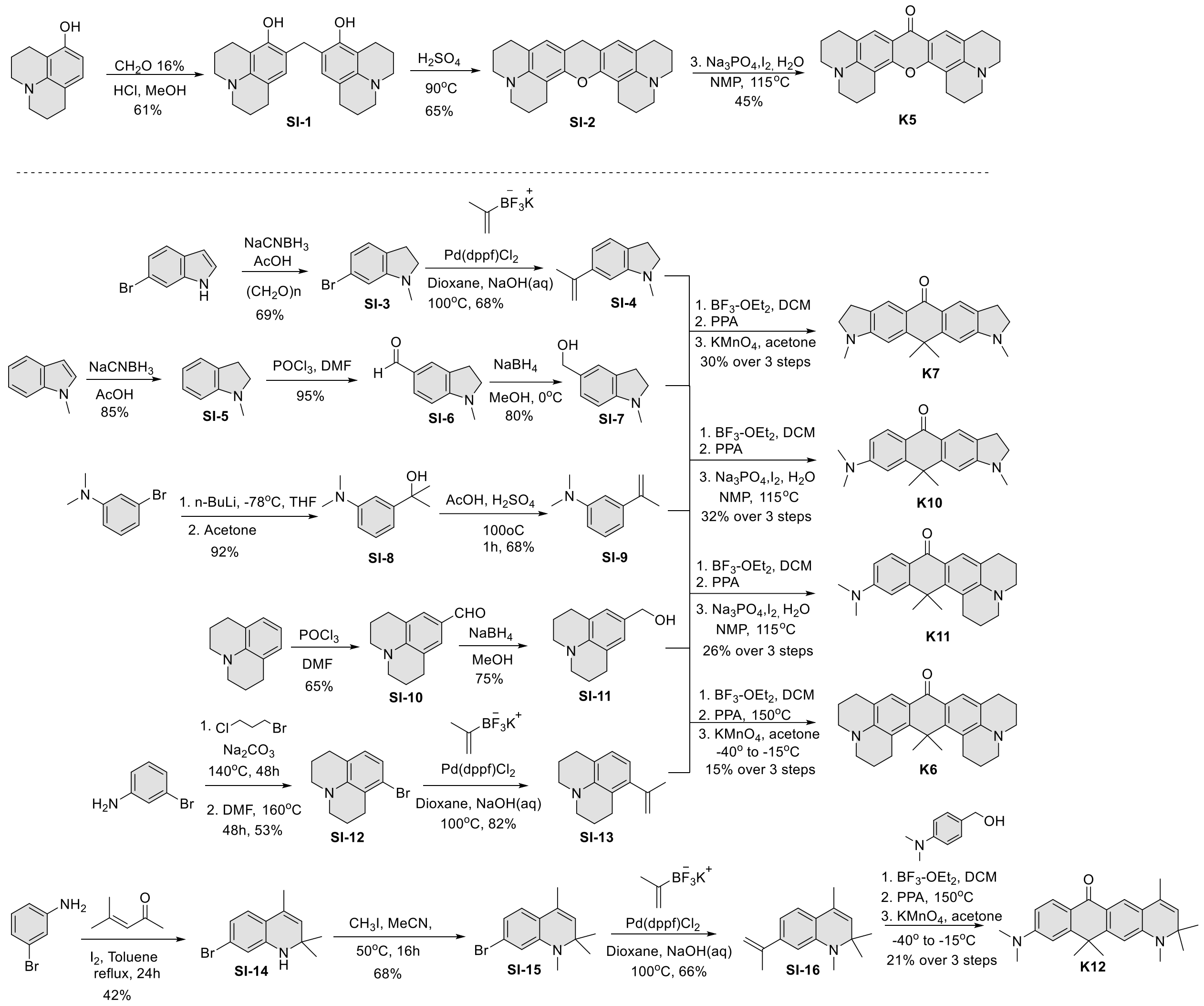

**Supplementary Scheme S1.** Synthetic routes to symmetrical (**K5**-**K7**) and unsymmetrical (**K10**-**K12**) xanthones with carbon bridging atom.

a) **Symmetrical**

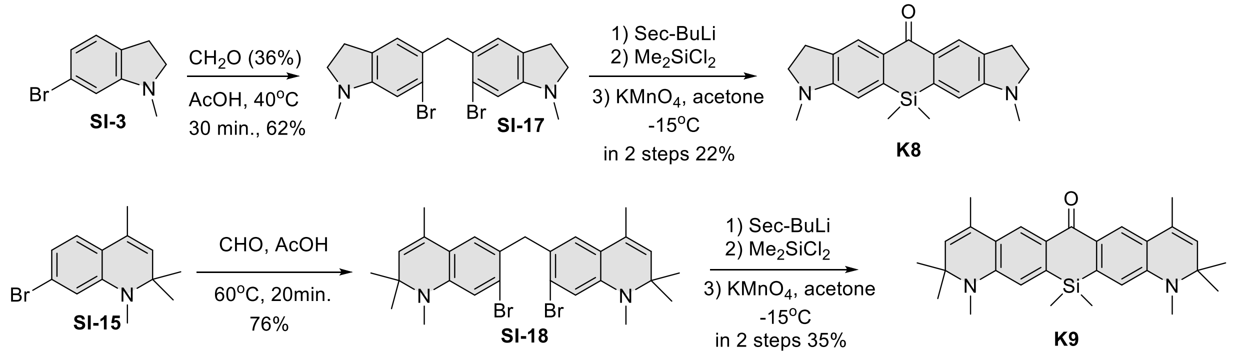

b) **Unsymmetrical**

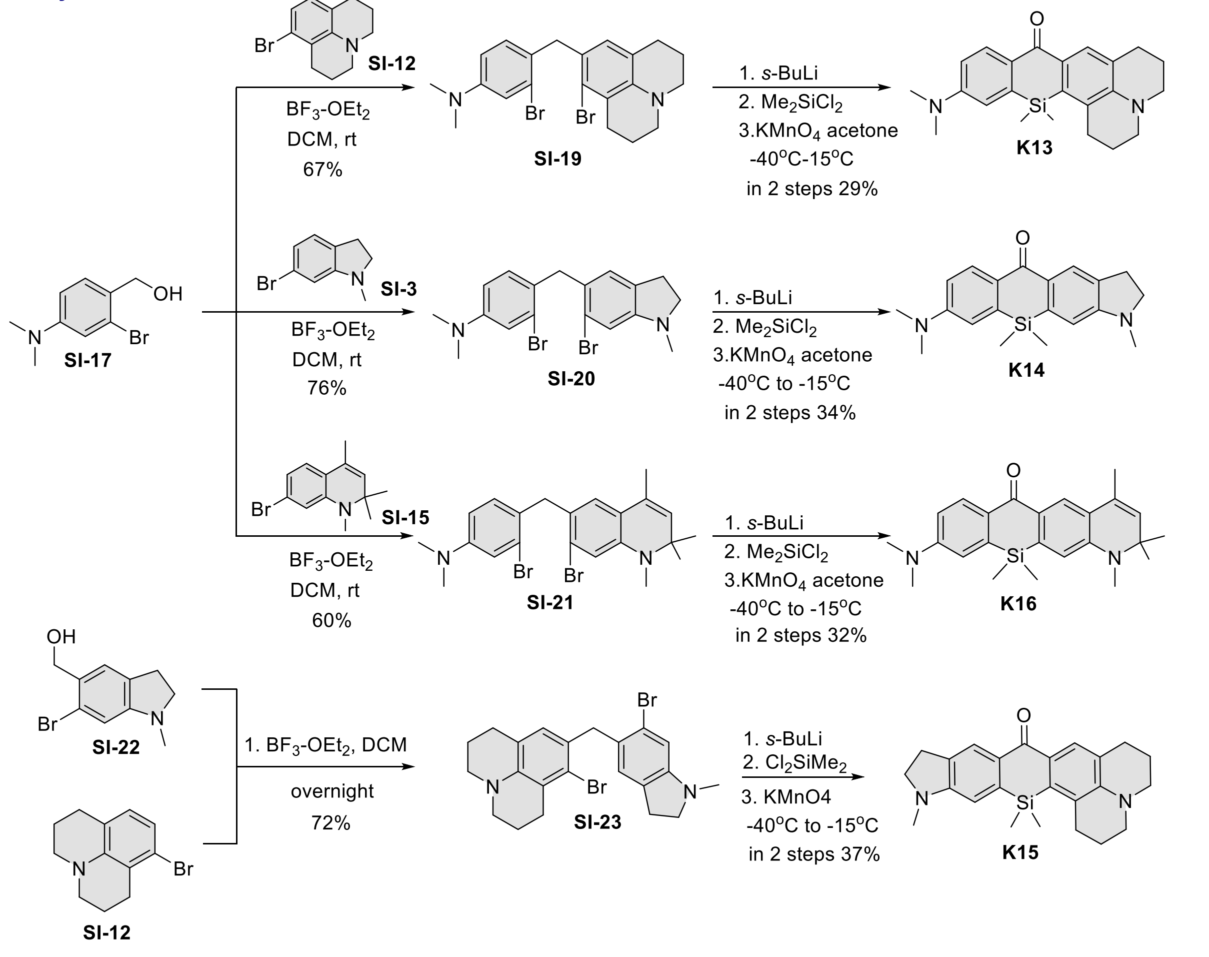

**Supplementary Scheme S2.** Synthetic routes to symmetrical (**K8-K9**) and unsymmetrical (**K13-K16**) xanthones with silicon bridging atom.

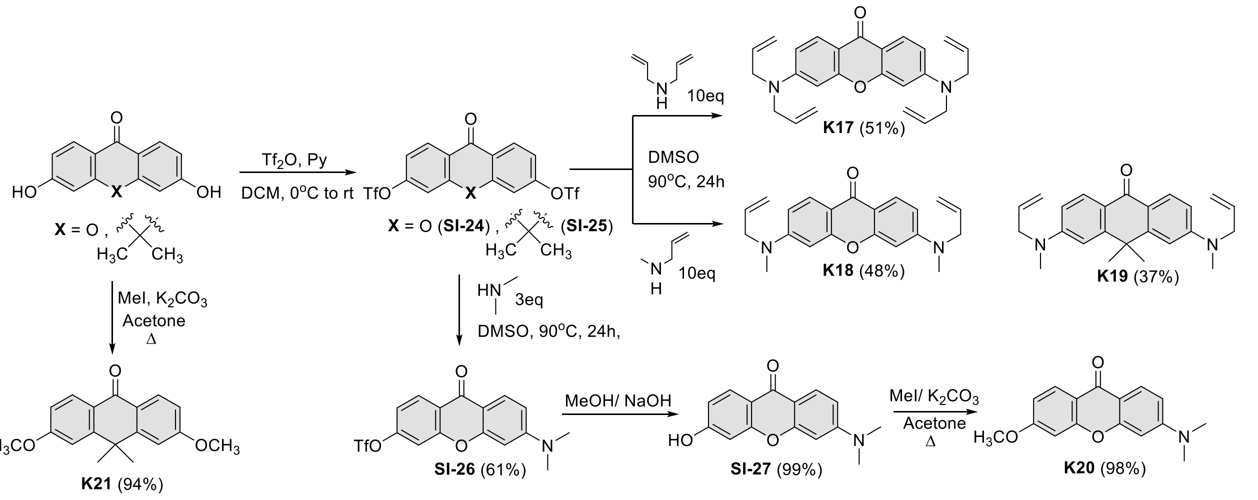

**Supplementary Scheme S3.** Synthetic routes to ketones **K17**-**K21**.

### Supplementary Figures

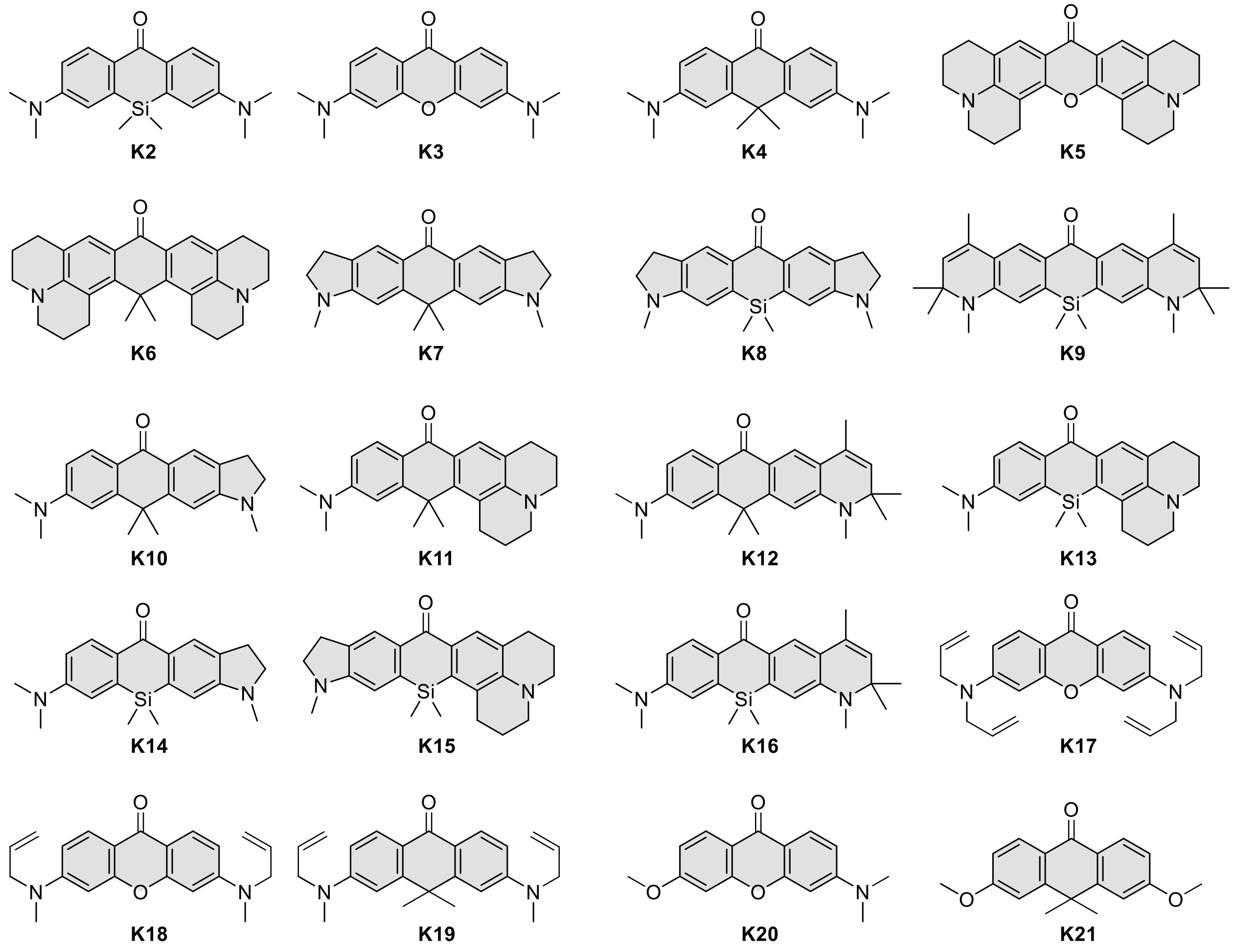

**Supplementary Figure S1.** Structures of ketones used in the reaction scope studies.

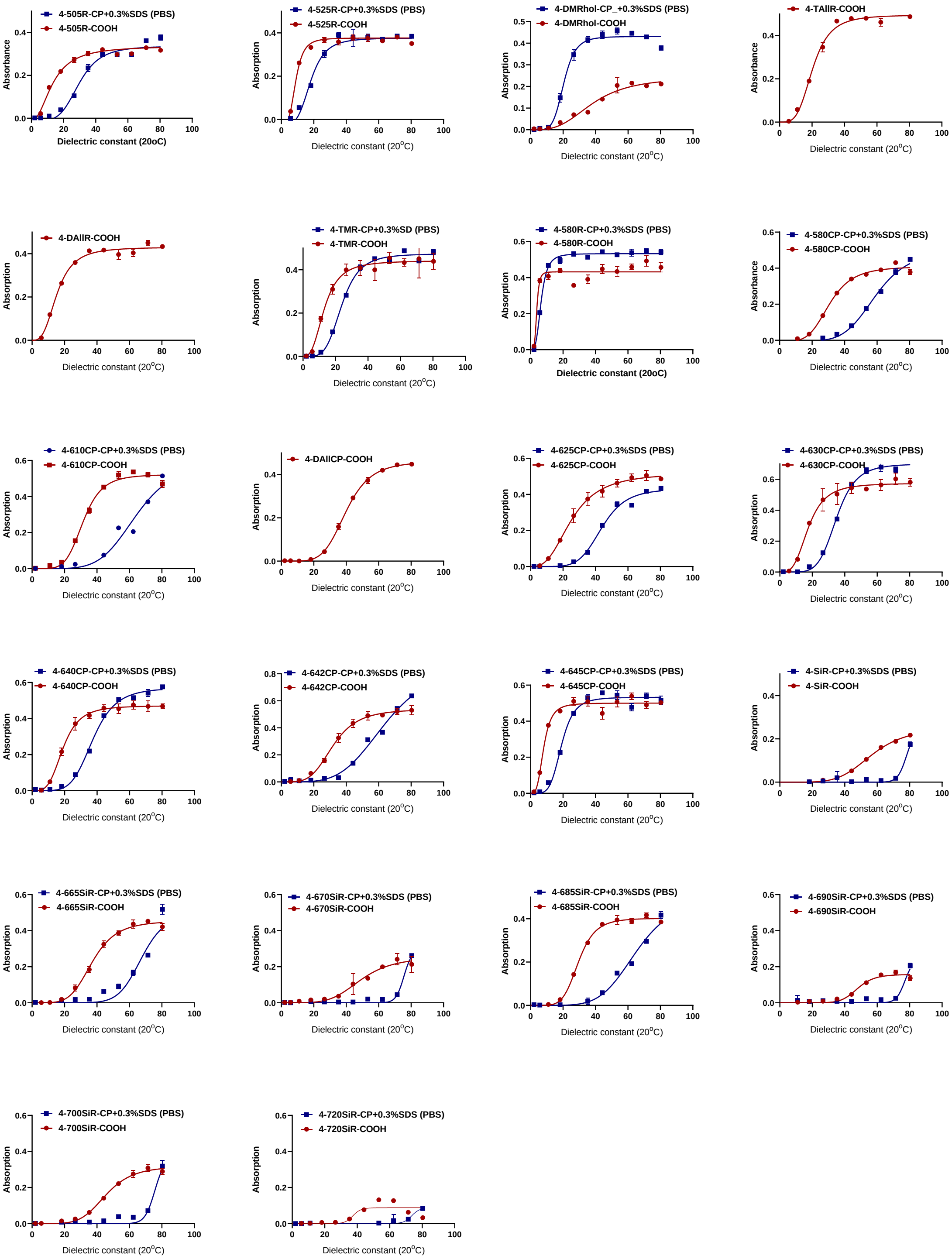

**Supplementary Figure S2.** Absorption of the free dyes and respective 4-(benzyloxy)-2-chloropyrimidine (CP) derivatives in 1, 4-dioxane-water mixtures. Plots show absorbance of free dye (red) or CP derivative (blue) at $\lambda_{max}$ versus dielectric constant of 1,4-dioxane-water mixtures. $D_{50}$ values, obtained by fitting the data to $EC_{50}$ dose-response equation, are listed in supplementary table S2. Data points are presented as mean ± s.d. of three independently repeated experiments (N=3). When titrating the probes, SDS was included to suppress their aggregation.

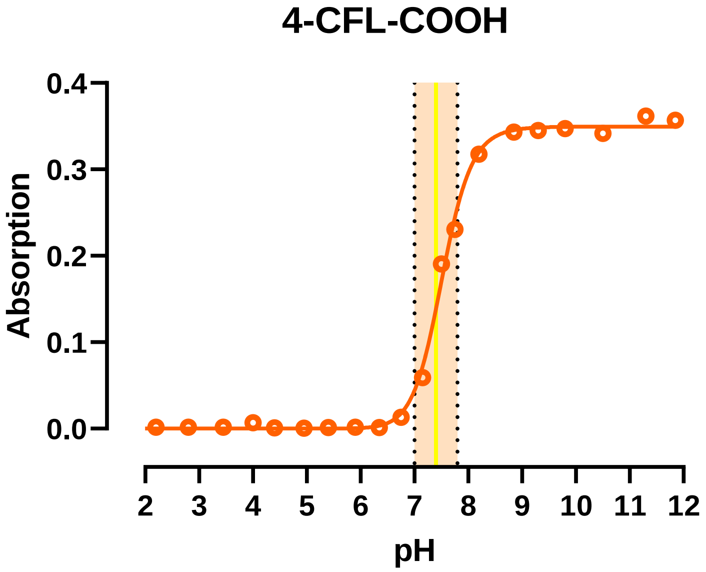

**Supplementary Figure S3**. Absorbance at $\lambda_{max}$ versus pH for **4-CFL-COOH**. Dashed lines mark interval pH 7.0-7.8 and yellow line indicates pH = 7.4. Measurements were performed in universal buffer for UV spectrophotometry which was prepared by taking 50mL solution consisting of 0.1 M citric acid (21.01 g/l), 0.1M $KH_2PO_4$ (13.61 g/l), 0.1 M sodium tetraborate (19.07 g/l), 0.1 M Tris (12.11 g/l), 0.1 M KCl (7.46 g/l) and adjusting the pH to the required values by adding x mL of 0.4 M HCl or 0.4 M NaOH, followed by the dilution to 200 ml. Buffers in the range of pH 2-12 were prepared in ~0.5 increments.

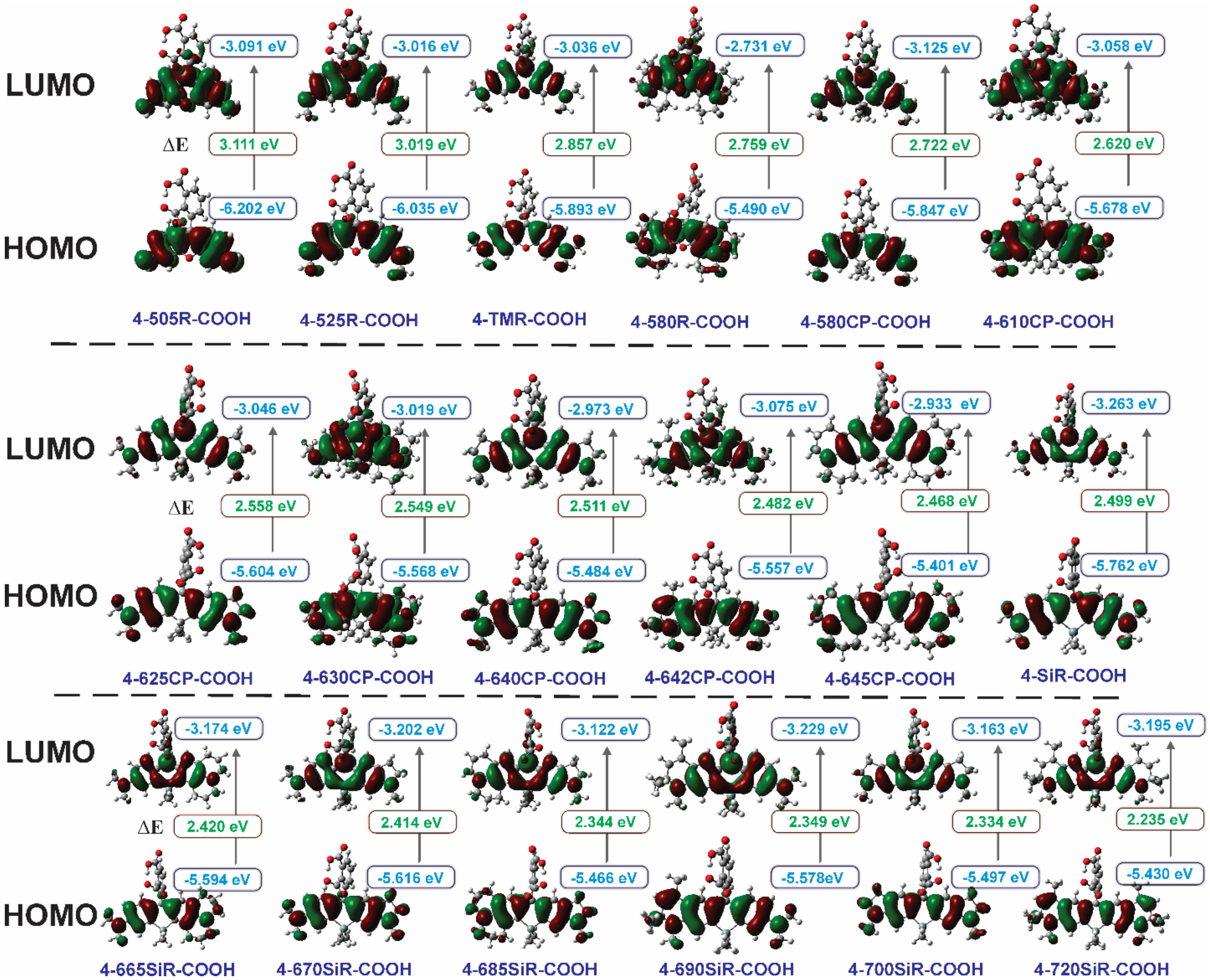

**Supplementary Figure S4.** Isodensity surface plots of frontier molecular orbitals of the synthesized 4-carboxyrhodamine dyes obtained by DFT calculations at B3LYP/6-311++G(d,p)/IEFPCM ($H_2O$) level of theory.

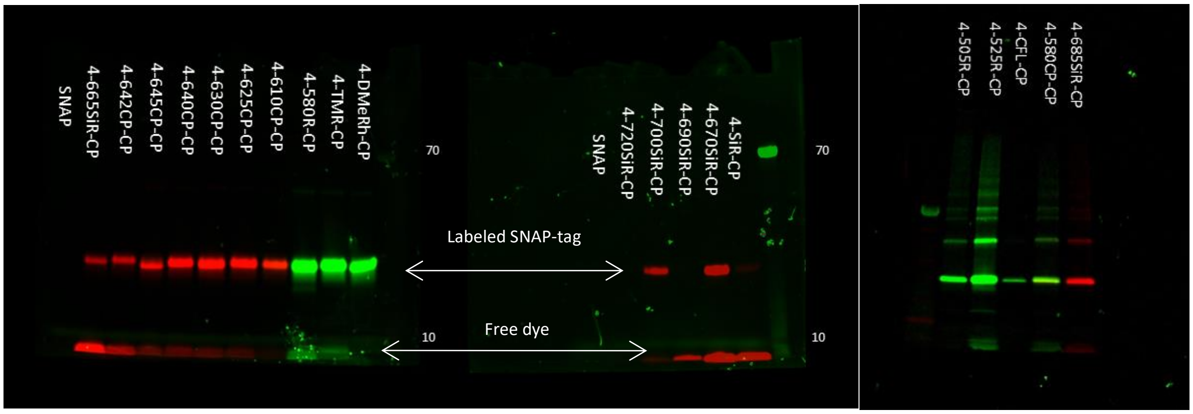

(Fluorescence)

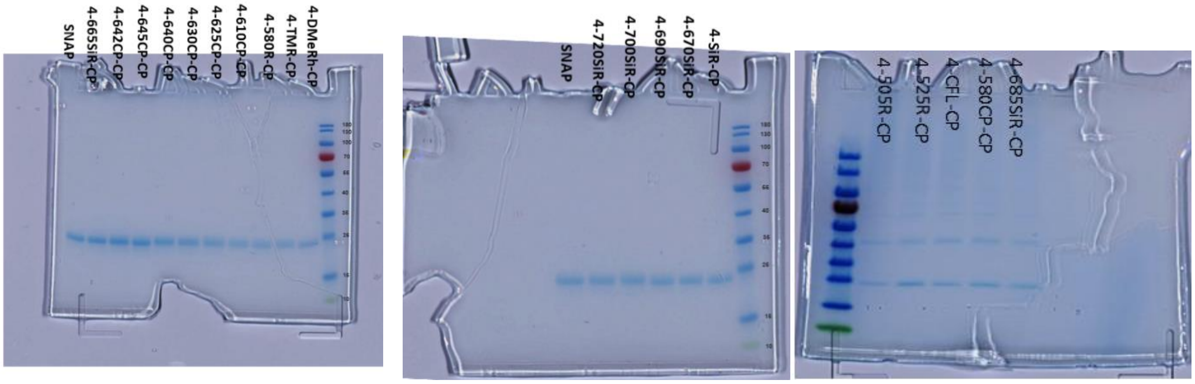

(Coomassie stain)

**Supplementary Figure S5.** SDS-PAGE gel images of SNAP tag protein labeled with 4-(benzyloxy)-2-chloropyrimidine (CP) derivatives **27**-**46**. Incubated for 3h at 37$^{o}$C.

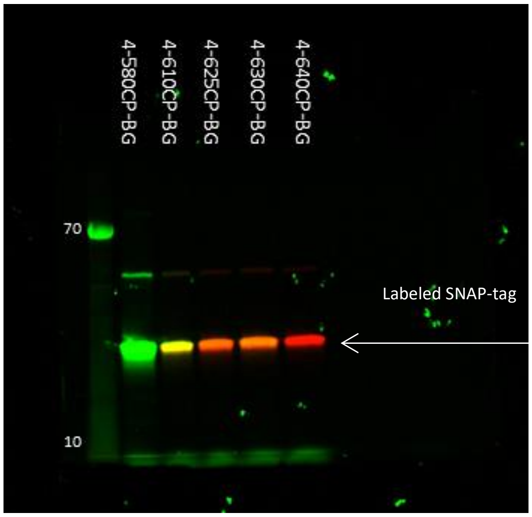

(Fluorescence)

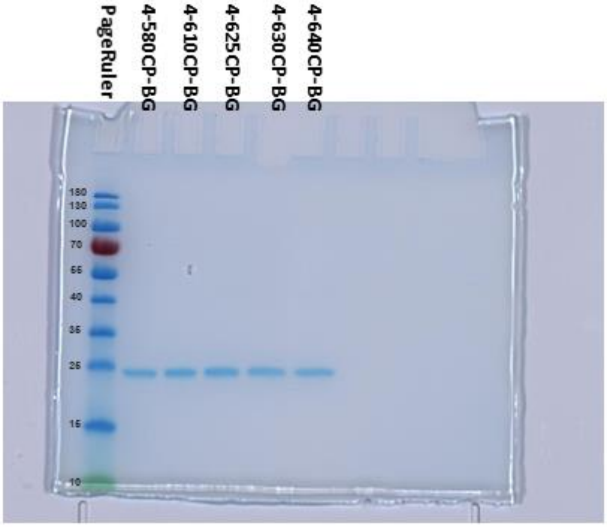

(Coomassie stain)

**Supplementary Figure S6.** SDS-PAGE gel images of SNAP tag protein labeled with O6-benzylguanine based (BG) SNAP-tag substrates **105**-**109**. Incubated for 3h at 37$^{o}$C.

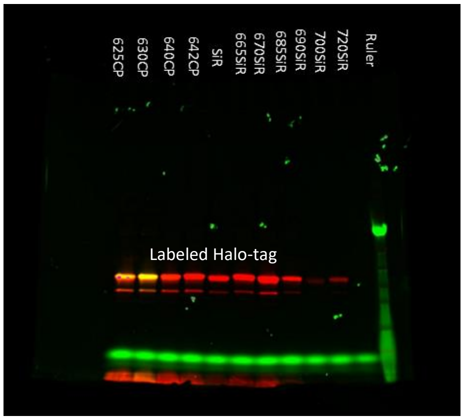

(Fluorescence

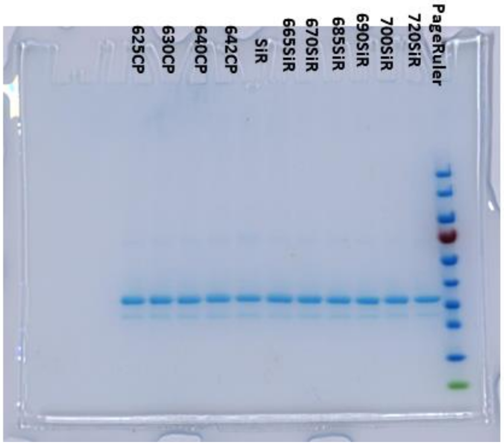

(Coomassie stain)

**Supplementary Figure S7.** SDS-PAGE gel images of Halo-tag protein labeled with Halo-tag substrates **68**-**79** with a SNAP tag protein. Incubated for 3h at 37$^{o}$C.

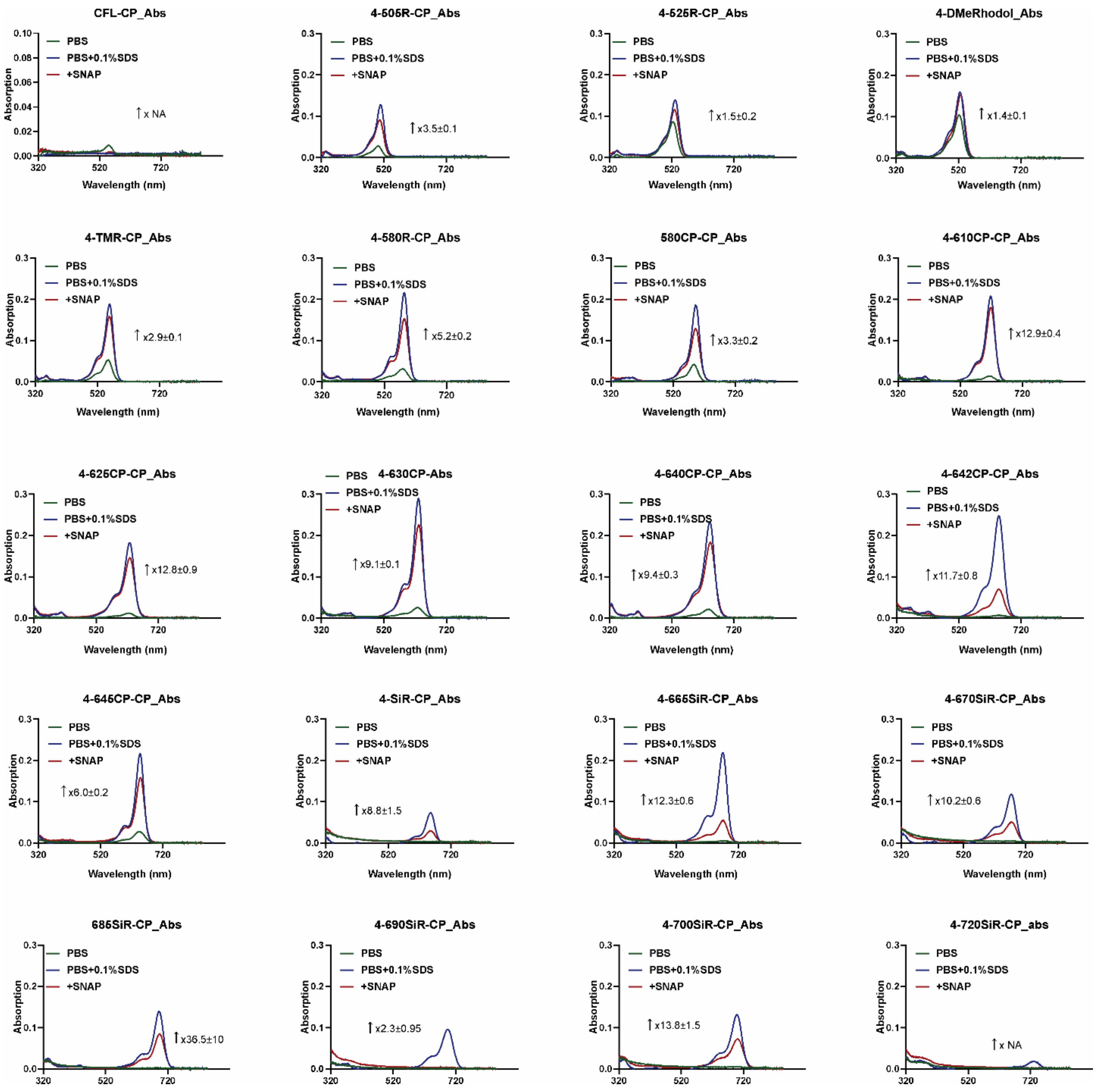

**Supplementary Figure S8**. Absorbance spectra of the chloropyrimidine (CP) based SNAP-tag substrates. Spectra were recorded after incubating 2.5 µM probes with 5 µM SNAP-tag protein (red), 0.1% SDS (blue) or without additives in PBS (green) at 37 °C for 3 h to ensure complete reaction. Spectra are presented as averages of three independently repeated experiments (N=3).

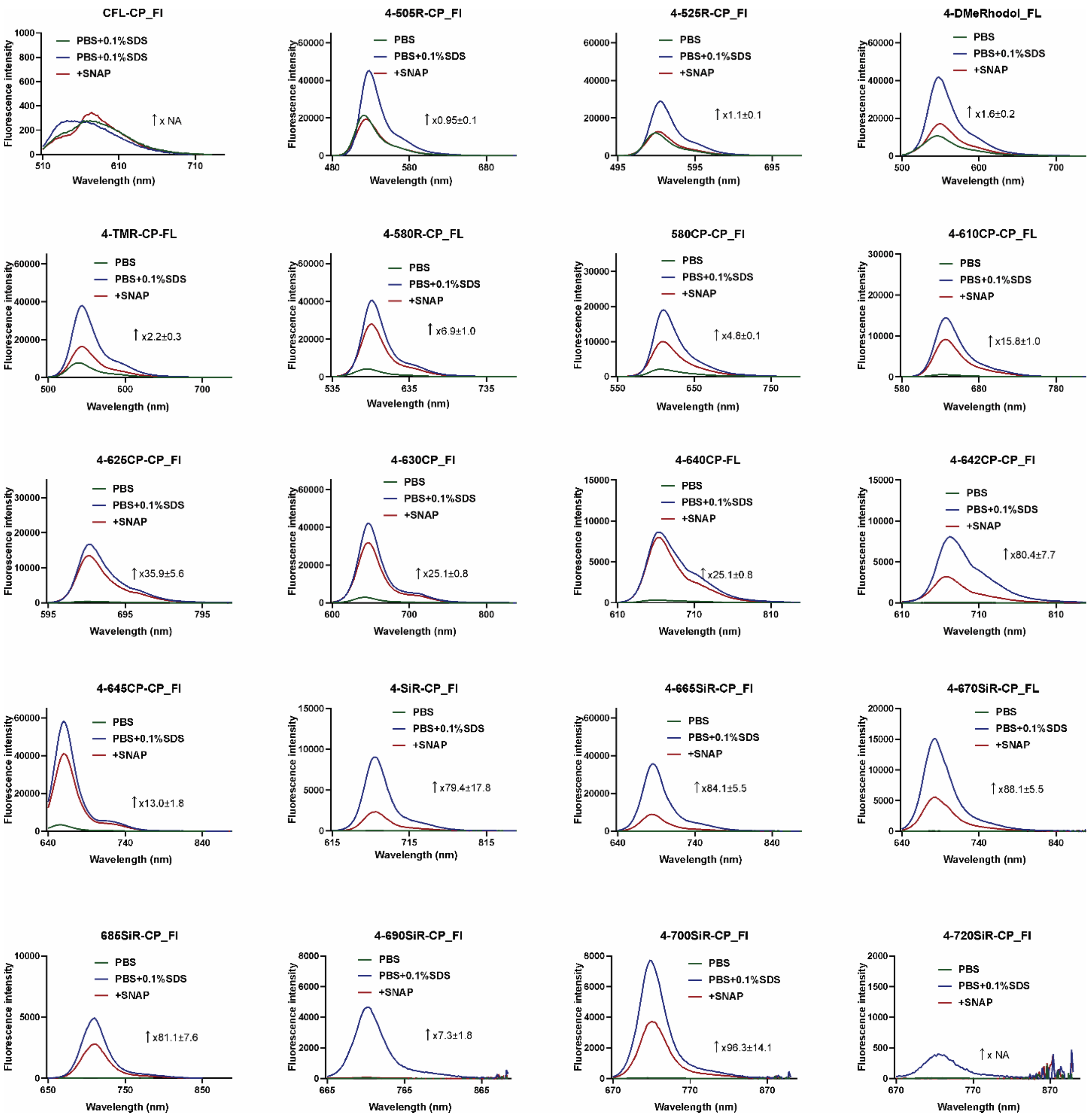

**Supplementary Figure S9**. Fluorescence spectra of the chloropyrimidine (CP) based SNAP-tag substrates. Spectra were recorded after incubating 2.5 µM probes with 5 µM SNAP-tag protein (red), 0.1% SDS (blue) or without additives in PBS (green) at 37 °C for 3 h to ensure complete reaction. Spectra are presented as averages of three independently repeated experiments (N=3).

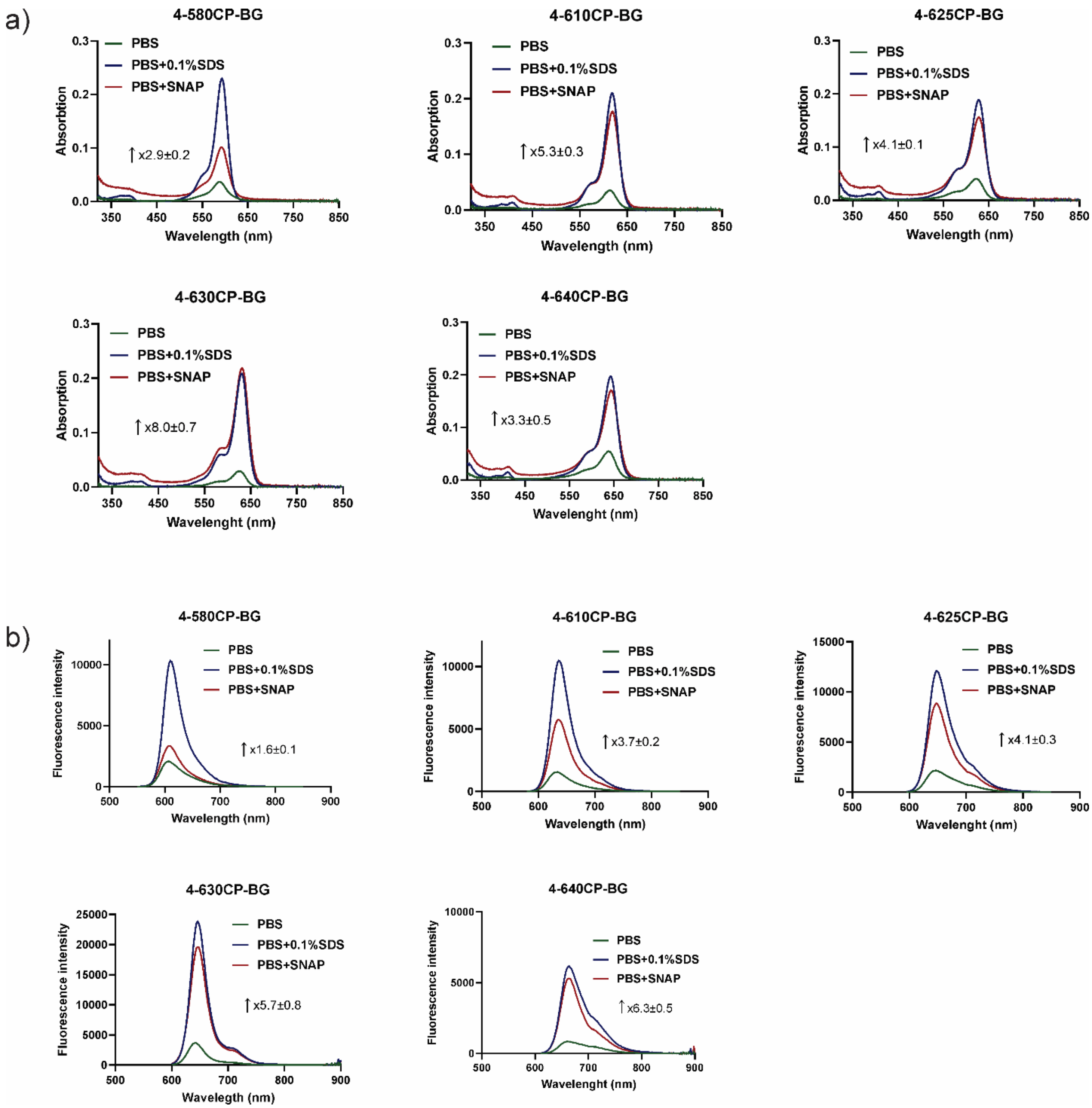

**Supplementary Figure S10**. Absorbance and fluorescence spectra of the O6-benzylguanine based (BG) SNAP-tag substrates. Absorbance (a) and fluorescence (b) spectra were recorded after incubating 2.5 µM probes with 5 µM Halo-tag protein (red), 0.1% SDS (blue) or without additives in PBS (green) at 37 °C for 3 h to ensure complete reaction. Spectra are presented as averages of three independently repeated experiments (N=3).

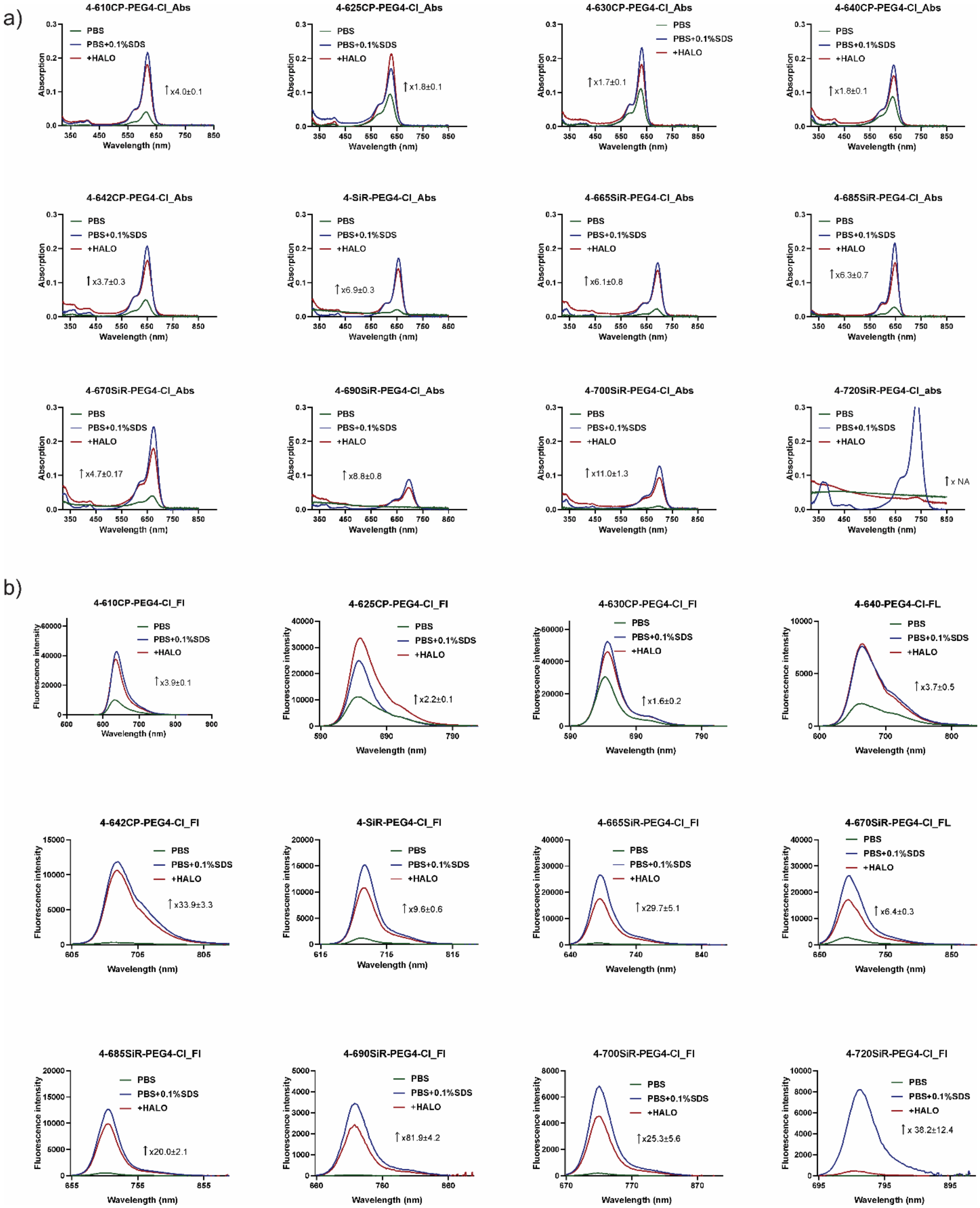

**Supplementary Figure S11**. Absorbance and fluorescence spectra of the Halo-tag substrates. Absorbance (a) and fluorescence (b) spectra were recorded after incubating 2.5 µM probes with 5 µM Halo-tag protein (red), 0.1% SDS (blue) or without additives in PBS (green) at 37 °C for 3 h to ensure complete reaction. Spectra are presented as averages of three independently repeated experiments (N=3).

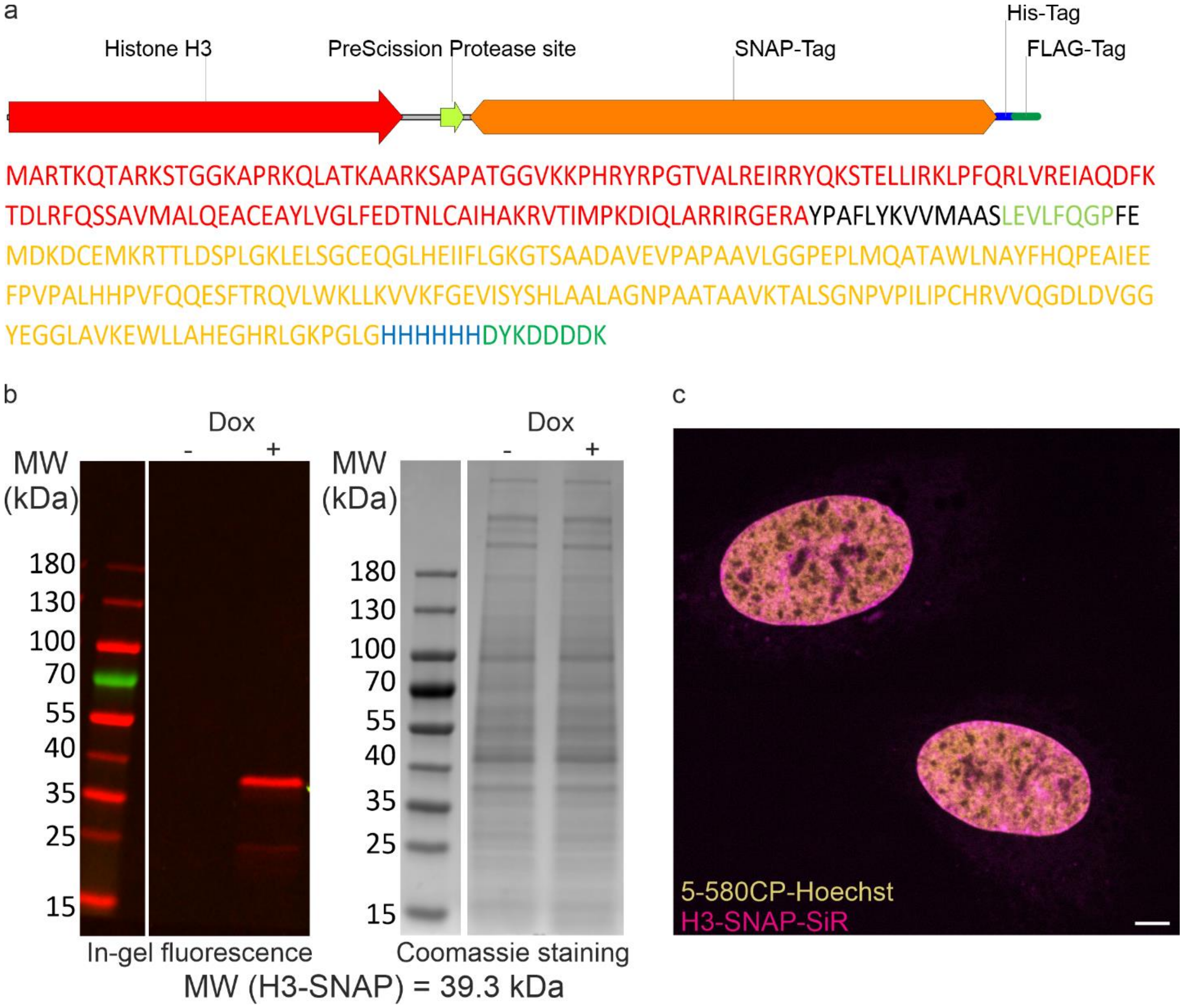

**Supplementary Figure S12**. Properties U-2 OS cell line expressing SNAP-tagged Histone H3. a. amino acid sequence of H3-SNAP protein. b. Inducible expression of H3-SNAP protein in U-2 OS cells. SNAP-tag labelled with 1 µM SiR-SNAP (NEB) substrate for 1h at 37°C in CelLytic™ M (Sigma-Aldrich) cell lysis reagent solution. The obtained protein sample supplemented with sample loading buffer (Bio-rad) and SDS-PAGE was performed. Protein expression induced with 0.1 µg/ml doxycycline for 24h before the lysis. c. Co-localization of H3-SNAP protein and nuclear DNA stain 5-580CP-Hoechst. Cells were stained with 100 nM SiR-SNAP (NEB) and 100 nM 5-580CP-Hoecshst in DMEM growth medium containing 10% FBS at 37 °C for 1 h, washed once with HBSS and imaged in HBSS on Abberior Expert line. Protein expression induced with 0.1 µg/ml doxycycline for 24h before imaging. Scale bar 10 µm. See supplementary materials and methods section for further details on imaging settings.

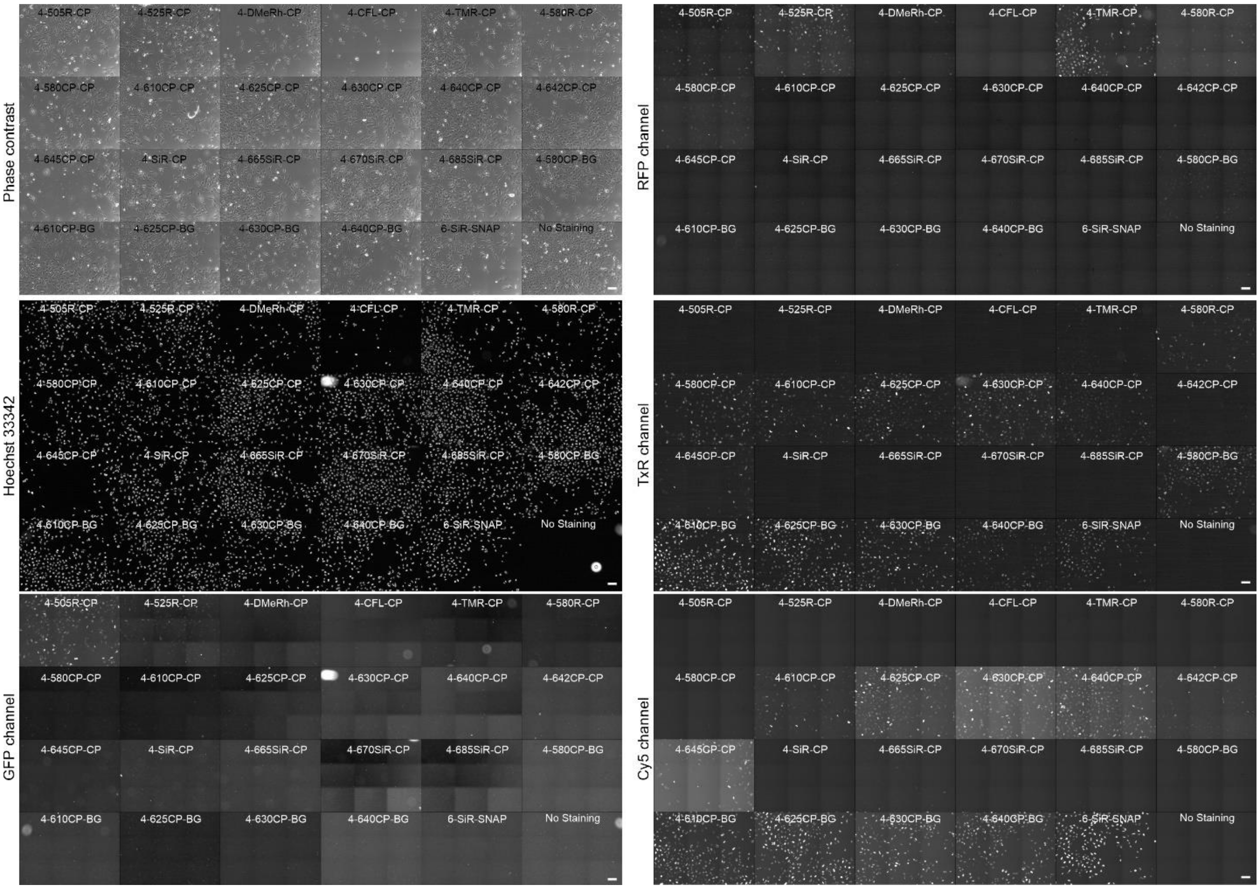

**Supplementary Figure S13.** Wide field fluorescence microscopy images of living U-2 OS cells expressing histone H3-SNAP protein. Cells were stained with a series of different SNAP-tag substrates at 100 nM concentration and 0.1 µg/ml Hoechst 33342 in DMEM growth medium containing 10% FBS at 37 °C for 1 h, washed once with HBSS and imaged in HBSS on Biotek Lionheart FX automated microscope. Note, DMEM growth medium shows strong autofluorescence in GFP channel and was replaced with HBSS to visualize probes in fluorescent in GFP channel. Scale bars 100 µm. See supplementary materials and methods section for further details on imaging settings.

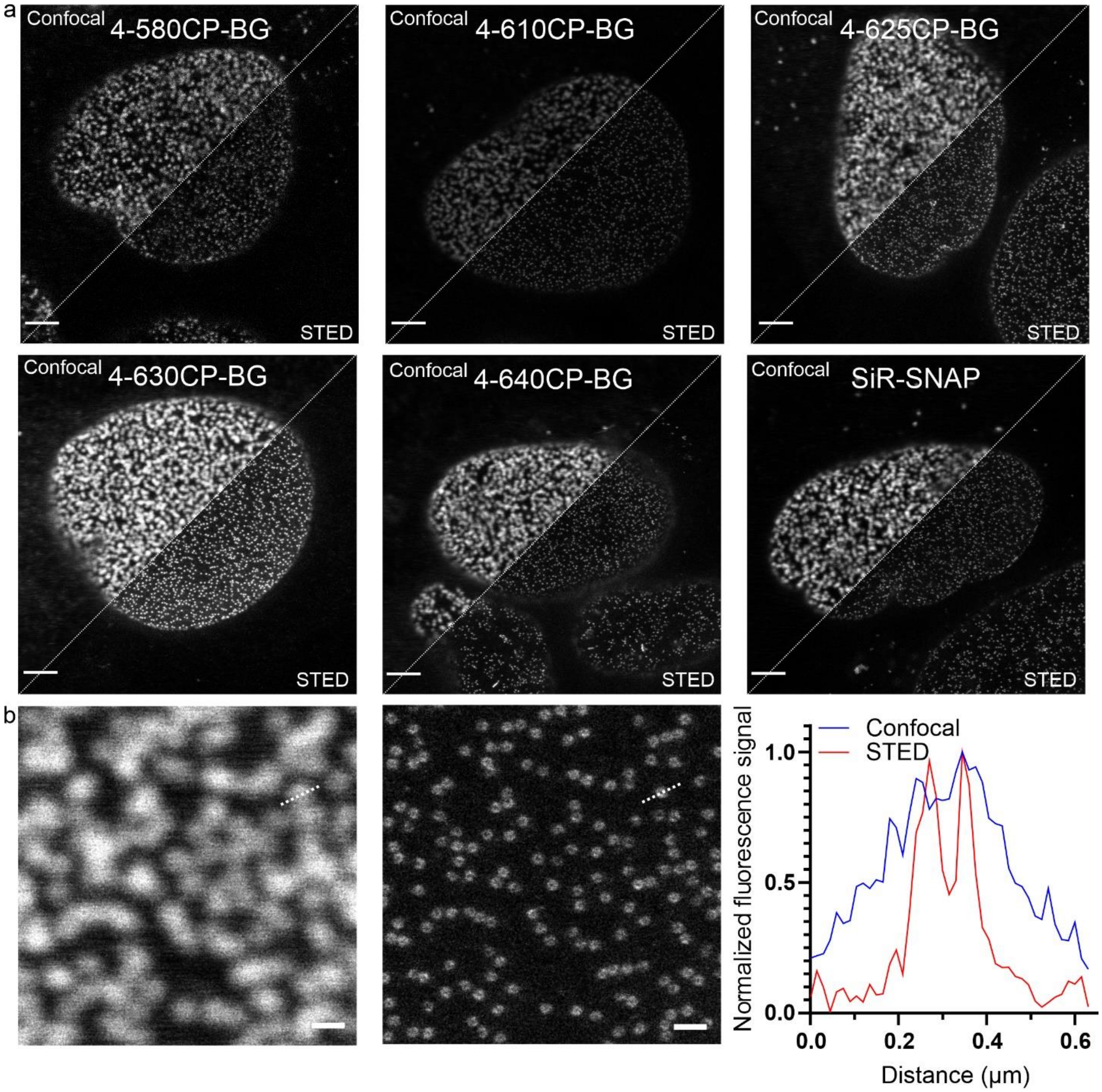

**Supplementary Figure S14. STED imaging of nucleopores tagged with SNAP-tag.** Performance of the new O6-Benzylguanine (BG) derivatives. a. Comparison of confocal and STED images of living U-2 OS cells expressing Nup96-SNAP stained with BG derivatives. Scale bars: 3 µm. b. Confocal (left) and STED (center) zoomed images of U-2 OS cells expressing Nup96-SNAP stained with **4-630CP-BG** probe. Line profile (right) showing normalized fluorescence signal along the dashed line in the images. Scale bar 0.5µm. Living cells were incubated with 100nM of the indicated probe in DMEM containing 10% FBS at 37 °C for 1 h, washed once with HBSS and imaged in DMEM containing 10% FBS. Images were acquired on Abberior Expert line scanning microscope equipped with 775 nm depletion laser. See supplementary materials and methods section for further details on imaging settings.

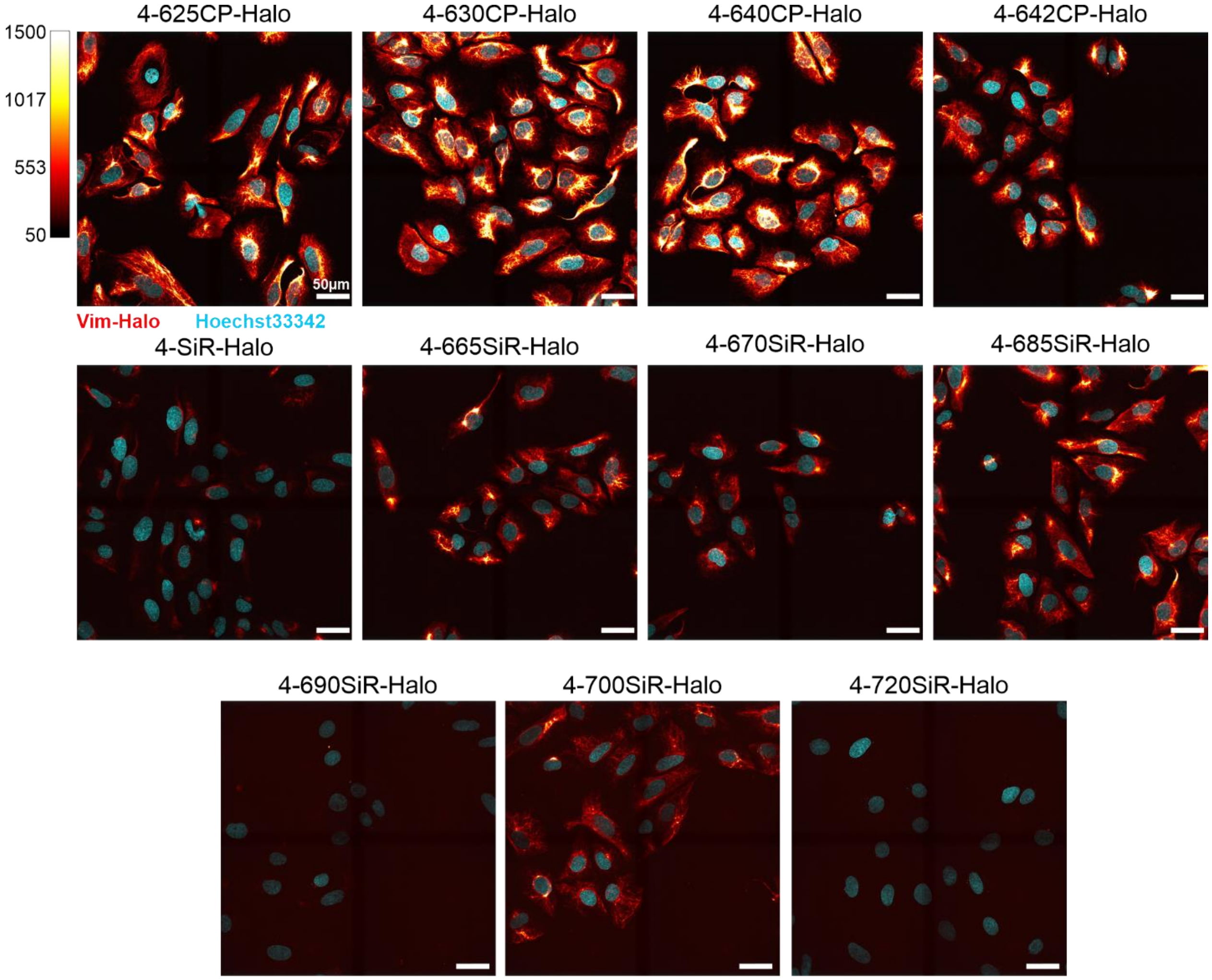

**Supplementary Figure S15. Staining of U-2 OS vimentin-Halo cells with new HaloTag substrates.** Cells were incubated with 100 nM dye in DMEM medium containing 10% FBS for 1h at 37°C, washed once with culture medium followed by wash with HBSS and subsequently imaged in culture medium on a Visitron spinning disk confocal microscope equipped with CFI Plan Apochromat Lambda 60x oil immersion objective (Nikon), using 640 nm laser for excitation and 665 nm long pass emission filter. Four fields of view were acquired as z-stacks with 200 nm step size and were merged with VisiView software. Max intensity projections are shown. Note, that due to the different spectra, staining intensities cannot be compared. See methods section for further details on imaging settings.

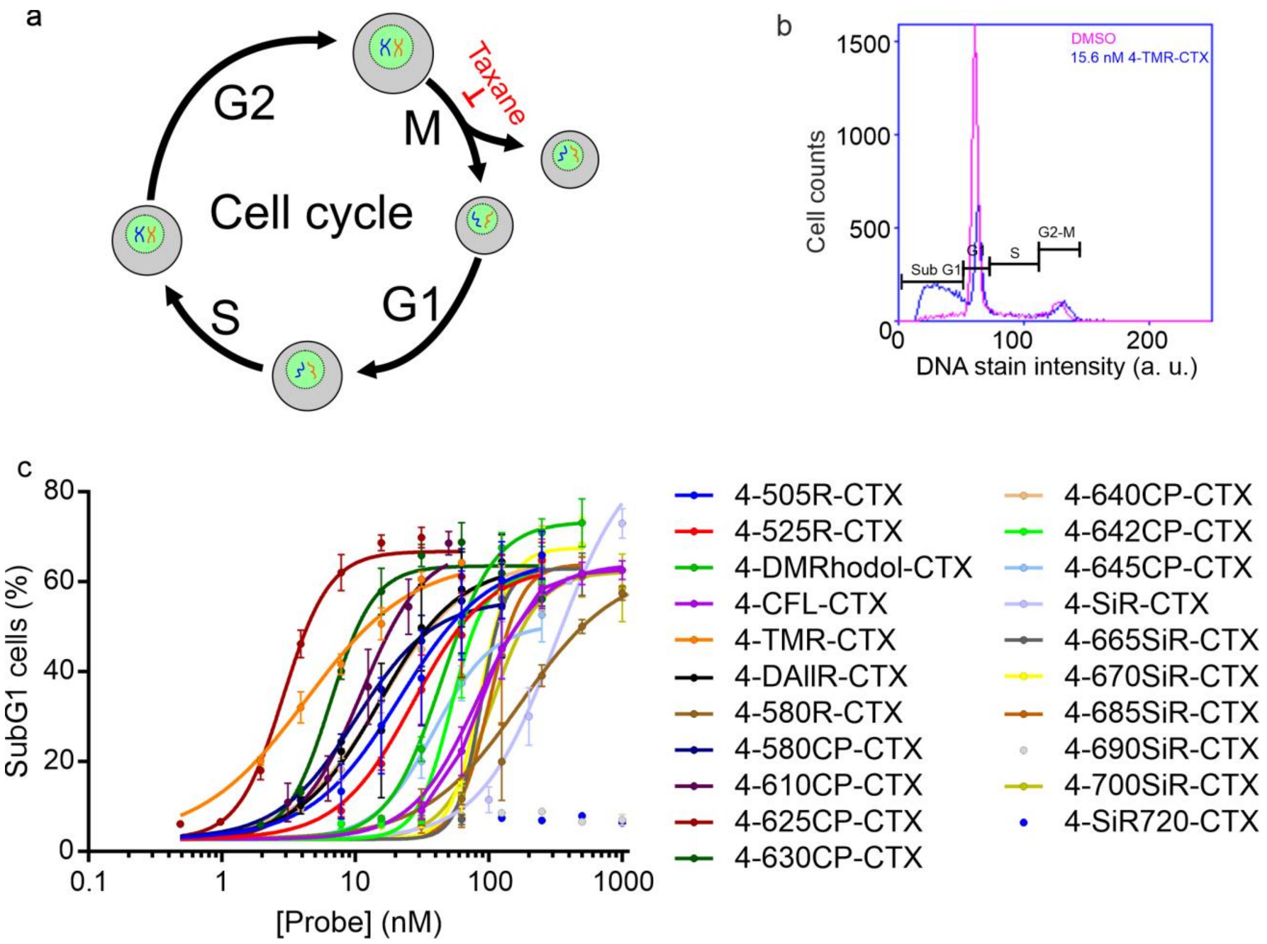

**Supplementary Figure S16.** Cell cycle perturbations in HeLa cells induced by fluorescent tubulin probes. a. Cytotoxicity of taxanes results from the inhibition of the cell cycle at the stage of mitosis (M). b. Representative histogram of DNA content distribution in HeLa cells treated with DMSO or 15.6 nM 4-TMR-CTX for 24 h. The cell cycle phases are identified by the amount of DNA per cell. c. Accumulation of subG1 phase HeLa cells upon treatment with tubulin probes. Data were fitted to the dose response curve to obtain $EC_{50}$.

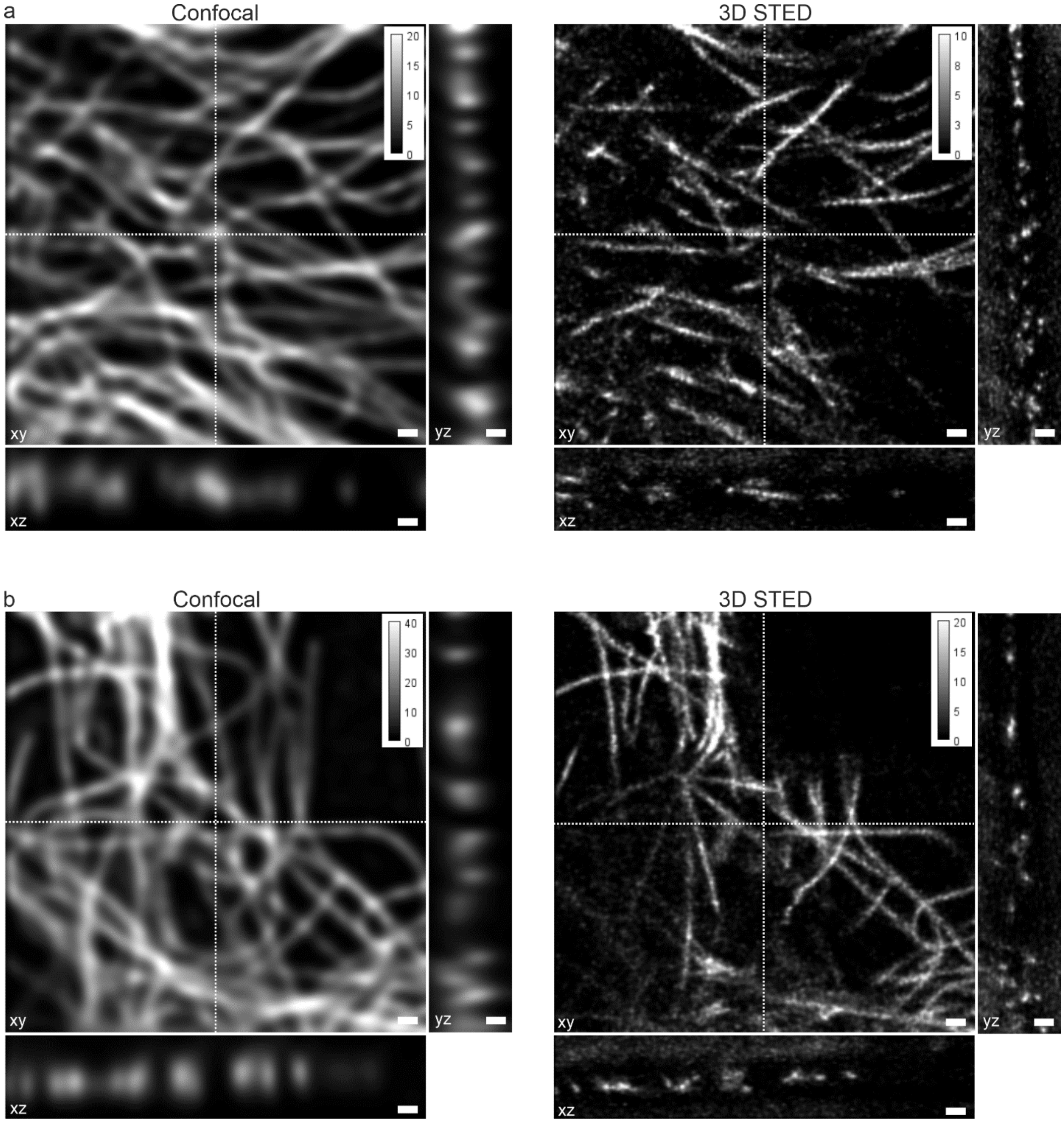

**Supplementary Figure S17.** Deconvolved confocal and 3D STED images of microtubules in living human fibroblasts stained for 1h at 37°C with a) 10 nM and b) 100nM **4-630CP-CTX** probe in the growth DMEM medium and imaged subsequently under no-wash conditions. Deconvolution performed using SVI Huygens software. Dashed white line indicates the position of xy, xz and yz sections. Voxel size: 40 x 40 x 40 nm. Scale bars: 0.5 µm. See supplementary materials and methods section for further details on imaging settings.

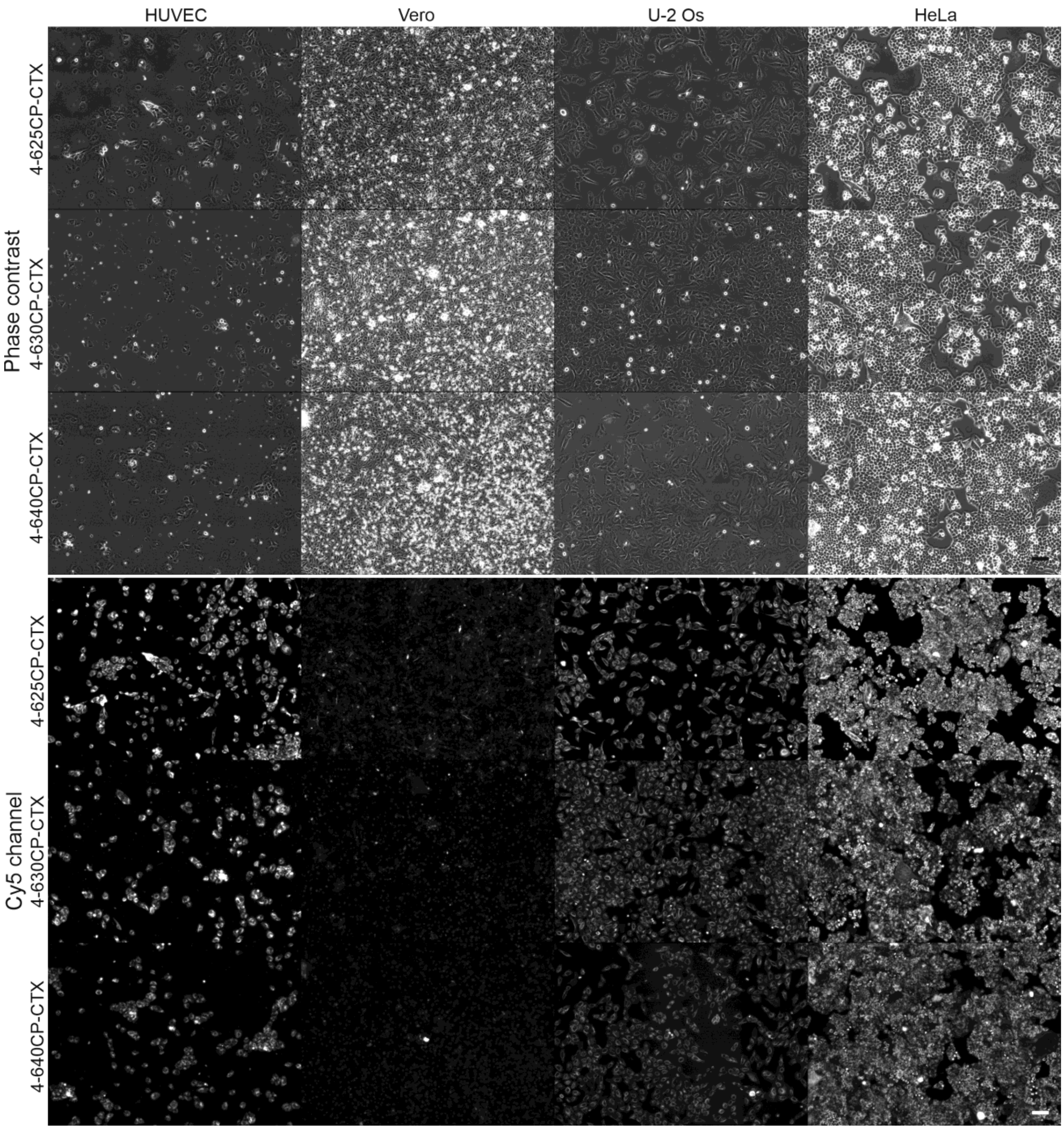

**Supplementary Figure S18.** Wide field fluorescence microscopy images of HUVEC, Vero, U-2 OS and HeLa cells stained with 100 nM **4-625CP-CTX**, **4-630CP-CTX** and **4-640CP-CTX** tubulin probes. Cells were stained in DMEM growth medium containing 10% FBS at 37 °C for 1 h, washed once with HBSS and imaged in DMEM growth medium containing 10% FBS on Biotek Lionheart FX automated microscope. Scale bars 100 µm. See methods section for further details on imaging settings. Note, that staining of Vero cells is the weakest due to high efflux activity of membrane pumps.

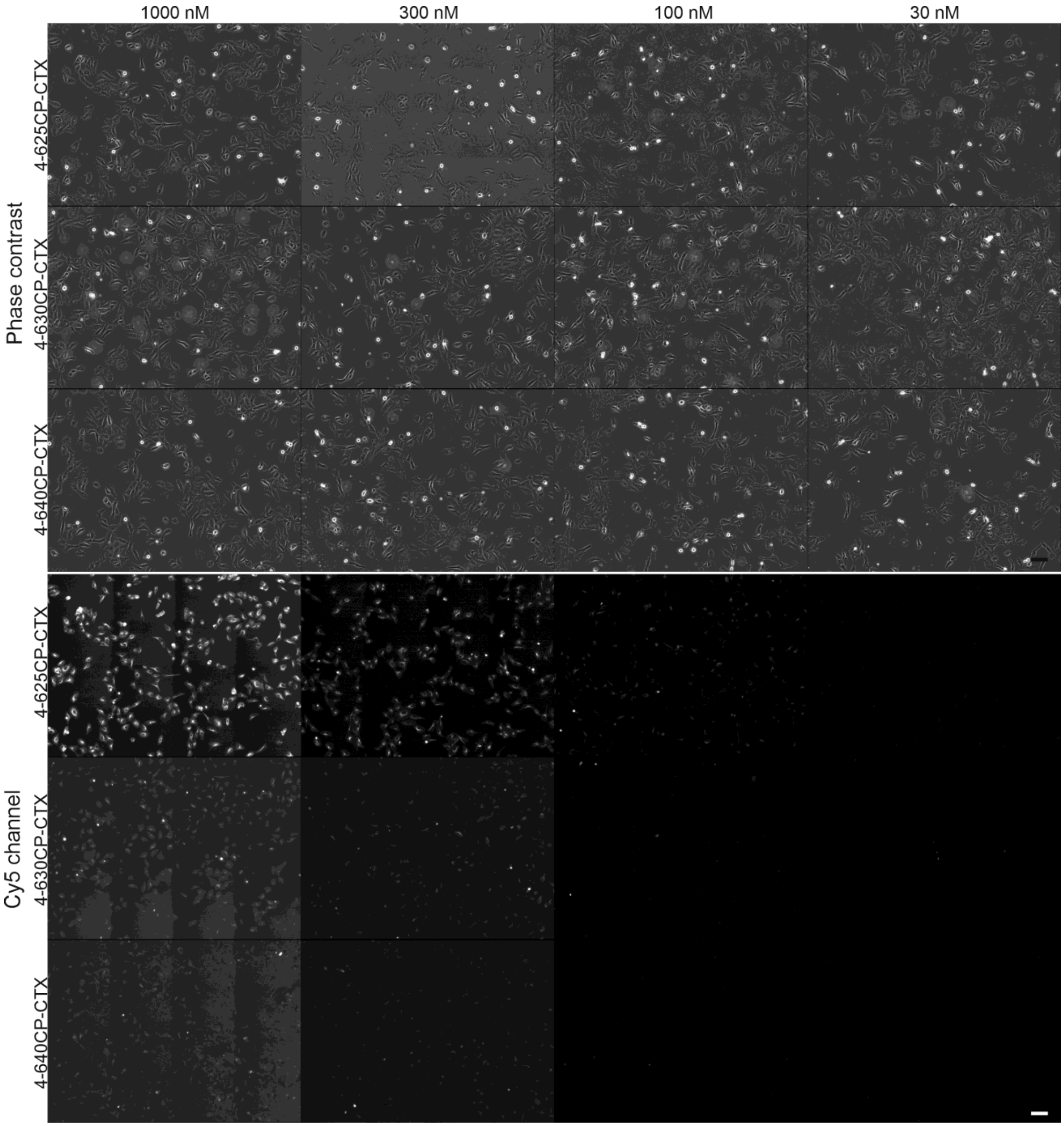

**Supplementary Figure S19.** Wide field fluorescence microscopy images of Vero cells stained with variable concentrations of **4-625CP-CTX**, **4-630CP-CTX** and **4-640CP-CTX** tubulin probes. Cells were stained in DMEM growth medium containing 10% FBS at 37 °C for 1 h, washed once with HBSS and imaged in DMEM growth medium containing 10% FBS on Biotek Lionheart FX automated microscope. Scale bars: 100 µm. See supplementary materials and methods section for further details on imaging settings.

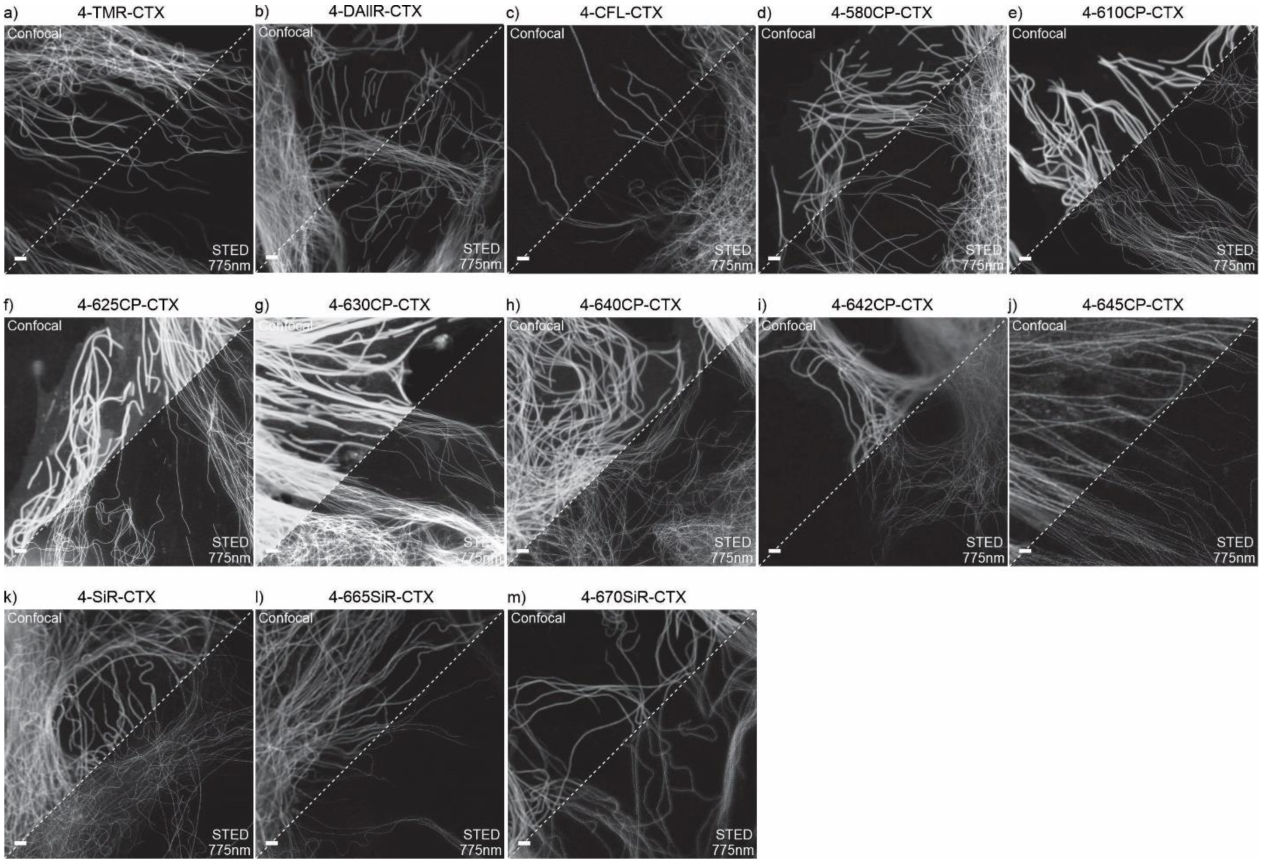

**Supplementary Figure S20.** Confocal and STED images of microtubules in living human fibroblasts under no-wash conditions. Dyes were excited with: a) - d) 3 mW 561 nm laser; e) - m) 0.3 mW 640 nm laser. STED images were acquired with 775 nm depletion laser. Living human fibroblasts were incubated with 100nM of the indicated tubulin probe in growth medium containing 10% FBS at 37 °C for 1 h and imaged subsequently. Images were acquired on Abberior Expert line scanning microscope equipped with 775 nm STED laser. Pixel size: 10 x 10 nm. Scale bars: 2 µm. See supplementary materials and methods section for further details on imaging settings.

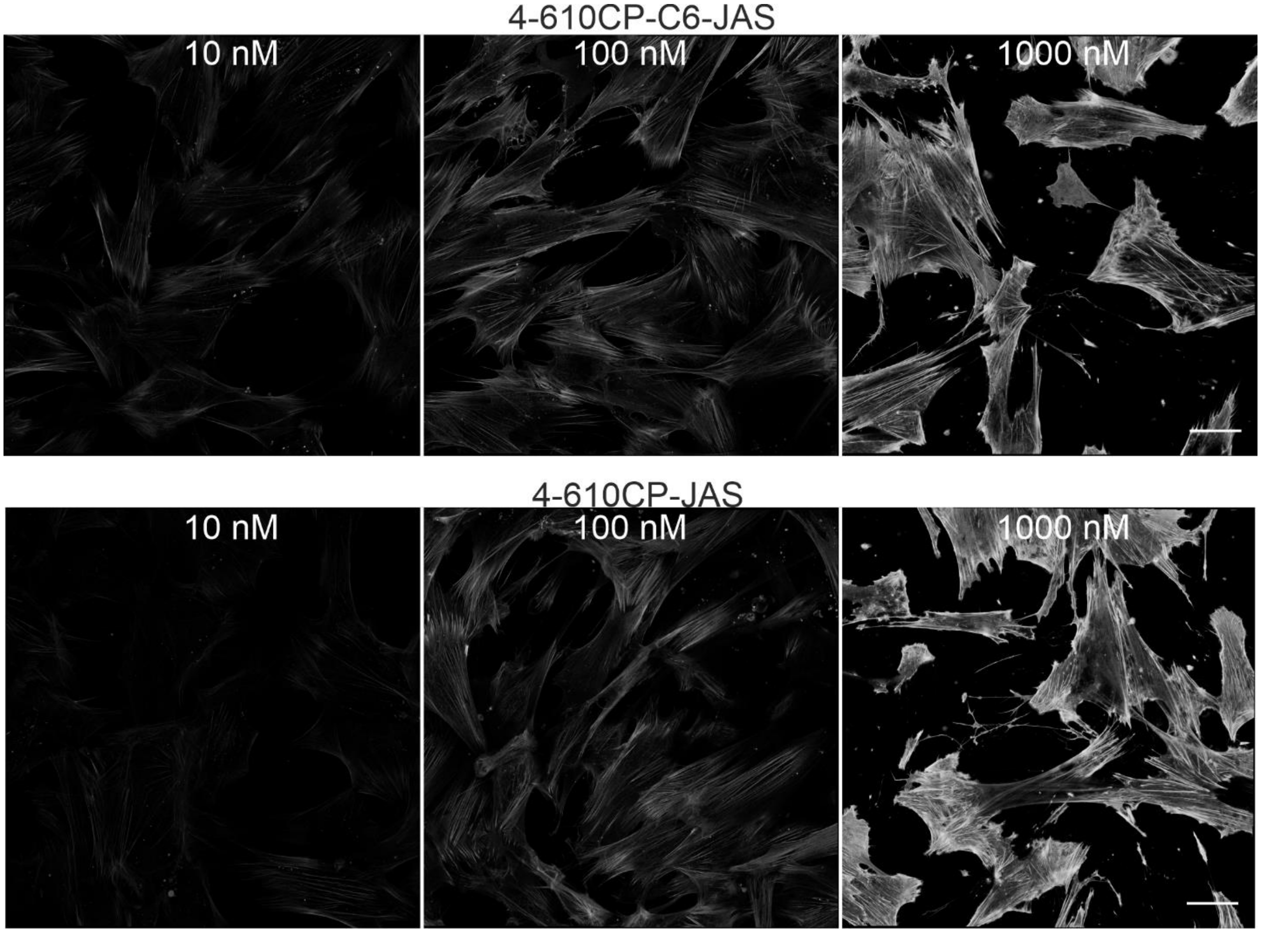

**Supplementary Figure S21.** Confocal microscopy images of actin in human fibroblasts stained with jasplakinolide - based probes containing different linker length probes– **4-610CP-C6-JAS** and **4-610CP-JAS**. Cells were incubated with probes at indicated concentration in DMEM growth medium containing 10% FBS at 37 °C for 1 h and subsequently imaged on Abberior Facility line without removal of the probes. Multiple fields of view were stitched with SVI Huygens software. Scale bars: 50 µm. Note, the performance of both probes is identical. See supplementary materials and methods section for further details on imaging settings.

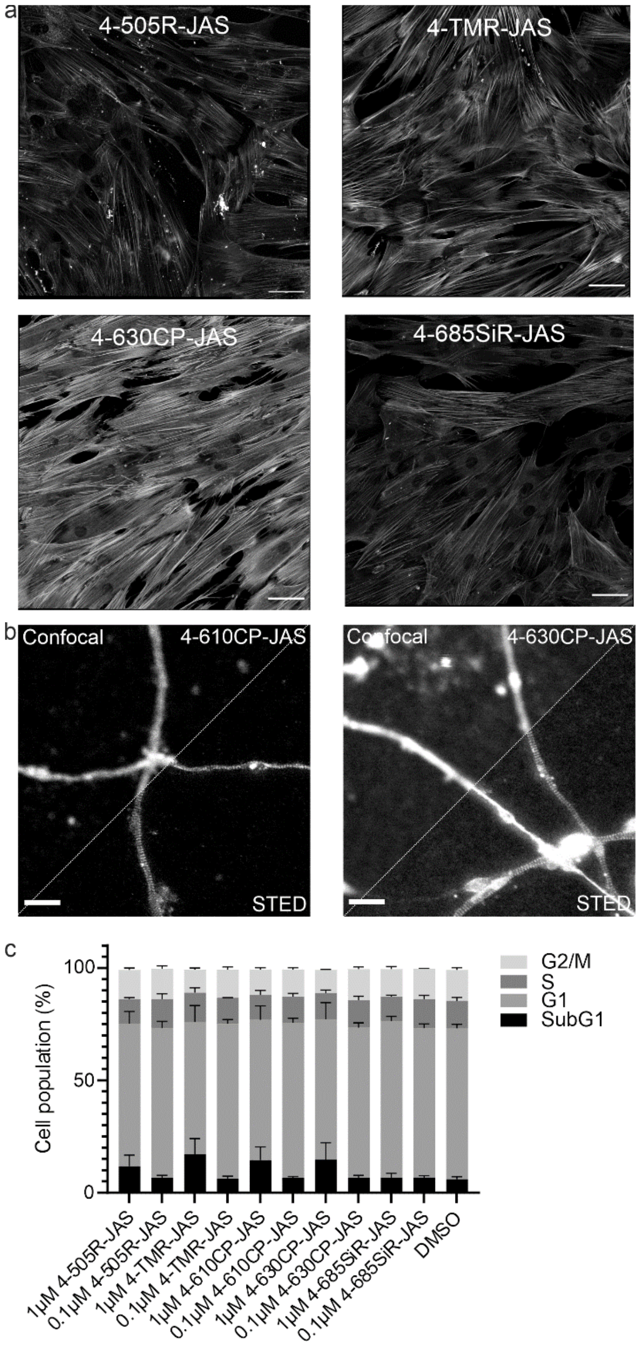

**Supplementary Figure S22.** Properties of the actin probes. a. Confocal microscopy images of live fibroblasts stained with 100 nM actin probes for 1h at 37°C. Cells were imaged in growth DMEM media without probe removal using Abberior Facility line microscope. Multiple fields of view were stitched with SVI Huygens software. Scale bars: 50 µm. b.Confocal and STED images of actin cytoskeleton in neurons stained with 100 nM **4-610CP-JAS** or **4-630CP-JAS** probes in growth media without probe removal. Images aquired on Abberior Expert line microscope. See methods section for further details on imaging settings.c. Cytotoxicity measurements of the actin probes. HeLa cells were incubated with the indicated concentrations of the probes at 37 °C for 24 h in a humidified 5% $CO_2$ incubator. Experimental data are averages of three independent experiments (N = 3, n ≥ 9000 cells) and presented as means with standard deviations.

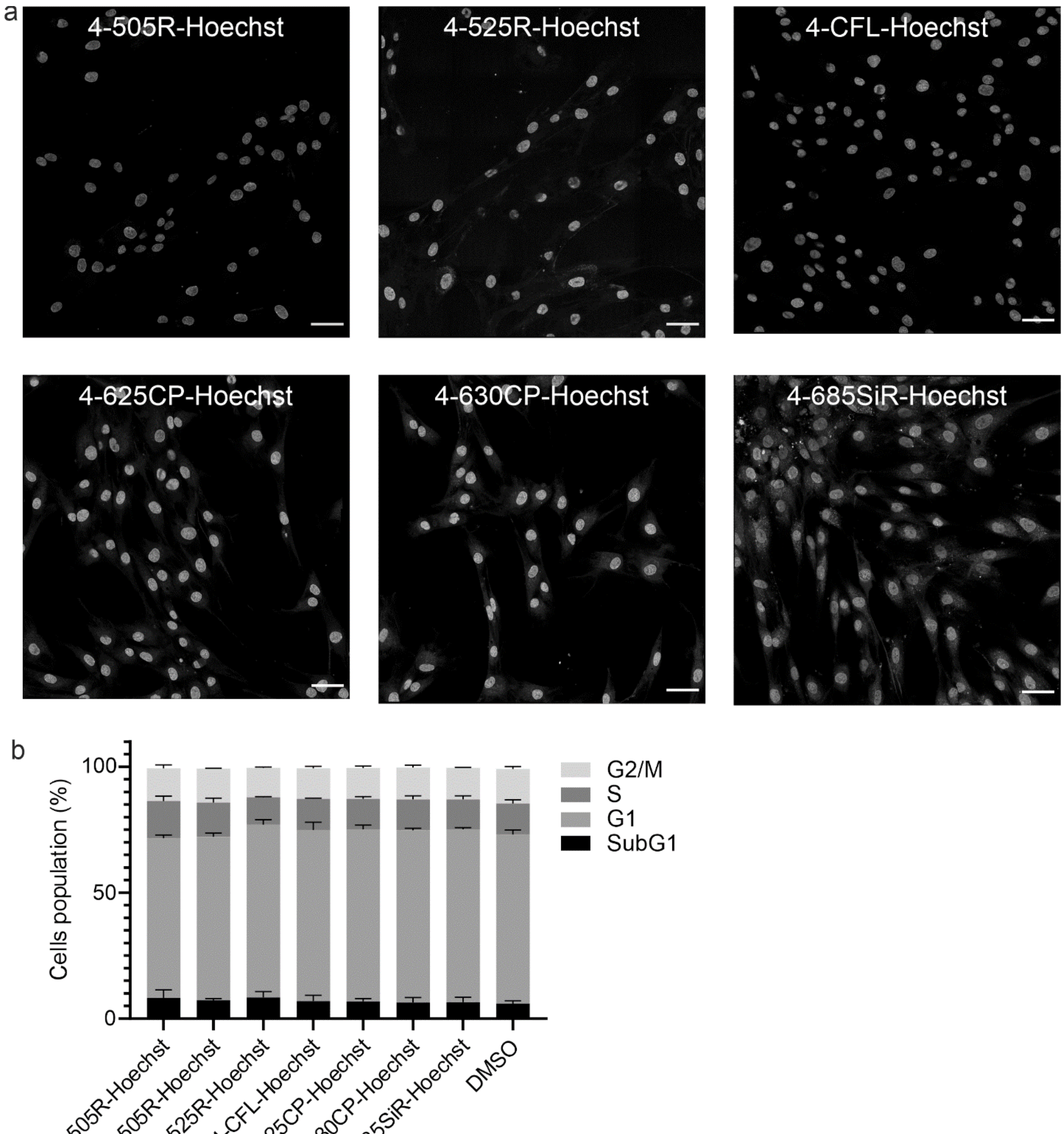

**Supplementary Figure S23.** Properties of DNA probes. a. Confocal microscopy images of live fibroblasts stained with 100 nM DNA probes for 1h at 37°C. Cells were washed once with HBSS and imaged in growth DMEM media on Abberior Facility line microscope. Multiple fields of view were stitched with Huygens software. Scale bars: 50 µm. See supplementary materials and methods section for further details on imaging settings. b. Cytotoxicity measurements of DNA probes. HeLa cells were incubated with the indicated concentrations of Hoechst derivatives at 37 °C for 24 h in a humidified 5% $CO_2$ incubator. Experimental data are averages of three independent experiments (N = 3, 9000 ≤ n ≤ 10000 cells) and presented as means with standard deviations.

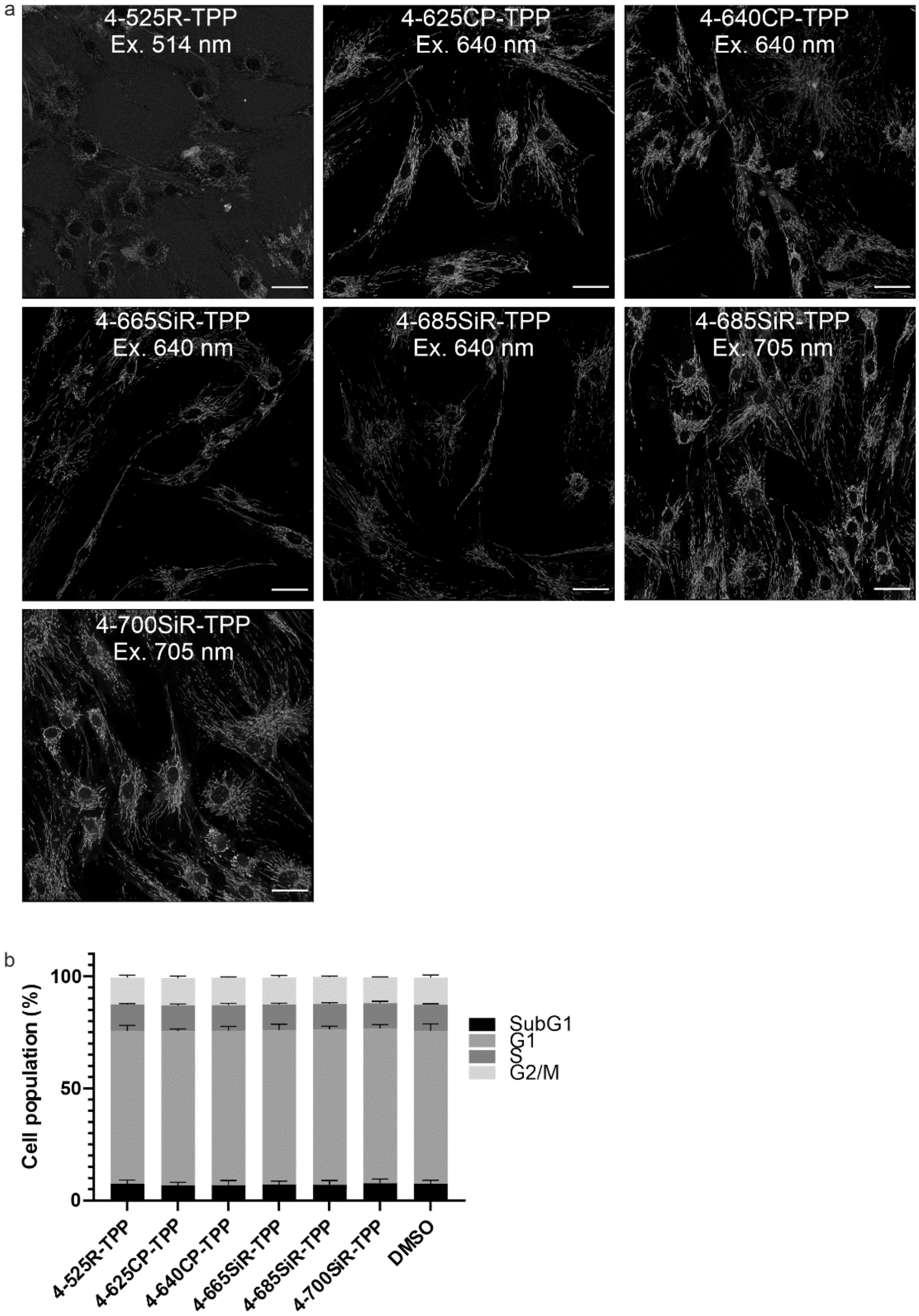

**Supplementary Figure S24. Properties of probes targeting mitochondria.** a. Confocal fluorescence microscopy images of human fibroblasts stained with 100nM triphenylphosphonium (TPP) mitochondrial probes. Cells were stained in DMEM growth medium containing 10% FBS at 37 °C for 1 h and imaged on Abberior Facility line without removal of the probes. Multiple fields of view were stitched with SVI Huygens software. Note, excitation laser wavelength is indicated. Scale bars 50 µm. b. Cytotoxicity measurements of mitochondrial probes. HeLa cells were incubated with 100 nM probes at 37 °C for 24 h in a humidified 5% $CO_2$ incubator. Experimental data are averages of three independent experiments (N = 3, 9000 ≤ n ≤ 10000 cells) and presented as means with standard deviations. See supplementary materials and methods section for further details on imaging settings.

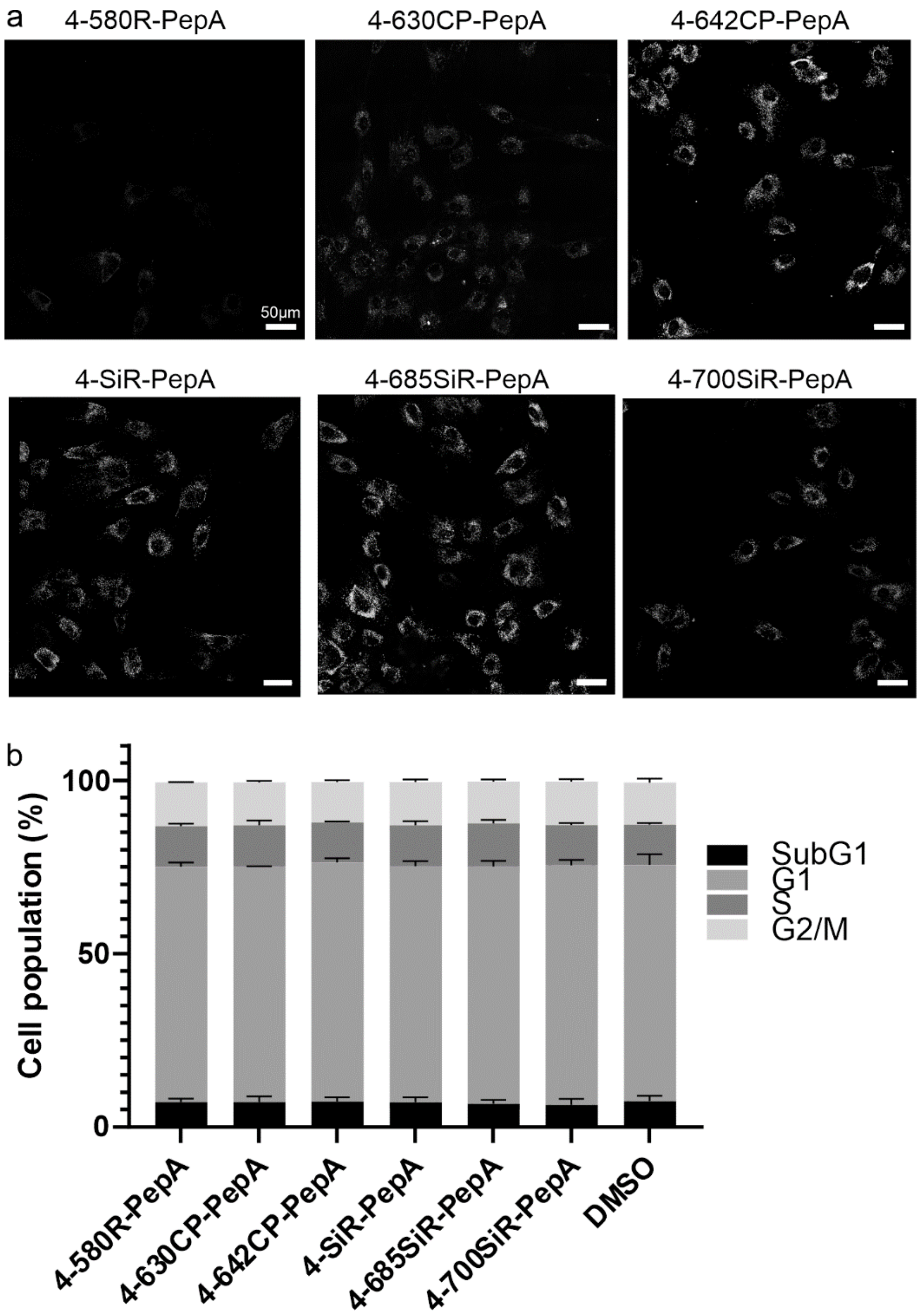

**Supplementary Figure S25. Properties of lysosomal probes.** a. Staining of human fibroblasts with new dye-pepstatin A conjugates. The cells were incubated with 100 nM probe in DMEM medium with FBS for 1h at 37°C, washed once with medium and once with HBSS and imaged in the same medium on a Abberior facility line microscope in a confocal imaging mode. 25 fields of view were acquired and were stitched with SVI Huygens software. Note, that due to the different spectra, staining intensities cannot be compared. b. Cytotoxicity measurements of lysosomal probes. HeLa cells were incubated with 100 nM probes at 37 °C for 24 h in a humidified 5% $CO_2$ incubator. Experimental data are averages of three independent experiments (N = 3, 9000 ≤ n ≤ 10000 cells) and presented as means with standard deviations. See supplementary materials and methods section for further details on imaging settings.

#### Supplementary Tables

**Supplementary Table S1.** Photophysical properties of the synthesized 4-carboxyrhodamines in PBS (pH =7.4) +0.1%SDS buffer.

| Dye (-COOH) | $\lambda_{max}^{abs}$ (nm) | $\lambda_{max}^{em}$ (nm) | ε x $10^2$ ($m^{-1}$ $cm^{-1}$) | QY (%) | τ (ns) |
|---|---|---|---|---|---|
| **4-505R (22)** | 504 | 524 | 672 ± 16 | 98 ± 2 | 4.16 ± 0.02 |
| **4-525R (23)** | 526 | 547 | 821 ± 26 | 98 ± 3 | 4.39 ± 0.06 |
| **4-DMRhol (25)** | 521 | 547 | 683 ± 44 | 20 ± 1 | – |
| **4-TAllR (17)** | 551 | 573 | 991 ± 104 | 90 ± 6 | 4.12 ± 0.04 |
| **4-DAllR (18)** | 554 | 574 | 816 ± 31 | 75 ± 2 | 3.86 ± 0.01 |
| **4-CFL (26)** | 549 | 574 | 132 ± 2.3 | 64 ± 4 | 3.59 ± 0.01 |
| **4-TMR (3)** | 554 | 575 | 778 ± 45 | 59 ± 1 | 2.99 ±0.01 |
| **4-580R (5)** | 580 | 598 | 896 ± 43 | 86 ± 1 | 4.82 ± 0.02 |
| **4-580CP (24)** | 591 | 613 | 1166 ± 55 | 66 ± 1 | 4.20 ± 0.01 |
| **4-610CP (4)** | 616 | 640 | 1055 ± 94 | 67 ± 1 | 4.23 ± 0.01 |
| **4-DAllCP (19)** | 617 | 640 | 1035 ± 185 | 75 ± 2 | 4.70 ± 0.04 |
| **4-625CP (10)** | 627 | 653 | 1032 ± 10 | 60 ± 1 | 4.19 ± 0.02 |
| **4-630CP (11)** | 629 | 647 | 1100 ± 74 | 77± 4 | 4.53 ± 0.04 |
| **4-640CP (7)** | 641 | 666 | 1092 ± 83 | 46 ± 1 | 3.79 ± 0.01 |
| **4-642CP (12)** | 648 | 680 | 1157 ± 31 | 40± 2 | 2.76 ± 0.01 |
| **4-645CP (6)** | 645 | 661 | 1086 ± 78 | 80 ± 3 | 4.61 ± 0.05 |
| **4-SiR (2)** | 655 | 674 | 733 ± 51 | 62 ± 1 | 3.95 ± 0.03 |
| **4-665SiR (13)** | 669 | 687 | 1004 ± 37 | 55 ± 2 | 3.84 ± 0.02 |
| **4-670SiR (14)** | 675 | 699 | 764 ± 78 | 41 ± 1 | 3.23 ± 0.01 |
| **4-685SiR (15)** | 693 | 712 | 1076 ± 100 | 44 ± 2 | 3.46 ± 0.04 |
| **4-690SiR (16)** | 696 | 724 | 730 ± 39 | 20 ± 1 | 1.96 ± 0.01 |
| **4-700SiR (8)** | 700 | 726 | 920 ± 26 | 24.6 ± 0.3 | 2.62 ± 0.01 |
| **4-720SiR (9)** | 732 | 764 | 636 ± 43 | 15 ± 1 | 1.74 ± 0.02 |

**Supplementary Table S2.** Measured $D_{50}$ and $K_{L\text{-}Z}$ values of the free dyes and respective 4-(benzyloxy)-2-chloropyrimidine (CP) probes.

| Dye | -COOH | -CONH- | -COOH | -CONH- |
|---|---|---|---|---|
| | $^{Dye}D_{50}$ | $^{Probe}D_{50}$ | $^{Dye}K_{L\text{-}Z}$ | $^{Probe}K_{L\text{-}Z}$ |
| **4-505R (22)** | 13 ± 1 | 30 ± 1 | 4.1 ± 1.1 | 2.0 ± 0.3 |
| **4-525R (23)** | 8.9 ± 0.3 | 18.1 ± 0.3 | 9.4 ± 1.3 | 7.2 ± 0.6 |
| **4-DMRhol (25)** | 38 ± 3 | 20.4 ± 0.5 | 1.0 ± 0.1 | 0.17 ± 0.01 |
| **4-TAllR (17)** | 20.4 ± 0.4 | –[a] | 3.5 ± 1.4 | –[a] |
| **4-DAllR (18)** | 15.4 ± 0.4 | –[a] | 9.8 ± 0.2 | –[a] |
| **4-CFL (26)** | –[b] | –[b] | –[b] | –[b] |
| **4-TMR (3)** | 14 ± 1 | 23.4 ± 0.3 | 8.5 ± 0.8 | 7.9 ± 0.5 |
| **4-580R (5)** | 3.8 ± 0.4 | 6.5 ± 0.2 | 10.22 ± 0.02 | 9.3 ± 2.8 |
| **4-580CP (24)** | 30.8 ± 0.7 | 59 ± 2 | 1.0 ± 0.1 | 0.23 ± 0.01 |
| **4-610CP (4)** | 32.1 ± 0.7 | 65 ± 2 | 0.99 ± 0.04 | 0.19 ± 0.08 |
| **4-DAllCP (19)** | 40.0 ± 0.4 | –[a] | 0.4 ± 0.1 | –[a] |
| **4-625CP (10)** | 25 ± 1 | 44.0 ± 0.7 | 5.7± 1.3 | 0.5 ± 0.1 |
| **4-630CP (11)** | 17.6 ± 0.6 | 35.0 ± 0.3 | 7.9 ± 2.2 | 1.75 ± 0.04 |
| **4-640CP (7)** | 19.1 ± 0.4 | 37.5 ± 0.4 | 6.5 ± 2.8 | 1.5 ± 0.2 |
| **4-642CP (12)** | 32.0 ± 0.6 | 65 ± 4 | 1.4 ± 0.4 | 0.5 ± 0.2 |
| **4-645CP (6)** | 8.1 ± 0.3 | 19.1 ±0.4 | 13.5 ± 2.8 | 12.0 ± 4.1 |
| **4-SiR (2)** | >80 | >80 | 0.04 ± 0.01 | 0.001 |
| **4-665SiR (13)** | 37.2 ± 0.6 | 68 ± 4 | 1.5 ± 0.8 | 0.13 ± 0.07 |
| **4-670SiR (14)** | >80 | >80 | 0.2 ± 0.1 | 0.001 |
| **4-685SiR (15)** | 29.6 ± 0.5 | 65 ± 3 | 1.6 ± 0.3 | 0.4 ± 0.2 |
| **4-690SiR (16)** | >80 | >80 | 0.06 ± 0.04 | 0.001 |
| **4-700SiR (8)** | >80 | >80 | 0.23 ± 0.04 | 0.001 |
| **4-720SiR (9)** | >80 | >80 | 0.4 ± 0.4 | 0.0001 |

[a] The corresponding 4-(benzyloxy)-2-chloropyrimidine (CP) derivatives were not synthesized. [b] The 4-CFL showed no change along increasing amount of water in 1,4-dioxane mixture and neither $D_{50}$ nor $K_{L\text{-}Z}$ values were obtained.

**Supplementary Table S3.** TDDFT 6-311++G (d,p)/IEFPCM (water) calculated vertical excitation energy and wavelength values before and after empirical correction of 0.4 eV and experimentally measured absorption maxima values of the studied dyes.

| | Calculated by TDDFT | | Empirically corrected | | Experimentally measured | |
|---|---|---|---|---|---|---|
| **Dye (-COOH)** | **$\Delta E_{H-L}$ (eV)** | **$\lambda_{abs}$ (nm)[a]** | **$\Delta E_{H-L}$ (-0.4 eV)** | **$\lambda_{abs}$ (nm)** | **$\lambda_{abs}$ (nm)** | **$\Delta E_{H-L}$ (ev)[b]** |
| **4-505R (22)** | 2.888 | 429 | 2.489 | 498 | 500 | 2.480 |
| **4-525R (23)** | 2.783 | 445 | 2.383 | 520 | 522 | 2.375 |
| **4-TMR (3)** | 2.635 | 470 | 2.236 | 555 | 551 | 2.250 |
| **4-580R (5)** | 2.536 | 489 | 2.137 | 580 | 579 | 2.141 |
| **4-580CP (24)** | 2.564 | 483 | 2.164 | 573 | 585 | 2.119 |
| **4-610CP (4)** | 2.460 | 503 | 2.061 | 602 | 611 | 2.029 |
| **4-625CP (10)** | 2.416 | 513 | 2.016 | 615 | 621 | 1.996 |
| **4-630CP (11)** | 2.416 | 513 | 2.017 | 615 | 624 | 1.987 |
| **4-640CP (7)** | 2.363 | 525 | 1.964 | 631 | 636 | 1.949 |
| **4-642CP (12)** | 2.305 | 538 | 1.906 | 651 | 641 | 1.934 |
| **4-645CP (6)** | 2.347 | 528 | 1.947 | 637 | 642 | 1.931 |
| **4-SiR (2)** | 2.353 | 527 | 1.954 | 635 | 649 | 1.910 |
| **4-665SiR (13)** | 2.295 | 540 | 1.895 | 654 | 663 | 1.870 |
| **4-670SiR (14)** | 2.280 | 544 | 1.880 | 659 | 668 | 1.856 |
| **4-685SiR (15)** | 2.219 | 559 | 1.820 | 681 | 687 | 1.804 |
| **4-690SiR (16)** | 2.165 | 573 | 1.770 | 702 | 687 | 1.805 |
| **4-700SiR (8)** | 2.200 | 564 | 1.800 | 689 | 694 | 1.786 |
| **4-720SiR (9)** | 2.039 | 608 | 1.639 | 756 | 721 | 1.719 |

Supplementary Table S4. **Properties of tubulin probes**

| Probe | $EC_{50}$ (nM) | Isat ($MW \cdot cm^2$) | App. $FWHM_{min}$ (nm) | Microtubule to cytosol signal ratio[b] |
|---|---|---|---|---|
| **4-505R-CTX (48)** | 28 ± 4 | n.a. | n.a. | n.a. |
| **4-525R-CTX (49)** | 34 ± 4 | n.a. | n.a. | n.a. |
| **4-DMRhol-CTX (50)** | 48 ± 3 | n.a. | n.a. | n.a. |
| **4-DAllR-CTX (52)** | 21 ± 3 | 16 ± 3 | 121 ± 32[a] | 17 ± 5 |
| **4-CFL-CTX (47)** | 91 ± 7 | 8 ± 2 | 96 ± 38 [a] | 27 ± 12 |
| **4-TMR-CTX (51)** | 6 ± 2 | 13 ± 1 | 102 ± 24 [a] | 50 ± 25 |
| **4-580R-CTX (53)** | 239 ± 30 | n.a. | n.a. | n.a. |
| **4-580CP-CTX (54)** | 12 ± 1 | 3.9 ± 0.5 | 77 ± 26 [a] | 16 ± 5 |
| **4-610CP-CTX (55)** | 15 ± 2 | 0.62 ± 0.07 | 34 ± 10 [a] | 35 ± 14 |
| **4-625CP-CTX (56)** | 3.1 ± 0.2 | 0.57 ± 0.05 | 37 ± 12 [a] | 16 ± 3 |
| **4-630CP-CTX (57)** | 7.0 ± 0.4 | 0.93 ± 0.09 | 45 ± 11 [a] | 9 ± 2 |
| **4-640CP-CTX (58)** | 19 ± 2 | 0.45 ± 0.04 | 32 ± 11 [a] | 7 ± 2 |
| **4-642CP-CTX (59)** | 61 ± 58 | 0.32 ± 0.02 | 27 ± 12 [a] | 15 ± 5 |
| **4-645CP-CTX (60)** | 46 ± 5 | 0.61 ± 0.03 | 34 ± 11 [a] | 17 ± 5 |
| **4-SiR-CTX (61)** | 248 ± 17 | 0.39 ± 0.04 | 24 ± 10 [a] | 79 ± 29 |
| **4-665SiR-CTX (62)** | 98 ± 11 | 0.21 ± 0.02 | 21 ± 9 [a] | 29 ± 10 |
| **4-670SiR-CTX (63)** | 93 ± 5 | 0.18 ± 0.01 | 29 ± 20[b] | 25 ± 10 |
| **4-685SiR-CTX (64)** | 114 ± 8 | n.a. | n.a. | n.a. |
| **4-690SiR-CTX (65)** | n.a. | n.a. | n.a. | n.a. |
| **4-700SiR-CTX (66)** | 96 ± 33 | n.a. | n.a. | n.a. |
| **4-720SiR-CTX (67)** | n.a. | n.a. | n.a. | n.a. |

a - values calculated from STED images (775 nm laser, 100% power).

b - measured at 50% STED laser power.

**Supplementary Table S5.** Comparison of microtubule diameter measured in cells stained with **4-625CP-CTX** and **4-630CP-CTX** in 3D confocal and 3D STED mode. Data represents mean ± SD (N=30)

| | **3D Confocal** | | | | **3D STED** | | | | |
|---|---|---|---|---|---|---|---|---|---|
| **Dye-CTX** | xz plane, nm | | zy plane, nm | | xz plane, nm | | zy plane, nm | | Microtubule to cytosol signal ratio |
| | **$FWHM_x$** | **$FWHM_z$** | **$FWHM_y$** | **$FWHM_z$** | **$FWHM_x$** | **$FWHM_z$** | **$FWHM_y$** | **$FWHM_z$** | |
| **4-625CP** (10 nM) | 396 ± 87 | 700 ± 122 | 349 ± 78 | 669 ± 92 | 112 ± 16 | 89 ± 16 | 111 ± 20 | 95 ± 14 | 20 ± 12 |
| **4-630CP** (10 nM) | 317 ± 88 | 607 ± 124 | 299 ± 86 | 554 ± 79 | 158 ± 32 | 108 ± 21 | 150 ± 34 | 126 ± 22 | 12 ± 7 |

**Supplementary Table S6. List of Videos and imaging parameters.**

| | | | excitation / emission | | | | |
|---|---|---|---|---|---|---|---|
| | | | 488 nm / 500-550 nm | 561 nm / 580-620 nm | 640 nm / 660-680 nm | 700 nm / 720-770 nm | line acculation |
| **Video S1** | 100 nM 4-700SiR-TPP | human fibroblasts | 10% | 15% | 20% | 10% | 2 |
| | 30 nM 4-642CP-PepA | | | | | | |
| | 30 nM 4-DAllR-CTX | | | | | | |
| | 1000 nM 4-505R-Hoe | | | | | | |
| **Video S2** | 100 nM 4-700SiR-TPP | HUVEC | 10% | 20% | 20% | 5% | 1 |
| | 10 nM 4-642CP-PepA | | | | | | |
| | 10 nM 4-DAllR-CTX | | | | | | |
| | 100 nM 4-505R-Hoe | | | | | | |
| **Video S3** | 100 nM 4-700SiR-TPP | human fibroblasts | 15% | 50% | 20% | 5% | 2 |
| | 30 nM 4-642CP-PepA | | | | | | |
| | 1000 nM 4-CFL-Hoe | | | | | | |
| | 1000 nM 4-505R-CTX | | | | | | |
| **Video S4** | 100 nM 4-700SiR-TPP | U-2 OS vimentin-HaloTag | 25% | 25% | 30% | 5% | 2 |
| | 100 nM 4-642CP-Halo | | | | | | |
| | 30 nM 4-DAllR-CTX | | | | | | |
| | 1000 nM 4-505R-Hoe | | | | | | |
| | | *line step #1* | | ✓ | | ✓ | |
| | | *line step #2* | ✓ | | ✓ | | |

#### Supplementary movies

**Video S1.** Human fibroblasts stained with **4-700SiR-TPP** (mitochondria), **4-642CP-PepA** (lysosomes), **4-DAllR-CTX** (microtubules) and **4-505R-Hoechst** (nucleus).

**Video S2.** HUVEC cells stained with **4-700SiR-TPP** (mitochondria), **4-642CP-PepA** (lysosomes), **4-DAllR-CTX** (microtubules) and **4-505R-Hoechst** (nucleus).

**Video S3**. Human fibroblasts stained with **4-700SiR-TPP** (mitochondria), **4-642CP-PepA** (lysosomes), **4-CFL-Hoechst** (nucleus) and **4-505R-CTX** (microtubules).

**Video S4.** U-2 OS vimentin-HaloTag cells stained with **4-700SiR-TPP** (mitochondria), **4-642CP-Halo** (vimentin), **4-DAllR-CTX** (microtubules) and **4-505R-Hoe** (nucleus).

Imaging parameters are listed in **Supplementary Table S6.** Cell staining and imaging are described in details in Supplementary materials and methods section.

### Supplementary materials and methods

#### Computation, molecular biology and biochemical methods

*Measurements of absorbance spectra in 1,4–dioxane-water mixtures*

Measurements of the absorbance changes in 1,4-dioxane-water mixtures were performed by pipetting 2µLof 1mM stock solutions of dyes or probes in DMSO into a 96 propylene bottom well plate (11 wells per sample) made from propylene (Corning 3364). To the wells going from right to left 300 µL of 1,4-dioxane-water mixtures containing 100%, 90%, 80%, 70%, 60%, 50%, 40%, 30%, 20%, 10% or 0% of 1,4-dioxane (if needed, mixtures with 0.3% SDS are used, with an exception in 100% dioxane were SDS is not soluble). After incubation for 1 hour at room temperature, absorption of solutions in each well was recorded from 320 nm to 850 nm with wavelength step size of 1 nm on a multiwell plate reader Spark® 20M (Tecan). The background absorption of the propylene bottom plate was measured in wells containing only 1,4–dioxane-water mixture with similar amount of DMSO and subtracted from the spectra of the samples. $D_{50}$ values were obtained by fitting plots to dose-response equation $EC_{50}$ (1) as implemented in GraphPad 9.0 software[1]:

$$A = A_0 + (A_{max} - A_0)/\left(1 + \left(\frac{D_{50}}{d}\right)^{Hill}\right) \qquad (1)$$

Where $A_o$ – absorbance at $\lambda_{max}$ at $\varepsilon_r = 0$, $A_{max}$ – the highest reached absorbance at $\lambda_{max}$. $d$ – dielectric constant of 1,4-dioxane-water mixture at a given point, *Hill* - Hill slope coefficient determining the steepness of a dose-response curve, $D_{50}$ - corresponds to d value that provokes half of the absorbance amplitude ($A_{max}$-$A_0$).

*Determination of* $K_{L\text{-}Z}$ values

Measurements of the absorbance changes in 1,4-dioxane-water 1:1 mixtures and ethanol + 0.1%TFA were performed by pipetting 2µL of 1mM stock solutions of dyes or probes in DMSO into a 96 propylene bottom well plate (Corning 3364) and adding 300 µL of corresponding solvent. After incubation for 1 hour at room temperature, absorption of solutions in each well was recorded from 320 nm to 850 nm with wavelength step size of 1 nm on a multiwell plate reader Spark® 20M (Tecan). The background absorption of the propylene bottom plate was measured in wells containing only 1,4–dioxane-water and ethanol+0.1%TFA mixture with similar amount of DMSO and subtracted from the spectra of the samples. The $K_{L\text{-}Z}$ were determined based on procedure published before[2] and was done as follows: we calculated $K_{L–Z}$ using the following equation: $K_{L–Z} = (\varepsilon_{dw}/\varepsilon_{max})/(1 - \varepsilon_{dw}/\varepsilon_{max})$. $\varepsilon_{dw}$ is the extinction coefficient of the dyes/probes in a 1:1 dioxane:water solvent mixture;. $\varepsilon_{max}$ refers to the maximal extinction coefficients measured in ethanol + 0.1%TFA. As noted previously[2] accurate determination of low $K_{L–Z}$ values ($\leq 10^{-3}$) is complicated by the relatively poor sensitivity of absorbance measurements. $K_{L–Z}$ = 10−3 was estimated if we observed a small but measurable absorbance signal in 1:1 dioxane:water solvent mixture over the dye-free blank; $K_{L–Z}$ of $10^{-4}$ was estimated if we observed no measurable absorbance of the dye solution.

*Quantum chemical calculations*

The initial geometries of the studied molecules were generated by using a molecular mechanics method (force field MMF94, steepest descent algorithm) and a systematic conformational analysis was carried out as implemented in Avogadro 1.1.1 software. The minimum energy conformer geometries found by molecular mechanics were further optimized with the Gaussian 09 program package[3] by means of density functional theory (DFT) using the Becke, 3-parameter, Lee–Yang–Parr (B3LYP)[4-5] exchange-correlation hybrid functional with 6-311++G(d,p) basis set[6], including the polarizable continuum model[7] in water. Further, harmonic vibrational frequencies were calculated to verify the stability of the optimized geometries. All the calculated vibrational frequencies were real (positive) indicating the true minimum of the calculated total potential energy of the optimized system. The computation was performed at the High Performance Computing center in Göttingen provided by GWDG.

The potential energy values for $D_{50}$ simulation were obtained by performing geometry optimizations at 6-311++G(d,p) IEFPCM (water) level of theory with additional input value eps= 2.1; 5.7; 11.0; 18.2; 26.6; 35.2; 44.2; 53.3; 62.4; 71.4; 80.4 to obtain 11 points against increasing dielectric constant. The calculations were performed for zwitterionic and spirolactone forms. Energy values were plotted against dielectric constant and the intersection point of zwitterion and spirolactone plots were considered as simulated $D_{50}$ value. The calculations were performed for the free **4-TMR-COOH** dye and for model **4-TMR-CONHMe** compound with truncated linker and ligand.

*Determination of Quantum Yields and Lifetimes*

The fluorescence quantum yields (absolute values) were obtained with a Quantaurus-QY absolute PL quantum yield spectrometer (model C11347-12, Hamamatsu) according to the manufacturer's instructions. Fluorescence lifetimes were measured with a Quantaurus-Tau fluorescence lifetime spectrometer (model C11367-32, Hamamatsu) according to the manufacturer's instructions.

*Plasmid construction*

Construction and characterization of pEBTet GW_SNAP plasmid was characterized previously [8-9]. This mammalian expression vector contain cytomegalovirus-type 2 tetracycline operator (tetO2)-tetO2 promoter which is "switched on" by addition of doxycycline. Histone H3 coding gene was PCR amplified from pmCherry-H3-23 template (Addgene, #55058), and cloned into pEBTet GW_SNAP vector using single tube BP/LR recombination protocol. Final construct sequence has been verified by sequencing.

*Maintenance and preparation of cells*

Human primary dermal fibroblasts were cultured in high-glucose DMEM (Thermo Fisher, #31053044) with 10% FBS (Thermo Fisher, #10082147) supplemented with of 1 mM Sodium pyruvate (Sigma, #S8636), 1% GlutaMax (Thermo Fisher, #35050038) and 1% Penicillin-Streptomycin (Sigma, #P0781) in a humidified 5% $CO_2$ incubator at 37 °C. The cells were split every 3-4 days or at confluence.

HeLa and Vero cells were cultured in high-glucose DMEM (Thermo Fisher, #31966047) with 10% FBS (BioSELL, #S0615) supplemented with 1% Penicillin-Streptomycin (Sigma, #P0781) in a humidified 5% $CO_2$ incubator at 37 °C. The cells were split every 3-4 days or at confluence.

U-2 OS cells were cultured in McCoys 5A medium (Thermo fisher #16600082) with 10% FBS (BioSELL, #S0615) supplemented with 1 mM Sodium pyruvate (Sigma, #S8636) and 1% of Penicillin-Streptomycin (Sigma #P0781) in a humidified 5% $CO_2$ incubator at 37 °C. U-2OS Nup96-SNAP expressing cells were cultivated under the same conditions, except growth medium was supplied with MEM Non-Essential Amino Acids Solution (Thermo fisher #11140050). The cells were split every 3-4 days or at confluence.

Human umbilical vein endothelial cells (HUVEC) cells were cultured in EMB medium (Lonza #CC-3129) and LVES (Thermo Fisher #C3008MP) supplemented with 1% of Penicillin-Streptomycin (Sigma #P0781) in a humidified 5% $CO_2$ incubator at 37 °C. Cells were split every 3-4 days or at confluence.

Cells were seeded in glass bottom 12-well (MatTek, #P12G-1.5-14-F) or 24-well (MatTek, #P24G-1.5-13-F) plates for wide-field imaging experiments on Biotek Lionheart FX automated microscope. Confocal and STED microscopy experiments were performed using µ-Slide 8 Well Glass Bottom dishes (Ibidi, #80827).

Inducible U-2 OS cell line expressing SNAP-tagged histone H3 was generated by transfecting cells with pEBTet H3-SNAP expression vector at ~70% confluence using Lipofectamine 2000 (Thermo Fisher Scientific, Cat. No. 11668027) following manufacturer’s recomendations. Transfected cells were cultivated in selection media composed of DMEM (Thermo Fisher Scientific, Cat. No. 31053-028) containing 10% FBS (Thermo Fisher Scientific, Cat. No. 10082139) and 1 µg/ml puromycin (Sigma, Cat. No. P9620) for approximately two weeks. Thereafter, selected cells were frozen in 10% DMSO and stored at -80°C. Expression of transgene was induced using 0.1 µg/ml doxycycline (Sigma, Cat. no. D9891) for 24 - 48 h before imaging experiment.

*Preparation and staining of mouse living primary cells and tissues*

*Animals*

All animal procedures were conducted in accordance with Directive 2010/63/EU of the European Parliament and the Council on the protection of animals used in research, as well as Animal Welfare Law of the Federal Republic of Germany (Tierschutzgesetz der Bundesrepublik Deutschland, TierSchG). All mice were housed with a 12 hours light/dark circadian cycle with *ad libitum* access to food and water.

*Primary neuronal isolation and staining*

The protocol for hippocampal neurons isolation has been described previously in [10]. Briefly, hippocampi were dissected from C57BL/6N mice of mixed gender at postnatal day P0-P1, treated with trypsin (0,25%; Gibco, cat. 15090046) for 25 minutes at 37°C and subsequently manually dissociated. Cells were plated on coverslips pre-coated with poly-L-ornithine (100 µg/ml; Sigma-Aldrich, cat. P3655) and laminin (1 µg/ml; BD Bioscience, cat. 354232) and cultivated in Neurobasal medium (Gibco, cat. 21103049) supplemented with B27 serum-free supplement (2%; Gibco, cat. 17504044), GlutaMax (1x; Gibco, cat. 35050061) and Penicillin-Streptomycin (100 units/ml and 100 µg/ml respectively; Gibco, cat. 15140122).

Living neurons were incubated in culture medium containing the indicated concentration of probe at 37°C for 30min and imaged without washing at room temperature thereafter.

*Tissue isolation and staining*

CD-1 mice of both genders were sedated with 3% isoflurane in a sealed container and quickly euthanized by cervical dislocation. The skull and the abdomen were opened, the organs were transferred to ice cold EGTA solution[11] and sliced manually using a scalpel blade. Slices were incubated in EGTA solution containing the indicated concentration of probe on ice for 30min and imaged without washing at room temperature thereafter.

*Cell cycle analysis by imaging flow cytometry and $EC_{50}$ determination*

The probes dissolved in DMSO (typically 500 - 2000× stock) were added into the media of cultured HeLa cells and incubated for 24 h at 37°C in humidified incubator with 5% $CO_2$. We used 6-well plates containing ~250,000 cells per well for this experiment. Afterwards, cells were processed according to the NucleoCounter® NC-3000™ two-step cell cycle analysis protocol for cells attached to T-flasks, cell culture plates or micro-carriers. We used NC-Slide A2™ slides (Chemometec, Cat. No. 942-0001) loaded with ~30 µl of the cell suspensions into a chamber of the slide. According to manufacturers recommendations ~10,000 cells in total were measured. The collected data were analyzed with ChemoMetec NucleoView NC-3000 software, version 2.1.25.8. The experiments were repeated three times on different days and the results are presented as means with standard deviations. The $EC_{50}$ values were determined by plotting percentage of subG1 phase cells and fitting the data in GraphPad Prism 8 to the following function:

$$Y = Y_{min} + (Y_{max} - Y_{min})/\left(1 + \left(\frac{EC_{50}}{X}\right)^{Hill}\right) \quad (2)$$

where $X$ – probe concentration, $Y_{min}$ – % subG1 in sample with DMSO (no probe added), $Y_{max}$ – the highest reachable % subG1, *Hill* - Hill coefficient determining the steepness of a dose-response curve, $EC_{50}$ - the concentration of probe that provoking halfway of subG1 cells in a population between the baseline ($Y_{min}$) and maximum response ($Y_{max}$).

*Wide field microscope and imaging parameters*

For wide-filed fluorescence microscopy images were aquired using Biotek Lionheart FX Automated Microscope equipped with Olympus dry 20× NA 0.45 objective and incubator set to +37°C. During acquisition cells were kept under atmosphere containing 5% of $CO_2$. The image acquisition parameters are listed below:

| Channel | LED | Filter cube | LED intensity | Integration time, ms | Camera gain |
|---|---|---|---|---|---|
| **Hoechst** | 365 nm | DAPI 377/447 | 5 | 10 | 30 |
| **GFP channel** | 465 nm | GFP 469/525 | 7 | 75 | 30 |
| **RFP channel** | 523 nm | RFP | 9 | 50 | 30 |
| **TxR channel** | 590 nm | Texas red | 9 | 50 | 30 |
| **Cy5 chennel** | 623 nm | Cy5 | 7 | 50 | 30 |

Parameters of image acquisition on a wide-field Lionheart FX Automated Microscope.

*Spinning disk confocal microscope and imaging parameters*

Spining disk confocal microscopy imaging was performed on an inverted microscope (Nikon Ti2) equipped with a 640 nm laser (Toptica, 150 mW) used for excitation and the oil-immersion objective Plan Apo Lambda 60x Oil NA 1.4 (MRD01605, Nikon). Emmsion light filtered using 665nm long pass filter. Images registered using back illuminated sCMOS camera Teledyne Photometrics Prime BSI with pixel size 6.5 x 6.5 µm corresponding to image pixel size 111 x 111 nm. All probes were used at 100 nM concentration. The image acquisition parameters are listed below:

| Probe | Laser power | Exposure (ms) |
|---|---|---|
| 4-625CP-(O4)Halo | 30% | 100 |
| 4-630CP-(O4)Halo | 30% | 100 |
| 4-640CP-(O4)Halo | 30% | 100 |
| 4-642CP-(O4)Halo | 30% | 100 |
| 4-SiR-(O4)Halo | 30% | 300 |
| 4-665SiR-(O4)Halo | 30% | 300 |
| 4-670SiR-(O4)Halo | 30% | 300 |
| 4-685SiR-(O4)Halo | 30% | 300 |
| 4-690SiR-(O4)Halo | 100% | 300 |
| 4-700SiR-(O4)Halo | 100% | 300 |
| 4-720SiR-(O4)Halo | 100% | 300 |

*Confocal/STED microscopes and imaging parameters*

Confocal and STED images were acquired using Abberior STED Expert Line, Abberior STED Falicilty Line scanning (Abberior Instruments GmbH) or TCS SP8 (Leica) microscopes. STED images were acquired using Abberior STED Expert Line or Abberior STED Falicilty Line scanning (Abberior Instruments GmbH) microscopes.

TCS SP8 confocal microscope equipped with 405, 458, 476, 488, 496, 514, 561 and 633 nm excitation lasers as well as HC PL APO CS2 63x/1.40 Oil objective (Leica) was used in described study. Microscope has three Hybrid and two PMT detectors which can be tuned to any detection window in the range 400 – 800 nm.Voxel size was 0.08µm x 0.08µm x 0.2µm (in xyz respectively) and pinhole size was set to 1.0 AU. Laser powers were optimized for each sample.

Abberior STED Expert Line equipped with 561 nm and 640 nm 40 MHz pulsed excitation lasers, a pulsed 775 nm 40 MHz 3W STED laser, and an UPlanSApo 100x/1.40 Oil objective. The following detection windows were used: for the TMR/580CP channel 615 / 20 nm, and for the 610CP/SiR channel 685 / 70 nm. Pixel size was 10 – 30 nm in the xy plane was used for 2D STED images. Laser powers were optimized for each sample. 3D STED images were acquired using pinhole set to 0.8 AU, voxel size set to 40 x 40 x 40 nm, 3D STED doughnut set to 95%, with single line accumulation and xzy scanning mode.

Abberior STED Facility Line equipped with 488, 515, 561, 640 and 700 nm 40 MHz pulsed excitation lasers, a pulsed 775 nm 40 MHz 3W STED laser, and an UPlanSApo 60x/1.42 Oil objective. Microscope has two APD and two MATRIX detectors which can be tuned to any detection window in the range 400 – 800 nm. Pixel size was 30 nm in the xy plane was used for 2D STED images and 80 nm in the xy plane for large field of view images. Laser powers were optimized for each sample.

Estimation of the STED effect was performed on Abberior STED Expert line microscope by varying the STED laser power from 0 to 100% while measuring cells stained with tubulin probes. Obtained data were fitted using GraphPad Prism 8 to the following equation:

$$Y = \frac{d_{conf}}{\sqrt{1+I/I_{sat}}} \quad (3),$$

where $d_{conf}$ – confocal resolution, $I$ – STED laser intensity power, $I_{sat}$ – saturating STED laser intensity power.

Settings used for confocal and STED imaging

| Figure | Probe | Concentration | Microscope | Excitation (nm) | Pixel dwell time (µs) | Objective | Pixel size (nm) | Emission (nm) |
|---|---|---|---|---|---|---|---|---|
| | **4-505R-CTX** | | | 488 | 2 | | | 505 – 608 |
| | **4-DMRh-CTX** | 100 nM | Abberior facility line | 518 | 2 | 60x NA 1.42 oil | 10 | 535 – 638 |
| | **4-525R-CTX** | | | 518 | 1 | | | 528 - 638 |
| | **4-TMR-CTX** | | | 561 | | | | 580 – 630 |
| | **4-DAllR-CTX** | | | 561 | | | | 580 – 630 |
| | **4-CFL-CTX** | | | 561 | | | | 580 – 630 |
| | **4-580CP-CTX** | | | 561 | | | | 580 – 630 |
| | **4-610CP-CTX** | | | 640 | | | | 650 – 720 |
| | **4-625CP-CTX** | | | 640 | | | | 650 – 720 |
| Figure 3a and S20 | **4-630CP-CTX** | 100 nM | Abberior expert line | 640 | 1 | 100x NA 1.4 oil | 10 | 650 – 720 |
| | **4-640CP-CTX** | | | 640 | | | | 650 – 720 |
| | **4-642CP-CTX** | | | 640 | | | | 650 – 720 |
| | **4-645CP-CTX** | | | 640 | | | | 650 – 720 |
| | **4-SiR-CTX** | | | 640 | | | | 650 – 720 |
| | **4-665SiR-CTX** | | | 640 | | | | 650 – 720 |
| | **4-670SiR-CTX** | | | 640 | | | | 650 – 720 |
| | **4-685SiR-CTX** | 100 nM | Abberior facility line | 705 | 2 | 60x NA 1.42 oil | 10 | 725 – 780 |
| | **4-700SiR-CTX** | | | 705 | | | | 725 - 780 |
| Figure 3c | **4-625CP-CTX** | 10 nM | Abberior expert line | 640 | 3 | 100x NA 1.4 oil | 20 | 650 - 720 |
| Figure 4a | **4-580CP-CTX** | 100 nM | Abberior expert line | 561 | 2 | 100x NA 1.4 oil | 30 | 580 – 630 |
| | **4-630CP**-(O4)Halo | 100 nM | | 640 | | | | 650 - 720 |
| Figure 4b | **4-580CP-Hoechst** | 100 nM | Abberior expert line | 561 | 2 | 100x NA 1.4 oil | 20 | 580 – 630 |
| | **4-625CP-BG** | 100 nM | | 640 | | | | 650 - 720 |
| | **4-DAllR-CTX** | 10 nM | | 561 | | | | 571 – 630 |
| Figure 4c | **4-630CP-JAS** | 10 nM | Abberior facility line | 640 | 3 | 60x NA 1.42 oil | 30 | 650 - 695 |
| | **4-700SiR-TPP** | 10 nM | | 705 | | | | 715 - 755 |
| Figure 4d-f | **4-505R-Hoechst** | 250 nM | TCS SP8 confocal microscope | 514 | 1.2 | 63x NA 1.4 oil | 80x80x20 | 525-550 |
| | **4-TMR-JAS** | 1 µM | | 561 | | | | 570-600 |
| | **4-685SiR-TPP** | 1.5 µM | | 630 | | | | 689-740 |
| Figure S12 | **5-580CP-Hoechst** | 100 nM | Abberior expert line | 561 | 3 | 100x NA 1.4 oil | 30 | 580 – 630 |
| | **SiR-SNAP** | 100 nM | | 640 | | | | 650 - 720 |
| Figure S14 | **4-580CP-BG** | 100 nM | Abberior expert line | 561 | 2 | 100x NA 1.4 oil | 15 | 580 - 630 |
| | **4-610CP-BG** | 100 nM | Abberior expert line | 640 | 2 | 100x NA 1.4 oil | 15 | 650 - 720 |
| | **4-625CP-BG** | 100 nM | Abberior expert line | 640 | 2 | 100x NA 1.4 oil | 15 | 650 - 720 |
| | **4-630CP-BG** | 100 nM | Abberior expert line | 640 | 2 | 100x NA 1.4 oil | 15 | 650 - 720 |
| | **4-640CP-BG** | 100 nM | Abberior expert line | 640 | 2 | 100x NA 1.4 oil | 15 | 650 - 720 |
| | **SiR-SNAP** | 100 nM | Abberior expert line | 640 | 2 | 100x NA 1.4 oil | 15 | 650 - 720 |
| Figure S17 | **4-630CP-CTX** | 10 nM | Abberior expert line | 640 | 1 | 100x NA 1.4 oil | 40 | 650 - 720 |
| | **4-630CP-CTX** | 100 nM | Abberior expert line | 640 | 1 | 100x NA 1.4 oil | 40 | 650 - 720 |
| Figure S21 | **4-610CP-JAS** | 10 nM | Abberior facility line | 640 | 3 | 60x NA 1.42 oil | 80 | 650 - 760 |
| | **4-610CP-JAS** | 100 nM | Abberior facility line | 640 | 3 | 60x NA 1.42 oil | 80 | 650 - 760 |
| | **4-610CP-JAS** | 1000 nM | Abberior facility line | 640 | 3 | 60x NA 1.42 oil | 80 | 650 - 760 |

| | | | | | | | | |
|---|---|---|---|---|---|---|---|---|
| | **4-610CP-C6-JAS** | 10 nM | Abberior facility line | 640 | 3 | 60x NA 1.42 oil | 80 | 650 - 760 |
| | **4-610CP-C6-JAS** | 100 nM | Abberior facility line | 640 | 3 | 60x NA 1.42 oil | 80 | 650 - 760 |
| | **4-610CP-C6-JAS** | 1000 nM | Abberior facility line | 640 | 3 | 60x NA 1.42 oil | 80 | 650 - 760 |
| Figure S22 | **4-TMR-JAS** | 100 nM | Abberior facility line | 561 | 3 | 60x NA 1.42 oil | 80 | 571 - 681 |
| | **4-685SiR-JAS** | 100 nM | Abberior facility line | 705 | 3 | 60x NA 1.42 oil | 80 | 715 - 800 |
| | **4-630CP-JAS** | 100 nM | Abberior facility line | 640 | 3 | 60x NA 1.42 oil | 80 | 650 -760 |
| | **4-505R-JAS** | 100 nM | Abberior facility line | 488 | 3 | 60x NA 1.42 oil | 80 | 498 - 608 |
| Figure S23 | **4-505R-Hoe** | 100 nM | Abberior facility line | 488 | 3 | 60x NA 1.42 oil | 80 | 500 - 550 |
| | **4-525R-Hoe** | 100 nM | Abberior facility line | 518 | 3 | 60x NA 1.42 oil | 80 | 530 - 580 |
| | **4-CFL-Hoe** | 100 nM | Abberior facility line | 561 | 3 | 60x NA 1.42 oil | 80 | 575 - 625 |
| | **4-625CP-Hoe** | 100 nM | Abberior facility line | 640 | 3 | 60x NA 1.42 oil | 80 | 650 - 700 |
| | **4-630CP-Hoe** | 100 nM | Abberior facility line | 640 | 3 | 60x NA 1.42 oil | 80 | 651 - 700 |
| | **4-685SiR-Hoe** | 100 nM | Abberior facility line | 705 | 3 | 60x NA 1.42 oil | 80 | 715 - 765 |
| Figure S24 | **4-625CP-TPP** | 100 nM | Abberior facility line | 640 | 3 | 60x NA 1.42 oil | 80 | 650 - 760 |
| | **4-640CP-TPP** | 100 nM | Abberior facility line | 640 | 3 | 60x NA 1.42 oil | 80 | 650 - 760 |
| | **4-665SiR-TPP** | 100 nM | Abberior facility line | 640 | 3 | 60x NA 1.42 oil | 80 | 650 - 760 |
| | **4-685SiR-TPP** | 100 nM | Abberior facility line | 640 | 3 | 60x NA 1.42 oil | 80 | 650 - 760 |
| | **4-525R-TPP** | 100 nM | Abberior facility line | 518 | 3 | 60x NA 1.42 oil | 80 | 531 - 630 |
| | **4-685SiR-TPP** | 100 nM | Abberior facility line | 705 | 3 | 60x NA 1.42 oil | 80 | 715 - 800 |
| | **4-700SiR-TPP** | 100 nM | Abberior facility line | 705 | 3 | 60x NA 1.42 oil | 80 | 715 - 800 |
| Figure S25 | **4-580R-PepA** | 100 nM | Abberior facility line | 561 | 5 | 60x NA 1.42 oil | 80 | 571- 681 |
| | **4-630CP-PepA** | 100 nM | Abberior facility line | 640 | 3 | 60x NA 1.42 oil | 80 | 650 - 760 |
| | **4-642CP-PepA** | 100 nM | Abberior facility line | 640 | 3 | 60x NA 1.42 oil | 80 | 650 - 760 |
| | **4-SiR-PepA** | 100 nM | Abberior facility line | 640 | 3 | 60x NA 1.42 oil | 80 | 650 - 760 |
| | **4-685SiR-PepA** | 100 nM | Abberior facility line | 705 | 3 | 60x NA 1.42 oil | 80 | 715 - 800 |
| | **4-700SiR-PepA** | 100 nM | Abberior facility line | 705 | 3 | 60x NA 1.42 oil | 80 | 715 - 800 |

*4-color time-lapse movies*

For recording multicolor movies, the cells were incubated with the indicated probes for 1h at 37°C and imaged without washing at 37°C with 5% $CO_2$ flow. In case of U-2 OS vimentin-HaloTag cell line, the cells were first incubated with **4-642CP-($O_4$)Halo** substrate for 1h at 37°C, washed twice, stained with the other probes and imaged without washing.

The time-lapse series were recorded on the Abberior Facility Line microscope in the confocal imaging mode under control of Imspector software. The 2-step sequential line scanning was used: 561 nm and 700 nm channels were recorded in the first step and 488 nm and 640 nm channels were recorded during the second step. Pixel size was 100 nm, pixel dwell time – 1 µs, frame rate – 0.33 fps. 200 frames of 50×50 µm were acquired for each movie and the final videos were rendered at 20 fps.

*Processing and visualization of acquired images*

All acquired or reconstructed images were processed and visualized using Fiji[12]. Line profiles were measured using the “straight line” tool with the line width set to 3 pixels. To define microtubule diameter, line profiles were fitted with Gaussian and Lorentzian distributions for confocal and STED images respectively. For 3D STED images resolution, perpendicular microtubule profile images in xz and zy planes were cropped using Fiji BigDataViewer Plugin and fitted using 2D Gaussian distribution. Fitting routines were optimized and performed with Python3 language. Stitching and deconvolution of images were performed using SVI Huygens Essential software package.

### General chemical experimental information and synthesis methods

NMR spectra were recorded at 25 °C with an Agilent 400-MR spectrometer at 400.06 MHz (1H) and 100.60 MHz ($^{13}C$) and Varian INOVA 600 (I600) spectrometer at 599.74 MHz ($^{1}H$) and are reported in ppm. All $^{1}H$ and $^{13}C$ spectra are referenced to tetramethylsilane (δ = 0 ppm) using the residual signals of the solvents according to the values reported in literature[13]. Multiplicities of signals are described as follows: s = singlet, d = doublet, t = triplet, q = quartet, p = pentet, m = multiplet or overlap of non-equivalent resonances; br = broad signal. Coupling constants (J) are given in Hz.

ESI-MS were recorded on a Varian 500-MS spectrometer (Agilent). ESI-HRMS were recorded on a MICROTOF spectrometer (Bruker) equipped with ESI ion source (Apollo) and direct injector with LC autosampler Agilent RR 1200.

Analytical LC–MS analysis was performed on an Agilent 1260 Infinity II LC/MS system equipped with an autosampler, diode array detector WR, fluorescence detector Spectra and Infinity Lab LC/MSD 6100 series quadruple with API electrospray. Analysis was done by using an Agilent Zorbax SB-C18 RRHT, 2.1 x 50 mm, 1.8 µm threaded column and SUPELCO Titan C18, 2.1 x 75 mm, 1.9 µm column with A: 25 mM $HCOONH_4$ (pH = 3.6) aqueous buffer and B: MeOH

Preparative HPLC was performed on a combined Agilent1260/1290 Infinity II preparative system equipped with an 1290 Infinity II open-bed sampler (G7169B)/fraction collector (G7159B), 1260 Infinity II preparative binary pump (G7161A), 1260 Infinity II multiple wavelength detector (G7165A) and with Agilent 5 Prep-C18, 5 µm, 100 x 50 mm preparative column.

**9,9'-methylenebis(2,3,6,7-tetrahydro-1H,5H-pyrido[3,2,1-ij]quinolin-8-ol)** (**SI-1**):

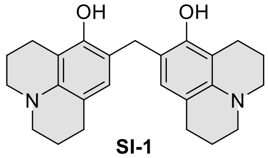

**SI-1**

To the ice bath cooled solution of 8-hydroxyjulolidine (1 g, 5.3 mmol) in methanol (10 mL) hydrochloric acid was added ( 0.5 mL, 32%) and stirred for 10 min. Formalin (1 mL, 16%) was then added to the reaction mixture and the resulting mixture was allowed to stand overnight. The mixture was poured into water (40 mL) and was neutralized with saturated $Na_2CO_3$ solution. The mixture was extracted with DCM (3x 50 mL), the combined extracts were dried over $Na_2SO_4$, filtered and solvent was removed under reduced pressure. The product was purified by flash column chromatography (Teledyne Isco RediSep Rf 40 g; gradient 2% to 40% hexane – EtOAc), fractions containing the product were evaporated to give 0.7g (68%) of off-white solid. And was immediately used in the next step. The obtained compound S

***2,3,6,7,12,13,16,17*-octahydro-1*H*,5*H*,9*H*,11*H*,15*H*-Xantheno[2,3,4-ij:5,6,7-i'j']diquinolizine (SI-2)**:

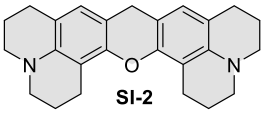

**SI-2**

Compound **SI-1** (0.7g, 1.8 mmol) was added to concentrated sulphuric acid (5 mL). The resulting solution was heated at 95$^o$C for 3 hours. The reaction was allowed to cool to room temperature before being poured onto ice. The pH was adjusted to 5 with 40% NaOH solution whilst keeping the mixture cold. Then mixture was extracted with DCM (3x 100 mL) the combined extracts were dried over $Na_2SO_4$, filtered and the solvent was removed under reduced pressure. The product was purified by flash column chromatography (Teledyne Isco RediSep Rf 40 g; gradient 2% to 40% hexane – EtOAc), fractions containing the product were evaporated to give 0.435g (65%) of off-white solid.

$^1$H NMR (400 MHz, $CDCl_3$) δ 6.69 (s, 2H), 6.08 (s, 2H), 3.63 (s, 2H), 3.09 – 2.99 (m, 8H), 2.69 (t, *J* = 6.6 Hz, 4H), 2.61 (t, *J* = 6.6 Hz, 4H), 2.02 – 1.93 (m, 8H).

$^{13}$C NMR (101 MHz, $CDCl_3$) δ 149.3, 127.8, 50.5, 49.8, 30.8, 26.8, 22.2, 21.5, 21.2. (not all aromatic peaks were registered due to partial protonation and dynamics related to it in CDCl3)

**6-Bromoindoline** (**SI-3**):

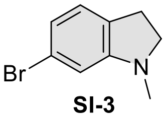

**SI-3**

6-Bromoindole (3.5 g, 17.8 mmol) was dissolved in glacial acetic acid (50 mL) and $NaCNBH_3$ was added carefully in small portions (intense bubbling). Once the addition was complete, paraformaldehyde (5.3 g, 178 mmol) was added and the reaction was stirred for 3-5 hours. The reaction course was monitored by TLC hex:EtOAce (8:2). Then the reaction mixture was poured into water (200 mL) and neutralized with saturated $Na_2CO_3$ solution. The mixture was extracted with DCM (3x 100 mL), combined extracts were dried with $Na_2SO_4$, filtered and solvent was removed under reduced pressure. The product was purified by flash column chromatography (Teledyne Isco RediSep Rf 80 g; gradient 2% to 40% hexane – EtOAc), fractions containing the product were evaporated to give 2.6 g (69%) of colorless oil.

$^{1}$H NMR (400 MHz, $CDCl_3$) δ 6.90 (dt, *J* = 7.7, 1.1 Hz, 1H), 6.75 (dd, *J* = 7.7, 1.7 Hz, 1H), 6.56 (d, *J* = 1.8 Hz, 1H), 3.33 (t, *J* = 8.3 Hz, 2H), 2.89 (td, *J* = 8.2, 1.1 Hz, 2H), 2.74 (s, 3H).

$^{13}$C NMR (101 MHz, $CDCl_3$) δ 154.9, 129.4, 125.4, 121.2, 120.2, 110.1, 56.2, 35.8, 28.4.

ESI-MS, positive mode: m/z = 212.0 $[M+H]^+$.

HRMS (ESI) calcd for $C_9H_{11}BrN$ $[M+H]^+$ 212.0069, found 212.0073.

**1-methyl-6-(prop-1-en-2-yl)indoline** (**SI-4**):

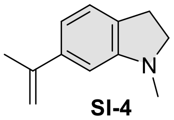

**SI-4**

A mixture of **SI-3** (2.6 g, 12.3 mmol), potassium isopropenyltrifluoroborate (2.2 g, 14.7 mmol, 1.2 equiv) and Pd(dppf)$Cl_2$ (301 mg, 0.37 mmol, 3 mol %) in 1,4-dioxane (45 mL) in a 100 mL round bottom flask was purged on a Schlenk line and filled with argon. Aqueous NaOH (5 mL of 2 M solution) was then injected, and the yellowish solution turned brown. The mixture was then heated up to 100 °C and stirred for 3 h, cooled down to rt, diluted with water (200 mL) and extracted with DCM (3×100 mL). The combined organic layers were washed with brine and dried over $Na_2SO_4$. The product was isolated by flash column chromatography (Teledyne Isco RediSep Rf 80 g; gradient 2% to 20% EtOAc – hexane, Teledyne Isco RediSep Rf 40 g, gradient 10% to 70% hexane – DCM) yellowish oil, yield 1.4 g (66%).

$^{1}$H NMR (400 MHz, $CDCl_3$) δ 7.13 – 7.01 (m, 1H), 6.82 (dd, *J* = 7.5, 1.6 Hz, 1H), 6.62 (d, *J* = 1.5 Hz, 1H), 5.38 – 5.30 (m, 1H), 5.05 (p, *J* = 1.6 Hz, 1H), 3.34 (t, *J* = 8.2 Hz, 2H), 2.96 (t, *J* = 8.2 Hz, 2H), 2.81 (s, 3H), 2.20 – 2.12 (m, 3H).

$^{13}$C NMR (101 MHz, $CDCl_3$) δ 144.2, 141.2, 130.1, 124.0, 115.9, 111.7, 104.8, 56.5, 36.4, 28.6, 22.3.

ESI-MS, positive mode: m/z = 174.1 $[M+H]^+$.

HRMS (ESI) calcd for $C_{12}H_{16}N$ $[M+H]^+$ 174.1277, found 174.1278.

**1-methylindoline** (**SI-5**):

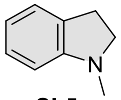

**SI-5**

The 1-methylindoline was prepared according to previously published procedure[14].

**1-methylindoline-5-carbaldehyde** (**SI-6**):

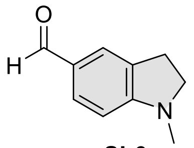

**SI-6**

The 1-methylindoline-5-carbaldehyde (**SI-6**) was prepared according to previously published procedure[15].

**(1-methylindolin-5-yl)methanol** (**SI-7**):

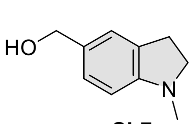

**SI-7**

1-methylindoline-5-carbaldehyde (**SI-6**) (6.2g, 38.5 mmol) was dissolved in MeOH (40 mL) and solution was cooled in an ice bath. $NaBH_4$ (1.3 g, 34.2 mmol) was added in small portions. After addition was complete, reaction mixture was left to warm to room temperature. After 1h of stirring at rt water (5 mL) was added and stirring was continued for 10 min. Then MeOH was removed on rotary evaporator, water was added to the residue and mixture was extracted with EtoAc (3x 100 mL), washed with brine and dried over $Na_2SO_4$. The solvent was evaporated and the product was purified by flash column chromatography (Teledyne Isco RediSep Rf 80 g; gradient 20% to 80% hexane – EtOAc). White solid, yield 5.7 g (91%).

$^1$H NMR (400 MHz, $d_6$-DMSO) δ 7.00 (s, 1H), 6.94 (d, *J* = 7.9 Hz, 1H), 6.43 (d, *J* = 7.9 Hz, 1H), 4.87 – 4.80 (m, 1H), 4.33 (d, *J* = 5.6 Hz, 2H), 3.19 (t, *J* = 8.2 Hz, 2H), 2.83 (t, *J* = 8.2 Hz, 2H), 2.66 (s, 3H).

$^{13}$C NMR (101 MHz, $d_6$-DMSO) δ 152.4, 131.6, 129.8, 126.0, 123.3, 106.5, 63.3, 55.9, 36.1, 28.2.

**2-(3-(dimethylamino)phenyl)propan-2-ol** (**SI-8**):

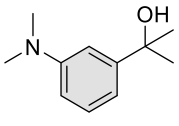

**SI-8**

Solution of 3-Bromo-dimethylaniline (4g, 20 mmol) in THF (50 mL) was cooled to -78°C in dry ice/acetone cooling bath. Then *n*-BuLi (8 mL, 2.5M) was added through a syringe and mixture was stirred for 30 min at -78°C followed by the addition of dry acetone (2 mL, 27 mmol). After addition was complete the reaction mixture was stirred at -78° for 15 min and then cooling bath was removed and the mixture was allowed to warm to rt. Stirring was continued for 30 min at rt. Then HCl (2mL, 1M) was added to the reaction mixture through a syringe. THF was partially evaporated, water was added and the mixture was extracted with EtOAc (4x 50 mL), washed with brine and dried over $Na_2SO_4$. The solvent was evaporated and the product was purified by flash column chromatography (Teledyne Isco RediSep Rf 80 g; gradient 10% to 70% hexane – EtOAc) to give 2.88 g of yellowish liquid in 80% yield.

$^1$H NMR (400 MHz, $CDCl_3$) δ 7.23 (t, *J* = 7.9 Hz, 1H), 6.97 (t, *J* = 2.2 Hz, 1H), 6.83 (d, *J* = 7.7 Hz, 1H), 6.68 (dd, *J* = 8.2, 2.3 Hz, 1H), 2.97 (s, 6H), 1.59 (s, 6H).

$^{13}$C NMR (101 MHz, $CDCl_3$) δ 150.5, 150.4, 129.0, 113.4, 111.4, 109.1, 72.9, 41.0, 31.9.

**N,N-dimethyl-3-(prop-1-en-2-yl)aniline** (**SI-9**):

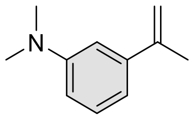

**SI-9**

Compound **SI-8** (2.88 g, 16.1 mmol) was dissolved in acetic acid (30 mL) and 1 mL of concentrated $H_2SO_4$ was added. The obtained mixture was heated to 100°C for 1h. Reaction course was monitored by TLC hex:EtOAce (8:2). Once reaction was complete mixture was poured to water and neutralized with saturated $Na_2CO_3$. Mixture was extracted with EtOAc (4x 100mL), washed with brine and dried over $Na_2SO_4$. The solvent was evaporated and product was purified by flash column chromatography (Teledyne Isco RediSep Rf 80 g; gradient 2% to 20% hexane – EtOAc) to give 2.0 g of light yellow liquid in a 77% yield.

$^1$H NMR (400 MHz, $CDCl_3$) δ 7.26 (t, *J* = 8.1 Hz, 1H), 6.95 – 6.86 (m, 2H), 6.77 – 6.71 (m, 1H), 5.39 (dt, *J* = 1.5, 0.8 Hz, 1H), 5.11 (p, *J* = 1.5 Hz, 1H), 3.02 (s, 6H), 2.21 (dd, *J* = 1.5, 0.8 Hz, 3H).

$^{13}$C NMR (101 MHz, $CDCl_3$) δ 150.7, 144.4, 142.5, 129.0, 114.6, 112.2, 112.2, 110.3, 40.9, 22.2.

**2,3,6,7-tetrahydro-1H,5H-pyrido[3,2,1-ij]quinoline-9-carbaldehyde** (**SI-10**):

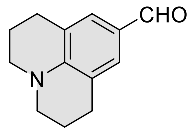

**SI-10**

Phosphorous oxychloride (2.14 mL, 22.9 mmol) was added dropwise to anhydrous DMF (6 mL) whilst mixture was cooled in an ice bath. The obtained mixture was stirred at room temperature for 30 minutes. Solution of julolidine (3.6 g, 20.8 mmol) in DMF (5 mL) was added and the reaction mixture was heated at 90°C for 4 hours. Then mixture was cooled and poured to 100 mL water and neutralized with saturated $Na_2CO_3$ solution. Obtained mixture was extracted with EtOAc (3x 100mL), washed with brine, dried over $Na_2SO_4$ and filtered. The solvent was evaporated and product was purified by flash column chromatography (Teledyne Isco RediSep Rf 40 g; gradient 10% to 70% hexane – EtOAc) to give 2.7 g of yellow solid in a 65% yield.

$^1$H NMR (400 MHz, *d$_6$*-DMSO) δ 9.50 (s, 1H), 7.21 (s, 2H), 3.27 (t, *J* = 5.7 Hz, 4H), 2.69 (t, *J* = 6.3 Hz, 4H), 1.89 – 1.82 (m, 4H).

$^{13}$C NMR (101 MHz, *d$_6$*-DMSO) δ 189.2, 147.5, 128.8, 123.3, 119.9, 49.3, 27.0, 20.7.

**(2,3,6,7-tetrahydro-1H,5H-pyrido[3,2,1-ij]quinolin-9-yl)methanol** (**SI-11**):

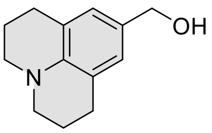

**SI-11**

Carbaldehyde **SI-10** (1.2 g, 5.97 mmol) was dissolved in 30 mL of MeOH and $NaBH_4$ (247 mg, 6.5 mmol) was added in small portions at room temperature over the course of 30 minutes. Once addition was complete the mixture was stirred for 1 hour. Reaction was quenched by addition of water (5 mL) and MeOH was evaporated on rotary evaporator. Water was poured on residue and the mixture was extracted with EtOAc (3x 50 mL), washed with brined, dried over $Na_2SO_4$ and filtered. The solvent was evaporated and product was purified by flash column chromatography (Teledyne Isco RediSep Rf 40 g; gradient 10% to 70% hexane – EtOAc) to give 0.93 g of white solid in a 77% yield. The compound easily oxidizes and we recommend to store it under inert gas at -15°C.

$^{1}$H NMR (400 MHz, $d_6$-DMSO) δ 6.63 (s, 2H), 4.72 (t, *J* = 5.6 Hz, 1H), 4.23 (d, *J* = 5.6 Hz, 2H), 3.05 (t, *J* = 5.6 Hz, 4H), 2.65 (t, *J* = 6.5 Hz, 4H), 1.89 – 1.83 (m, 4H).

$^{13}$C NMR (101 MHz, $d_6$-DMSO) δ 141.6, 129.3, 125.6, 120.6, 63.1, 49.4, 27.2, 21.8.

**8-bromo-2,3,6,7-tetrahydro-1H,5H-pyrido[3,2,1-ij]quinolone** (**SI-12**):

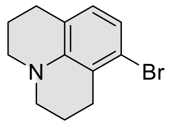

**SI-12**

3-Bromoaniline (6.3 mL, 58 mmol), $Na_2CO_3$ (24.6 g, 232 mmol) and 1-bromo-3-chloropropane (110 mL, 1.11 mol) were heated with stirring at 140°C for 48 hours. Then mixture was cooled, water (250 mL) was added and products were extracted with DCM (3x 150 mL). Combine organic extracts were dried over $Na_2SO_4$ and filtered. Filtrate was concentrated and excess of 1-bromo-3-chloropropane was removed. The crude mixture was dissolved in DMF (30 mL) and heated at 160°C for 48 hours. After cooling to room temperature solution of NaOH was added (200 mL, 1M) and mixture was extracted with EtOAc (3x 250 mL). The combined extracts were dried over $Na_2SO_4$, filtered and solvent was removed on rotary evaporator. The product was purified by flash column chromatography (Teledyne Isco RediSep Rf 80 g; gradient 1% to 10% hexane – EtOAc) to give 4.6 g of slightly yellow solid in a 31% yield.

$^{1}$H NMR (400 MHz, $d_6$-DMSO) δ 6.69 – 6.60 (m, 2H), 3.12 – 3.06 (m, 4H), 2.63 (dt, *J* = 11.7, 6.8 Hz, 4H), 1.91 – 1.81 (m, 4H).

$^{13}$C NMR (101 MHz, $d_6$-DMSO) δ 144.5, 127.8, 122.0, 120.4, 119.5, 118.5, 49.2, 48.8, 28.0, 27.0, 21.3, 21.2.

**8-(prop-1-en-2-yl)-2,3,6,7-tetrahydro-1H,5H-pyrido[3,2,1-ij]quinoline** (**SI-13**):

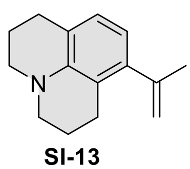

**SI-13**

A mixture of **SI-12** (2.5 g, 9.9 mmol), potassium isopropenyltrifluoroborate (1.75 g, 11.9 mmol, 1.2 equiv) and Pd(dppf)$Cl_2$ (245 mg, 0.3 mmol, 3 mol %) in 1,4-dioxane (45 mL) in a 100 mL round bottom flask was purged on a Schlenk line and filled with argon. Aqueous NaOH (5 mL of 2 M solution) was then injected, and the yellowish solution turned brown. The mixture was then heated up to 100 °C and stirred for 3 h, cooled down to rt, diluted with water (200 mL) and extracted with DCM (3× 100 mL). The combined organic layers were washed with brine and dried over $Na_2SO_4$. The product was isolated by flash column chromatography (Teledyne Isco RediSep Rf 80 g; gradient 1% to 10% EtOAc – hexane, Teledyne Isco RediSep Rf 40 g, gradient 10% to 70% hexane – DCM) resulting in light yellow oil, yield 1.7 g (82%).

$^1H$ NMR (400 MHz, $d_6$-DMSO) δ 6.64 (d, *J* = 7.5 Hz, 1H), 6.21 (d, *J* = 7.5 Hz, 1H), 5.09 (dq, *J* = 3.0, 1.5 Hz, 1H), 4.69 (dq, *J* = 2.6, 0.8 Hz, 1H), 3.08 (q, *J* = 5.8 Hz, 4H), 2.61 (dt, *J* = 14.5, 6.8 Hz, 4H), 1.91 (dd, *J* = 1.5, 0.9 Hz, 3H), 1.88 – 1.79 (m, 4H).

$^{13}C$ NMR (101 MHz, $d_6$-DMSO) δ 146.4, 142.9, 141.6, 126.5, 120.1, 117.8, 115.3, 114.2, 50.1, 49.7, 27.8, 25.4, 24.8, 22.1.

ESI-MS, positive mode: m/z = 214.2 [M+H]+.

HRMS (ESI) calcd for $C_{16}H_{19}N$ $[M+H]^+$ 214.1590, found 214.1592.

**7-bromo-2,2,4-trimethyl-1,2-dihydroquinoline** (**SI-14**):

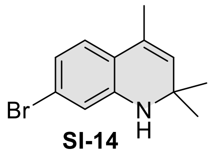

SI-14

3-Bromoaniline (8.65 mL, 80 mmol), mesityl oxide (1.3 mL, 160 mmol), $I_2$ (254 mg, 1 mmol) was dissolved in toluene (40 mL) and refluxed with stirring for 8h. Then another portion of $I_2$ (254 mg, 1 mmol) was added and reflux was continued for another 16h. Reaction course was monitored by TLC (hexane:EtOAc 9:1). Once reaction was complete the solvent and the excess of mesityl oxide was evaporated and the product was purified by flash column chromatography (Teledyne Isco RediSep Rf 120 g; gradient 1% to 10% EtOAc – hexane) to give 8.5 g of yellow oil in 42% yield.

$^1H$ NMR (400 MHz, $d_6$-DMSO) δ 6.83 (d, *J* = 7.9 Hz, 1H), 6.59 (d, *J* = 2.0 Hz, 1H), 6.54 (dd, *J* = 8.1, 2.0 Hz, 1H), 6.11 (s, 1H), 5.33 – 5.27 (m, 1H), 1.86 (d, *J* = 1.5 Hz, 3H), 1.19 (s, 6H).

$^{13}C$ NMR (101 MHz, $d_6$-DMSO) δ 145.7, 128.6, 126.8, 124.9, 120.9, 119.2, 117.4, 114.0, 51.4, 31.0, 18.1.

**7-bromo-1,2,2,4-tetramethyl-1,2-dihydroquinoline** (**SI-15**):

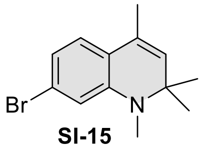

SI-15

A mixture of **SI-14** (5g, 19.8 mmol) $CH_3I$ (2.5 mL, 40 mmol) and $K_2CO_3$ (5.5g, 40 mmol) in MeCN (50mL) was stirred at 50°C overnight. Reaction course was monitored by TLC (hexane:EtOAc 95:5). Reaction Mixture was cooled and filtered. Filtrate was collected and evaporated, product was purified by flash column chromatography (Teledyne Isco RediSep Rf 120 g; gradient 0% to 10% EtOAc – hexane) to give 3.6g of yellow oil in 68% yield.

$^1H$ NMR (400 MHz, $CD_2Cl_2$) δ 6.87 (d, *J* = 8.0 Hz, 1H), 6.71 (dd, *J* = 8.0, 1.9 Hz, 1H), 6.60 (d, *J* = 1.9 Hz, 1H), 5.31 (q, *J* = 1.4 Hz, 2H), 2.76 (s, 3H), 1.94 (d, *J* = 1.4 Hz, 3H), 1.29 (s, 6H).

$^{13}C$ NMR (101 MHz, $CD_2Cl_2$) δ 146.5, 130.2, 127.3, 124.2, 122.2, 122.0, 118.3, 113.0, 56.5, 30.6, 27.1, 18.1.

**1,2,2,4-tetramethyl-7-(prop-1-en-2-yl)-1,2-dihydroquinoline** (**SI-16**):

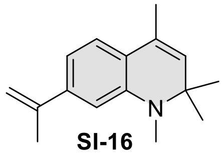

**SI-16**

A mixture of **SI-15** (1.5 g, 5.6 mmol), potassium isopropenyltrifluoroborate (1.0 g, 6.8 mmol, 1.2 eq) and Pd(dppf)$Cl_2$ (137 mg, 0.17 mmol, 3 mol %) in 1,4-dioxane (45 mL) in a 100 mL round bottom flask was purged on a Schlenk line and filled with argon. Aqueous NaOH (5 mL of 2 M solution) was then injected, and the yellowish solution turned brown. The mixture was then heated up to 100°C and stirred for 5 h, cooled down to rt, diluted with water (200 mL) and extracted with DCM (3× 100 mL). The combined organic layers were washed with brine and dried over $Na_2SO_4$. The product was isolated by flash column chromatography (Teledyne Isco RediSep Rf 80 g; gradient 1% to 10% EtOAc – hexane, Teledyne Isco RediSep Rf 80 g, gradient 1% to 10% hexane – DCM) to give 0.84 g of light yellow oil in a 66% yield.

$^1$H NMR (400 MHz, $CD_2Cl_2$) δ 7.00 (d, *J* = 7.8 Hz, 1H), 6.75 (dd, *J* = 7.8, 1.8 Hz, 1H), 6.61 (d, *J* = 1.8 Hz, 1H), 5.35 (dd, *J* = 1.7, 0.8 Hz, 1H), 5.31 (q, *J* = 1.6 Hz, 1H), 5.03 (p, *J* = 1.5 Hz, 1H), 2.82 (s, 3H), 2.14 (dd, *J* = 1.5, 0.8 Hz, 3H), 1.97 (d, *J* = 1.4 Hz, 4H), 1.29 (s, 6H).

$^{13}$C NMR (101 MHz, $CD_2Cl_2$) δ 145.1, 144.1, 141.3, 130.1, 127.8, 122.8, 122.7, 113.5, 111.1, 107.7, 56.1, 30.4, 26.8, 21.6, 18.2.

ESI-MS, positive mode: m/z = 228.2 [M+H]+.

HRMS (ESI) calcd for $C_{16}H_{22}N$ $[M+H]^+$ 228.1747, found 228.1749.

**Bis(6-bromo-1-methylindolin-5-yl)methane** (**SI-17**):

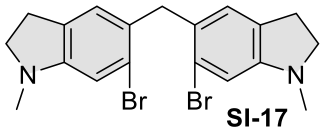

**SI-17**

Compound **SI-3** (2.1 g, 10 mmol) was dissolved in 20 mL of glacial acetic acid then formaldehyde solution was added (4.5 mL, 37%). Reaction mixture was heated to 40°C for 15-20 min. The color of mixture changed from colorless to green/yellow. Reaction completion was confirmed by TLC (Hexane:EtOAc 9:1). Reaction mixture was cooled, poured into water and neutralized with NaOH solution. Mixture was extracted with DCM (3x 50 mL), washed with brine, dried over $Na_2SO_4$ and filtered. Filtrate was concentrated on rotary evaporator, the mixture was deposited on celite and product was purified by flash column chromatography (Büchi Reveleris HP silica 40 g, gradient 5% to50% hexane – EtOAc) to give 1.35 g of white solid in a 62% yield.

$^1$H NMR (400 MHz, *d*$_6$-DMSO) δ 6.70 (s, 1H), 6.68 (s, 1H), 3.84 (s, 1H), 3.24 (t, *J* = 8.2 Hz, 2H), 2.76 (t, *J* = 8.2 Hz, 2H), 2.68 (s, 3H).

$^{13}$C NMR (101 MHz, *d*$_6$-DMSO) δ 153.2, 130.1, 126.8, 125.7, 122.3, 110.2, 55.6, 40.0, 35.5, 27.7.

**Bis(7-bromo-1,2,2,4-tetramethyl-1,2-dihydroquinolin-6-yl)methane** (**SI-18**):

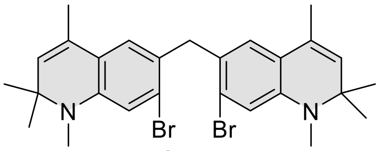

**SI-18**

Compound **SI-15** (3.7 g, 13.9 mmol) was dissolved in AcOH (30 mL) and solution of formaldehyde (5.6 mL, 37%) was added. The reaction mixture was stirred at 60°C for 30 min. Reaction course was followed by TLC (Hexane:EtOAc 95:5). Once reaction was complete reaction mixture was cooled, poured into water and neutralized with NaOH solution. Mixture was extracted with DCM (3x 50 mL), washed with brine, dried over $Na_2SO_4$ and filtered. Filtrate was concentrated on rotary evaporator, the mixture was deposited on celite and product was purified by flash column chromatography (Büchi Reveleris HP silica 80 g, gradient 0% to 10% hexane – EtOAc) to give 2.9 g of off-white solid in a 76% yield.

$^1$H NMR (400 MHz, $CDCl_3$) δ 6.77 (s, 1H), 6.71 (s, 1H), 5.28 (q, *J* = 1.5 Hz, 1H), 3.98 (s, 1H), 2.77 (s, 3H), 1.84 (d, *J* = 1.4 Hz, 3H), 1.28 (s, 6H).

$^{13}$C NMR (101 MHz, $CDCl_3$) δ 144.8, 130.6, 127.9, 126.9, 125.3, 124.7, 123.0, 114.5, 56.5, 40.0, 30.8, 27.2, 18.5.

ESI-MS, positive mode: m/z = 589.0 $[M+H]^+$.

HRMS (ESI) calcd for $C_{26}H_{37}Br_2N_2Si$ $[M+H]^+$ 587.1087, found 587.1014.

**3-bromo-4-((8-bromo-2,3,6,7-tetrahydro-1H,5H-pyrido[3,2,1-ij]quinolin-9-yl)methyl)-N,N-dimethylaniline** (**SI-19**):

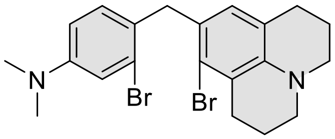

**SI-19**

Compound **SI-12** (1.5g, 5.95mmol), (2-bromo-4-(dimethylamino)phenyl)methanol[16] (1.37 g, 5.95 mmol) and $BF_3 \cdot OEt_2$ (1.5 mL, 11.9 mmol) were dissolved in DCM (40 mL), and the mixture was stirred at room temperature for 5h. The reaction was quenched with water and the solution was extracted with DCM (3x 50mL). The organic layer was washed with brine, dried over $Na_2SO_4$, filtered and filtrate was evaporated to dryness. The residue was purified by flash column chromatography (Teledyne Isco RediSep Rf 80 g; gradient 5% to 50% EtOAc – hexane + 5% constant DCM additive) to give 1.85 g of white solid in a 67% yield.

$^1$H NMR (400 MHz, $CDCl_3$) δ 6.93 (d, *J* = 2.6 Hz, 1H), 6.82 (d, *J* = 8.6 Hz, 1H), 6.58 (dd, *J* = 8.6, 2.7 Hz, 1H), 6.45 (s, 1H), 3.94 (s, 2H), 3.14 – 3.04 (m, 4H), 2.91 (s, 6H), 2.80 (t, *J* = 6.7 Hz, 2H), 2.62 (t, *J* = 6.6 Hz, 2H), 2.06 – 1.85 (m, 4H).

$^{13}$C NMR (101 MHz, $CDCl_3$) δ 150.7, 143.7, 131.1, 129.0, 127.6, 127.1, 126.0, 125.8, 121.7, 121.4, 116.5, 112.3, 50.6, 50.0, 41.4, 40.8, 30.0, 28.1, 22.8, 22.5.

ESI-MS, positive mode: m/z = 465.0 $[M+H]^+$.

HRMS (ESI) calcd for $C_{21}H_{25}Br_2N_2$ $[M+H]^+$ 465.0360, found 465.0360.

**3-bromo-4-((6-bromo-1-methylindolin-5-yl)methyl)-N,N-dimethylaniline** (**SI-20**):

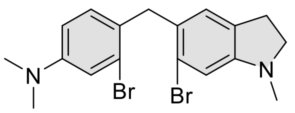

**SI-20**

Compound **SI-3** (2.5 g, 11.8 mmol), (2-bromo-4-(dimethylamino)phenyl)methanol[16] (2.7 g, 11.8 mmol) and $BF_3{\cdot}OEt_2$ (2.95 mL, 23.6 mmol) were dissolved in DCM (100 mL), and the mixture was stirred at room temperature for 5h. The reaction was quenched with water and the solution was extracted with DCM (3x 100mL). The organic layer was washed with brine, dried over $Na_2SO_4$, filtered and filtrate was evaporated to dryness. The residue was purified by flash column chromatography (Teledyne Isco RediSep Rf 80 g; gradient 2% to 40% EtOAc – hexane + 5% constant DCM additive) to give 3.8 g of white solid in a 76% yield.

$^1$H NMR (400 MHz, $CD_2Cl_2$) δ 6.93 (d, *J* = 2.7 Hz, 1H), 6.84 (d, *J* = 8.5 Hz, 1H), 6.70 (s, 1H), 6.64 (s, 1H), 6.59 (dd, *J* = 8.6, 2.7 Hz, 1H), 3.96 (s, 2H), 3.29 (t, *J* = 8.2 Hz, 2H), 2.91 (s, 6H), 2.81 (td, *J* = 8.2, 1.1 Hz, 2H), 2.72 (s, 3H).

$^{13}$C NMR (101 MHz, $CD_2Cl_2$) δ 153.8, 150.8, 131.2, 131.0, 128.3, 127.5, 126.5, 125.9, 123.4, 116.5, 112.3, 111.1, 56.8, 40.8, 40.7, 36.3, 28.9.

ESI-MS, positive mode: m/z = 425.0 $[M+H]^+$.

HRMS (ESI) calcd for $C_{18}H_{21}Br_2N_2$ $[M+H]^+$ 425.0046, found 425.0045.

**3-bromo-4-((7-bromo-1,2,2,4-tetramethyl-1,2-dihydroquinolin-6-yl)methyl)-N,N-dimethylaniline** (**SI-21**):

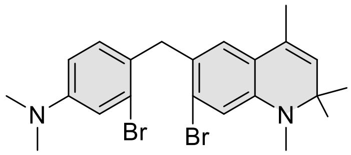

**SI-21**

Compound **SI-15** (1.5 g, 5.6 mmol), (2-bromo-4-(dimethylamino)phenyl)methanol[16] (1.3 g, 5.6 mmol) and $BF_3{\cdot}OEt_2$ (1.4 mL, 11.2 mmol) were dissolved in DCM (50 mL), and the mixture was stirred at room temperature for 2h. The reaction was quenched with water and the solution was extracted with DCM (3x 100mL). The organic layer was washed with brine, dried over $Na_2SO_4$, filtered and filtrate was evaporated to dryness. The residue was purified by flash column chromatography (Teledyne Isco RediSep Rf 80 g; gradient 2% to 40% EtOAc – hexane + 5% constant DCM additive) to give 1.6 g of yellow glassy solid material in a 60% yield.

$^1$H NMR (400 MHz, $CD_2Cl_2$) δ 6.93 (d, *J* = 2.6 Hz, 1H), 6.81 (d, *J* = 8.6 Hz, 1H), 6.78 (s, 1H), 6.68 (s, 1H), 6.58 (dd, *J* = 8.6, 2.7 Hz, 1H), 5.31 (q, *J* = 1.5 Hz, 1H), 3.96 (s, 2H), 2.90 (s, 6H), 2.76 (s, 3H), 1.85 (d, *J* = 1.4 Hz, 3H), 1.29 (s, 6H).

$^{13}C$ NMR (101 MHz, $CD_2Cl_2$) δ 150.7, 145.5, 131.2, 130.9, 127.9, 127.5, 126.7, 126.0, 125.8, 125.1, 123.3, 116.6, 114.6, 112.3, 57.0, 40.8, 40.4, 31.2, 27.6, 18.7.

ESI-MS, positive mode: m/z = 479.1 [M+H]+.

HRMS (ESI) calcd for $C_{22}H_{27}N_2Br_2$ $[M+H]^+$ 477.0536, found 477.0527.

**(6-bromo-1-methylindolin-5-yl)methanol** (**SI-22**):

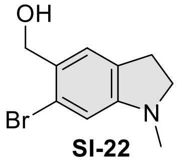

**SI-22**

The (6-bromo-1-methylindolin-5-yl)methanol (**SI-22**)was prepared according to previously published procedure[16].

**8-bromo-9-((6-bromo-1-methylindolin-5-yl)methyl)-2,3,6,7-tetrahydro-1H,5H-pyrido[3,2,1-ij]quinolone** (**SI-23**):

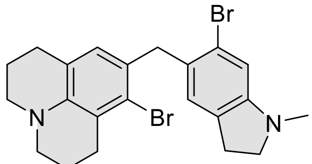

**SI-23**

Compound **SI-22** (1.0 g, 4.0 mmol), compound **SI-12** (0.97 g, 4.0 mmol) and $BF_3{\cdot}OEt_2$ (1.0 mL, 8 mmol) were dissolved in DCM (50 mL), and the mixture was stirred at room temperature for 2h. The reaction was quenched with water and the solution was extracted with DCM (3x 50mL). The organic layer was washed with brine, dried over $Na_2SO_4$, filtered and filtrate was evaporated to dryness. The residue was purified by flash column chromatography (Teledyne Isco RediSep Rf 80 g; gradient 5% to 40% EtOAc – hexane) to give 1.37 g of white solid in a 72% yield.

$^1H$ NMR (400 MHz, $CDCl_3$) δ 6.71 (s, 1H), 6.66 (s, 1H), 6.48 (s, 1H), 3.97 (s, 2H), 3.29 (t, $J$ = 8.1 Hz, 2H), 3.13 – 3.06 (m, 4H), 2.87 – 2.80 (m, 4H), 2.74 (s, 3H), 2.65 (t, $J$ = 6.5 Hz, 2H), 2.04 – 1.91 (m, 4H).

$^{13}C$ NMR (101 MHz, $CDCl_3$) δ 153.1, 143.1, 130.2, 128.7, 128.5, 127.2, 126.2, 125.7, 123.2, 121.4, 120.9, 111.0, 56.5, 50.2, 49.7, 41.4, 36.2, 29.5, 28.5, 27.6, 22.4, 22.1.

ESI-MS, positive mode: m/z = 477.0 $[M+H]^+$.

HRMS (ESI) calcd for C22H25Br2N2 $[M+H]^+$ 477.0360, found 477.0357.

**9-oxo-9H-xanthene-3,6-diyl bis(trifluoromethanesulfonate)** (**SI-24**):

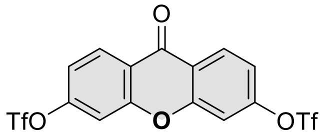

**SI-24**

Dry pyridine (8.9 mL, 110 mmol) was added to a stirred suspension of 3,6-dihydroxy-9H-xanthen-9- one (2.5 g, 11 mmol) in dry dichloromethane (50 mL) at 0 °C. The reaction mixture was stirred for another 15 min at this temperature, and $Tf_2O$ (5.55 mL, 33.0 mmol) was added dropwise during 20 min. The mixture was allowed to warm to room temperature and stirred for additional 2h. After no starting material was observed (TLC), water (30 mL) was added, and the organic layer was separated, washed with aq HCl (1 M, 3 × 30 mL), brine (30 mL) and dried over MgSO4. The solvent was removed under reduced pressure and dried under vacuum to give 5.3g (99%) of pure product as white solid.

$^1$H NMR (400 MHz, $CDCl_3$) δ 8.44 (dd, *J* = 8.9, 0.4 Hz, 1H), 7.49 (d, *J* = 2.3 Hz, 1H), 7.35 (dd, *J* = 8.8, 2.3 Hz, 1H).

$^{13}$C NMR (101 MHz, $CDCl_3$) δ 174.7, 156.7, 153.5, 129.8, 121.5, 118.8 (q, $^1J_{C\text{-}F}$ = 318 Hz, $-CF_3$), 118.3, 111.6.

ESI-MS, positive mode: m/z = 514.9 $[M+Na]^+$.

HRMS (ESI) calcd for $C_{15}H_6F_6O_8S_2Na$ $[M+H]^+$ 514.9300, found 514.9294.

**9,9-dimethyl-10-oxo-9,10-dihydroanthracene-2,7-diyl bis(trifluoromethanesulfonate)** (**SI-25**):

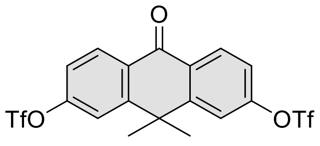

**SI-25**

Dry pyridine (4.8 mL, 59 mmol) was added to a stirred suspension of 3,6-dihydroxy-10,10-dimethyl-9(10H)-anthracenone[17] (1.5 g, 5.9 mmol) in dry dichloromethane (50 mL) at 0°C. The reaction mixture was stirred for another 15 min at this temperature, and $Tf_2O$ (3.0 mL, 17.7 mmol) was added dropwise during 20 min. The mixture was allowed to warm to room temperature and stirred for additional 2h. After no starting material was observed (TLC), water (30 mL) was added, and the organic layer was separated, washed with aq HCl (1 M, 3 × 30 mL), brine (30 mL) and dried over MgSO4. The solvent was removed under reduced pressure and dried under vacuum to give 3.0g (99%) of pure product as white solid.

$^1$H NMR (400 MHz, $CDCl_3$) δ 8.46 (d, *J* = 8.8 Hz, 2H), 7.58 (d, *J* = 2.4 Hz, 2H), 7.38 (dd, *J* = 8.8, 2.4 Hz, 2H), 1.78 (s, 6H)

$^{13}$C NMR (101 MHz, $CDCl_3$) δ 180.9, 153.3, 152.6, 130.9, 129.4, 120.5, 120.0, 118.9 (q, $^1J_{C\text{-}F}$ = 319 Hz, $-CF_3$), 38.7, 33.0.

ESI-MS, positive mode: m/z = 519.0 $[M+H]^+$.

HRMS (ESI) calcd for $C_{18}H_{12}F_6O_7S_2$ $[M+H]^+$ 519.0001, found 518.9996.

**6-(dimethylamino)-9-oxo-9H-xanthen-3-yl trifluoromethanesulfonate** (**SI-26**):

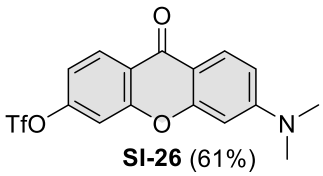

**SI-26** (61%)

*N,N*-Dimethylamine (1.83 mL, 3.66 mmol, 2M in THF) and compound **SI-24** (0.6g, 1.22 mmol) were dissolved in dry DMSO (6 mL). Mixture was stirred and heated in a sealed vial at 90°C overnight. Reaction mixture was cooled, poured into water and extracted with EtOAc (4x 50 mL), washed with brine, dried over $Na_2SO_4$ and filtered. Filtrate was concentrated on rotary evaporator, the residue was deposited on celite and products were purified by flash column chromatography (Büchi

Reveleris HP silica 40 g, gradient 5% to 50% DCM – EtOAc) to give 288 mg of target compound in 61% yield as off-white solid.

$^1$H NMR (400 MHz, $CDCl_3$) δ 8.38 (d, *J* = 8.8 Hz, 1H), 8.13 (d, *J* = 9.1 Hz, 1H), 7.34 (d, *J* = 2.3 Hz, 1H), 7.22 (dd, *J* = 8.8, 2.4 Hz, 1H), 6.74 (dd, *J* = 9.1, 2.4 Hz, 1H), 6.48 (d, *J* = 2.5 Hz, 1H), 3.13 (s, 6H).

$^{13}$C NMR (101 MHz, $CDCl_3$) δ 174.3, 158.5, 156.6, 155.3, 152.3, 129.2, 128.2, 122.3, 118.8 (q, $^1J_{C-F}$ = 319 Hz, $-CF_3$), 116.6, 111.5, 110.8, 110.3, 96.8, 40.3.

ESI-MS, positive mode: m/z = 388.0 $[M+H]^+$.

HRMS (ESI) calcd for $C_{16}H_{13}NO_5F_3S$ $[M+H]^+$ 388.0461, found 388.0464.

**3-(dimethylamino)-6-hydroxy-9H-xanthen-9-one** (**SI-27**):

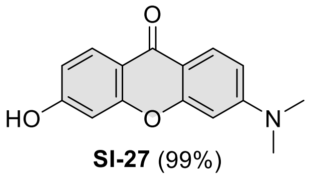

**SI-27** (99%)

To a solution of **SI-26** (288 mg, 0.74 mmol) in *i*-IPrOH (10 mL) solution of NaOH (1 mL, 1M) was added and the obtained mixture was stirred at 60°C for 1h. Reaction mixture was cooled, neutralized with HCl (1 mL, 1M), diluted with water and cooled in an ice bath. The formed precipitate was filtered and dried to give 187 mg of pure compound as light yellow solid in a quantitative yield.

$^1$H NMR (400 MHz, *$d_6$*-DMSO) δ 10.7 (s, 1H), 7.9 (dd, *J* = 21.0, 8.8 Hz, 2H), 6.9 – 6.7 (m, 3H), 6.6 (d, *J* = 1.9 Hz, 1H), 3.1 (s, 6H).

$^{13}$C NMR (101 MHz, *$d_6$*-DMSO) δ 173.5, 162.9, 157.6, 157.3, 154.5, 127.5, 126.9, 114.3, 113.1, 110.4, 109.6, 101.9, 96.6, 39.7.

**3,7-bis(dimethylamino)-5,5-dimethyldibenzo[b,e]silin-10(5H)-one** (**K2**):

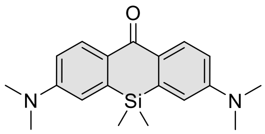

The 3,7-bis(dimethylamino)-5,5-dimethyldibenzo[b,e]silin-10(5H)-one (**K2**)was prepared according to previously published procedure[18].

**3,6-bis(dimethylamino)-9H-xanthen-9-one** (**K3**):

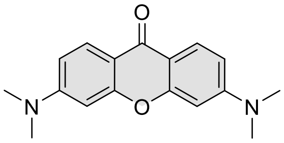

The 3,6-bis(dimethylamino)-9H-xanthen-9-one (**K3**) was prepared according to previously published procedure[19].

**3,6-bis(dimethylamino)-10,10-dimethylanthracen-9(10H)-one** (**K4**):

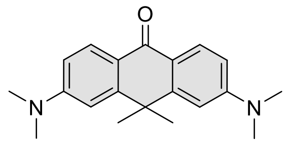

The 3,6-bis(dimethylamino)-10,10-dimethylanthracen-9(10H)-one (**K4**) was prepared according to previously published procedure[18].

Compound **K5**:

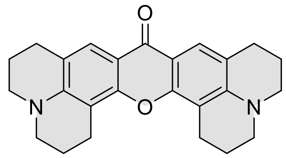

**K5**

Compound **SI-1** (0.7 g, 1.8 mmol) was added to concentrated $H_2SO_4$ (10 mL) and was heated to 90°C for 3 hours. The reaction was cooled and poured onto ice and neutralized with NaOH solution while maintaining temperature with addition of ice, the pH was adjusted to 8-9. The mixture was extracted with DCM (4x 50 mL), the combined organic extracts were dried over $Na_2SO_4$, filtered and solvent was removed under reduced pressure. The residue was dissolved in NMP (20 mL) transferred to Ace pressure tube and $Cs_2CO_3$ (1.2 g, 3.6 mmol), $I_2$ (0.46 g, 1.8 mmol) and water (1 mL) was added. The vial was sealed and heated at 115°C for 2 hours, during the time mixture color changed from brown to purple and back to brown. Mixture was cooled and another portion of $Cs_2CO_3$ (0.59 g, 1.8 mmol) and $I_2$ (0.46 g, 1.8 mmol) was added, same was repeated 2h afterwards and then mixture was left at 115°C overnight. After cooling to rt reaction mixture was filtered through celite, celite was washed with DCM:MeOH (8:2). The filtrate was concentrated on rotavap, NMP was removed by vacuum distillation. The obtained residue was dissolved in DCM:MeOH and celite was added followed by removal of the solvent. The product was purified by flash column chromatography (Büchi Reveleris HP silica 40g; gradient 5% to 40% DCM – EtOAc), fractions containing the product were evaporated to give 0.35g (50% in 2 steps) of brown solid.

$^1$H NMR (400 MHz, $CDCl_3$) δ 7.74 (s, 2H), 3.26 (s, 8H), 2.90 (t, *J* = 6.0 Hz, 4H), 2.81 (t, *J* = 6.0 Hz, 4H), 2.09 – 1.91 (m, 8H).

$^{13}$C NMR (101 MHz, $CDCl_3$) δ 174.8, 153.4, 147.3, 123.5, 118.4, 110.6, 105.6, 50.3, 49.7, 27.6, 21.8, 21.0, 20.7.

ESI-MS, positive mode: m/z = 409.2 $[M+Na]^+$.

HRMS (ESI) calcd for C25H26N2O2Na $[M+Na]^+$ 409.1886, found 409.1879.

Compound **K6:**

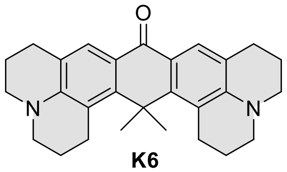

K6

Solution consisting of **SI-11** (500 mg, 2.5 mmol, 1 eq) and **SI-13** (530 mg, 2.5 mmol, 1 eq) in dry DCM was cooled to -78°C in dry ice/acetone cooling bath. Then $BF_3$-$OEt_2$ (1.17 mL, 9.3 mmol) was injected at once and the cooling bath was removed allowing the reaction mixture to warm to room temperature. After stirring for 1h at room temperature reaction was complete (TLC control, hexane : EtoAc 8:2) and the solvent was evaporated. To the crude residue ~40g of polyphosphoric acid (prepared by phosphoric acid + $P_2O_5$ method) was added and the reaction mixture was heated to 145°C. Heating was continued for 3-4h till the reaction was complete and then mixture was poured on ice and neutralized with concentrated NaOH solution. The mixture was extracted with DCM, washed with brine and dried over $Na_2SO_4$. The organic solvent was evaporated and the residue was dissolved in acetone (50 mL). The solution was cooled to -40°C (controlled dry ice/acetone bath) and $KMnO_4$ (395 mg, 2.5 mmol) was added all at once. The temperature of the cooling bath was allowed to slowly rise to -15°C in the course of 1h. TLC (8:2:1 - hexane:EtOAc:DCM) showed incomplete reaction. Reaction mixture was cooled to -40°C and another portion of $KMnO_4$ (100 mg, 0.65 mmol) was introduced. Again temperature of the cooling bath was allowed to slowly rise to -15°C in the course of 1h. Once reaction was complete the reaction was quenched by pouring to cold (-78°C) DCM (100 mL). Celite was added to the mixture, stirred and filtered through another thin layer of celite and washed with DCM. Filtrate was concentrated on a rotary evaporator, the mixture was deposited on celite and product was purified by flash column chromatography (Büchi Reveleris HP silica 40 g, gradient 10% to 60% DCM – EtOAc; Büchi Reveleris HP silica 24g, gradient 20% to 100% Hexane – EtOAc with a constant 5% DCM adittive) and lyophilized from MeCN/$H_2O$ mixture to give 154 mg (15% in 3 steps) of yellow solid.

$^{1}$H NMR (400 MHz, $CD_2Cl_2$) δ 7.76 (s, 1H), 3.29 (q, *J* = 6.1, 5.7 Hz, 4H), 2.84 (dt, *J* = 50.2, 5.9 Hz, 4H), 1.95 (q, *J* = 5.2 Hz, 4H).

$^{13}$C NMR (101 MHz, $CD_2Cl_2$) δ 181.8, 152.2, 148.6, 126.0, 121.2, 118.4, 118.0, 51.3, 50.7, 37.2, 32.2, 28.7, 28.6, 22.4, 22.4.

ESI-MS, positive mode: m/z = 413.3 $[M+H]^+$.

HRMS (ESI) calcd for $C_{28}H_{33}N_2O$ $[M+H]^+$ 413.2587, found 413.2586.

**1,9,11,11-tetramethyl-2,3,7,8,9,11-hexahydrobenzo[1,2-f:5,4-f']diindol-5(1H)-one** (**K7**):

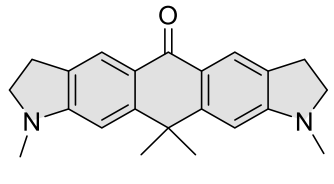

**K7**

Solution consisting of **SI-4** (500 mg, 2.89 mmol, 1eq) and **SI-7** (470 mg, 2.89 mmol, 1 eq) in dry DCM was cooled to -78°C in dry ice/acetone cooling bath. Then $BF_3$-$OEt_2$ (544 μL, 4.33 mmol, 1.5 eq) was injected at once and the cooling bath was removed allowing the reaction mixture to warm to room temperature. After stirring for 1h at room temperature reaction was complete (TLC control, hexane : EtoAc 8:2) and the solvent was evaporated. To the crude residue ~25g of polyphosphoric acid (prepared by phosphoric acid + $P_2O_5$ method) was added and the reaction mixture was heated to 110°C. Heating was continued for 1.5h and then mixture was poured on ice and neutralized with concentrated NaOH solution. The mixture was extracted with DCM, washed with brine and dried over $Na_2SO_4$. The organic solvent was evaporated and the residue was dissolved in acetone (50 mL). The solution was cooled to -15°C (*i*-PrOH:water 1:1, dry ice) and $KMnO_4$ (460 mg, 2.91 mmol) was added in small portions. After 1h, TLC (8:2 - hexane:EtoAc) showed incomplete reaction and another portion of $KMnO_4$ (460 mg, 2.91 mmol) was introduced in small portions. Once reaction was complete the reaction was quenched by pouring to cold (-78°C) DCM (100 mL). Celite was added to the mixture, stirred and filtered through another thin layer of celite, washed with DCM. Filtrate was concentrated on rotary evaporator, the mixture was deposited on celite and product was purified by flash column chromatography (Büchi Reveleris HP silica 40g; gradient 2% to 40% DCM – EtOAc) to give 288 mg (30% in 3 steps) of yellow solid.

$^1$H NMR (400 MHz, $CDCl_3$) δ 8.06 (t, *J* = 1.3 Hz, 2H), 6.47 (s, 2H), 3.49 (t, *J* = 8.3 Hz, 4H), 3.03 (td, *J* = 8.3, 1.3 Hz, 4H), 2.90 (s, 6H), 1.68 (s, 6H).

$^{13}$C NMR (101 MHz, $CDCl_3$) δ 181.1, 156.4, 152.6, 129.4, 123.2, 121.2, 102.0, 55.3, 38.7, 34.7, 33.5, 27.7.

ESI-MS, positive mode: m/z = 355.2 $[M+Na]^+$.

HRMS (ESI) calcd for $C_{22}H_{24}N_2ONa$ $[M+Na]^+$ 355.1781, found 355.1781.

**1,9,11,11-tetramethyl-2,3,7,8,9,11-hexahydrosilino[3,2-f:5,6-f']diindol-5(1H)-one** (**K8**):

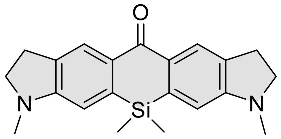

**K8**

In a 250 mL round-bottom flask, a degassed solution of **SI-17** (1.7 g, 3.9 mmol) in anhydrous THF (50 mL) was cooled to -78°C (dry ice – acetone cooling bath). *s*-Butyllithium (7.0 mL of 1.4M in cyclohexane, 9.75 mmol) was introduced through a needle to the reaction mixture and stirring was continued at -78°C for 1.5h. Then $Cl_2SiMe_2$ (470 μL, 3.9 mmol, 2 eq) was slowly injected to the reaction mixture and stirred for 10 min at -78°C before the cooling bath was removed and mixture was allowed to slowly warm to room temperature. Stirring was continued for 1h at rt then it was quenched with 1 mL of saturated $NH_4Cl$ solution. THF was partially evaporated on rotary evaporator. Water was added to the residue and mixture was extracted with DCM (4x 50 mL), washed with brine, dried over $Na_2SO_4$ and filtered. Filtrate was concentrated on rotary evaporator and obtained crude compound was dissolved in 50 mL of acetone. Obtained solution was cooled to -15°C (*i*-PrOH:$H_2O$ 1:1 dry ice cooling bath) and $KMnO_4$ (1.2g ,7.8 mmol) was added in small portions in the course of 3-4h. Reaction course was monitored by TLC (9:1 DCM:EtOAc). Once reaction was complete it was quenched by pouring reaction mixture to -78°C cooled DCM (100 mL). Celite was added to the mixture, stirred and filtered through another thin layer of celite, washed with DCM. Filtrate was concentrated on rotary evaporator, the mixture was deposited on celite and product was purified by flash column chromatography (Büchi Reveleris HP silica 40 g, gradient 2% to 20% DCM – EtOAc) to give 300 mg (22% in 2 steps) of yellow solid.

$^1H$ NMR (400 MHz, $CDCl_3$) δ 8.21 (s, 1H), 6.51 (s, 1H), 3.47 (t, *J* = 8.4 Hz, 2H), 3.05 (td, *J* = 8.7, 1.0 Hz, 2H), 2.90 (s, 3H), 0.45 (s, 3H).

$^{13}C$ NMR (101 MHz, $CDCl_3$) δ 185.2, 154.9, 140.1, 132.3, 131.7, 126.2, 108.0, 55.0, 34.7, 28.2, -0.9.

ESI-MS, positive mode: m/z = 349.2 $[M+H]^+$.

HRMS (ESI) calcd for $C_{21}H_{25}N_2OSi$ $[M+H]^+$ 349.1731, found 349.1733.

**1,2,2,4,8,10,10,11,13,13-decamethyl-2,10,11,13-tetrahydrosilino[3,2-g:5,6-g']diquinolin-6(1H)-one** (**K9**):

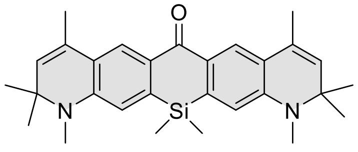

**K9**

In a 250 mL round-bottom flask, a degassed solution of **SI-18** (2 g, 3.7 mmol) in anhydrous THF (50 mL) was cooled to -78°C (dry ice – acetone cooling bath). *s*-Butyllithium (6.6 mL of 1.4M in cyclohexane, 9.25 mmol) was introduced through a needle to the reaction mixture and stirring was continued at -78°C for 1.5h. Then $Cl_2SiMe_2$ (445 µL, 3.7 mmol, 2 eq) was slowly injected to the reaction mixture and stirred for 10 min at -78°C before the cooling bath was removed and mixture was allowed to slowly warm to room temperature. Stirring was continued for 1h at rt then it was quenched with 1 mL of saturated $NH_4Cl$ solution. THF was partially evaporated on rotary evaporator. Water was added to the residue and mixture was extracted with DCM (4x 50 mL), washed with brine, dried over $Na_2SO_4$ and filtered. Filtrate was concentrated on rotary evaporator and obtained crude compound was dissolved in 50 mL of acetone. Obtained solution was cooled to -20°C (Ethylene glycol+10% EtOH dry ice cooling bath) and $KMnO_4$ (1.75 g, 11.1 mmol) was added in small portions in the course of 3-4h. Reaction course was monitored by TLC (8:2 hexane:EtOAc). Once reaction was complete it was quenched by pouring reaction mixture to -78°C cooled DCM (100 mL). Celite was added to the mixture, stirred and filtered through another thin layer of celite, washed with DCM. Filtrate was concentrated on rotary evaporator, the mixture was deposited on celite and product was purified by flash column chromatography (Büchi Reveleris HP silica 40 g, gradient 5% to 70% hexane– EtOAc) to give 590 mg (35% in 2 steps) of yellow solid.

$^1$H NMR (400 MHz, $CDCl_3$) δ 8.19 (s, 1H), 6.57 (s, 1H), 5.32 (q, *J* = 1.4 Hz, 1H), 2.94 (s, 3H), 2.08 (d, *J* = 1.4 Hz, 3H), 1.38 (s, 6H), 0.46 (s, 3H).

$^{13}$C NMR (101 MHz, $CDCl_3$) δ 185.3, 146.9, 140.8, 129.9, 129.9, 128.2, 124.9, 123.4, 112.5, 57.3, 31.2, 28.7, 18.8, -0.9.

ESI-MS, positive mode: m/z = 457.3 $[M+H]^+$.

HRMS (ESI) calcd for $C_{29}H_{36}N_2OSi$ $[M+H]^+$ 457.2670, found 457.2671.

**8-(dimethylamino)-1,10,10-trimethyl-1,2,3,10-tetrahydro-5H-naphtho[2,3-f]indol-5-one** (**K10**):

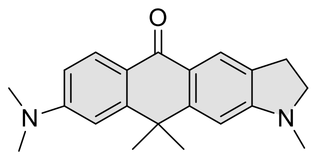

**K10**

Solution consisting of **SI-9** (1 g, 6.2 mmol, 1eq) and **SI-7** (1.02 g, 6.2 mmol, 1 eq) in dry DCM was cooled to -78°C in dry ice/acetone cooling bath. Then $BF_3$-$OEt_2$ (1.17 mL, 9.3 mmol) was injected at once and the cooling bath was removed allowing the reaction mixture to warm to room temperature. After stirring for 1h at room temperature reaction was complete (TLC control, hexane : EtoAc 8:2) and the solvent was evaporated. To the crude residue ~40g of polyphosphoric acid (prepared by phosphoric acid + $P_2O_5$ method) was added and the reaction mixture was heated to 110°C. Heating was continued for 2h and then mixture was poured on ice and neutralized with concentrated NaOH solution. The mixture was extracted with DCM, washed with brine and dried over $Na_2SO_4$. The organic solvent was evaporated and the residue was dissolved in acetone (100 mL). The solution was cooled to -15°C (*i*-PrOH:water 1:1, dry ice) and $KMnO_4$ (980 mg, 6.2 mmol) was added in small portions. After 1h, TLC (8:2 - hexane:EtoAc) showed incomplete reaction and another portion of $KMnO_4$ (490 mg, 3.1 mmol) was introduced in small portions. Once reaction was complete the reaction was quenched by pouring to cold (-78°C) DCM (100 mL). Celite was added to the mixture, stirred and filtered through another thin layer of celite, washed with DCM. Filtrate was concentrated on rotary evaporator, the mixture was deposited on celite and product was purified by flash column chromatography (Büchi Reveleris HP silica 40g, gradient 2% to 50% DCM – EtOAc; Büchi Reveleris HP silica 24g, gradient 20% to 100% Hexane – EtOAc ) to give 635 mg (32% in 3 steps) of yellow solid.

$^1$H NMR (400 MHz, *d$_6$*-DMSO) δ 7.97 (d, *J* = 8.8 Hz, 1H), 7.77 (t, *J* = 1.2 Hz, 1H), 6.85 (d, *J* = 2.5 Hz, 1H), 6.77 (dd, *J* = 8.9, 2.5 Hz, 1H), 6.72 (s, 1H), 3.45 (t, *J* = 8.2 Hz, 2H), 3.06 (s, 6H), 2.96 (td, *J* = 8.2, 1.2 Hz, 2H), 2.89 (s, 3H), 1.65 (s, 6H).

$^{13}$C NMR (101 MHz, *d$_6$*-DMSO) δ 179.4, 156.5, 152.9, 152.5, 151.9, 129.0, 128.0, 121.8, 119.9, 118.9, 110.8, 108.0, 102.4, 66.3, 54.5, 39.8, 38.0, 34.2, 33.2, 27.0.

ESI-MS, positive mode: m/z = 321.2 $[M+H]^+$.

HRMS (ESI) calcd for $C_{21}H_{25}N_2O$ $[M+H]^+$ 321.1961, found 321.1961.

**12-(dimethylamino)-14,14-dimethyl-2,3,5,6,7,14-hexahydro-1H,9H-naphtho[2,3-f]pyrido[3,2,1-ij]quinolin-9-one** (**K11**):

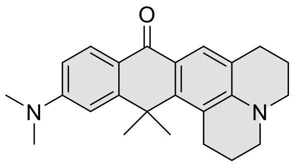

**K11**

Solution consisting of **SI-9** (395 mg, 2.46 mmol, 1 eq) and **SI-11** (500 mg, 2.46 mmol, 1 eq) in dry DCM was cooled to -78°C in dry ice/acetone cooling bath. Then $BF_3$-$OEt_2$ (465 µL, 3.69 mmol) was injected at once and the cooling bath was removed allowing the reaction mixture to warm to room temperature. After stirring for 1h at room temperature reaction was complete (TLC control, hexane : EtoAc 8:2) and the solvent was evaporated. To the crude residue ~25 g of polyphosphoric acid (prepared by phosphoric acid + $P_2O_5$ method) was added and the reaction mixture was heated to 140°C. Heating was continued for 2h and then mixture was poured on ice and neutralized with concentrated NaOH solution. The mixture was extracted with DCM, washed with brine and dried over $Na_2SO_4$. The residue was dissolved in NMP (20 mL) transferred to Ace pressure tube and $Na_3PO_4$-$6H_20$ (2.3 g, 4.92 mmol), $I_2$ (625 mg, 2.46 mmol) and water (0.5 mL) was added. The vial vas sealed and heated at 110°C for 2 hours, during the time mixture color changed from brown to blue and back to brown. Mixture was cooled and another portion of $Na_3PO_4$-$6H_20$ (1.15 g, 2.46 mmol) and $I_2$ (625 mg, 2.46 mmol) was addeds and then mixture was left at 115°C overnight. After cooling to rt reaction mixture was filtered through celite, celite was washed with DCM:MeOH (8:2). The filtrate was concentrated on rotary evaporator, NMP was removed by vacuum distillation. The obtained residue was dissolved in DCM:MeOH and celite was added followed by removal of the solvent. The product was purified by flash column chromatography (Büchi Reveleris HP silica 40g; gradient 20% to 80% Hexane – EtOAc), fractions containing the product were evaporated to give 230 mg(26% in 3 steps) of yellow solid.

$^1H$ NMR (400 MHz, $CD_2Cl_2$) δ 8.07 (d, $J$ = 8.8 Hz, 1H), 7.87 (s, 1H), 6.74 (dd, $J$ = 8.8, 2.5 Hz, 1H), 6.70 (d, $J$ = 2.5 Hz, 1H), 3.32 – 3.27 (m, 4H), 3.08 (s, 6H), 3.04 – 3.00 (m, 2H), 2.82 – 2.78 (m, 2H), 1.99 – 1.93 (m, 4H), 1.82 (s, 6H).

$^{13}C$ NMR (101 MHz, $CD_2Cl_2$) δ 181.4, 156.8, 153.8, 148.5, 146.9, 128.4, 126.9, 121.5, 119.7, 119.4, 118.8, 111.3, 109.0, 51.3, 50.7, 40.6, 38.4, 31.6, 28.7, 28.3, 22.5, 22.3.

ESI-MS, positive mode: m/z = 361.2 $[M+H]^+$.

HRMS (ESI) calcd for $C_{24}H_{28}N_2O$ $[M+H]^+$ 361.2274, found 361.2268.

**9-(dimethylamino)-1,2,2,4,11,11-hexamethyl-2,11-dihydronaphtho[2,3-g]quinolin-6(1H)-one** (**K12**):

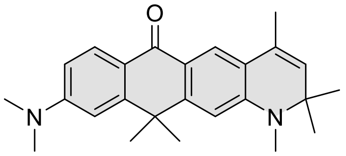

**K12**

Solution consisting of 4-(dimethylamino)benzyl alcohol (432 mg, 2.86 mmol, 1 eq) and **SI-16** (650 mg, 2.86 mmol, 1 eq) in dry DCM was cooled in an ice bath and $BF_3$-$OEt_2$ (540 µL, 4.3 mmol) was injected at once. The cooling bath was removed allowing the reaction mixture to warm to room temperature. After stirring for 3h at room temperature the reaction was complete (TLC control, hexane : EtoAc 8:2) and the solvent was evaporated. To the crude residue ~40g of polyphosphoric acid (prepared by phosphoric acid + $P_2O_5$ method) was added and the reaction mixture was heated to 120°C. Heating was continued for 2h till the reaction was complete and then the mixture was poured on ice and neutralized with concentrated NaOH solution. The mixture was extracted with DCM, washed with brine and dried over $Na_2SO_4$. The organic solvent was evaporated and the residue was dissolved in acetone (50 mL). The solution was cooled to -40°C (controlled dry ice/acetone bath) and $KMnO_4$ (452 mg, 2.86 mmol) was added all at once. The temperature of the cooling bath was allowed to slowly rise to -25°C and was held at this temperature for 2h. Then mixture was allowed to warm to -15 °C and stirred for additional 1h. Reaction course was monitored by TLC (7:2:1 - hexane:EtOAc:DCM). Once reaction was complete it was quenched by pouring mixture to cold (-78°C) DCM (100 mL). Celite was added to the mixture, stirred and filtered through another thin layer of celite, washed with DCM. Filtrate was concentrated on rotary evaporator, the mixture was deposited on celite and product was purified by flash column chromatography (Büchi Reveleris HP silica 40 g, gradient 10% to 70% hexane – EtOAc with a constant 5% DCM additive) to give 225mg (21% in 3 steps) of yellow solid.

$^1$H NMR (400 MHz, $CD_2Cl_2$) δ 8.15 (d, *J* = 9.0 Hz, 1H), 7.91 (s, 1H), 6.79 – 6.74 (m, 2H), 6.57 (s, 1H), 5.36 – 5.33 (m, 1H), 3.09 (s, 6H), 2.95 (s, 3H), 2.07 (d, *J* = 1.4 Hz, 3H), 1.70 (s, 6H), 1.38 (s, 6H).

$^{13}$C NMR (101 MHz, $CD_2Cl_2$) δ 181.0, 153.7, 153.2, 152.8, 149.1, 130.2, 129.1, 127.9, 122.2, 121.7, 120.4, 120.0, 111.3, 108.5, 106.8, 57.9, 40.6, 38.6, 33.9, 31.6, 28.9, 19.0.

ESI-MS, positive mode: m/z = 397.2 $[M+Na]^+$.

HRMS (ESI) calcd for $C_{25}H_{30}N_2ONa$ $[M+Na]^+$ 397.2250, found 397.2249.

**12-(dimethylamino)-14,14-dimethyl-2,3,5,6,7,14-hexahydro-1H,9H-benzo[5,6]silino[2,3-f]pyrido[3,2,1-ij]quinolin-9-one** (**K13**):

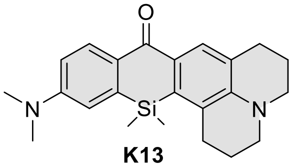

**K13**

In a 250 mL round-bottom flask, a degassed solution of **SI-19** (1.1 g, 2.37 mmol) in anhydrous THF (50 mL) was cooled to -78°C (dry ice – acetone cooling bath). *s*-Butyllithium (4.2 mL of 1.4M in cyclohexane, 5.92 mmol) was introduced through a needle to the reaction mixture and stirring was continued at -78°C for 1.5h. Then $Cl_2SiMe_2$ (362 µL, 3.0 mmol,) was slowly injected to the reaction mixture and stirred for 10 min at -78°C before the cooling bath was removed and mixture was allowed to slowly warm to room temperature. Stirring was continued for 1h at rt then it was quenched with 1 mL of saturated $NH_4Cl$ solution. THF was partially evaporated on rotary evaporator. Water was added to the residue and mixture was extracted with DCM (4x 50 mL), washed with brine, dried over $Na_2SO_4$ and filtered. Filtrate was concentrated on rotary evaporator and obtained crude compound was dissolved in 50 mL of acetone. Obtained solution was cooled to -40°C (controlled dry ice/acetone bath) and $KMnO_4$ (395 mg, 2.5 mmol) was added all at once. The temperature of the cooling bath was allowed to slowly rise to -15°C in the course of 1h. TLC (8:2:1 - hexane:EtOAc:DCM) showed incomplete reaction. Reaction mixture was cooled to -40°C and another portion of $KMnO_4$ (300 mg, 1.95 mmol) was introduced. Again temperature of the cooling bath was allowed to slowly rise to -15°C in the course of 1-2h. Once reaction was complete the reaction was quenched by pouring to cold (-78°C) DCM (100 mL). Celite was added to the mixture, stirred and filtered through another thin layer of celite, washed with DCM. Filtrate was concentrated on rotary evaporator, the mixture was deposited on celite and product was purified by flash column chromatography (Büchi Reveleris HP silica 40 g, gradient 5% to 80% hexane – EtOAc +5% constant DCM additive) to give 258 mg (29% in 3 steps) of yellow solid.

$^1H$ NMR (400 MHz, $CD_2Cl_2$) δ 8.24 (dd, *J* = 8.6, 0.7 Hz, 1H), 7.98 (d, *J* = 0.5 Hz, 1H), 6.84 – 6.79 (m, 2H), 3.32 – 3.26 (m, 4H), 3.08 (s, 6H), 2.96 – 2.90 (m, 2H), 2.84 – 2.78 (m, 2H), 2.05 – 1.92 (m, 4H), 0.52 (s, 6H).

$^{13}C$ NMR (101 MHz, $CD_2Cl_2$) δ 185.4, 152.2, 145.8, 141.9, 135.6, 131.2, 131.2, 130.2, 129.7, 129.3, 125.3, 123.4, 114.7, 113.5, 50.9, 50.4, 40.4, 29.5, 28.8, 22.5, 22.2, 0.1.

ESI-MS, positive mode: m/z = 377.2 $[M+H]^+$.

HRMS (ESI) calcd for $C_{23}H_{29}N_2OSi$ $[M+H]^+$ 377.2044, found 377.2038.

**8-(dimethylamino)-1,10,10-trimethyl-1,2,3,10-tetrahydro-5H-benzo[5,6]silino[3,2-f]indol-5-one** (**K14**):

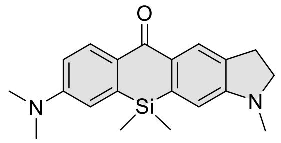

**K14**

In a 250 mL round-bottom flask, a degassed solution of **SI-19** (1.0 g, 2.36 mmol) in anhydrous THF (50 mL) was cooled to -78°C (dry ice – acetone cooling bath). *s*-Butyllithium (4.2 mL of 1.4M in cyclohexane, 5.92 mmol) was introduced through a needle to the reaction mixture and stirring was continued at -78°C for 1.5h. Then $Cl_2SiMe_2$ (355 µL, 2.95 mmol,) was slowly injected to the reaction mixture and stirred for 10 min at -78°C before the cooling bath was removed and mixture was allowed to slowly warm to room temperature. Stirring was continued for 1h at rt then it was quenched with 1 mL of saturated $NH_4Cl$ solution. THF was partially evaporated on rotary evaporator. Water was added to the residue and mixture was extracted with DCM (4x 50 mL), washed with brine, dried over $Na_2SO_4$ and filtered. Filtrate was concentrated on rotary evaporator and obtained crude compound was dissolved in 50 mL of acetone. Obtained solution was cooled to -40°C (controlled dry ice/acetone bath) and $KMnO_4$ (553 mg, 3.5 mmol) was added all at once. The temperature of the cooling bath was allowed to slowly rise to -15°C in the course of 1h. TLC (8:2:1 - hexane:EtOAc:DCM) showed incomplete reaction. Reaction mixture was cooled to -40°C and another portion of $KMnO_4$ (240 mg, 1.52 mmol) was introduced. Again temperature of the cooling bath was allowed to slowly rise to -15°C in the course of 1-2h. Once reaction was complete the reaction was quenched by pouring to cold (-78°C) DCM (100 mL). Celite was added to the mixture, stirred and filtered through another thin layer of celite, washed with DCM. Filtrate was concentrated on rotary evaporator, the mixture was deposited on celite and product was purified by flash column chromatography (Büchi Reveleris HP silica 40 g, gradient 5% to 50% hexane – EtOAc +5% constant DCM additive) to give 270 mg (34% in 2 steps) of yellow solid.

$^1$H NMR (400 MHz, $CD_2Cl_2$) δ 8.28 (dd, *J* = 8.4, 0.9 Hz, 1H), 8.13 (dt, *J* = 1.3, 0.7 Hz, 1H), 6.84 – 6.81 (m, 2H), 6.54 (s, 1H), 3.47 (t, *J* = 8.5 Hz, 2H), 3.08 (s, 6H), 3.04 (td, *J* = 8.4, 1.1 Hz, 2H), 2.90 (s, 3H), 0.45 (s, 6H).

$^{13}$C NMR (101 MHz, $CD_2Cl_2$) δ 185.1, 155.5, 152.1, 140.8, 140.6, 132.9, 131.9, 131.7, 130.1, 126.2, 114.8, 113.5, 108.5, 55.3, 40.4, 34.9, 28.6, -0.8.

ESI-MS, positive mode: m/z = 337.2 [M+H]+.

HRMS (ESI) calcd for $C_{20}H_{25}N_2OSi$ $[M+H]^+$ 337.1731, found 337.1731.

**Compound K15**:

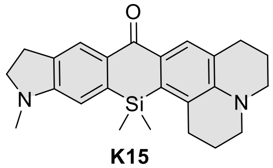

K15

In a 250 mL round-bottom flask, a degassed solution of **SI-23** (1.1 g, 2.3 mmol) in anhydrous THF (50 mL) was cooled to -78°C (dry ice – acetone cooling bath). *s*-Butyllithium (4.1 mL of 1.4M in cyclohexane, 5.75 mmol) was introduced through a needle to the reaction mixture and stirring was continued at -78°C for 1.5h. Then $Cl_2SiMe_2$ (360 µL, 2.9 mmol,) was slowly injected to the reaction mixture and stirred for 10 min at -78°C before the cooling bath was removed and mixture was allowed to slowly warm to room temperature. Stirring was continued for 1h at rt then it was quenched with 1 mL of saturated $NH_4Cl$ solution. THF was partially evaporated on rotary evaporator. Water was added to the residue and mixture was extracted with DCM (4x 50 mL), washed with brine, dried over $Na_2SO_4$ and filtered. Filtrate was concentrated on rotary evaporator and obtained crude compound was dissolved in 70 mL of acetone. Obtained solution was cooled to -40°C (controlled dry ice/acetone bath) and $KMnO_4$ (360 mg, 2.3 mmol) was added all at once. The temperature of the cooling bath was allowed to slowly rise to -15°C in the course of 2h. TLC (8:2 - hexane:EtOAc) showed incomplete reaction. Reaction mixture was cooled to -40°C and another portion of $KMnO_4$ (320 mg, 2 mmol) was introduced. Again temperature of the cooling bath was allowed to slowly rise to -25°C in the course of 1-2h, in the meantime additional amount of $KMnO_4$ (50 mg, 0.32 mmol) was added. Once reaction was complete the reaction was quenched by pouring to cold (-78°C) DCM (100 mL). Celite was added to the mixture, stirred and filtered through another thin layer of celite, washed with DCM. Filtrate was concentrated on rotary evaporator, the mixture was deposited on celite and product was purified by flash column chromatography (Büchi Reveleris HP silica 40 g, gradient 2% to 40% hexane – EtOAc +5% constant DCM additive) to give 335 mg (37% in 2 steps) of yellow solid.

$^1H$ NMR (400 MHz, $CD_2Cl_2$) δ 8.07 (s, 1H), 7.97 (s, 1H), 6.52 (s, 1H), 3.46 (t, *J* = 8.2 Hz, 2H), 3.30 – 3.25 (m, 4H), 3.03 (td, *J* = 8.3, 1.3 Hz, 2H), 2.92 (t, *J* = 6.1 Hz, 2H), 2.89 (s, 3H), 2.81 (t, *J* = 6.3 Hz, 2H), 2.04 – 1.93 (m, 4H), 0.51 (s, 6H).

$^{13}C$ NMR (101 MHz, $CD_2Cl_2$) δ 185.3, 155.6, 145.7, 141.6, 135.5, 132.8, 131.5, 130.2, 129.3, 125.7, 125.2, 123.3, 108.3, 55.4, 50.9, 50.4, 35.0, 29.4, 28.7, 28.6, 22.5, 22.2, 0.0.

ESI-MS, positive mode: m/z = 411.2 $[M+Na]^+$.

HRMS (ESI) calcd for $C_{24}H_{28}N_2OSiNa$ $[M+Na]^+$ 411.1863, found 411.1853.

**9-(dimethylamino)-1,2,2,4,11,11-hexamethyl-2,11-dihydrobenzo[5,6]silino[3,2-g]quinolin-6(1H)-one** (**K16**):

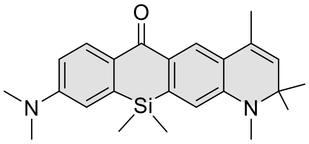

**K16**

In a 250 mL round-bottom flask, a degassed solution of **SI-21** (1.6 g, 3.35 mmol) in anhydrous THF (50 mL) was cooled to -78°C (dry ice – acetone cooling bath). *s*-Butyllithium (6.0 mL of 1.4M in cyclohexane, 8.37 mmol) was introduced through a needle to the reaction mixture and stirring was continued at -78°C for 1.5h. Then $Cl_2SiMe_2$ (506 µL, 4.2 mmol,) was slowly injected to the reaction mixture and stirred for 10 min at -78°C before the cooling bath was removed and mixture was allowed to slowly warm to room temperature. Stirring was continued for 1h at rt then it was quenched with 1 mL of saturated $NH_4Cl$ solution. THF was partially evaporated on rotary evaporator. Water was added to the residue and mixture was extracted with DCM (4x 50 mL), washed with brine, dried over $Na_2SO_4$ and filtered. Filtrate was concentrated on rotary evaporator and obtained crude compound was dissolved in 50 mL of acetone. Obtained solution was cooled to -40°C (controlled dry ice/acetone bath) and $KMnO_4$ (530 mg, 3.35 mmol) was added all at once. The temperature of the cooling bath was allowed to slowly rise to -15°C in the course of 2h. TLC (8:2:1 - hexane:EtOAc:DCM) showed incomplete reaction. Reaction mixture was cooled to -40°C and another portion of $KMnO_4$ (320 mg, 2 mmol) was introduced. Again temperature of the cooling bath was allowed to slowly rise to -15°C in the course of 1-2h. Once reaction was complete the reaction was quenched by pouring to cold (-78°C) DCM (100 mL). Celite was added to the mixture, stirred and filtered through another thin layer of celite, washed with DCM. Filtrate was concentrated on rotary evaporator, the mixture was deposited on celite and product was purified by flash column chromatography (Büchi Reveleris HP silica 40 g, gradient 5% to 50% hexane – EtOAc +5% constant DCM additive) to give 418 mg (32% in 2 steps) of yellow solid.

$^{1}$H NMR (400 MHz, $CD_2Cl_2$) δ 8.29 (dd, *J* = 8.5, 0.8 Hz, 1H), 8.08 (s, 1H), 6.85 – 6.79 (m, 2H), 6.60 (s, 1H), 5.36 (d, *J* = 1.3 Hz, 1H), 3.08 (s, 6H), 2.94 (s, 3H), 2.07 (d, *J* = 1.4 Hz, 3H), 1.37 (s, 6H), 0.46 (s, 6H).

$^{13}$C NMR (101 MHz, $CD_2Cl_2$) δ 185.1, 152.1, 147.5 (2 peaks based on HMBC), 141.3, 140.9, 131.6, 130.7, 130.1, 128.3, 124.9, 123.9, 114.9, 113.5, 113.2, 57.7, 40.4, 31.5, 28.8, 19.0, -0.8.

ESI-MS, positive mode: m/z = 391.3 $[M+H]^+$.

HRMS (ESI) calcd for C24H31N2OSi $[M+H]^+$ 391.2200, found 391.2196.

**3,6-bis(diallylamino)-9H-xanthen-9-one** (**K17**):

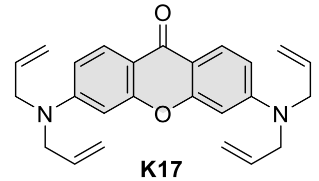

**K17**

Diallylamine (2.5 mL, 20.3 mmol) and compound **SI-24** (1 g, 2.03 mmol) were dissolved in dry DMSO (6 mL). Mixture was stirred and heated in a sealed vial at 100°C for 24h. Reaction course was monitored by TLC (hexane:EtOAc 3:7). Once reaction was finished mixture was poured to water and extracted with EtOAc (4x 50 mL), washed with brine, dried over $Na_2SO_4$ and filtered. Filtrate was concentrated on rotary evaporator, the residue was deposited on celite and products were purified by flash column chromatography (Büchi Reveleris HP silica 40 g, gradient 5% to 40% hexane – EtOAc +10% constant DCM additive) to give 400 mg of target compound in 51% yield as yellow solid.

$^{1}$H NMR (400 MHz, *d₆*-DMSO) δ 7.84 (d, *J* = 9.0 Hz, 2H), 6.74 (dd, *J* = 9.0, 2.4 Hz, 2H), 6.53 (d, *J* = 2.4 Hz, 2H), 5.97 – 5.80 (m, 4H), 5.23 – 5.10 (m, 8H), 4.06 (d, *J* = 4.7 Hz, 8H).

$^{13}$C NMR (101 MHz, $d_6$-DMSO) δ 173.0, 157.4, 152.7, 133.3, 126.8, 116.2, 111.0, 109.4, 97.1, 52.5.

ESI-MS, positive mode: m/z = 409.2 $[M+Na]^+$.

HRMS (ESI) calcd for $C_{25}H_{26}N_2O_2Na$ $[M+Na]^+$ 409.1886, found 409.1881.

**3,6-bis(allyl(methyl)amino)-9H-xanthen-9-one** (**K18**):

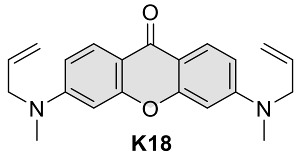

**K18**

*N*-Allylmethylamine (1.95 mL, 20.3 mmol) and compound **SI-24** (1 g, 2.03 mmol) were dissolved in dry DMSO (6 mL). Mixture was stirred and heated in a sealed vial at 100°C for 24h. Reaction course was monitored by TLC (hexane:EtOAc 3:7). Once reaction was finished mixture was poured to water and extracted with EtOAc (4x 50 mL), washed with brine, dried over $Na_2SO_4$ and filtered. Filtrate was concentrated on rotary evaporator, the residue was deposited on celite and products were purified by flash column chromatography (Büchi Reveleris HP silica 40 g, gradient 20% to 80% hexane – EtOAc +5% constant DCM additive) to give 325 mg of target compound in 48% yield as yellow solid.

$^1$H NMR (400 MHz, $d_6$-DMSO) δ 7.86 (d, $J$ = 9.0 Hz, 1H), 6.76 (dd, $J$ = 9.0, 2.4 Hz, 1H), 6.53 (d, $J$ = 2.4 Hz, 1H), 5.85 (ddt, $J$ = 17.1, 10.4, 4.8 Hz, 1H), 5.21 – 5.04 (m, 2H), 4.17 – 4.00 (m, 2H), 3.05 (s, 3H).

$^{13}$C NMR (101 MHz, $d_6$-DMSO) δ 173.1, 157.5, 153.4, 132.9, 126.8, 116.1, 110.9, 109.3, 96.8, 53.9, 38.1.

ESI-MS, positive mode: m/z = 335.2 $[M+H]^+$.

HRMS (ESI) calcd for $C_{21}H_{23}N_2O_2$ $[M+H]^+$ 335.1754, found 335.1754.

**3,6-bis(allyl(methyl)amino)-10,10-dimethylanthracen-9(10H)-one** (**K19**):

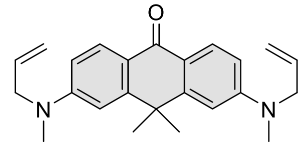
**K19**

*N*-Allylmethylamine (1.85 mL, 19.3 mmol) and compound **SI-25** (1 g, 1.93 mmol) were dissolved in dry DMSO (6 mL). Mixture was stirred and heated in a sealed vial at 100°C for 24h. Reaction course was monitored by TLC (hexane:EtOAc 3:7). Once reaction was finished mixture was poured into water and extracted with EtOAc (4x 50 mL), washed with brine, dried over $Na_2SO_4$ and filtered. Filtrate was concentrated on rotary evaporator, the residue was deposited on celite and products were purified by flash column chromatography (Büchi Reveleris HP silica 40 g, gradient 5% to 40% hexane – EtOAc +5% constant DCM additive) to give 257 mg of target compound in 37% yield as yellow solid.

$^1$H NMR (400 MHz, $d_6$-DMSO) δ 7.96 (d, $J$ = 8.9 Hz, 2H), 6.85 (d, $J$ = 2.5 Hz, 2H), 6.77 (dd, $J$ = 8.9, 2.5 Hz, 2H), 5.92 – 5.81 (m, 2H), 5.19 – 5.11 (m, 4H), 4.10 (d, $J$ = 5.0 Hz, 4H), 3.06 (s, 6H), 1.64 (s, 6H).

$^{13}$C NMR (101 MHz, $d_6$-DMSO) δ 179.4, 152.1, 152.1, 133.4, 128.1, 118.8, 116.2, 110.8, 108.2, 53.9, 38.0, 37.7, 33.3.

ESI-MS, positive mode: m/z = 361.4 $[M+H]^+$.

HRMS (ESI) calcd for $C_{24}H_{29}N_2O$ $[M+H]^+$ 361.2279, found 361.2283.

**3-(dimethylamino)-6-methoxy-9H-xanthen-9-one** (**K20**):

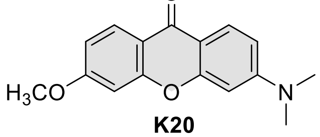
**K20**

Iodomethane (174 µL, 2.8 mmol) was added to a stirred suspension of $K_2CO_3$ (193 mg, 1.4 mmol) and **SI-27** (180 mg, 0.7 mmol) in acetone (10 mL). The reaction mixture was sealed with septa and stirred for 2h at 45°C. Then acetone was evaporated on rotary evaporator, water was added and mixture was extracted with DCM (3x 25 mL), washed with brine, dried over $Na_2SO_4$ and filtered. The filtrate was concentrated on rotary evaporator to give 370 mg of pure compound as a white solid in a quantitative yield.

$^{1}H$ NMR (400 MHz, $CDCl_3$) δ 8.20 (d, *J* = 8.8 Hz, 1H), 8.13 (d, *J* = 9.0 Hz, 1H), 6.88 (dd, *J* = 8.8, 2.4 Hz, 1H), 6.81 (d, *J* = 2.4 Hz, 1H), 6.70 (dd, *J* = 9.0, 2.5 Hz, 1H), 6.47 (d, *J* = 2.4 Hz, 1H), 3.90 (s, 3H), 3.10 (s, 6H).

$^{13}C$ NMR (101 MHz, $CDCl_3$) δ 175.4, 164.3, 158.4, 158.0, 154.7, 128.1, 127.9, 116.3, 112.2, 111.8, 109.6, 100.3, 97.0, 55.8, 40.3.

ESI-MS, positive mode: m/z = 292.1 $[M+Na]^+$.

HRMS (ESI) calcd for $C_{16}H_{15}NO_3Na$ $[M+Na]^+$ 292.0944, found 292.0943.

**3,6-dimethoxy-10,10-dimethylanthracen-9(10H)-one** (**K21**):

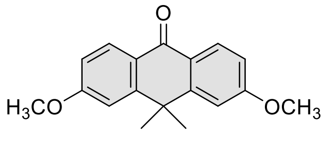

**K21** (94%)

Iodomethane (445 µL, 7.1 mmol) was added to a stirred suspension of $K_2CO_3$ (650 mg, 4.7 mmol) and 3,6-dihydroxy-10,10-dimethyl-9(10H)-anthracenone[18] (300 mg, 1.18 mmol) in acetone (20 mL). The reaction mixture was sealed with septa and stirred overnight at 50°C. Then acetone was evaporated on rotary evaporator, water was added and mixture was extracted with EtOAc (3x 25 mL), washed with brine, dried over $Na_2SO_4$ and filtered. The filtrate was concentrated on rotary evaporator, the residue was deposited on celite and product was purified by flash column chromatography (Büchi Reveleris HP silica 40 g, gradient 2% to 50% hexane – EtOAc) to give 312 mg of target compound in 94% yield as white solid.

$^{1}H$ NMR (400 MHz, $CDCl_3$) δ 8.34 (d, *J* = 8.8 Hz, 1H), 7.10 (d, *J* = 2.4 Hz, 1H), 6.97 (dd, *J* = 8.8, 2.5 Hz, 1H), 3.92 (s, 3H), 1.71 (s, 3H).

$^{13}C$ NMR (101 MHz, $CDCl_3$) δ 181.9, 163.6, 152.8, 130.1, 124.0, 112.8, 111.7, 55.6, 38.3, 33.3.

ESI-MS, positive mode: m/z = 305.1 $[M+Na]^+$.

HRMS (ESI) calcd for $C_{18}H_{19}O_3$ $[M+H]^+$ 283.1329, found 283.1331.

<u>General procedure **A** for the compounds **2-26**:</u>

In a 50 mL round-bottom flask, a degassed solution of 3-bromophthalic acid (100 mg, 0.408 mmol, 1 eq) in anhydrous THF (5 mL) was cooled to -78°C (dry ice – acetone cooling bath). *n*-Butyllithium (640 µL, 1.6M in hexane, 1.02 mmol, 2.5 eq) was introduced through a needle to the reaction mixture dropwise in 3-5 min, the color of reaction mixture starts to change to light orange once ~ the last 0.5 eq are being introduced .Once addition was complete mixture color changed to orange and stirring was continued at -78°C for 20 min. Meantime a corresponding ketone **K2**-**K21** (0.136 mmol, 0.33 eq) was dissolved in minimal amount of THF (1-30 mL, varies depending on ketone) and slowly injected to the reaction mixture with a syringe. The mixture was stirred for additional 10 min. at -78°C and then cooling bath was removed and reaction mixture was allowed to warm to room temperature. Stirring was continued for 30 min at rt and then 1 mL of glacial acetic acid (1 mL, 17.5 mmol) was injected with a syringe resulting in immediate change of color. After stirring for 5 min the THF was partially removed on rotary evaporator and water (20 mL) and solution of HCl was added (2 mL, 1M) and mixture was extracted with DCM or DCM/MeOH (9:1) mixture (5x 25 mL). The combined organic extracts were dried over $Na_2SO_4$ and filtered. Filtrate was concentrated on rotary evaporator, the residue was deposited on celite and products were purified by flash column chromatography.

*NOTE: reaction is highly sensitive to the ratio of reagents used .As even small amount of water can affect the concentration of alkyllithium reagents, we highly recommend to use a freshly distilled anhydrous THF and if needed predetermine exact n-BuLi concentration prior usage by titration.*

<u>General procedure **B** for the compounds **2-26**:</u>

In a 50 mL round-bottom flask, a degassed solution of 3-bromophthalic acid (200 mg, 0.816 mmol, 1 eq) in anhydrous THF (5 mL) was cooled to -78°C (dry ice – acetone cooling bath). *n*-Butyllithium (816 µL, 2.5M in hexane, 2.04 mmol, 2.5 eq) was introduced through a needle to the reaction mixture dropwise in 3-5 min, the color of reaction mixture starts to change to light orange once ~ the last 0.5 eq are being introduced .Once addition was complete mixture color changed to orange and stirring was continued at -78°C for 20 min. Meantime a corresponding ketone **K5**-**K7** (0.136 mmol, 0.17 eq) was partialy dissolved in THF (30 mL) and the obtained suspension was slowly injected to the reaction mixture with a syringe. The mixture was stirred for additional 10 min. at -78$^{o}$C and then cooling bath was removed and reaction mixture was allowed to warm to room temperature. During the warm up stage the suspension slowly turns into solution. Stirring was continued for 1h at rt and then 1 mL of glacial acetic acid (1 mL, 17.5 mmol) was introduced resulting in immediate change of color. After stirring for 5 min the THF was partially removed on rotary evaporator and water (20 mL) and solution of HCl was added (2 mL, 1M) and mixture was extracted with DCM or DCM/MeOH (9:1) mixture (5x 25 mL). The combined organic extracts were dried over $Na_2SO_4$ and filtered. Filtrate was concentrated on rotary evaporator, the residue was deposited on celite and products were purified by flash column chromatography.

**4-SiR-COOH** (**2**):

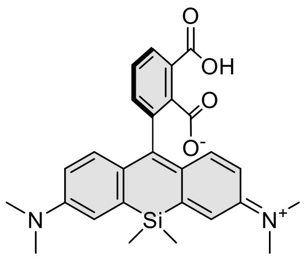

**4-SiR-COOH** (**2**)

Purified by flash column chromatography (Büchi Reveleris HP silica 40 g, gradient 5% to 60% hexane – EtOAc). Obtained 58 mg of target compound in 90% yield as a greenish solid by general method A. Analysis data conforms to previously published data[20].

$^1$H NMR (400 MHz, $d_5$-Pyridine) δ 8.17 (dd, *J* = 7.7, 0.9 Hz, 1H), 7.77 (t, *J* = 7.7 Hz, 1H), 7.54 (dd, *J* = 7.7, 0.9 Hz, 1H), 7.18 (d, *J* = 2.9 Hz, 2H), 7.04 (d, *J* = 9.0 Hz, 2H), 6.54 (dd, *J* = 9.0, 2.9 Hz, 2H), 2.85 (s, 12H), 0.73 (s, 3H), 0.65 (s, 3H).

$^{13}$C NMR (101 MHz, $d_5$-Pyridine) δ δ 170.1, 169.4, 150.4 (overlapped with pyridine, visible in HMBC spectra), 157.1, 137.4, 135.6, 134.9, 132.3, 129.7, 129.2, 127.0, 124.2 (overlapped with pyridine, visible in HMBC spectra), 117.4, 114.6, 92.3, 40.4, 0.8, -0.8.

**4-TMR-COOH** (**3**):

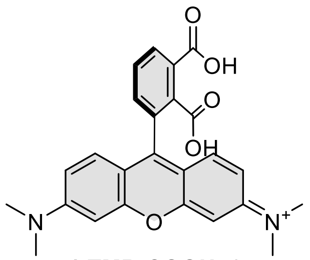

**4-TMR-COOH** (**3**)

Purified by flash column chromatography (Büchi Reveleris HP silica 24 g, gradient 5% to 50% DCM – DCM:MeOH (1:1) +0.1 constant additive of HCOOH). The compound was dissolved in MeCN and filtered through 0.45 µm PTFE filter and lyophilized from MeCN/$H_2O$ mixture to obtain 53 mg of target compound in 84% yield as a dark red solid by general method A. Analysis data conforms to previously published data[20].

$^1$H NMR (400 MHz, $CD_3OD$) δ 8.25 (dd, *J* = 7.8, 1.2 Hz, 1H), 7.82 (t, *J* = 7.8 Hz, 1H), 7.60 (dd, *J* = 7.8, 1.2 Hz, 1H), 7.21 (d, *J* = 9.5 Hz, 2H), 7.08 (dd, *J* = 9.5, 2.4 Hz, 2H), 6.87 (d, *J* = 2.4 Hz, 2H), 3.29 (s, 12H).

$^{13}$C NMR (101 MHz, $CD_3OD$) δ 170.5, 168.6, 159.0, 159.0, 157.0, 136.8, 134.1, 132.8, 132.5, 132.2, 132.0, 131.0, 115.5, 115.2, 97.4, 41.0.

<u>Gram scale experimental procedure of </u>**4-TMR-COOH** (**3**) synthesis:

In a 500 mL round-bottom flask, a degassed solution of 3-bromophthalic acid (4.0 g, 16.3 mmol, 1 eq) in anhydrous THF (100 mL) was cooled to -78°C (dry ice – acetone cooling bath). *n*-Butyllithium (25.1 mL, 2.5M in hexane, 40.2 mmol, 2.5 eq) was introduced through a needle to the reaction mixture dropwise in 3-5 min, the color of reaction mixture starts to change to light orange once ~ the last 0.5 eq were being introduced .Once addition was complete mixture color changed to orange and stirring was continued at -78°C for 20 min. In meantime reaction mixture became cloudy and some of the lithium organics started to crash out of the solution. Ketone **K3** (1.5 g, 5.4 mmol, 0.33 eq) was dissolved in THF (70 mL) and slowly injected to the reaction mixture with a syringe. The mixture was stirred for additional 10 min. at -78°C and then cooling bath was removed and reaction mixture was allowed to warm to room temperature. Stirring was continued for 1h at rt and then glacial acetic acid (5 mL, 76.2 mmol) was injected with a syringe resulting in immediate change of color to dark violet. After stirring for 5 min the THF was partially removed on rotary evaporator and water (70 mL) and solution of HCl was added (15 mL, 1M) and mixture was extracted with DCM/MeOH (9:1) mixture (7x 100 mL). The combined organic extracts were dried over $Na_2SO_4$ and filtered. Filtrate was concentrated on rotary evaporator, the residue was deposited on celite and products were purified by flash column chromatography (Büchi Reveleris HP silica 80 g, gradient 5% to 50% DCM – DCM:MeOH (1:1) +0.1% constant additive of formic acid). The compound was dissolved in 50 mL MeCN and filtered through 0.45 µm PTFE filter. Acetonitrile was evaporated on rotary evaporator to yield 1.7 g compound in 74% yield as a dark violet solid.

*NOTE: The gram scale conditions were not fully optimized. The amount of solvent was reduced to minimal level and some crashing out of reactants from the reaction mixture was evident, thus we recommend to use larger volumes of THF for the ArLi generation and for dissolving **K3** ketone. Increase of solvent might result in better yield.*

**4-610CP-COOH** (**4**):

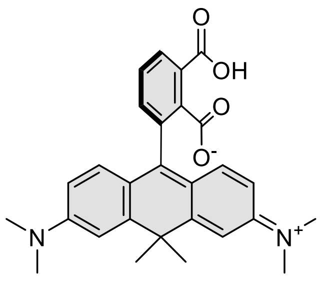

**4-610CP-COOH** (**4**)

Purified by flash column chromatography (Büchi Reveleris HP silica 24 g, gradient 20% to 100% DCM – DCM:MeOH 9:1) The compound was dissolved in MeCN and filtered through 0.45 µm PTFE filter and lyophilized from MeCN/$H_2O$ mixture to give 45 mg of target compound in 72% yield as a dark violet solid by general method A. Analysis data conforms to previously published data[20].

$^1H$ NMR (400 MHz, $CD_3OD$) δ 8.17 (dd, *J* = 7.8, 1.2 Hz, 1H), 7.77 (t, *J* = 7.8 Hz, 1H), 7.53 (dd, *J* = 7.7, 1.2 Hz, 1H), 7.23 (d, *J* = 2.5 Hz, 2H), 7.09 (d, *J* = 9.4 Hz, 2H), 6.84 (dd, *J* = 9.4, 2.5 Hz, 2H), 3.34 (s, 12H), 1.86 (s, 3H), 1.72 (s, 3H).

$^{13}C$ NMR (101 MHz, $CD_3OD$) δ 170.6, 168.9, 158.3, 157.9, 138.9, 136.6, 135.9, 134.2, 131.8, 131.7, 130.6, 122.0, 114.0, 112.1, 48.6, 43.2, 41.0, 36.2, 32.3.

**4-580R-COOH (5)**:

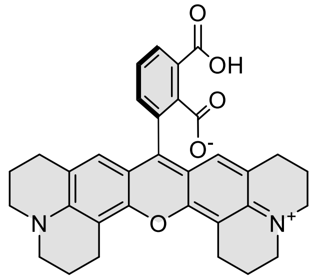

4-580R-COOH (5)

Purified by flash column chromatography (Büchi Reveleris HP silica 24 g, gradient 5% to 60% DCM – DCM:MeOH 1:1 + constant 0.1% HCOOH additive). The compound was dissolved in MeCN and filtered through 0.45 µm PTFE filter and lyophilized from MeCN/$H_2O$ mixture to obtain 27 mg of target compound in 37% yield general method A and 56 mg in 77% yield by general method B as deep violet solid.

$^{1}$H NMR (400 MHz, $d_6$-DMSO, + $CF_3COOD$) δ 8.08 (dd, *J* = 7.8, 1.2 Hz, 1H), 7.77 (t, *J* = 7.7 Hz, 1H), 7.54 (dd, *J* = 7.7, 1.2 Hz, 1H), 6.57 (s, 2H), 3.56 – 3.44 (m, 8H), 2.96 (td, *J* = 6.2, 2.4 Hz, 4H), 2.69 – 2.56 (m, 4H), 2.02 – 1.93 (m, 4H), 1.83 (t, *J* = 6.0 Hz, 4H).

$^{13}$C NMR (101 MHz, $d_6$-DMSO + $CF_3COOD$, open form) δ 168.1, 167.3, 163.2, 151.4, 151.0, 135.3, 133.4, 131.2, 130.8, 130.5, 130.2, 126.2, 123.8, 119.9, 117.0, 114.1, 112.5, 105.1, 50.6, 50.1, 27.1, 20.3, 19.5, 19.4.

ESI-MS, positive mode: m/z = 535.3 $[M+H]^+$.

HRMS (ESI) calcd for C33H31N2O5 $[M+H]^+$ 535.2227, found 535.2224

**4-645CP-COOH (6)**:

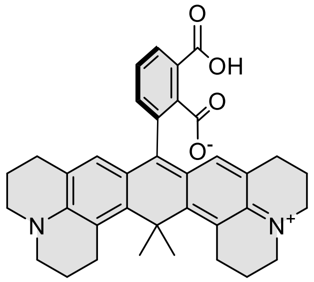

4-645CP-COOH (6)

Purified by flash column chromatography (Büchi Reveleris HP silica 24 g, gradient 5% to 40% DCM – DCM:MeOH 1:1 + constant 0.1% HCOOH additive). The compound was dissolved in MeCN and filtered through 0.45 µm PTFE filter and lyophilized from MeCN/$H_2O$ mixture to obtain 9 mg of target compound in 12% yield general method A and 25 mg in 33% yield by general method B as dark blue solid.

$^{1}$H NMR (400 MHz, $d_6$-DMSO + $CF_3COOD$ ) δ 8.02 (dd, *J* = 7.8, 1.2 Hz, 1H), 7.71 (t, *J* = 7.7 Hz, 1H), 7.44 (dd, *J* = 7.7, 1.2 Hz, 1H), 6.38 (s, 2H), 3.50 (dt, *J* = 21.3, 5.6 Hz, 8H), 3.10 – 2.88 (m, *J* = 5.4 Hz, 4H), 2.50 – 2.36 (m, 4H), 2.13 – 1.90 (m, 7H), 1.89 (s, 3H), 1.85-1.73 (m, 4H).

$^{13}$C NMR (101 MHz, $d_6$-DMSO + $CF_3COOD$, open form) δ 168.3, 167.7, 163.4, 159.2, 154.1, 152.7, 135.8, 135.4, 133.5, 133.1, 130.7, 130.3, 129.8, 122.4, 121.7, 118.7, 117.5, 114.6, 51.8, 51.3, 39.6, 31.3, 30.5, 27.6, 27.5, 20.5, 20.4.

ESI-MS, positive mode: m/z = 561.2 $[M+H]^+$.

HRMS (ESI) calcd for $C_{36}H_{37}N_2O_4$ $[M+H]^+$ 561.2748, found 561.2737.

**4-640CP-COOH** (**7**):

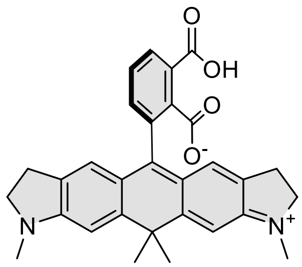

**4-640CP-COOH** (**7**)

Purified by flash column chromatography (Interchim puriflash 15 µm silica 25 g, gradient 5% to 30% DCM – DCM:MeOH 1:1 + constant 0.1% HCOOH additive). The compound was dissolved in MeCN and filtered through 0.45 µm PTFE filter and lyophilized from MeCN/$H_2O$ mixture to obtain 21 mg of target compound in 32% yield general method A and 35 mg in 54% yield by general method B as blue solid.

$^1$H NMR (400 MHz, $CD_3OD$) δ 8.16 (dd, *J* = 7.9, 1.2 Hz, 1H), 7.75 (t, *J* = 7.8 Hz, 1H), 7.49 (dd, *J* = 7.7, 1.2 Hz, 1H), 7.02 (s, 2H), 6.70 – 6.62 (m, 2H), 3.84 (t, *J* = 8.0 Hz, 4H), 3.21 (s, 6H), 2.98 (tdd, *J* = 8.1, 4.6, 1.4 Hz, 4H), 1.82 (s, 3H), 1.69 (s, 3H).

$^{13}$C NMR (101 MHZ, $CD_3OD$) δ 170.8, 168.9, 160.8, 160.6, 159.2, 136.8, 136.7, 134.4, 133.1, 131.6, 131.5, 130.7, 130.1, 122.8, 105.9, 55.9, 43.9, 36.0, 33.6, 32.3, 26.8.

ESI-MS, positive mode: m/z = 481.3 $[M+H]^+$.

HRMS (ESI) calcd for C30H29N2O4 $[M+H]^+$ 481.2122, found 481.2117

**4-SiR700-COOH** (**8**):

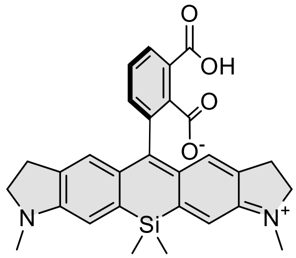

**4-SiR700-COOH** (**8**)

Purified by flash column chromatography (Büchi Reveleris HP silica 40 g, gradient 20% to 80% DCM – DCM:MeOH 9:1). Lyophilized from MeCN/$H_2O$ mixture to give 46 mg of target compound in 69% yield as a yellow-greenish solid by general method A. Analysis data conforms to previously published data.

$^1$H NMR (400 MHz, $d_6$-DMSO) δ 7.71 (t, *J* = 7.5 Hz, 1H), 7.65 (d, *J* = 7.4 Hz, 1H), 7.18 (d, *J* = 7.6 Hz, 1H), 6.78 (s, 2H), 6.53 (s, 2H), 3.27 – 3.16 (m, 4H), 2.85 – 2.67 (m, 10H), 0.58 (s, 3H), 0.49 (s, 3H).

$^{13}$C NMR (101 MHz, $d_6$-DMSO) δ 168.5, 167.2, 156.6, 152.4, 134.9, 133.6, 133.3, 132.5, 132.1, 127.9, 125.1, 122.5, 120.7, 110.0, 90.5, 54.9, 35.3, 28.0, -0.2, -0.5.

ESI-MS, positive mode: m/z = 497.2 $[M+H]^+$.

HRMS (ESI) calcd for C29H29N2O4Si $[M+H]^+$ 497.1891, found 497.1889.

**4-SiR720-COOH** (**9**):

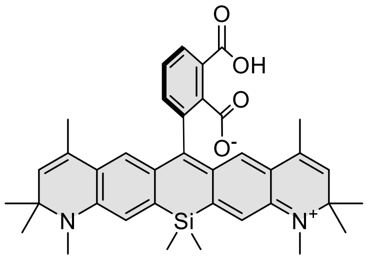

**4-SiR720-COOH** (**9**)

Purified by flash column chromatography (Büchi Reveleris HP silica 24g, gradient 5% to 50% DCM – DCM:MeOH 9:1). Lyophilized from MeCN/$H_2O$ mixture to give 55 mg of target compound in 67% yield as a yellow solid by general method A[21].

$^1$H NMR (400 MHz, $d_6$-DMSO) δ 7.83 (t, *J* = 7.5 Hz, 1H), 7.78 (dd, *J* = 7.5, 1.1 Hz, 1H), 7.45 (dd, *J* = 7.6, 1.1 Hz, 1H), 6.75 (s, 2H), 6.38 (s, 2H), 5.35 (d, *J* = 1.6 Hz, 2H), 2.81 (s, 6H), 1.56 (d, *J* = 1.4 Hz, 6H), 1.25 (s, 6H), 1.24 (s, 6H), 0.62 (s, 3H), 0.51 (s, 3H).

$^{13}$C NMR (101 MHz, $d_6$-DMSO) δ 168.0, 167.0, 154.8, 143.8, 136.1 (2 peaks based on HMBC), 134.4, 132.8, 131.2, 130.5, 128.4, 126.1, 122.8, 122.1, 120.8, 114.2, 90.8, 56.2, 30.5, 27.7, 27.6, 17.4, 0.2, -1.6.

ESI-MS, positive mode: m/z = 605.3 $[M+H]^+$.

HRMS (ESI) calcd for $C_{37}H_{41}N_2O_4Si$ $[M+H]^+$ 605.2830, found 605.2828.

**4-625CP-COOH** (**10**):

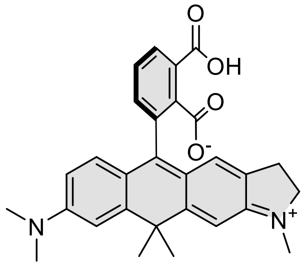

**4-625CP-COOH** (**10**)

Purified by flash column chromatography (Büchi Reveleris HP silica 40g, gradient 5% to 80% DCM – DCM:MeOH 9:1). Lyophilized from MeCN/$H_2O$ mixture to give 45 mg of target compound in 71% yield as a violet solid by general method A.

$^1$H NMR (400 MHz, $CD_3OD$) δ 8.16 (dd, *J* = 7.8, 1.2 Hz, 1H), 7.75 (t, *J* = 7.8 Hz, 1H), 7.51 (dd, *J* = 7.7, 1.2 Hz, 1H), 7.15 (d, *J* = 2.6 Hz, 1H), 7.13 (s, 1H), 6.97 (d, *J* = 9.3 Hz, 1H), 6.77 – 6.71 (m, 2H), 3.93 (t, *J* = 7.7 Hz, 2H), 3.29 (s, 3H), 3.26 (s, 6H), 3.06 – 2.97 (m, 2H), 1.84 (s, 3H), 1.71 (s, 3H).

$^{13}$C NMR (101 MHz, $CD_3OD$) δ 170.7, 168.9, 162.9, 161.8, 160.1, 156.6, 156.3, 137.5, 136.7, 136.3, 134.3, 134.1, 131.7, 131.6, 130.8, 130.6, 123.6, 121.3, 113.1, 111.3, 107.0, 56.4, 43.6, 40.7, 36.1, 33.8, 32.3, 26.5.

ESI-MS, positive mode: m/z = 469.3 $[M+H]^+$.

HRMS (ESI) calcd for C29H29N2O4 $[M+H]^+$ 469.2122, found 469.2118

**4-630CP-COOH** (**11**):

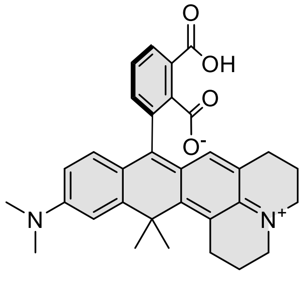

**4-630CP-COOH** (**11**)

Purified by flash column chromatography (Büchi Reveleris HP silica 24g, gradient 5% to 80% DCM – DCM:MeOH 9:1). Lyophilized from MeCN/$H_2O$ mixture to give 42 mg of target compound in 62% yield as a violet solid by general method A.

$^1$H NMR (400 MHz, $d_6$-DMSO) δ 7.63 – 7.55 (m, 2H), 6.95 (dd, $J$ = 7.1, 1.4 Hz, 1H), 6.78 (d, $J$ = 2.6 Hz, 1H), 6.55 (dd, $J$ = 8.9, 2.5 Hz, 1H), 6.39 (d, $J$ = 8.8 Hz, 1H), 6.09 (s, 1H), 3.13 (dt, $J$ = 15.4, 6.4 Hz, 4H), 2.95 – 2.86 (m, 8H), 2.46 – 2.33 (m, 2H), 1.87 (s, 5H), 1.80 (s, 3H), 1.72 (dd, $J$ = 8.6, 4.5 Hz, 2H).

$^{13}$C NMR (101 MHz, $d_6$-DMSO) δ 169.1, 168.2, 157.2, 151.0, 150.1, 145.0, 140.7, 135.3, 128.2, 128.2, 127.9, 126.1, 124.8, 121.9, 121.6, 120.8, 117.5, 116.9, 112.4, 110.0, 88.9, 50.4, 49.7, 40.5, 37.8, 33.1, 32.4, 28.0, 27.9, 22.3, 21.6.

ESI-MS, positive mode: m/z = 509.1 $[M+H]^+$.

HRMS (ESI) calcd for $C_{32}H_{33}N_2O_4$ $[M+H]^+$ 509.2435, found 509.2423.

**4-642CP-COOH (12):**

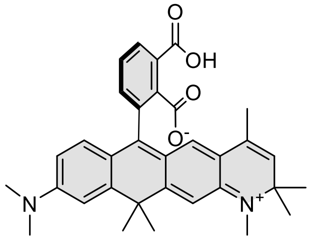

**4-642CP-COOH** (**12**)

Purified by flash column chromatography (Büchi Reveleris HP silica 40g, gradient 5% to 80% DCM – DCM:MeOH 9:1). Lyophilized from MeCN/$H_2O$ mixture to give 48 mg of target compound in 66% yield as a blue solid by general method A.

$^1$H NMR (400 MHz, $d_6$-DMSO) δ 7.76 – 7.70 (m, 2H), 7.15 (dd, $J$ = 7.0, 1.6 Hz, 1H), 6.91 (d, $J$ = 2.6 Hz, 1H), 6.67 (s, 1H), 6.59 (dd, $J$ = 8.9, 2.6 Hz, 1H), 6.48 (d, $J$ = 8.8 Hz, 1H), 6.15 (s, 1H), 5.33 (d, $J$ = 1.6 Hz, 1H), 2.93 (s, 6H), 2.84 (s, 3H), 1.81 (s, 3H), 1.70 (s, 3H), 1.54 (d, $J$ = 1.3 Hz, 3H), 1.26 (s, 6H).

$^{13}$C NMR (101 MHz, $d_6$-DMSO) δ 168.1, 167.1, 155.9, 150.6, 146.2, 146.1, 145.2, 135.1, 132.4, 132.4, 130.7, 128.4, 128.0, 125.5, 122.2, 121.2, 121.0, 118.1, 117.4, 111.8, 109.1, 107.0, 87.2, 56.3, 40.0, 38.0, 34.5, 33.0, 30.7, 27.9, 27.6, 17.5.

ESI-MS, positive mode: m/z = 523.3 [M+H]+.

HRMS (ESI) calcd for $C_{33}H_{35}N_2O_4$ $[M+H]^+$ 523.2591, found 523.2590.

**4-SiR665-COOH** (**13**):

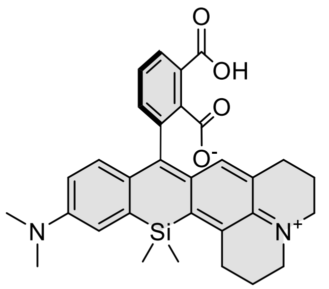

**4-SiR665-COOH** (**13**)

Purified by flash column chromatography (Büchi Reveleris HP silica 40g, gradient 20% to 80% DCM – DCM:MeOH 9:1). Lyophilized from MeCN/$H_2O$ mixture to give 46 mg of target compound in 65% yield as a light blue solid by general method A.

$^1$H NMR (400 MHz, $d_6$-DMSO) δ 7.7 (t, *J* = 7.5 Hz, 1H), 7.6 (dd, *J* = 7.5, 1.1 Hz, 1H), 7.2 (dd, *J* = 7.7, 1.1 Hz, 1H), 6.9 (d, *J* = 2.7 Hz, 1H), 6.7 (dd, *J* = 9.0, 2.7 Hz, 1H), 6.6 (d, *J* = 8.9 Hz, 1H), 6.3 (s, 1H), 3.2 (t, *J* = 5.8 Hz, 2H), 3.1 – 3.1 (m, 2H), 2.9 (s, 8H), 2.5 – 2.4 (m, 2H), 2.0 – 1.9 (m, 2H), 1.8 – 1.7 (m, 2H), 0.7 (s, 3H), 0.6 (s, 3H).

$^{13}$C NMR (101 MHz, $d_6$-DMSO) δ 168.5, 167.1, 157.0, 149.2, 142.1, 135.6, 134.8, 133.2, 130.2, 129.9, 129.5, 127.8, 127.0, 126.2, 125.6, 125.1, 122.9, 120.5, 115.9, 114.1, 90.7, 49.4, 48.9, 39.8, 28.9, 27.7, 21.6, 21.1, 0.8, 0.7.

ESI-MS, positive mode: m/z = 525.3 $[M+H]^+$.

HRMS (ESI) calcd for $C_{31}H_{33}N_2O_4Si$ $[M+H]^+$ 525.2204, found 525.2202.

**4-SiR670-COOH** (**14**):

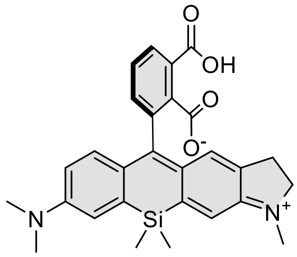

**4-SiR670-COOH** (**14**)

Purified by flash column chromatography (Büchi Reveleris HP silica 40g, gradient 10% to 80% DCM – DCM:MeOH 9:1). Lyophilized from MeCN/$H_2O$ mixture to give 48 mg of target compound in 73% yield as a yellowish-green solid by general method A.

$^1$H NMR (400 MHz, $d_6$-DMSO) δ 7.7 (t, *J* = 7.6 Hz, 1H), 7.7 (dd, *J* = 7.4, 1.0 Hz, 1H), 7.2 (dd, *J* = 7.7, 1.0 Hz, 1H), 7.0 (dd, *J* = 2.2, 1.1 Hz, 1H), 6.8 (s, 1H), 6.7 – 6.6 (m, 2H), 6.5 (s, 1H), 3.2 (dt, *J* = 16.1, 8.2 Hz, 2H), 2.9 (s, 6H), 2.7 (s, 5H), 0.6 (s, 3H), 0.5 (s, 3H).

$^{13}$C NMR (101 MHz, $d_6$-DMSO) δ 168.3, 167.2, 156.0, 152.5, 149.1, 135.3, 134.8, 133.8, 133.5, 132.4, 132.0, 130.5, 127.9, 127.6, 125.4, 122.6, 121.1, 116.1, 113.8, 110.2, 90.5, 54.9, 39.8, 35.3, 28.0, -0.1, -0.8.

ESI-MS, positive mode: m/z = 485.2 $[M+H]^+$.

HRMS (ESI) calcd for C28H29N2O4Si $[M+H]^+$ 485.1891, found 485.1886.

**4-685SiR-COOH** (**15**):

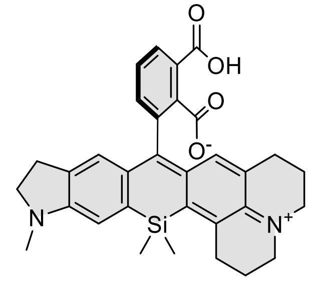

**4-685SiR-COOH** (**15**)

Purified by flash column chromatography (Büchi Reveleris HP silica 40g, gradient 1% to 15% DCM – DCM:MeOH). Lyophilized from MeCN/$H_2O$ mixture to give 40 mg of target compound in 55% yield as a light-blue solid by general method A.

$^1$H NMR (400 MHz, $d_6$-DMSO) δ 7.69 (t, *J* = 7.5 Hz, 1H), 7.64 (dd, *J* = 7.5, 1.2 Hz, 1H), 7.13 (dd, *J* = 7.5, 1.2 Hz, 1H), 6.75 (s, 1H), 6.50 (s, 1H), 6.30 (s, 1H), 3.26 – 3.07 (m, 6H), 2.90 (t, *J* = 6.3 Hz, 2H), 2.83 – 2.67 (m, 5H), 2.50 – 2.37 (m, 2H), 1.99 – 1.90 (m, 2H), 1.80 – 1.70 (m, 2H), 0.65 (s, 3H), 0.54 (s, 3H).

$^{13}$C NMR (101 MHz, $d_6$-DMSO) δ 168.7, 167.0, 157.4, 152.5, 142.0, 135.0, 133.8, 132.9, 132.7, 131.3, 129.8, 129.6, 127.9, 126.1, 125.5, 125.1, 123.0, 121.8, 120.3, 109.8, 90.9, 66.3, 54.9, 49.4, 48.9, 35.3, 28.8, 28.0, 27.7, 21.6, 21.1, 0.8, 0.7.

ESI-MS, positive mode: m/z = 537.1 $[M+H]^+$.

HRMS (ESI) calcd for $C_{32}H_{33}N_2O_4Si$ $[M+H]^+$ 537.2204, found 537.2209.

**4-SiR690-COOH** (**16**):

Purified by flash column chromatography (Büchi Reveleris HP silica 24g, gradient 20% to 80% hexane – EtOAc). Lyophilized from MeCN/$H_2O$ mixture to give 47 mg of target compound in 65% yield as a yellowish-green solid by general method A.

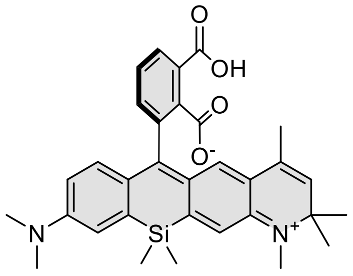

**4-SiR690-COOH** (**16**)

$^1$H NMR (400 MHz, $d_6$-DMSO) δ 7.8 (t, *J* = 7.6 Hz, 1H), 7.7 (dd, *J* = 7.5, 1.0 Hz, 1H), 7.4 (dd, *J* = 7.7, 1.0 Hz, 1H), 7.0 (dd, *J* = 2.4, 0.9 Hz, 1H), 6.8 (s, 1H), 6.7 – 6.6 (m, 2H), 6.4 (s, 1H), 5.3 (d, *J* = 1.6 Hz, 1H), 2.9 (s, 6H), 2.8 (s, 3H), 1.6 (d, *J* = 1.4 Hz, 3H), 1.2 (d, *J* = 4.8 Hz, 6H), 0.6 (s, 3H), 0.5 (s, 3H).

$^{13}$C NMR (101 MHz, $d_6$-DMSO) δ 168.5, 167.6, 155.5, 149.7, 144.3, 136.4, 136.4, 134.9, 133.9, 131.7, 131.1, 131.0, 128.6, 127.9, 126.6, 126.2, 123.4, 122.2, 121.3, 116.9, 114.6, 114.0, 91.0, 56.7, 40.2, 30.9, 28.1, 28.0, 17.9, 0.6, -1.0.

ESI-MS, positive mode: m/z = 539.2 $[M+H]^+$.

HRMS (ESI) calcd for $C_{32}H_{35}N_2O_4Si$ $[M+H]^+$ 539.2361, found 539.2357.

**4-TAllR-COOH** (**17**):

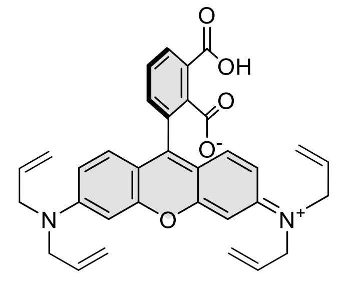

4-TAllR-COOH (17)

Purified by flash column chromatography (Büchi Reveleris HP silica 24g, gradient 1% to 25% DCM – MeOH). Lyophilized from MeCN/$H_2O$ mixture to give 46 mg of target compound in 63% yield as a pink solid by general method A.

$^{1}H$ NMR (400 MHz, $CD_3OD$) δ 8.11 (dd, *J* = 7.8, 1.2 Hz, 1H), 7.64 (t, *J* = 7.7 Hz, 1H), 7.38 (dd, *J* = 7.6, 1.2 Hz, 1H), 7.22 (d, *J* = 9.4 Hz, 2H), 6.99 – 6.93 (m, 2H), 6.90 (d, *J* = 2.5 Hz, 2H), 5.93 (ddt, *J* = 17.1, 10.0, 4.7 Hz, 4H), 5.28 – 5.18 (m, 8H), 4.28 – 4.17 (m, 8H).

$^{13}C$ NMR (101 MHz, $CD_3OD$) δ 172.7, 170.8, 158.7, 157.7, 139.1, 135.3, 135.2, 133.8, 133.1, 132.8, 132.4, 132.4, 130.1, 117.7, 115.0, 114.7, 98.2, 54.5.

ESI-MS, positive mode: m/z = 535.2 $[M+H]^+$.

HRMS (ESI) calcd for $C_{33}H_{31}N_2O_5$ $[M+H]^+$ 535.2227, found 535.2224.

**4-DAllR-COOH** (**18**):

Purified by flash column chromatography (Büchi Reveleris HP silica 40g, gradient 1% to 25% DCM – MeOH). Lyophilized from MeCN/$H_2O$ mixture to give 49 mg of target compound in 75% yield as a pink solid by general method A.

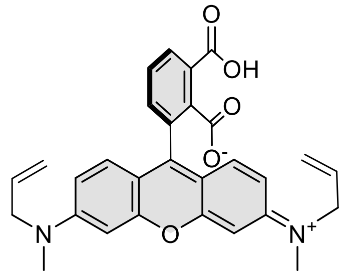

4-DAllR-COOH (18)

$^{1}H$ NMR (400 MHz, $CD_3OD$) δ 8.19 (dd, *J* = 7.8, 1.2 Hz, 1H), 7.78 (t, *J* = 7.8 Hz, 1H), 7.56 (dd, *J* = 7.7, 1.2 Hz, 1H), 7.18 (d, *J* = 9.5 Hz, 2H), 7.05 (dd, *J* = 9.5, 2.5 Hz, 2H), 6.92 (d, *J* = 2.4 Hz, 2H), 5.89 (ddt, *J* = 17.2, 10.0, 4.8 Hz, 2H), 5.25 – 5.11 (m, 4H), 4.31 – 4.20 (m, 4H), 3.26 (s, 6H).

$^{13}C$ NMR (101 MHz, $CD_3OD$) δ 170.4, 168.7, 159.2, 158.9, 157.4, 136.7, 134.1, 133.0, 132.5, 132.2, 132.2, 132.1, 131.1, 117.8, 115.8, 115.5, 97.8, 56.2, 39.6.

ESI-MS, positive mode: m/z = 483.2 [M+H]+.

HRMS (ESI) calcd for $C_{29}H_{27}N_2O_5$ $[M+H]^+$ 483.1914, found 483.1910.

**4-DAllCP-COOH 19**:

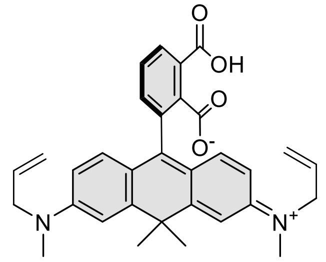

**4-DAllCP-COOH (19)**

Purified by flash column chromatography (Büchi Reveleris HP silica 40g, gradient 10% to 100% DCM – DCM:MeOH 9:1). Lyophilized from MeCN/$H_2O$ mixture to give 50 mg of target compound in 73% yield as a dark violet solid by general method A.

$^1$H NMR (400 MHz, $CD_3OD$) δ 8.03 (dd, *J* = 7.6, 1.0 Hz, 1H), 7.72 (t, *J* = 7.7 Hz, 1H), 7.22 (dd, *J* = 7.7, 1.0 Hz, 1H), 7.00 (d, *J* = 2.6 Hz, 2H), 6.71 (d, *J* = 9.0 Hz, 2H), 6.62 (dd, *J* = 9.0, 2.6 Hz, 2H), 5.87 (ddt, *J* = 17.1, 10.3, 4.9 Hz, 2H), 5.19 – 5.11 (m, 4H), 4.09 – 4.02 (m, 4H), 3.06 (s, 6H), 1.82 (s, 3H), 1.71 (s, 3H).

$^{13}$C NMR (101 MHz, $CD_3OD$) δ 172.6, 169.6, 152.8, 150.6, 134.8, 134.3, 133.4, 132.0, 131.6, 131.3, 129.2, 128.0, 120.0, 116.7, 113.1, 111.0, 56.0, 40.4, 38.8, 35.7, 33.1.

ESI-MS, positive mode: m/z = 509.2 $[M+H]^+$.

HRMS (ESI) calcd for $C_{32}H_{33}N_2O_4$ $[M+H]^+$ 509.2435, found 509.2424.

Compound **20**:

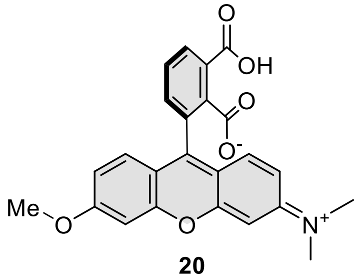

**20**

Purified by flash column chromatography (Büchi Reveleris HP silica 40g, gradient 1% to 20% DCM – DCM:MeOH 1:1 +constant 0.1% additive of HCOOH). Lyophilized from MeCN/$H_2O$ mixture to give 43 mg of target compound in 76% yield as a white solid by general method A.

$^1$H NMR (400 MHz, $d_6$-DMSO) δ 7.80 (d, *J* = 4.3 Hz, 2H), 7.32 (t, *J* = 4.3 Hz, 1H), 6.93 – 6.86 (m, 1H), 6.70 (s, 2H), 6.61 – 6.43 (m, 3H), 3.81 (s, 3H), 2.95 (s, 6H).

$^{13}$C NMR (101 MHz, $d_6$-DMSO) δ 167.1, 166.9, 161.0, 153.3, 152.1, 152.0, 151.8, 135.5, 132.8, 129.1, 129.0, 128.4, 125.7, 122.5, 111.6, 111.0, 109.3, 105.2, 100.8, 97.9, 82.8, 55.6, 39.8.

ESI-MS, negative: m/z = 416.1 $[M-H]^-$.

HRMS (ESI) calcd for $C_{24}H_{18}NO_6$ $[M-H]^-$ 416.1140, found 416.1153.

Compound **21**:

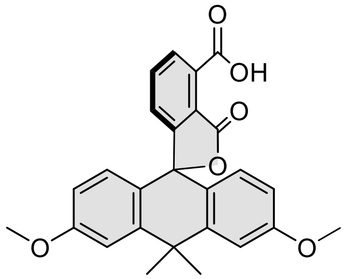

**21**

Purified by flash column chromatography (Büchi Reveleris HP silica 40g, gradient 20% to 100% DCM – DCM:MeOH 9:1). Lyophilized from MeCN/$H_2O$ mixture to give 53 mg of target compound in 91% yield as a white solid by general method A.

$^1$H NMR (400 MHz, *d$_6$*-DMSO) δ 7.79 – 7.71 (m, 2H), 7.28 (d, *J* = 2.6 Hz, 2H), 7.15 (dd, *J* = 7.2, 1.5 Hz, 1H), 6.83 (dd, *J* = 8.8, 2.6 Hz, 2H), 6.65 (d, *J* = 8.8 Hz, 2H), 3.80 (s, 6H), 1.83 (s, 3H), 1.73 (s, 3H).

$^{13}$C NMR (101 MHz, *d$_6$*-DMSO) δ 167.8, 166.9, 159.8, 155.9, 146.5, 135.6, 132.4, 128.8, 128.7, 125.5, 122.6, 121.6, 113.7, 111.6, 85.1, 55.3, 38.1, 34.1, 33.3.

ESI-MS, negative mode: m/z = 429.1 [M-H]$^-$.

HRMS (ESI) calcd for $C_{26}H_{21}O_6$ [M-H]$^-$ 429.1344, found 429.1342.

Compound **22**:

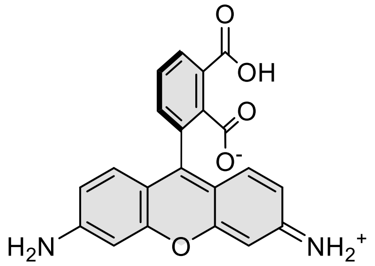

**4-505R-COOH (22)**

1,3-dimethylbarbituric acid (100 mg, 0.64 mmol) and $Pd(PPh_3)_4$ (15 mg, 0.013 mmol) were added to a solution of **4-TAllR-COOH** (**17**) (50 mg, 0.094 mmol) in MeCN (10 mL). The reaction mixture was purged with argon and the vial was sealed. The mixture was stirred at 45°C. The reaction course was monitored by LC/MS analysis and reaction was complete in 10h of heating. The vial was opened and water (10 mL), MeCN (5 mL) and HCOOH (200 µL) were added to dissolve the formed precipitants. The obtained mixture was filtered through 0.45 µm PTFE filter to remove palladium. The solution was concentrated and dissolved in minimum amount of MeCN/$H_20$ mixture and purified by the preparative HPLC (preparative column: Agilent 5 Prep-C18, 5 µm, 100 x 50 mm; solvent A: H2O + 0.2% v/v HCOOH, solvent B MeCN; temperature 25 °C, gradient A:B - 2 min 80:20 isocratic, 2-15 min 80:20 to 20:80 gradient, 15-20 20:80 isocratic). Fractions containing the product were collected, solvent was removed and obtained residue was lyophilized from 50:50 MeCN:$H_2O$ mixture to obtain 21 mg of product as orange-brown solid in 60% yield.

$^1$H NMR (400 MHz, $CD_3OD$) δ 8.21 (dd, *J* = 7.8, 1.2 Hz, 1H), 7.81 (t, *J* = 7.8 Hz, 1H), 7.60 (dd, *J* = 7.7, 1.2 Hz, 1H), 7.11 (d, *J* = 9.1 Hz, 2H), 6.84 (dd, *J* = 9.0, 2.1 Hz, 2H), 6.81 (d, *J* = 2.1 Hz, 2H).

$^{13}C$ NMR (101 MHz, $CD_3OD$) δ 170.4, 168.7, 161.6, 159.7, 157.6, 136.7, 134.0, 133.6, 132.4, 132.1, 131.0, 117.9, 115.1, 98.3.

ESI-MS, positive mode: m/z = 375.3 $[M+H]^+$.

HRMS (ESI) calcd for $C_{21}H_{15}N_2O_5$ $[M+H]^+$ 375.0975, found 375.0965.

**4-525R-COOH** (**23**):

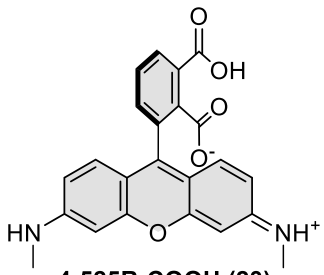

4-525R-COOH (23)

1,3-dimethylbarbituric acid (50 mg, 0.32 mmol) and $Pd(PPh_3)_4$ (10 mg, 0.0086 mmol) were added to a solution of **4-DAllR-COOH** (**18**) (65 mg, 0.135 mmol) in MeCN (10 mL). The reaction mixture was purged with argon and the vial was sealed. The mixture was stirred at 40°C overnight. The reaction completion was confirmed by LC/MS analysis. The vial was and opened and acetonitrile was evaporated to dryness on rotary evaporator. The crude mixture was deposited on celite and product was purified by flash column chromatography (Büchi Reveleris HP silica 40g, gradient 2% to 80% DCM –MeOH). The compound was dissolved in MeCN and filtered through 0.45 µm PTFE filter and lyophilized from MeCN/$H_2O$ mixture to give 39 mg of target compound in 72% yield.

$^1H$ NMR (400 MHz, $d_6$-DMSO) δ 7.62 (t, $J$ = 7.5 Hz, 1H), 7.53 (d, $J$ = 7.4 Hz, 1H), 7.04 (d, $J$ = 7.5 Hz, 1H), 6.47 (d, $J$ = 8.5 Hz, 2H), 6.43 (s, 2H), 6.39 – 6.33 (m, 4H), 2.71 (s, 6H).

$^{13}C$ NMR (101 MHz, $d_6$-DMSO) δ 168.6, 167.8, 153.0, 152.4, 151.0, 134.0, 128.5, 128.4, 127.9, 123.3, 110.2, 110.2, 106.8, 96.1, 29.5.

ESI-MS, positive mode: m/z = 403.1 $[M+H]^+$.

HRMS (ESI) calcd for $C_{23}H_{19}N_2O_5$ $[M+H]^+$ 403.1288, found 403.1288.

**4-580CP-COOH** (**24**):

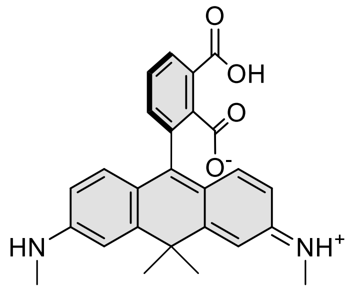

**4-580CP-COOH** (**24**)

1,3-dimethylbarbituric acid (50 mg, 0.32 mmol) and $Pd(PPh_3)_4$ (10 mg, 0.0086 mmol) were added to a solution of **4-DAllCP-COOH** (**19**) (55 mg, 0.108 mmol) in MeCN (10 mL). The reaction mixture was purged with argon and the vial was sealed. The mixture was stirred at 40°C overnight. The reaction completion was confirmed by LC/MS analysis. The vial was opened and acetonitrile was evaporated to dryness on rotary evaporator. The crude mixture was deposited on celite and product was purified by flash column chromatography (Büchi Reveleris HP silica 40g, gradient 2% to 40% DCM –MeOH). The compound was dissolved in MeCN and filtered through 0.45 µm PTFE filter and lyophilized from MeCN/$H_2O$ mixture to give 41 mg of target compound in 89% yield.

$^1$H NMR (400 MHz, $CD_3OD$) δ 8.17 (dd, *J* = 7.9, 1.1 Hz, 1H), 7.76 (t, *J* = 7.7 Hz, 1H), 7.52 (dd, *J* = 7.7, 1.2 Hz, 1H), 7.11 (d, *J* = 2.2 Hz, 2H), 7.03 (d, *J* = 9.2 Hz, 2H), 6.64 (dd, *J* = 9.2, 2.2 Hz, 2H), 3.06 (s, 6H), 1.81 (s, 3H), 1.68 (s, 3H).

$^{13}$C NMR (101 MHz, $cd_3od$) δ 170.6, 168.9, 162.5, 159.4, 158.6, 139.6, 136.6, 136.0, 134.2, 131.7, 131.7, 130.6, 122.1, 112.1 (visible from HSQC), 42.7, 35.8, 31.9, 30.1.

**4-DMRhol-COOH** (**25**):

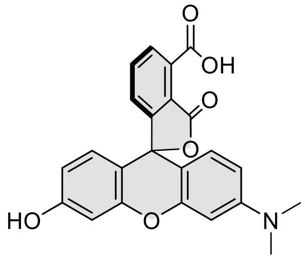

**4-DMRhol-COOH** (**25**)

Compound **20** (60 mg, 0.144 mmol) was suspended in dry 1,2-dichloroethane (20 mL) in ace pressure tube and 6 ml of 1M $BBr_3$ (6 mmol) solution in DCM was added. The tube was sealed and stirred at 55°C (silicon oil bath temperature) for 24h. Reaction completion was confirmed by LC/MS analysis. The mixture was cooled to room temperature and the tube was carefully opened and poured into water (20 mL) and the mixture was extracted with DCM/MeOH (9:1) mixture (7x 25 mL). The extracts were combined, dried over $Na_2SO_4$ and filtered. Filtrate was evaporated and the obtained crude mixture was dissolved in MeCN/$H_2O$ + 0.2 HCOOH 1:1 mixture and purified by the preparative HPLC (preparative column: Agilent 5 Prep-C18, 5 µm, 100 x 50 mm; solvent A: H2O + 0.2% v/v HCOOH, solvent B MeCN; temperature 25 °C, gradient A:B - 2 min 80:20 isocratic, 2-15 min 80:20 to 20:80 gradient, 15-20 20:80 isocratic). Fractions containing the product were collected, solvent was removed and obtained residue was lyophilized from 50:50 MeCN:$H_2O$ mixture to obtain 41 mg of product as red solid in 70% yield.

$^1$H NMR (400 MHz, $CD_3OD$) δ 8.24 (dd, *J* = 7.9, 1.2 Hz, 1H), 7.84 (t, *J* = 7.8 Hz, 1H), 7.63 (dd, *J* = 7.7, 1.2 Hz, 1H), 7.37 – 7.26 (m, 3H), 7.07 (dd, *J* = 9.7, 2.3 Hz, 2H), 7.03 (dd, *J* = 9.1, 2.3 Hz, 1H), 3.40 (s, 6H).

$^{13}C$ NMR (101 MHz, $CD_3OD$) δ 170.3, 169.3, 168.7, 160.7, 160.1, 158.9, 158.5, 136.5, 133.9, 133.7, 133.4, 132.6, 132.4, 132.1, 131.2, 118.5, 118.2, 117.7, 116.5, 103.3, 97.6, 41.6.

ESI-MS, negative mode: m/z = 402.1 $[M-H]^-$.

HRMS (ESI) calcd for $C_{23}H_{16}NO_6$ $[M-H]^-$ 402.0983, found 402.0983.

**4-CFL-COOH** (**26**):

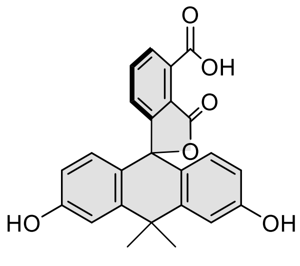

**4-CFL-COOH** (**26**)

Compound **21** (70 mg, 0.163 mmol) was suspended in dry 1,2-dichloroethane (20 mL) in ace pressure tube and 6 ml of 1M $BBr_3$ (6 mmol) solution in DCM was added. The tube was sealed and stirred at 55°C (silicon oil bath temperature) for 72h. Reaction completion was confirmed by LC/MS analysis. The mixture was cooled to room temperature and the tube was carefully opened and poured into water (20 mL) and the mixture was extracted with DCM/MeOH (9:1) mixture (7x 25 mL). The extracts were combined, dried over $Na_2SO_4$ and filtered. Filtrate was evaporated and the obtained crude mixture was deposited on celite and purified by flash column chromatography (Büchi Reveleris HP silica 40g, gradient 2% to 25% DCM –MeOH) to give 43 mg in 65% yield as white solid.

$^1H$ NMR (400 MHz, $d_6$-DMSO) δ 9.94 (s, 2H), 7.72 – 7.58 (m, 2H), 7.07 (dd, $J$ = 9.9, 1.9 Hz, 3H), 6.62 (dd, $J$ = 8.6, 2.4 Hz, 2H), 6.50 (d, $J$ = 8.6 Hz, 2H), 1.72 (s, 3H), 1.62 (s, 3H).

$^{13}C$ NMR (101 MHz, $d_6$-DMSO) δ 168.1, 167.6, 158.1, 155.8, 146.5, 140.7, 135.2, 128.8, 128.1, 124.6, 121.5, 121.4, 114.9, 112.6, 85.5, 37.6, 34.4, 33.1.

ESI-MS, positive mode: m/z = 403.1 $[M+H]^+$.

HRMS (ESI) calcd. for $C_{24}H_{19}O_6$ $[M+H]^+$ 403.1176, found 403.1170.

<u>General procedure for the synthesis of compounds **27**-**46**:</u>

Into a solution of corresponding rhodamine dye (5 µmol, 1 eq; **2-16** and **22-26**) and DIPEA (5 µL) in DMSO (100 µL) a solution of HATU (6.5 µmol, 1.3 eq) in DMSO (100 µL) was added and the mixture was mixed for 1 min. Then a solution of 4-((4-(aminomethyl)benzyl)oxy)-6-chloropyrimidin-2-amine (7.5 µmol ,1.5 eq) in DMSO (100 µL) was added at once. Reactions were usually over after 30 min and their course was monitored by LC/MS analysis. Once reaction was finished it was quenched with 20 µL of formic acid and diluted with water and acetonitrile to 2 mL volume and was further purified by the means of preparative HPLC (preparative column: Agilent 5 Prep-C18, 5 µm, 100 x 50 mm).

**4-CFL-CP** (**27**): JB603

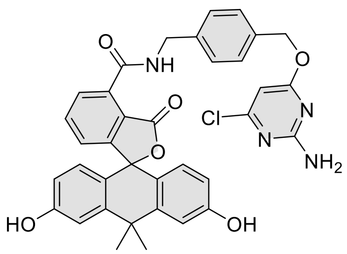

Compound was purified by preparative HPLC (solvent A: $H_2O$ + 0.2% v/v HCOOH, solvent B: acetonitrile; temperature 25 °C, gradient A:B - 3 min 70:30 isocratic, 4-20 min 70:30 to 0:100 gradient, 20-25 min 0:100 isocratic, 40 mL/min flow). Fractions containing the product were collected, evaporated and lyophilized from acetonitrile: water mixture. Yield 42% (1.4 mg) of white solid.

$^1$H NMR (400 MHz, $d_6$-DMSO) δ 9.72 (s, 2H), 9.49 (t, *J* = 5.8 Hz, 1H), 7.80 (dd, *J* = 7.5, 1.0 Hz, 1H), 7.72 (t, *J* = 7.6 Hz, 1H), 7.49 (d, *J* = 8.3 Hz, 2H), 7.44 (d, *J* = 8.3 Hz, 2H), 7.13 – 7.04 (m, 5H), 6.62 (d, *J* = 1.4 Hz, 4H), 6.14 (s, 1H), 5.32 (s, 2H), 4.59 (d, *J* = 5.8 Hz, 2H), 1.73 (s, 3H), 1.64 (s, 3H).

ESI-MS, positive mode: m/z = 671.2 $[M+Na]^+$.

HRMS (ESI) calcd for $C_{36}H_{29}ClN_4O_6Na$ $[M+H]^+$ 671.1668, found 671.1651.

**4-505R-CP** (**28**):

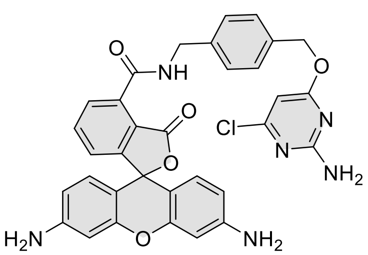

Compound was purified by preparative HPLC (solvent A: $H_2O$ + 10mM $NH_4COOH$ pH = 3.6, solvent B: acetonitrile; temperature 25 °C, gradient A:B - 3 min 80:20 isocratic, 4-20 min 80:20 to 0:100 gradient, 20-25 min 0:100 isocratic, 40 mL/min flow). Fractions containing the product were collected, evaporated and lyophilized from acetonitrile water mixture. The obtained solid was dissolved in 700 µL of $d_6$-DMSO. The samples from DMSO solution were diluted x100 in PBS +0.1%SDS and concentration was determined spectroscopically with Nanodrop. Determined stock concentration was 4.1 mM which constitutes to 58% yield (1.8 mg).

$^1$H NMR (400 MHz, *$d_6$*-DMSO) δ 9.52 (t, *J* = 5.8 Hz, 1H), 7.85 (dd, *J* = 7.5, 1.1 Hz, 1H), 7.78 (t, *J* = 7.6 Hz, 1H), 7.48 (d, *J* = 8.2 Hz, 2H), 7.43 (d, *J* = 8.3 Hz, 2H), 7.26 (dd, *J* = 7.6, 1.1 Hz, 1H), 7.10 (s, 2H), 6.45 (d, *J* = 8.6 Hz, 2H), 6.38 (d, *J* = 2.2 Hz, 2H), 6.29 (dd, *J* = 8.5, 2.2 Hz, 2H), 6.14 (s, 1H), 5.58 (s, 4H), 5.31 (s, 2H), 4.58 (d, *J* = 5.7 Hz, 2H).

ESI-MS, positive mode: m/z = 621.2 $[M+H]^+$.

HRMS (ESI) calcd for $C_{33}H_{31}ClN_6O_5$ $[M+H]^+$ 621.1648, found 621.1639.

**4-525R-CP** (**29**):

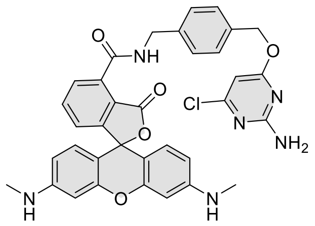

Compound was purified by preparative HPLC (solvent A: $H_2O$ + 10mM $NH_4COOH$ pH = 3.6, solvent B: acetonitrile; temperature 25 °C, gradient A:B - 3 min 80:20 isocratic, 4-20 min 80:20 to 0:100 gradient, 20-25 min 0:100 isocratic, 40 mL/min flow). Fractions containing the product were collected, evaporated and lyophilized from acetonitrile water mixture. The obtained solid was dissolved in 700 µL of $d_6$-DMSO. The samples from DMSO solution were diluted x100 in PBS +0.1%SDS and concentration was determined spectroscopically with Nanodrop. Determined stock concentration was 3.5 mM which constitutes to 49% yield (1.6 mg).

$^1$H NMR (400 MHz, $d_6$-DMSO) δ 9.53 (t, $J$ = 5.8 Hz, 1H), 7.86 (dd, $J$ = 7.5, 1.1 Hz, 1H), 7.78 (t, $J$ = 7.6 Hz, 1H), 7.48 (d, $J$ = 8.2 Hz, 2H), 7.43 (d, $J$ = 8.3 Hz, 2H), 7.25 (dd, $J$ = 7.6, 1.0 Hz, 1H), 7.14 – 7.06 (m, 2H), 6.54 – 6.47 (m, 2H), 6.37 – 6.25 (m, 4H), 6.18 (q, $J$ = 4.9 Hz, 2H), 6.14 (s, 1H), 5.31 (s, 2H), 4.59 (d, $J$ = 5.8 Hz, 2H), 2.69 (d, $J$ = 4.9 Hz, 6H).

ESI-MS, positive mode: m/z = 649.2 $[M+H]^+$.

HRMS (ESI) calcd for $C_{35}H_{30}ClN_6O_5$ $[M+H]^+$ 649.1961, found 649.1955.

**4-DMRhodol-CP** (**30**):

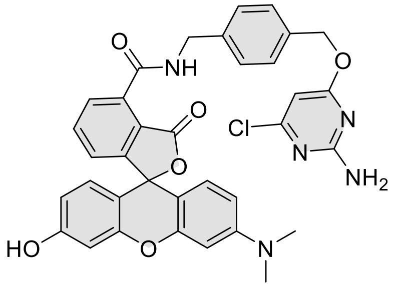

Compound was purified by preparative HPLC (solvent A: $H_2O$ + 0.2% v/v HCOOH, solvent B: acetonitrile; temperature 25 °C, gradient A:B - 3 min 70:30 isocratic, 4-20 min 70:30 to 0:100 gradient, 20-25 min 0:100 isocratic, 40 mL/min flow). Fractions containing the product were collected, evaporated and lyophilized from acetonitrile: water mixture. Yield 52% (1.7 mg) of light red solid.

$^1$H NMR (400 MHz, $d_6$-DMSO) δ 10.13 (br s, 1H), 9.50 (t, $J$ = 5.7 Hz, 1H), 7.84 (dd, $J$ = 7.6, 1.0 Hz, 1H), 7.76 (t, $J$ = 7.6 Hz, 1H), 7.47 (d, $J$ = 8.3 Hz, 2H), 7.41 (d, $J$ = 8.3 Hz, 2H), 7.26 (dd, $J$ = 7.7, 1.1 Hz, 1H), 7.08 (s, 2H), 6.67 (d, $J$ = 8.7 Hz, 1H), 6.65 – 6.60 (m, 2H), 6.53 – 6.47 (m, 3H), 6.13 (s, 1H), 5.30 (s, 2H), 4.57 (d, $J$ = 5.7 Hz, 2H), 2.93 (s, 6H).

ESI-MS, positive mode: m/z = 650.2 $[M+H]^+$.

HRMS (ESI) calcd for $C_{35}H_{29}N_5O_6Cl$ $[M+H]^+$ 650.1801, found 650.1785.

**4-TMR-CP** (**31**):

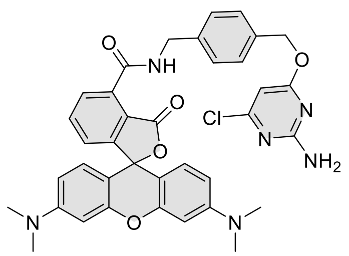

Compound was purified by preparative HPLC (solvent A: $H_2O$ + 10mM $NH_4COOH$ pH = 3.6, solvent B: acetonitrile; temperature 25 °C, gradient A:B - 3 min 80:20 isocratic, 4-20 min 80:20 to 0:100 gradient, 20-25 min 0:100 isocratic, 40 mL/min flow). Fractions containing the product were collected, evaporated and lyophilized from acetonitrile water mixture. The obtained solid was dissolved in 700 µL of $d_6$-DMSO. The samples from DMSO solution were diluted x100 in PBS +0.1%SDS and concentration was determined spectroscopically with Nanodrop. Determined stock concentration was 5.5 mM which constitutes to 77% yield (2.6 mg).

$^1$H NMR (400 MHz, , $d_6$-DMSO) δ 9.51 (t, *J* = 5.8 Hz, 1H), 7.85 (dd, *J* = 7.5, 1.0 Hz, 1H), 7.77 (t, *J* = 7.6 Hz, 1H), 7.47 (d, *J* = 8.3 Hz, 2H), 7.42 (d, *J* = 8.4 Hz, 2H), 7.23 (dd, *J* = 7.7, 1.0 Hz, 1H), 7.09 (s, 2H), 6.66 – 6.59 (m, 2H), 6.52 – 6.40 (m, 4H), 6.13 (s, 1H), 5.30 (s, 2H), 4.58 (d, *J* = 5.7 Hz, 2H), 2.93 (s, 12H).

ESI-MS, positive mode: m/z = 677.2 $[M+H]^+$.

HRMS (ESI) calcd for $C_{37}H_{34}N_6O_5Cl$ $[M+H]^+$ 677.2274, found 677.2262.

**4-580R-CP** (**32**):

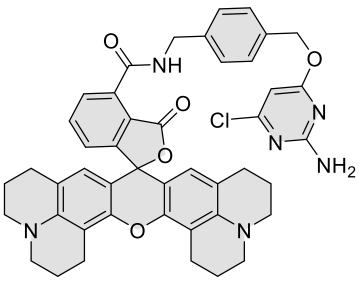

Compound was purified by preparative HPLC (solvent A: $H_2O$ + 10mM $NH_4COOH$ pH = 3.6, solvent B: acetonitrile; temperature 25 °C, gradient A:B - 3 min 80:20 isocratic, 4-20 min 80:20 to 0:100 gradient, 20-25 min 0:100 isocratic, 40 mL/min flow). Fractions containing the product were collected, evaporated and lyophilized from acetonitrile water mixture. The obtained solid was dissolved in 700 µL of $d_6$-DMSO. The samples from DMSO solution were diluted x100 in PBS +0.1%SDS and concentration was determined spectroscopically with Nanodrop. Determined stock concentration was 3.2 mM which constitutes to 45% yield (1.6 mg).

$^1$H NMR (400 MHz, $d_6$-DMSO) δ 9.65 (t, *J* = 5.6 Hz, 1H), 7.88 (dd, *J* = 7.5, 0.9 Hz, 1H), 7.76 (t, *J* = 7.6 Hz, 1H), 7.48 (d, *J* = 8.1 Hz, 2H), 7.42 (d, *J* = 8.2 Hz, 2H), 7.27 (dd, *J* = 7.7, 0.8 Hz, 1H), 7.10 (s, 2H), 6.14 (s, 1H), 6.12 (s, 2H), 5.31 (s, 2H), 4.58 (d, *J* = 5.8 Hz, 2H), 3.13 (s, 8H), 2.85 (t, *J* = 6.4 Hz, 4H), 2.49 – 2.35 (m, 4H), 1.99 – 1.91 (m, 4H), 1.78 (p, *J* = 6.1 Hz, 4H).

ESI-MS, positive mode: m/z = 781.3 $[M+H]^{+.}$

HRMS (ESI) calcd for $C_{45}H_{42}ClN_6O_5$ $[M+H]^+$ 781.2900, found 781.2896.

**4-580CP-CP** (**33**):

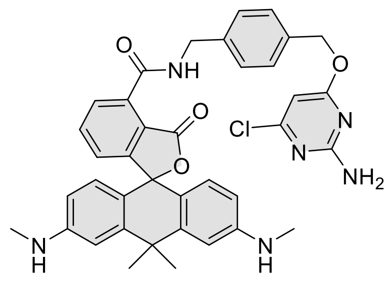

Compound was purified by preparative HPLC (solvent A: $H_2O$ + 10mM $NH_4COOH$ pH = 3.6, solvent B: acetonitrile; temperature 25 °C, gradient A:B - 3 min 80:20 isocratic, 4-20 min 80:20 to 0:100 gradient, 20-25 min 0:100 isocratic, 40 mL/min flow). Fractions containing the product were collected, evaporated and lyophilized from acetonitrile water mixture. The obtained solid was dissolved in 700 µL of $d_6$-DMSO. The samples from DMSO solution were diluted x100 in PBS +0.1%SDS and concentration was determined spectroscopically with Nanodrop. Determined stock concentration was 5.4 mM which constitutes to 76% yield (2.7 mg).

$^1$H NMR (400 MHz, $d_6$-DMSO) δ 9.58 (t, *J* = 5.8 Hz, 1H), 7.79 (dd, *J* = 7.5, 1.0 Hz, 1H), 7.70 (t, *J* = 7.6 Hz, 1H), 7.48 (d, *J* = 8.3 Hz, 2H), 7.42 (d, *J* = 8.3 Hz, 2H), 7.09 (s, 2H), 7.06 (dd, *J* = 7.7, 1.0 Hz, 1H), 6.74 (d, *J* = 2.3 Hz, 2H), 6.46 (d, *J* = 8.6 Hz, 2H), 6.36 (dd, *J* = 8.7, 2.3 Hz, 2H), 6.13 (s, 1H), 5.83 (q, *J* = 5.0 Hz, 2H), 5.30 (s, 2H), 4.58 (d, *J* = 5.7 Hz, 2H), 2.68 (d, *J* = 4.9 Hz, 6H), 1.74 (s, 3H), 1.64 (s, 3H).

ESI-MS, positive mode: m/z = 675.3 $[M+H]^{+\cdot}$

HRMS (ESI) calcd for $C_{38}H_{36}ClN_6O_4$ $[M+H]^+$ 675.2481, found 675.2468.

**4-610CP-CP** (**34**):

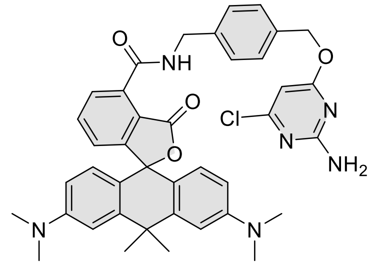

Compound was purified by preparative HPLC (solvent A: $H_2O$ + 10mM $NH_4COOH$ pH = 3.6, solvent B: acetonitrile; temperature 25 °C, gradient A:B - 3 min 80:20 isocratic, 4-20 min 80:20 to 0:100 gradient, 20-25 min 0:100 isocratic, 40 mL/min flow). Fractions containing the product were collected, evaporated and lyophilized from acetonitrile water mixture. The obtained solid was dissolved in 700 µL of $d_6$-DMSO. The samples from DMSO solution were diluted x100 in PBS +0.1%SDS and concentration was determined spectroscopically with Nanodrop. Determined stock concentration was 4.6 mM which constitutes to 65% yield (2.1 mg)

$^1$H NMR (400 MHz, $d_6$-DMSO) δ 9.58 (t, *J* = 5.8 Hz, 1H), 7.80 (dd, *J* = 7.5, 1.0 Hz, 1H), 7.70 (t, *J* = 7.6 Hz, 1H), 7.49 (d, *J* = 8.2 Hz, 2H), 7.44 (d, *J* = 8.3 Hz, 2H), 7.10 (s, 2H), 7.05 (dd, *J* = 7.7, 1.0 Hz, 1H), 6.95 – 6.86 (m, 2H), 6.69 – 6.48 (m, 4H), 6.14 (s, 1H), 5.32 (s, 2H), 4.60 (d, *J* = 5.8 Hz, 2H), 2.94 (s, 12H), 1.82 (s, 3H), 1.72 (s, 3H).

ESI-MS, positive mode: m/z = 703.3 $[M+H]^{+\cdot}$

HRMS (ESI) calcd for $C_{40}H_{40}ClN_6O_4$ $[M+H]^+$ 703.2794, found 703.2788.

**4-625CP-CP** (**35**):

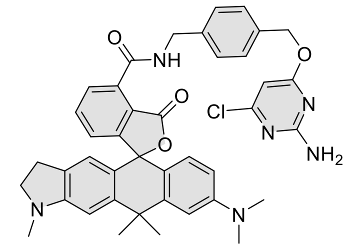

Compound was purified by preparative HPLC (solvent A: $H_2O$ + 10mM $NH_4COOH$ pH = 3.6, solvent B: acetonitrile; temperature 25 °C, gradient A:B - 3 min 80:20 isocratic, 4-20 min 80:20 to 0:100 gradient, 20-25 min 0:100 isocratic, 40 mL/min flow). Fractions containing the product were collected, evaporated and lyophilized from acetonitrile water mixture. The obtained solid was dissolved in 700 µL of $d_6$-DMSO. The samples from DMSO solution were diluted x100 in PBS +0.1%SDS and concentration was determined spectroscopically with Nanodrop. Determined stock concentration was 3.8 mM which constitutes to 53% yield (1.9 mg)

$^1$H NMR (400 MHz, $d_6$-DMSO) δ 9.59 (t, *J* = 5.8 Hz, 1H), 7.80 (dd, *J* = 7.5, 1.0 Hz, 1H), 7.70 (t, *J* = 7.6 Hz, 1H), 7.50 (d, *J* = 8.1 Hz, 2H), 7.44 (d, *J* = 8.1 Hz, 2H), 7.10 (s, 2H), 7.05 (dd, *J* = 7.7, 0.9 Hz, 1H), 6.89 (d, *J* = 2.1 Hz, 1H), 6.76 (s, 1H), 6.58 – 6.53 (m, 2H), 6.39 (s, 1H), 6.14 (s, 1H), 5.32 (s, 2H), 4.60 (d, *J* = 5.8 Hz, 2H), 3.28 – 3.20 (m, 2H), 2.93 (s, 6H), 2.79 (s, 3H), 2.77 – 2.66 (m, 2H), 1.79 (s, 3H), 1.71 (s, 3H).

ESI-MS, positive mode: m/z = 715.2 $[M+H]^+$.

HRMS (ESI) calcd for $C_{41}H_{40}ClN_6O_4$ $[M+H]^+$ 715.2794, found 715.2782.

**4-630CP-CP** (**36**):

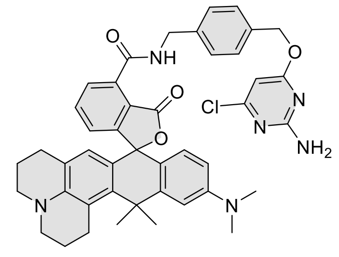

Compound was purified by preparative HPLC (solvent A: $H_2O$ + 10mM $NH_4COOH$ pH = 3.6, solvent B: acetonitrile; temperature 25 °C, gradient A:B - 3 min 80:20 isocratic, 4-20 min 80:20 to 0:100 gradient, 20-25 min 0:100 isocratic , 40 mL/min flow). Fractions containing the product were collected, evaporated and lyophilized from acetonitrile water mixture. The obtained solid was dissolved in 700 µL of $d_6$-DMSO. The samples from DMSO solution were diluted x100 in PBS +0.1%SDS and concentration was determined spectroscopically with Nanodrop. Determined stock concentration was 3.1 mM which constitutes to 44% yield (1.7 mg)

$^1$H NMR (400 MHz, $d_6$-DMSO) δ 9.62 (t, *J* = 5.8 Hz, 1H), 7.79 (dd, *J* = 7.5, 1.0 Hz, 1H), 7.68 (t, *J* = 7.6 Hz, 1H), 7.50 (d, *J* = 8.3 Hz, 2H), 7.44 (d, *J* = 8.3 Hz, 2H), 7.10 (s, 2H), 7.01 (dd, *J* = 7.7, 1.0 Hz, 1H), 6.79 (d, *J* = 2.5 Hz, 1H), 6.56 (dd, *J* = 8.9, 2.5 Hz, 1H), 6.50 (d, *J* = 8.9 Hz, 1H), 6.17 (s, 1H), 6.15 (s, 1H), 5.32 (s, 2H), 4.60 (d, *J* = 5.8 Hz, 2H), 3.20 – 3.12 (m, 4H), 2.98 – 2.93 (m, 2H), 2.92 (s, 6H), 2.47 – 2.36 (m, 2H), 1.95 – 1.89 (m, 2H), 1.89 (s, 3H), 1.83 (s, 3H), 1.79 – 1.72 (m, 2H).

ESI-MS, positive mode: m/z = 755.3 $[M+H]^+$.

HRMS (ESI) calcd for $C_{44}H_{44}ClN_6O_4$ $[M+H]^+$ 755.3107, found 755.3100.

**4-640CP-CP** (**37**):

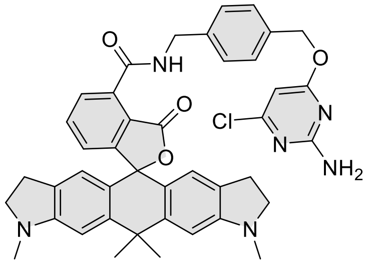

Compound was purified by preparative HPLC (solvent A: $H_2O$ + 10mM $NH_4COOH$ pH = 3.6, solvent B: acetonitrile; temperature 25 °C, gradient A:B - 3 min 80:20 isocratic, 4-20 min 80:20 to 0:100 gradient, 20-25 min 0:100 isocratic , 40 mL/min flow). Fractions containing the product were collected, evaporated and lyophilized from acetonitrile water mixture. The obtained solid was dissolved in 700 µL of $d_6$-DMSO. The samples from DMSO solution were diluted x100 in PBS +0.1%SDS and concentration was determined spectroscopically with Nanodrop. Determined stock concentration was 4.6 mM which constitutes to 65% yield (2.4 mg).

$^1H$ NMR (400 MHz, $d_6$-DMSO) δ 9.61 (t, *J* = 5.8 Hz, 1H), 7.79 (dd, *J* = 7.5, 1.0 Hz, 1H), 7.70 (t, *J* = 7.6 Hz, 1H), 7.50 (d, *J* = 8.2 Hz, 2H), 7.44 (d, *J* = 8.2 Hz, 2H), 7.10 (s, 2H), 7.05 (dd, *J* = 7.7, 0.9 Hz, 1H), 6.74 (s, 2H), 6.36 (s, 2H), 6.14 (s, 1H), 5.32 (s, 2H), 4.60 (d, *J* = 5.8 Hz, 2H), 3.27 – 3.18 (m, 4H), 2.78 (s, 6H), 2.75 – 2.60 (m, 4H), 1.76 (s, 3H), 1.69 (s, 3H).

ESI-MS, positive mode: m/z = 727.3 $[M+H]^+$.

HRMS (ESI) calcd for $C_{42}H_{40}ClN_6O_4$ $[M+H]^+$ 727.2794, found 727.2786.

**4-642CP-CP** (**38**):

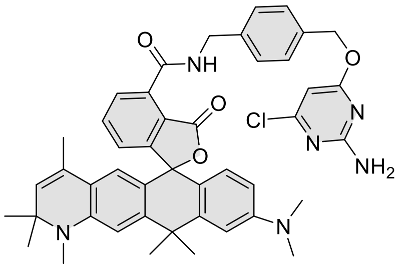

Compound was purified by preparative HPLC (solvent A: $H_2O$ + 10mM $NH_4COOH$ pH = 3.6, solvent B: acetonitrile; temperature 25 °C, gradient A:B - 3 min 30:70 isocratic, 4-20 min 30:70 to 0:100 gradient, 20-25 min 0:100 isocratic, 40 mL/min flow). Fractions containing the product were collected, evaporated and lyophilized from acetonitrile water mixture. The obtained solid was dissolved in 700 µL of $d_6$-DMSO. The samples from DMSO solution were diluted x100 in PBS +0.1%SDS and concentration was determined spectroscopically with Nanodrop. Determined stock concentration was 5.1 mM which constitutes to 72% yield (2.7 mg).

$^1H$ NMR (400 MHz, $d_6$-DMSO) δ 9.62 (t, *J* = 5.8 Hz, 1H), 7.85 (dd, *J* = 7.5, 1.0 Hz, 1H), 7.74 (t, *J* = 7.6 Hz, 1H), 7.49 (d, *J* = 8.2 Hz, 2H), 7.43 (d, *J* = 8.4 Hz, 2H), 7.15 – 7.07 (m, 3H), 6.91 (d, *J* = 2.3 Hz, 1H), 6.67 (s,

1H), 6.60 – 6.54 (m, 2H), 6.18 (s, 1H), 6.14 (s, 1H), 5.34 – 5.29 (m, 3H), 4.60 (t, *J* = 5.9 Hz, 2H), 2.94 (s, 6H), 2.84 (s, 3H), 1.81 (s, 3H), 1.71 (s, 3H), 1.55 (d, *J* = 1.4 Hz, 3H), 1.26 (d, *J* = 4.4 Hz, 6H).

ESI-MS, positive mode: m/z = 769.3 $[M+H]^+$.

HRMS (ESI) calcd for $C_{45}H_{46}ClN_6O$ $[M+H]^+$ 769.3264, found 769.3261.

**4-645CP-CP** (**39**):

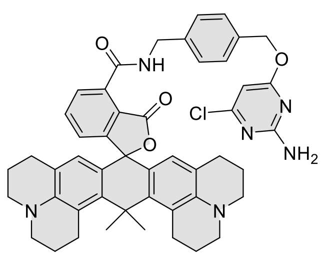

Compound was purified by preparative HPLC (solvent A: $H_2O$ + 10 mM $NH_4COOH$ pH = 3.6, solvent B: acetonitrile; temperature 25 °C, gradient A:B - 3 min 40:60 isocratic, 4-20 min 40:60 to 0:100 gradient, 20-25 min 0:100 isocratic, 40 mL/min flow). Fractions containing the product were collected, evaporated and lyophilized from acetonitrile water mixture. The obtained solid was dissolved in 700 µL of $d_6$-DMSO. The samples from DMSO solution were diluted x100 in PBS +0.1%SDS and concentration was determined spectroscopically with Nanodrop. Determined stock concentration was 2.9 mM which constitutes to 40% yield (1.6 mg).

$^1H$ NMR (400 MHz, $d_6$-DMSO) δ 9.66 (t, *J* = 5.8 Hz, 1H), 7.79 (dd, *J* = 7.5, 1.0 Hz, 1H), 7.66 (d, *J* = 7.6 Hz, 1H), 7.50 (d, *J* = 8.3 Hz, 2H), 7.44 (d, *J* = 8.2 Hz, 2H), 7.10 (s, 2H), 6.98 (dd, *J* = 7.7, 1.0 Hz, 1H), 6.14 (s, 1H), 6.06 (s, 2H), 5.32 (s, 2H), 4.60 (d, *J* = 5.8 Hz, 2H), 3.19 – 3.10 (m, 8H), 2.84 (dd, *J* = 7.3, 4.2 Hz, 4H), 2.47 – 2.34 (m, 4H), 1.98 (s, 3H), 1.92 (s, 3H), 1.89 (dd, *J* = 6.8, 4.9 Hz, 4H), 1.74 (tq, *J* = 9.4, 5.9, 5.5 Hz, 5H).

ESI-MS, positive mode: m/z = 807.3 $[M+H]^+$.

HRMS (ESI) calcd for $C_{48}H_{48}ClN_6O_4$ $[M+H]^+$ 807.3420, found 807.3399.

**4-SiR-CP** (**40**):

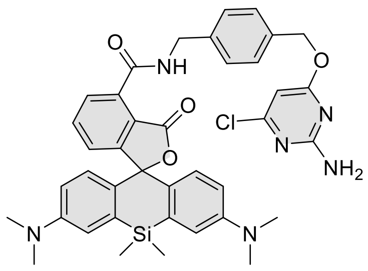

Compound was purified by preparative HPLC (solvent A: $H_2O$ + 10 mM $NH_4COOH$ pH = 3.6, solvent B: acetonitrile; temperature 25 °C, gradient A:B - 3 min 40:60 isocratic, 4-20 min 40:60 to 0:100 gradient, 20-25 min 0:100 isocratic, 40 mL/min flow). Fractions containing the product were collected, evaporated and lyophilized from acetonitrile water mixture. The obtained solid was dissolved in 700 µL of $d_6$-DMSO. The samples from DMSO solution were diluted x100 in PBS +0.1%SDS and concentration was determined spectroscopically with Nanodrop. Determined stock concentration was 4.3 mM which constitutes to 61% yield (2.2 mg).

$^{1}$H NMR (400 MHz, $d_6$-DMSO) δ 9.50 – 9.43 (m, 1H), 7.83 – 7.75 (m, 2H), 7.47 (d, *J* = 8.3 Hz, 2H), 7.42 (d, *J* = 8.5 Hz, 2H), 7.28 (dd, *J* = 5.6, 3.1 Hz, 1H), 7.09 (s, 4H), 6.73 (q, *J* = 8.6 Hz, 4H), 6.14 (s, 1H), 5.31 (s, 2H), 4.56 (d, *J* = 5.8 Hz, 2H), 2.95 (s, 12H), 0.63 (s, 3H), 0.53 (s, 3H).

ESI-MS, positive mode: m/z = 719.3 $[M+H]^+$.

HRMS (ESI) calcd for $C_{39}H_{40}ClN_6O_4Si$ $[M+H]^+$ 719.2563, found 719.2543.

**4-665SiR-CP** (**41**):

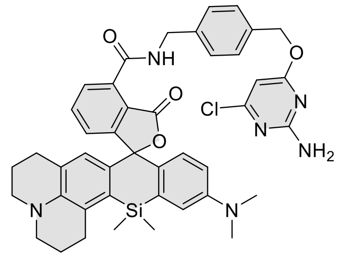

Compound was purified by preparative HPLC (solvent A: $H_2O$ + 10 mM $NH_4COOH$ pH = 3.6, solvent B: acetonitrile; temperature 25 °C, gradient A:B - 3 min 30:70 isocratic, 4-20 min 30:70 to 0:100 gradient, 20-25 min 0:100 isocratic, 40 mL/min flow). Fractions containing the product were collected, evaporated and lyophilized from acetonitrile water mixture. The obtained solid was dissolved in 700 µL of $d_6$-DMSO. The samples from DMSO solution were diluted x100 in PBS +0.1%SDS and concentration was determined spectroscopically with Nanodrop. Determined stock concentration was 4.3 mM which constitutes to 61% yield (2.2 mg).

$^{1}$H NMR (400 MHz, $d_6$-DMSO) δ 9.51 (t, *J* = 5.8 Hz, 1H), 7.74 – 7.67 (m, 2H), 7.46 (d, *J* = 8.3 Hz, 2H), 7.41 (d, *J* = 8.3 Hz, 2H), 7.12 (dd, *J* = 7.2, 1.5 Hz, 1H), 7.09 (s, 2H), 6.92 (d, *J* = 2.7 Hz, 1H), 6.68 (d, *J* = 9.0 Hz, 1H), 6.63 (dd, *J* = 9.1, 2.7 Hz, 1H), 6.33 (s, 1H), 6.13 (s, 1H), 5.30 (s, 2H), 4.56 (d, *J* = 5.8 Hz, 2H), 3.15 (t, *J* = 5.8 Hz, 2H), 3.10 (t, *J* = 5.5 Hz, 3H), 2.89 (s, 8H), 2.46 – 2.38 (m, 2H), 2.01 – 1.86 (m, 4H), 1.74 (q, *J* = 6.0 Hz, 4H), 0.65 (s, 3H), 0.54 (s, 3H).

ESI-MS, positive mode: m/z = 771.2 $[M+H]^+$.

HRMS (ESI) calcd for $C_{43}H_{44}ClN_6O_4Si$ $[M+H]^+$ 771.2876, found 771.2865.

**4-670SiR-CP** (**42**):

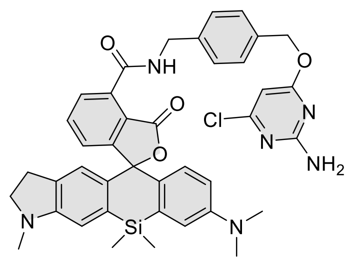

Compound was purified by preparative HPLC (solvent A: $H_2O$ + 10 mM $NH_4COOH$ pH = 3.6, solvent B: acetonitrile; temperature 25 °C, gradient A:B - 3 min 30:70 isocratic, 4-20 min 30:70 to 0:100 gradient, 20-25 min 0:100 isocratic, 40 mL/min flow). Fractions containing the product were collected, evaporated and lyophilized from acetonitrile water mixture. The obtained solid was dissolved in 700 µL of $d_6$-DMSO. The samples from DMSO solution were diluted x100 in PBS +0.1%SDS and concentration was determined spectroscopically with Nanodrop. Determined stock concentration was 4.7 mM which constitutes to 66% yield (2.3 mg).

$^1$H NMR (400 MHz, *d*$_6$-DMSO) δ 9.49 (t, *J* = 5.8 Hz, 1H), 7.78 – 7.70 (m, 2H), 7.46 (d, *J* = 8.3 Hz, 2H), 7.41 (d, *J* = 8.2 Hz, 2H), 7.20 (dd, *J* = 6.6, 2.1 Hz, 1H), 7.09 (s, 2H), 6.96 (d, *J* = 2.8 Hz, 1H), 6.79 (s, 1H), 6.70 (d, *J* = 9.0 Hz, 1H), 6.62 (dd, *J* = 9.0, 2.8 Hz, 1H), 6.58 (s, 1H), 6.13 (s, 1H), 5.30 (s, 2H), 4.56 (d, *J* = 5.8 Hz, 2H), 3.24 – 3.15 (m, 2H), 2.89 (s, 6H), 2.81 – 2.65 (m, 5H), 0.58 (s, 3H), 0.49 (s, 3H).

ESI-MS, positive mode: m/z = 731.3 $[M+H]^+$.

HRMS (ESI) calcd for $C_{40}H_{40}ClN_6O_4Si$ $[M+H]^+$ 731.2563, found 731.2552.

**4-685SiR-CP (43):**

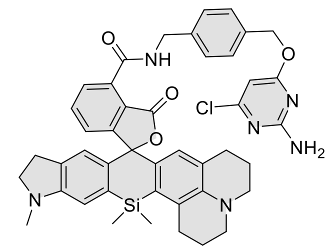

Compound was purified by preparative HPLC (solvent A: $H_2O$ + 10 mM $NH_4COOH$ pH = 3.6, solvent B: acetonitrile; temperature 25 °C, gradient A:B - 3 min 40:60 isocratic, 4-20 min 40:60 to 0:100 gradient, 20-25 min 0:100 isocratic, 40 mL/min flow). Fractions containing the product were collected, evaporated and lyophilized from acetonitrile water mixture. The obtained solid was dissolved in 700 µL of d$_6$-DMSO. The samples from DMSO solution were diluted x100 in PBS +0.1%SDS and concentration was determined spectroscopically with Nanodrop. Determined stock concentration was 3.5 mM which constitutes to 49% yield (1.9 mg).

$^1$H NMR (400 MHz, *d*$_6$-DMSO) δ 9.52 (t, *J* = 5.8 Hz, 1H), 7.72 (dd, *J* = 7.5, 1.3 Hz, 1H), 7.68 (t, *J* = 7.5 Hz, 1H), 7.47 (d, *J* = 8.2 Hz, 2H), 7.42 (d, *J* = 8.2 Hz, 2H), 7.12 – 7.05 (m, 3H), 6.73 (s, 1H), 6.55 (s, 1H), 6.32 (s, 1H), 6.13 (s, 1H), 5.30 (s, 2H), 4.56 (d, *J* = 5.8 Hz, 2H), 3.23 – 3.08 (m, 6H), 2.89 (t, *J* = 6.4 Hz, 2H), 2.84 – 2.58 (m, 7H), 2.47 – 2.36 (m, 2H), 1.94 (p, *J* = 6.1 Hz, 3H), 1.77 – 1.70 (m, 2H), 0.63 (s, 3H), 0.52 (s, 3H).

ESI-MS, positive mode: m/z = 783.3 $[M+H]^+$.

HRMS (ESI) calcd for $C_{44}H_{44}ClN_6O_4Si$ $[M+H]^+$ 783.2876, found 783.2866.

**4-690SiR-CP** (**44**):

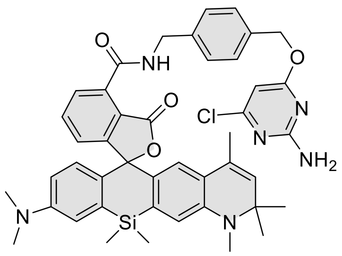

Compound was purified by preparative HPLC (solvent A: $H_2O$ + 10 mM $NH_4COOH$ pH = 3.6, solvent B: acetonitrile; temperature 25 °C, gradient A:B - 3 min 30:70 isocratic, 4-20 min 30:70 to 0:100 gradient, 20-25 min 0:100 isocratic, 40 mL/min flow). Fractions containing the product were collected, evaporated and lyophilized from acetonitrile water mixture. The obtained solid was dissolved in 700 µL of $d_6$-DMSO. The samples from DMSO solution were diluted x100 in PBS +0.1%SDS and concentration was determined spectroscopically with Nanodrop. Determined stock concentration was 2.9 mM which constitutes to 41% yield (1.6 mg).

$^1$H NMR (400 MHz, $d_6$-DMSO) δ 9.48 (t, *J* = 5.7 Hz, 1H), 7.86 – 7.79 (m, 2H), 7.46 (d, *J* = 8.4 Hz, 2H), 7.41 (d, *J* = 8.5 Hz, 2H), 7.39 – 7.35 (m, 1H), 7.10 (s, 2H), 7.01 (d, *J* = 2.8 Hz, 1H), 6.76 (s, 1H), 6.70 (d, *J* = 8.9 Hz, 1H), 6.61 (dd, *J* = 9.0, 2.8 Hz, 1H), 6.41 (s, 1H), 6.14 (s, 1H), 5.34 (d, *J* = 1.5 Hz, 1H), 5.30 (s, 2H), 4.61 – 4.51 (m, 2H), 2.91 (s, 6H), 2.81 (s, 3H), 1.57 (d, *J* = 1.4 Hz, 3H), 1.24 (d, *J* = 4.1 Hz, 7H), 0.63 (s, 3H), 0.51 (s, 3H).

ESI-MS, positive mode: m/z = 785.3 $[M+H]^+$.

HRMS (ESI) calcd for $C_{44}H_{46}ClN_6O_4Si$ $[M+H]^+$ 785.3033, found 785.3025.

**4-700SiR-CP** (**45**):

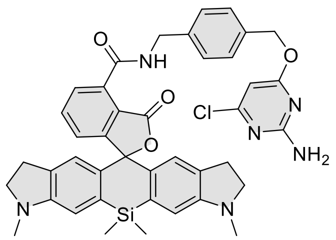

Compound was purified by preparative HPLC (solvent A: $H_2O$ + 10 mM $NH_4COOH$ pH = 3.6, solvent B: acetonitrile; temperature 25 °C, gradient A:B - 3 min 30:70 isocratic, 4-20 min 30:70 to 0:100 gradient, 20-25 min 0:100 isocratic, 40 mL/min flow). Fractions containing the product were collected, evaporated and lyophilized from acetonitrile water mixture. The obtained solid was dissolved in 700 µL of $d_6$-DMSO. The samples from DMSO solution were diluted x100 in PBS +0.1%SDS and concentration was determined spectroscopically with Nanodrop. Determined stock concentration was 3.3 mM which constitutes to 46% yield (1.7 mg).

$^1$H NMR (400 MHz, $d_6$-DMSO) δ 9.52 (t, *J* = 5.8 Hz, 1H), 7.78 – 7.71 (m, 2H), 7.49 (d, *J* = 8.2 Hz, 2H), 7.43 (d, *J* = 8.2 Hz, 2H), 7.17 (dd, *J* = 7.1, 1.6 Hz, 1H), 7.10 (s, 2H), 6.78 (s, 2H), 6.59 (s, 2H), 6.14 (s, 1H), 5.32 (s, 2H), 4.58 (d, *J* = 5.8 Hz, 2H), 3.25 – 3.18 (m, 4H), 2.76 (s, 10H), 0.57 (s, 3H), 0.49 (s, 3H).

ESI-MS, positive mode: m/z = 743.3 $[M+H]^+$.

HRMS (ESI) calcd for $C_{41}H_{40}ClN_6O_4Si$ $[M+H]^+$ 743.2563, found 743.2551.

**4-720SiR-CP** (**46**):

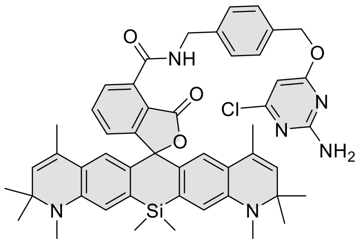

Compound was purified by preparative HPLC (solvent A: $H_2O$ + 10 mM $NH_4COOH$ pH = 3.6, solvent B: acetonitrile; temperature 25 °C, gradient A:B - 3 min 30:70 isocratic, 4-20 min 30:70 to 0:100 gradient, 20-25 min 0:100 isocratic, 40 mL/min flow). Fractions containing the product were collected, evaporated and lyophilized from acetonitrile water mixture. The obtained solid was dissolved in 700 µL of $d_6$-DMSO. The samples from DMSO solution were diluted x100 in PBS +0.1%SDS and concentration was determined spectroscopically with Nanodrop. Determined stock concentration was 4.9 mM which constitutes to 69% yield (2.9 mg).

$^1$H NMR (400 MHz, $d_6$-DMSO) δ 9.48 (t, $J$ = 5.8 Hz, 1H), 7.88 – 7.83 (m, 2H), 7.48 (t, $J$ = 4.3 Hz, 1H), 7.45 (d, $J$ = 8.5 Hz, 2H), 7.40 (d, $J$ = 8.4 Hz, 2H), 7.09 (s, 2H), 6.77 (s, 2H), 6.40 (s, 2H), 6.13 (s, 1H), 5.33 (d, $J$ = 1.5 Hz, 2H), 5.30 (s, 2H), 4.55 (d, $J$ = 5.8 Hz, 2H), 2.81 (s, 6H), 1.56 (d, $J$ = 1.4 Hz, 6H), 1.24 (d, $J$ = 2.4 Hz, 12H), 0.63 (s, 3H), 0.50 (s, 3H).

ESI-MS, positive mode: m/z = 851.3 $[M+H]^+$.

HRMS (ESI) calcd for $C_{49}H_{52}ClN_6O_4Si$ $[M+H]^+$ 851.3502, found 851.3495.

General procedure for the synthesis of compounds **47**-**67**:

Into a solution of corresponding rhodamine dye (5 µmol, 1 eq; **2-17** and **22-26**) and DIPEA (5 µL) in DMSO (100 µL) a solution of HATU (6.5 µmol, 1.3 eq) in DMSO (100 µL) was added and the mixture was mixed for 1 min. Then a solution of **CTX-C8-NH₂**[20] (7.5 µmol ,1.5 eq) in DMSO (100 µL) was added at once. Reactions were usually over after 30 min and their course was monitored by LC/MS analysis. Once reaction was finished it was quenched with 20 µL of formic acid and diluted with water and acetonitrile to 2 mL volume and was further purified by the means of preparative HPLC (preparative column: Agilent 5 Prep-C18, 5 µm, 100 x 50 mm).

**4-CFL-CTX** (**47**):

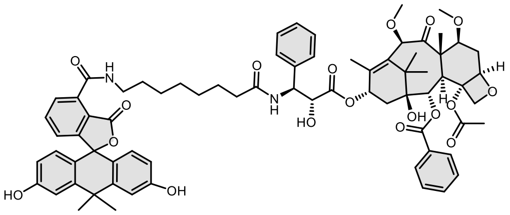

Compound was purified by preparative HPLC (solvent A: $H_2O$ +0.2% HCOOH, solvent B: acetonitrile; temperature 25 °C, gradient A:B - 3 min 70:30 isocratic, 4-20 min 70:30 to 0:100 gradient, 20-25 min 0:100 isocratic, 40 mL/min flow). Fractions containing the product were collected, evaporated and lyophilized from acetonitrile water mixture to obtain product in 38% yield (2.4 mg).

$^1$H NMR (400 MHz, $d_6$-DMSO) δ 9.72 (br s, 2H), 9.01 (t, *J* = 5.6 Hz, 1H), 8.39 (d, *J* = 9.1 Hz, 1H), 8.02 – 7.96 (m, 2H), 7.76 (dd, *J* = 7.5, 1.0 Hz, 1H), 7.71 (d, *J* = 7.6 Hz, 1H), 7.69 – 7.65 (m, 1H), 7.59 (t, *J* = 7.5 Hz, 2H), 7.41 – 7.30 (m, 4H), 7.21 (t, *J* = 7.1 Hz, 1H), 7.08 – 7.06 (m, 2H), 6.65 – 6.53 (m, 4H), 5.96 (dd, *J* = 8.5, 4.2 Hz, 2H), 5.39 (d, *J* = 7.2 Hz, 1H), 5.29 (dd, *J* = 9.0, 5.8 Hz, 1H), 4.96 (dd, *J* = 9.9, 2.0 Hz, 1H), 4.71 (s, 1H), 4.65 (s, 1H), 4.43 (t, *J* = 6.2 Hz, 1H), 4.02 (s, 2H), 3.76 (dd, *J* = 10.7, 6.5 Hz, 1H), 3.63 (d, *J* = 7.2 Hz, 1H), 3.38 – 3.35 (m, overlapped with water peak, 2H), 3.30 (s, overlapped with water peak, 3H), 3.21 (s, 3H), 2.68 – 2.63 (m, 1H), 2.25 (s, 3H), 2.18 (t, *J* = 7.3 Hz, 2H), 2.01 – 1.78 (m, 6H), 1.73 (s, 3H), 1.64 (s, 3H), 1.61 – 1.41 (m, 8H), 1.31 (m, 6H), 1.03 (s, 3H), 0.97 (s, 3H).

ESI-MS, positive mode: m/z = 1283.5 $[M+Na]^+$.

HRMS (ESI) calcd for $C_{72}H_{80}N_2O_{18}Na$ $[M+Na]^+$ 1283.5298, found 1283.5292.

**4-505R-CTX** (**48**):

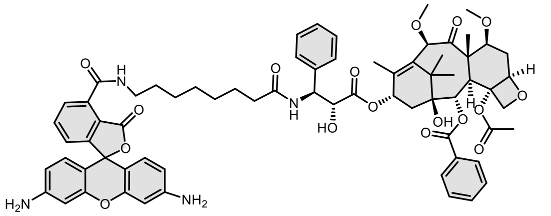

Compound was purified by preparative HPLC (solvent A: 10mM $NH_4COOH$ pH = 3.6, solvent B: acetonitrile; temperature 25 °C, gradient A:B - 3 min 70:30 isocratic, 4-20 min 70:30 to 0:100 gradient, 20-25 min 0:100 isocratic, 40 mL/min flow). Fractions containing the product were collected, evaporated and lyophilized from acetonitrile water mixture. The obtained solid was dissolved in 700 µL of $d_6$-DMSO. The samples from DMSO solution were diluted x100 in PBS +0.1%SDS and concentration was determined spectroscopically with Nanodrop. Determined stock concentration was 2.1 mM which constitutes to 29% yield (1.8 mg).

$^1$H NMR (600 MHz, DMSO-$d_6$) δ 9.06 (t, *J* = 5.6 Hz, 1H), 8.43 (d, *J* = 9.0 Hz, 1H), 7.99 – 7.97 (m, 2H), 7.82 (d, *J* = 7.4 Hz, 1H), 7.76 (t, *J* = 7.6 Hz, 1H), 7.68 (t, *J* = 7.7 Hz, 1H), 7.61 – 7.58 (m, 2H), 7.37 (t, *J* = 7.5 Hz, 2H), 7.32 (d, *J* = 7.6 Hz, 2H), 7.25 – 7.19 (m, 2H), 6.43 (d, *J* = 8.5 Hz, 2H), 6.38 (d, *J* = 2.2 Hz, 2H), 6.29

(dd, $J$ = 8.5, 2.2 Hz, 2H), 6.03 (s, 1H), 5.94 (t, $J$ = 9.2 Hz, 1H), 5.59 (s, 4H), 5.39 (d, $J$ = 7.1 Hz, 1H), 5.28 (dd, $J$ = 9.0, 5.9 Hz, 1H), 4.96 (d, $J$ = 9.6 Hz, 1H), 4.70 (s, 1H), 4.65 (s, 1H), 4.42 (d, $J$ = 6.0 Hz, 1H), 4.02 (s, 2H), 3.75 (dd, $J$ = 10.6, 6.6 Hz, 1H), 3.63 (d, $J$ = 7.1 Hz, 1H), 3.38 – 3.37 (m, overlapped with water peak, 2H), 3.30 (s, overlapped with water peak, 3H), 3.21 (s, 3H), 2.68 – 2.63 (m, 1H), 2.24 (s, 3H), 2.17 (t, $J$ = 7.2 Hz, 2H), 1.98 – 1.94 (m, 1H), 1.88 – 1.83 (m, 4H), 1.57 – 1.49 (m, 8H), 1.37 – 1.25 (m, 6H), 1.03 (s, 3H), 0.97 (s, 3H).

ESI-MS, positive mode: m/z = 1233.3 $[M+H]^+$.

HRMS (ESI) calcd for $C_{69}H_{77}N_4O_{17}$ $[M+H]^+$ 1233.5278, found 1233.5272.

**4-525R-CTX** (**49**):

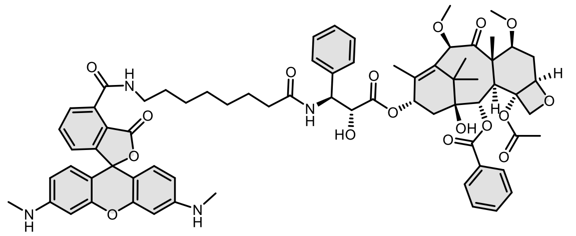

Compound was purified by preparative HPLC (solvent A: $H_2O$ 10mM $NH_4COOH$ pH = 3.6, solvent B: acetonitrile; temperature 25 °C, gradient A:B - 3 min 70:30 isocratic, 4-20 min 70:30 to 0:100 gradient, 20-25 min 0:100 isocratic, 40 mL/min flow). Fractions containing the product were collected, evaporated and lyophilized from acetonitrile water mixture. The obtained solid was dissolved in 700 µL of $d_6$-DMSO. The samples from DMSO solution were diluted x100 in PBS +0.1%SDS and concentration was determined spectroscopically with Nanodrop. Determined stock concentration was 2.6 mM which constitutes to 37% yield (2.3 mg).

$^1$H NMR (600 MHz, $d_6$-DMSO) δ 9.07 (t, $J$ = 5.6 Hz, 1H), 8.43 (d, $J$ = 2.9 Hz, 1H), 7.98 (d, $J$ = 7.6 Hz, 2H), 7.83 (d, $J$ = 7.5 Hz, 1H), 7.76 (t, $J$ = 7.6 Hz, 1H), 7.68 (t, $J$ = 7.7 Hz, 1H), 7.62 – 7.58 (m, 2H), 7.36 (t, $J$ = 7.5 Hz, 2H), 7.32 (d, $J$ = 7.7 Hz, 2H), 7.24 – 7.18 (m, 2H), 6.48 (d, $J$ = 8.5 Hz, 2H), 6.43 – 6.23 (m, 4H), 6.18 (q, $J$ = 5.0 Hz, 2H), 6.03 (br s, 1H), 5.94 (t, $J$ = 9.4 Hz, 1H), 5.39 (d, $J$ = 7.1 Hz, 1H), 5.28 (dd, $J$ = 9.0, 5.9 Hz, 1H), 4.95 (d, $J$ = 9.7 Hz, 1H), 4.70 (s, 1H), 4.65 (s, 1H), 4.42 (s, 1H), 4.02 (s, 2H), 3.75 (dd, $J$ = 10.7, 6.6 Hz, 1H), 3.63 (d, $J$ = 7.1 Hz, 1H), 3.37 – 3.37 (m, overlapped with water peak, 2H), 3.30 (s, overlapped with water peak, 3H), 3.21 (s, 3H), 2.69 (d, $J$ = 4.9 Hz, 6H), 2.66 – 2.60 (m, 1H), 2.24 (s, 3H), 2.17 (t, $J$ = 7.6 Hz, 2H), 1.98 – 1.82 (m, 5H), 1.57 – 1.48 (m, 8H), 1.38 – 1.26 (m, 6H), 1.03 (s, 3H), 0.97 (s, 3H).

ESI-MS, positive mode: m/z = 1261.6 $[M+H]^+$.

HRMS (ESI) calcd for $C_{71}H_{81}N_4O_{17}$ $[M+H]^+$ 1261.5591, found 1261.5587.

**4-DMRhodol-CTX** (**50**):

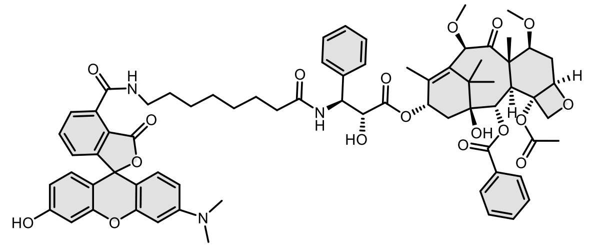

Compound was purified by preparative HPLC (solvent A: $H_2O$ +0.2% HCOOH, solvent B: acetonitrile; temperature 25 °C, gradient A:B - 3 min 70:30 isocratic, 4-20 min 70:30 to 0:100 gradient, 20-25 min 0:100 isocratic, 40 mL/min flow). Fractions containing the product were collected, evaporated and lyophilized from acetonitrile water mixture to obtain product in 46% yield (2.9 mg).

$^1$H NMR (400 MHz, $d_6$-DMSO) δ 10.12 (s, 1H), 9.05 (t, $J$ = 5.5 Hz, 1H), 8.39 (d, $J$ = 9.0 Hz, 1H), 8.00 – 7.95 (m, 2H), 7.83 (dd, $J$ = 7.5, 1.0 Hz, 1H), 7.75 (t, $J$ = 7.6 Hz, 1H), 7.68 (t, $J$ = 7.3 Hz, 1H), 7.59 (t, $J$ = 7.5 Hz, 2H), 7.38 – 7.30 (m, 4H), 7.25 (dd, $J$ = 7.6, 1.0 Hz, 1H), 7.23 – 7.19 (m, 1H), 6.67 (d, $J$ = 8.7 Hz, 1H), 6.65 – 6.60 (m, 2H), 6.55 – 6.48 (m, 3H), 5.99 – 5.92 (m, 2H), 5.39 (d, $J$ = 7.2 Hz, 1H), 5.28 (dd, $J$ = 9.0, 5.8 Hz, 1H), 4.95 (d, $J$ = 9.4 Hz, 1H), 4.70 (s, 1H), 4.65 (s, 1H), 4.42 (t, $J$ = 6.4 Hz, 1H), 4.02 (s, 2H), 3.75 (dd, $J$ = 10.6, 6.6 Hz, 1H), 3.63 (d, $J$ = 7.1 Hz, 1H), 3.31 – 3.27 (m, 5H), 3.21 (s, 3H), 2.95 (s, 6H), 2.69 – 2.64 (m, 1H), 2.24 (s, 3H), 2.17 (t, $J$ = 7.4 Hz, 2H), 2.00 – 1.94 (m, 1H), 1.89 – 1.80 (m, 4H), 1.62 – 1.45 (m, 8H), 1.40 – 1.25 (m, 6H), 1.03 (s, 3H), 0.97 (s, 3H).

ESI-MS, positive mode: m/z = 1262.5 $[M+H]^+$.

HRMS (ESI) calcd for $C_{71}H_{80}N_3O_{18}$ $[M+H]^+$ 1262.5431, found 1262.5426.

**4-TMR-CTX** (**51**):

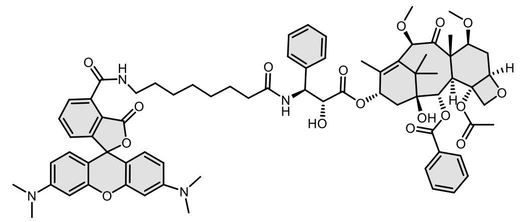

Compound was purified by preparative HPLC (solvent A: $H_2O$ +0.2% HCOOH, solvent B: MeOH; temperature 25 °C, gradient A:B - 3 min 70:30 isocratic, 4-20 min 70:30 to 0:100 gradient, 20-25 min 0:100 isocratic, 40 mL/min flow). Fractions containing the product were collected, evaporated and lyophilized from acetonitrile water mixture. The obtained solid was dissolved in 700 µL of $d_6$-DMSO. The samples from DMSO solution were diluted x100 in PBS +0.1%SDS and concentration was determined spectroscopically with Nanodrop. Determined stock concentration was 3.4 mM which constitutes to 48% yield (3.1 mg).

$^1$H NMR (400 MHz, $d_6$-DMSO) δ 9.06 (t, $J$ = 5.5 Hz, 1H), 8.38 (d, $J$ = 9.1 Hz, 1H), 7.98 (d, $J$ = 7.0 Hz, 2H), 7.84 (dd, $J$ = 7.5, 1.1 Hz, 1H), 7.76 (t, $J$ = 7.6 Hz, 1H), 7.70 – 7.64 (m, 1H), 7.59 (t, $J$ = 7.5 Hz, 2H), 7.35

(ddd, *J* = 15.1, 8.2, 6.8 Hz, 4H), 7.24 – 7.19 (m, 2H), 6.61 (dd, *J* = 9.4, 1.6 Hz, 2H), 6.53 – 6.45 (m, 4H), 5.94 (dd, *J* = 8.1, 5.4 Hz, 2H), 5.39 (d, *J* = 7.0 Hz, 1H), 5.32 – 5.28 (m, 1H), 4.95 (d, *J* = 9.1 Hz, 1H), 4.70 (s, 1H), 4.65 (s, 1H), 4.43 (dd, *J* = 6.9, 5.8 Hz, 1H), 4.02 (s, 2H), 3.75 (dd, *J* = 10.9, 6.4 Hz, 1H), 3.63 (d, *J* = 7.1 Hz, 1H), 3.31 – 3.28 (m, 5H), 3.21 (s, 3H), 2.94 (s, 12H), 2.69 – 2.61 (m, 1H), 2.25 (s, 3H), 2.18 (t, *J* = 7.8 Hz, 2H), 1.96 – 1.81 (m, 5H), 1.53 (q, *J* = 6.7, 5.7 Hz, 8H), 1.30 – 1.26 (m, 6H), 1.03 (s, 3H), 0.97 (s, 3H).

ESI-MS, positive mode: m/z = 1289.6 $[M+H]^+$.

HRMS (ESI) calcd for $C_{73}H_{85}N_4O_{17}$ $[M+H]^+$ 1289.5904, found 1289.5919.

**4-DAllR-CTX (52):**

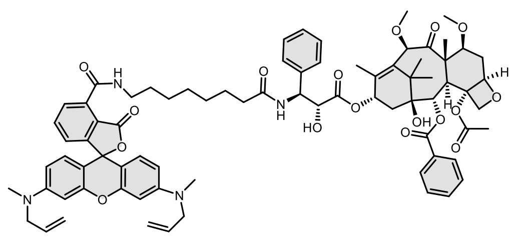

Compound was purified by preparative HPLC (solvent A: $H_2O$ +0.2% HCOOH, solvent B: MeOH; temperature 25 °C, gradient A:B - 3 min 70:30 isocratic, 4-20 min 70:30 to 0:100 gradient, 20-25 min 0:100 isocratic, 40 mL/min flow). Fractions containing the product were collected, evaporated and lyophilized from acetonitrile water mixture. The obtained solid was dissolved in 700 µL of $d_6$-DMSO. The samples from DMSO solution were diluted x100 in PBS +0.1%SDS and concentration was determined spectroscopically with Nanodrop. Determined stock concentration was 3.9 mM which constitutes to 55% yield (3.8 mg).

$^1$H NMR (400 MHz, DMSO-$d_6$) δ 9.05 (t, *J* = 5.6 Hz, 1H), 8.38 (d, *J* = 9.0 Hz, 1H), 8.01 – 7.95 (m, 2H), 7.84 (dd, *J* = 7.6, 1.1 Hz, 1H), 7.76 (t, *J* = 7.6 Hz, 1H), 7.68 (tt, *J* = 7.4, 1.5 Hz, 1H), 7.59 (t, *J* = 7.5 Hz, 2H), 7.39 – 7.31 (m, 4H), 7.26 (dd, *J* = 7.7, 1.0 Hz, 1H), 7.21 (tt, *J* = 7.2, 1.5 Hz, 1H), 6.60 – 6.55 (m, 2H), 6.50 – 6.45 (m, 4H), 5.94 (t, *J* = 8.3 Hz, 2H), 5.87 – 5.76 (m, 2H), 5.39 (d, *J* = 7.2 Hz, 1H), 5.29 (dd, *J* = 9.0, 5.8 Hz, 1H), 5.14 – 5.07 (m, 4H), 4.95 (dd, *J* = 9.7, 2.0 Hz, 1H), 4.70 (s, 1H), 4.65 (s, 1H), 4.43 (t, *J* = 5.8 Hz, 1H), 4.02 (s, 2H), 3.98 (d, *J* = 4.9 Hz, 4H), 3.75 (dd, *J* = 10.6, 6.5 Hz, 1H), 3.63 (d, *J* = 7.2 Hz, 1H), 3.39 – 3.36 (m, 2H), 3.30 (s, 3H), 3.21 (s, 3H), 2.95 (s, 6H), 2.70 – 2.61 (m, 1H), 2.25 (s, 3H), 2.18 (t, *J* = 7.3 Hz, 2H), 2.00 – 1.86 (m, 2H), 1.83 (s, 3H), 1.57 – 1.48 (m, 8H), 1.39 – 1.26 (m, 6H), 1.03 (s, 3H), 0.97 (s, 3H).

ESI-MS, positive mode: m/z = 1341.6 $[M+H]^+$.

HRMS (ESI) calcd for $C_{77}H_{89}N_4O_{17}$ $[M+H]^+$ 1341.6217, found 1341.6212.

**4-580R-CTX** (**53**):

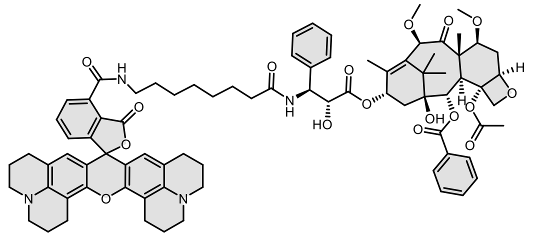

Compound was purified by preparative HPLC (solvent A: $H_2O$ 10mM $NH_4COOH$ pH = 3.6, solvent B: MeOH; temperature 25 °C, gradient A:B - 3 min 60:40 isocratic, 4-20 min 60:40 to 0:100 gradient, 20-25 min 0:100 isocratic, 40 mL/min flow). Fractions containing the product were collected, evaporated and lyophilized from acetonitrile water mixture. The obtained solid was dissolved in 700 µL of $d_6$-DMSO. The samples from DMSO solution were diluted x100 in PBS +0.1% SDS and concentration was determined spectroscopically with Nanodrop. Determined stock concentration was 3.4 mM which constitutes to 23% yield (1.6 mg).

$^1$H NMR (400 MHz, $d_6$-DMSO) δ 9.18 (t, *J* = 5.5 Hz, 1H), 8.40 (d, *J* = 9.0 Hz, 1H), 8.00 – 7.94 (m, 2H), 7.85 (dd, *J* = 7.6, 0.9 Hz, 1H), 7.73 (t, *J* = 7.6 Hz, 1H), 7.66 (t, *J* = 7.3 Hz, 1H), 7.58 (t, *J* = 7.5 Hz, 2H), 7.39 – 7.27 (m, 4H), 7.24 (dd, *J* = 7.7, 1.0 Hz, 1H), 7.19 (t, *J* = 7.1 Hz, 1H), 6.10 (s, 2H), 5.99 (d, *J* = 7.0 Hz, 1H), 5.93 (t, *J* = 9.2 Hz, 1H), 5.37 (d, *J* = 7.2 Hz, 1H), 5.26 (dd, *J* = 9.0, 5.7 Hz, 1H), 4.94 (d, *J* = 10.9 Hz, 1H), 4.69 (s, 1H), 4.63 (s, 1H), 4.40 (t, *J* = 6.1 Hz, 1H), 4.01 (s, 2H), 3.74 (dd, *J* = 10.6, 6.6 Hz, 1H), 3.62 (d, *J* = 7.2 Hz, 1H), 3.35 – 3.33 (m, 2H), 3.29 (s, 3H), 3.20 (s, 3H), 3.16 (t, *J* = 5.6 Hz, 2H), 3.11 (t, *J* = 5.6 Hz, 2H), 2.84 (t, *J* = 6.6 Hz, 2H), 2.64 – 2.59 (m, 1H), 2.46 – 2.38 (m, 2H), 2.23 (s, 3H), 2.16 (t, *J* = 7.2 Hz, 2H), 2.02 – 1.89 (m, 5H), 1.88 – 1.66 (m, 8H), 1.59 – 1.41 (m, 8H), 1.40 – 1.17 (m, 14H), 1.02 (s, 3H), 0.96 (s, 3H).

ESI-MS, positive mode: m/z = 1393.7 $[M+H]^+$.

HRMS (ESI) calcd for $C_{81}H_{93}N_4O_{17}$ $[M+H]^+$ 1393.6530, found 1393.6527.

**4-580CP-CTX** (**54**):

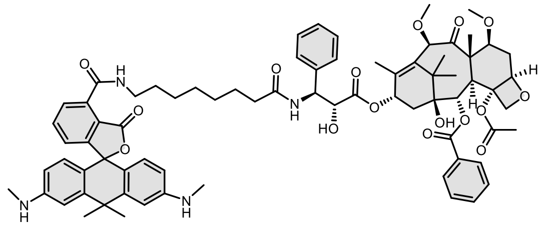

Compound was purified by preparative HPLC (solvent A: $H_2O$ 10mM $NH_4COOH$ pH = 3.6, solvent B: MeOH; temperature 25 °C, gradient A:B - 3 min 60:40 isocratic, 4-20 min 60:40 to 0:100 gradient, 20-25 min 0:100 isocratic, 40 mL/min flow). Fractions containing the product were collected, evaporated and lyophilized from acetonitrile water mixture. The obtained solid was dissolved in 700 µL of $d_6$-DMSO. The samples from DMSO solution were diluted x100 in PBS +0.1% SDS and concentration was determined spectroscopically with Nanodrop. Determined stock concentration was 3.8 mM which constitutes to 53% yield (3.4 mg).

$^1$H NMR (400 MHz, $d_6$-DMSO) δ 9.13 (t, *J* = 5.5 Hz, 1H), 8.38 (d, *J* = 9.1 Hz, 1H), 8.01 – 7.94 (m, 2H), 7.81 – 7.77 (m, 1H), 7.72 – 7.65 (m, 2H), 7.59 (t, *J* = 7.4 Hz, 2H), 7.40 – 7.29 (m, 4H), 7.21 (t, *J* = 7.1 Hz, 1H),

7.05 (dd, *J* = 7.7, 1.0 Hz, 1H), 6.76 (d, *J* = 2.3 Hz, 2H), 6.45 (d, *J* = 8.6 Hz, 2H), 6.38 (dd, *J* = 8.7, 2.3 Hz, 2H), 5.98 – 5.91 (m, 2H), 5.85 (d, *J* = 5.1 Hz, 2H), 5.39 (d, *J* = 7.1 Hz, 1H), 5.32 – 5.27 (m, 1H), 4.95 (d, *J* = 9.4 Hz, 1H), 4.71 (s, 1H), 4.65 (s, 1H), 4.43 (t, *J* = 6.4 Hz, 1H), 4.02 (s, 2H), 3.75 (dd, *J* = 10.6, 6.5 Hz, 1H), 3.63 (d, *J* = 7.1 Hz, 1H), 3.35 – 3.33 (m, 2H), 3.30 (s, 3H), 3.21 (s, 3H), 2.70 (d, *J* = 4.7 Hz, 6H), 2.66 – 2.62 (m, 1H), 2.25 (s, 3H), 2.17 (t, *J* = 7.5 Hz, 2H), 2.00 – 1.94 (m, 1H), 1.90 – 1.80 (m, 4H), 1.75 (s, 3H), 1.65 (s, 3H), 1.61 – 1.44 (m, 8H), 1.39 – 1.25 (m, 6H), 1.03 (s, 3H), 0.97 (s, 3H).

ESI-MS, positive mode: m/z = 1287.6 $[M+H]^+$. .

HRMS (ESI) calcd for $C_{74}H_{87}N_4O_{16}$ $[M+H]^+$ 1287.6112, found 1287.6094.

**4-610CP-CTX** (**55**):

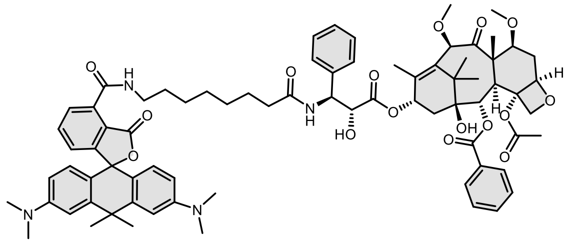

Compound was purified by preparative HPLC (solvent A: 10mM $NH_4COOH$ pH = 3.6, solvent B: MeOH; temperature 25 °C, gradient A:B - 3 min 60:40 isocratic, 4-20 min 60:40 to 0:100 gradient, 20-25 min 0:100 isocratic, 40 mL/min flow). Fractions containing the product were collected, evaporated and lyophilized from acetonitrile water mixture. The obtained solid was dissolved in 700 µL of $d_6$-DMSO. The samples from DMSO solution were diluted x100 in PBS +0.1% SDS and concentration was determined spectroscopically with Nanodrop. Determined stock concentration was 3.4 mM which constitutes to 48% yield (3.1 mg). Analysis data conforms to previously published data[20].

$^1$H NMR (400 MHz, $d_6$-DMSO) δ 9.09 (t, *J* = 5.5 Hz, 1H), 8.37 (d, *J* = 9.0 Hz, 1H), 7.98 – 7.91 (m, 2H), 7.75 (dd, *J* = 7.5, 1.0 Hz, 1H), 7.68 – 7.61 (m, 2H), 7.61 – 7.51 (m, 2H), 7.36 – 7.26 (m, 4H), 7.18 (tt, *J* = 7.1, 1.6 Hz, 1H), 7.01 (dd, *J* = 7.7, 1.0 Hz, 1H), 6.88 (d, *J* = 2.0 Hz, 2H), 6.60 – 6.49 (m, 4H), 6.00 – 5.87 (m, 2H), 5.36 (d, *J* = 7.1 Hz, 1H), 5.26 (dd, *J* = 9.1, 5.8 Hz, 1H), 4.92 (dd, *J* = 9.6, 2.1 Hz, 1H), 4.68 (s, 1H), 4.62 (s, 1H), 4.40 (t, *J* = 6.2 Hz, 1H), 3.99 (s, 2H), 3.72 (dd, *J* = 10.6, 6.6 Hz, 1H), 3.60 (d, *J* = 7.1 Hz, 1H), 3.34 – 3.31 (m, 2H), 3.27 (s, 3H), 3.18 (s, 3H), 2.91 (s, 12H), 2.69 – 2.57 (m, 1H), 2.21 (s, 3H), 2.15 (t, *J* = 7.3 Hz, 2H), 1.97 – 1.91 (m, 1H), 1.88 – 1.82 (m, 1H), 1.80 (s, 3H), 1.79 (s, 3H), 1.69 (s, 3H), 1.56 – 1.43 (m, 8H), 1.37 – 1.23 (m, 6H), 1.00 (s, 3H), 0.94 (s, 3H).

ESI-MS, positive mode: m/z = 1315.6 $[M+H]^+$. .

HRMS (ESI) calcd for $C_{76}H_{91}N_4O_{16}$ $[M+H]^+$ 1315.6425, found 1315.6409.

**4-625CP-CTX** (**56**):

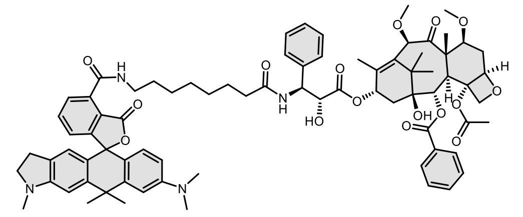

Compound was purified by preparative HPLC (solvent A: 10mM $NH_4COOH$ pH = 3.6, solvent B: MeOH; temperature 25 °C, gradient A:B - 3 min 60:40 isocratic, 4-20 min 60:40 to 0:100 gradient, 20-25 min 0:100 isocratic, 40 mL/min flow). Fractions containing the product were collected, evaporated and lyophilized from acetonitrile water mixture. The obtained solid was dissolved in 700 µL of $d_6$-DMSO. The samples from DMSO solution were diluted x100 in PBS +0.1% SDS and concentration was determined spectroscopically with Nanodrop. Determined stock concentration was 2.8 mM which constitutes to 39% yield (2.6 mg).

$^1$H NMR (400 MHz, $d_6$-DMSO) δ 9.13 (t, *J* = 5.6 Hz, 1H), 8.41 (d, *J* = 9.4 Hz, 1H), 8.00 – 7.94 (m, 2H), 7.78 (dd, *J* = 7.6, 1.0 Hz, 1H), 7.68 (td, *J* = 7.5, 1.9 Hz, 2H), 7.59 (t, *J* = 7.5 Hz, 2H), 7.40 – 7.29 (m, 4H), 7.21 (t, *J* = 7.2 Hz, 1H), 7.03 (dd, *J* = 7.7, 1.0 Hz, 1H), 6.89 (d, *J* = 2.4 Hz, 1H), 6.76 (s, 1H), 6.57 (dd, *J* = 9.0, 2.3 Hz, 1H), 6.53 (d, *J* = 8.8 Hz, 1H), 6.37 (s, 1H), 5.96 (t, *J* = 13.8 Hz, 2H), 5.39 (d, *J* = 7.1 Hz, 1H), 5.29 (dd, *J* = 9.0, 5.8 Hz, 1H), 4.95 (d, *J* = 9.2 Hz, 1H), 4.71 (s, 1H), 4.65 (s, 1H), 4.43 (s, 1H), 4.02 (s, 2H), 3.76 (dd, *J* = 10.6, 6.6 Hz, 1H), 3.63 (d, *J* = 7.1 Hz, 1H), 3.35 (s, 2H), 3.30 (s, 3H), 3.27 – 3.23 (m, 2H), 3.21 (s, 3H), 2.93 (s, 6H), 2.79 (s, 3H), 2.76 – 2.64 (m, 3H), 2.24 (s, 3H), 2.18 (t, *J* = 7.4 Hz, 2H), 1.98 – 1.94 (m, 1H), 1.92 – 1.82 (m, 4H), 1.79 (s, 3H), 1.70 (s, 3H), 1.65 – 1.48 (m, 8H), 1.41 – 1.27 (m, 6H), 1.03 (s, 3H), 0.97 (s, 3H).

ESI-MS, positive mode: m/z = 1327.6 $[M+H]^+$.

HRMS (ESI) calcd for $C_{77}H_{91}N_4O_16$ $[M+H]^+$ 1327.6425, found 1327.6433.

**4-630CP-CTX** (**57**):

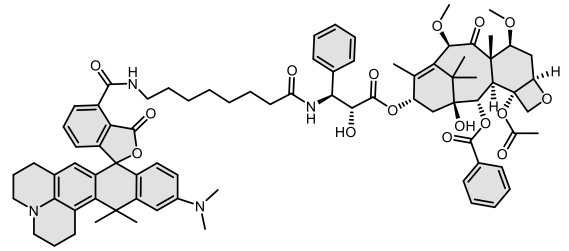

Compound was purified by preparative HPLC (solvent A: 10mM $NH_4COOH$ pH = 3.6, solvent B: MeOH; temperature 25 °C, gradient A:B - 3 min 50:50 isocratic, 4-20 min 50:50 to 0:100 gradient, 20-25 min 0:100 isocratic, 40 mL/min flow). Fractions containing the product were collected, evaporated and lyophilized from acetonitrile water mixture. The obtained solid was dissolved in 700 µL of $d_6$-DMSO. The samples from DMSO solution were diluted x100 in PBS +0.1% SDS and concentration was determined spectroscopically with Nanodrop. Determined stock concentration was 1.9 mM which constitutes to 27% yield (1.8 mg).

$^1$H NMR (400 MHz, $d_6$-DMF) δ 9.69 (t, *J* = 5.4 Hz, 1H), 8.46 (d, *J* = 9.2 Hz, 1H), 8.15 – 8.07 (m, 3H), 7.79 (t, *J* = 7.6 Hz, 1H), 7.71 (t, *J* = 7.4 Hz, 1H), 7.62 (t, *J* = 7.5 Hz, 2H), 7.50 (d, *J* = 7.3 Hz, 2H), 7.41 (t, *J* = 7.7

Hz, 2H), 7.27 (t, *J* = 7.3 Hz, 1H), 7.15 (dd, *J* = 7.7, 1.0 Hz, 1H), 6.93 (d, *J* = 2.5 Hz, 1H), 6.65 – 6.61 (m, 1H), 6.58 (d, *J* = 8.8 Hz, 1H), 6.31 (s, 1H), 6.17 (t, *J* = 9.2 Hz, 1H), 6.06 (d, *J* = 5.9 Hz, 1H), 5.60 (d, *J* = 7.1 Hz, 1H), 5.56 (dd, *J* = 9.1, 5.1 Hz, 1H), 5.03 (s, 1H), 4.87 (s, 1H), 4.68 (t, *J* = 4.9 Hz, 1H), 4.13 (q, *J* = 8.1 Hz, 2H), 3.92 (dd, *J* = 10.6, 6.5 Hz, 1H), 3.82 (d, *J* = 7.1 Hz, 1H), 3.48 – 3.45 (m, 2H), 3.41 (s, 3H), 3.31 (s, 3H), 3.27 – 3.15 (m, 4H), 3.09 – 3.03 (m, 1H), 3.00 (s, 6H), 2.54 – 2.38 (m, 5H), 2.34 – 2.08 (m, 6H), 2.07 – 1.91 (m, 8H), 1.91 (s, 3H), 1.82 – 1.75 (m, 2H), 1.72 – 1.52 (m, 8H), 1.45 – 1.29 (m, 6H), 1.19 (s, 3H), 1.15 (s, 3H).

ESI-MS, positive mode: m/z = 1367.7 $[M+H]^+$.

HRMS (ESI) calcd for $C_{80}H_{95}N_4O_{16}$ $[M+H]^+$ 1367.6738, found 1367.6745.

**4-640CP-CTX** (**58**):

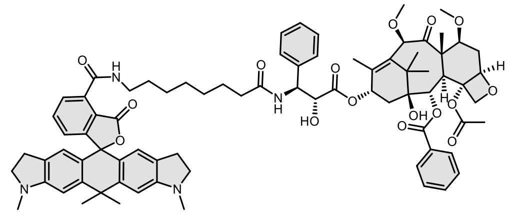

Compound was purified by preparative HPLC (solvent A: 10mM $NH_4COOH$ pH = 3.6, solvent B: MeOH; temperature 25 °C, gradient A:B - 3 min 50:50 isocratic, 4-20 min 50:50 to 0:100 gradient, 20-25 min 0:100 isocratic, 40 mL/min flow). Fractions containing the product were collected, evaporated and lyophilized from acetonitrile water mixture. The obtained solid was dissolved in 700 µL of $d_6$-DMSO. The samples from DMSO solution were diluted x100 in PBS +0.1% SDS and concentration was determined spectroscopically with Nanodrop. Determined stock concentration was 2.8 mM which constitutes to 39% yield (2.6 mg).

$^1$H NMR (400 MHz, $d_6$-DMSO) δ 9.15 (t, *J* = 5.6 Hz, 1H), 8.45 (d, *J* = 9.0 Hz, 1H), 8.00 – 7.96 (m, 2H), 7.77 (dd, *J* = 7.5, 1.0 Hz, 1H), 7.68 (t, *J* = 7.6 Hz, 2H), 7.59 (dd, *J* = 8.2, 6.7 Hz, 2H), 7.35 (ddd, *J* = 14.8, 8.2, 6.7 Hz, 4H), 7.23 – 7.18 (m, 1H), 7.03 (dd, *J* = 7.7, 1.0 Hz, 1H), 6.74 (s, 2H), 6.34 (s, 2H), 6.07 (d, *J* = 6.7 Hz, 1H), 5.95 (t, *J* = 9.0 Hz, 1H), 5.39 (d, *J* = 7.1 Hz, 1H), 5.28 (dd, *J* = 9.0, 5.9 Hz, 1H), 4.95 (dd, *J* = 9.6, 2.1 Hz, 1H), 4.70 (s, 1H), 4.65 (s, 1H), 4.43 (t, *J* = 5.7 Hz, 1H), 4.02 (s, 2H), 3.76 (dd, *J* = 10.6, 6.6 Hz, 1H), 3.63 (d, *J* = 7.1 Hz, 1H), 3.34 – 3.33 (m, 2H), 3.30 (s, 3H), 3.27 – 3.17 (m, 7H), 2.78 (s, 6H), 2.75 – 2.60 (m, 5H), 2.24 (s, 3H), 2.18 (t, *J* = 7.4 Hz, 2H), 1.98 – 1.83 (m, 5H), 1.76 (s, 3H), 1.69 (s, 3H), 1.60 – 1.46 (m, 8H), 1.39 – 1.25 (m, 6H), 1.03 (s, 3H), 0.97 (s, 3H).

ESI-MS, positive mode: m/z = 1339.6 $[M+H]^+$.

HRMS (ESI) calcd for $C_{78}H_{91}N_4O_{16}$ $[M+H]^+$ 1339.6425, found 1339.6428.

**4-642CP-CTX** (**59**):

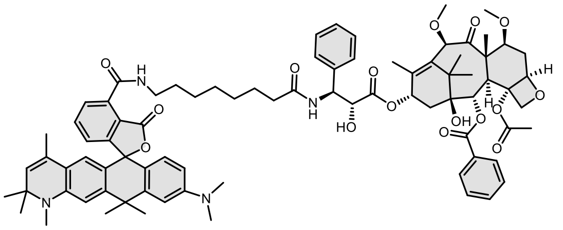

Compound was purified by preparative HPLC (solvent A: 10mM $NH_4COOH$ pH = 3.6, solvent B: MeOH; temperature 25 °C, gradient A:B - 3 min 50:50 isocratic, 4-20 min 50:50 to 0:100 gradient, 20-25 min 0:100 isocratic, 40 mL/min flow). Fractions containing the product were collected, evaporated and lyophilized from acetonitrile water mixture. The obtained solid was dissolved in 700 µL of $d_6$-DMSO. The samples from DMSO solution were diluted x100 in PBS +0.1% SDS and concentration was determined spectroscopically with Nanodrop. Determined stock concentration was 3.4 mM which constitutes to 48% yield (3.3 mg).

$^1$H NMR (400 MHz, $d_6$-DMSO) δ 9.17 (t, *J* = 5.6 Hz, 1H), 8.40 (d, *J* = 9.0 Hz, 1H), 8.01 – 7.94 (m, 2H), 7.85 – 7.82 (m, 1H), 7.76 – 7.63 (m, 3H), 7.59 (t, *J* = 7.4 Hz, 2H), 7.41 – 7.28 (m, 4H), 7.20 (t, *J* = 7.1 Hz, 1H), 7.10 (dd, *J* = 7.7, 1.0 Hz, 1H), 6.91 (s, 1H), 6.67 (s, 1H), 6.56 (d, *J* = 1.3 Hz, 2H), 6.16 (s, 1H), 6.01 – 5.91 (m, 2H), 5.38 (d, *J* = 7.1 Hz, 1H), 5.32 (d, *J* = 1.5 Hz, 1H), 5.28 (dd, *J* = 9.0, 5.8 Hz, 1H), 4.95 (d, *J* = 9.2 Hz, 1H), 4.70 (s, 1H), 4.65 (s, 1H), 4.42 (t, *J* = 4.9 Hz, 1H), 4.02 (s, 2H), 3.75 (dd, *J* = 10.6, 6.6 Hz, 2H), 3.63 (d, *J* = 7.1 Hz, 1H), 3.36 – 3.36 (m, 2H), 3.30 (s, 3H), 3.21 (s, 3H), 2.94 (s, 6H), 2.84 (s, 3H), 2.68 – 2.63 (m, 1H), 2.24 (s, 3H), 2.18 (t, *J* = 7.3 Hz, 2H), 2.02 – 1.77 (m, 8H), 1.70 (s, 3H), 1.69 – 1.39 (m, 11H), 1.37 – 1.19 (m, 12H), 1.03 (s, 3H), 0.97 (s, 3H).

ESI-MS, positive mode: m/z = 1381.7 $[M+H]^+$.

HRMS (ESI) calcd for $C_{81}H_{97}N_4O_{16}$ $[M+H]^+$ 1381.6894, found 1381.6904.

**4-645CP-CTX** (60):

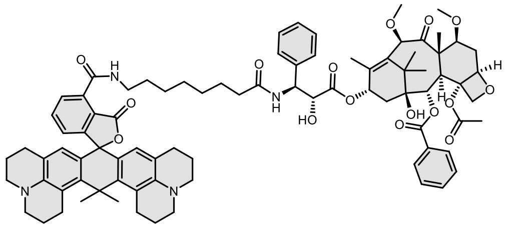

Compound was purified by preparative HPLC (solvent A: 10mM $NH_4COOH$ pH = 3.6, solvent B: MeOH; temperature 25 °C, gradient A:B - 3 min 50:50 isocratic, 4-20 min 50:50 to 0:100 gradient, 20-25 min 0:100 isocratic, 40 mL/min flow). Fractions containing the product were collected, evaporated and lyophilized from acetonitrile water mixture. The obtained solid was dissolved in 700 µL of $d_6$-DMSO. The samples from DMSO solution were diluted x100 in PBS +0.1% SDS and concentration was determined spectroscopically with Nanodrop. Determined stock concentration was 1.6 mM which constitutes to 22% yield (1.6 mg).

$^1$H NMR (400 MHz, $d_6$-DMSO) δ 9.75 (t, *J* = 5.3 Hz, 1H), 8.51 (d, *J* = 8.8 Hz, 1H), 8.13 – 8.08 (m, 2H), 7.77 (t, *J* = 7.6 Hz, 2H), 7.70 (tt, *J* = 7.4, 1.2 Hz, 1H), 7.62 (t, *J* = 7.4 Hz, 2H), 7.50 (d, *J* = 7.1 Hz, 2H), 7.41 (t, *J*

= 7.7 Hz, 2H), 7.27 (tt, *J* = 7.3, 1.2 Hz, 1H), 7.10 (dd, *J* = 7.7, 1.0 Hz, 1H), 6.20 (s, 2H), 6.16 (d, *J* = 9.0 Hz, 2H), 5.60 (d, *J* = 7.1 Hz, 1H), 5.55 (dd, *J* = 9.1, 5.1 Hz, 1H), 5.05 – 4.99 (m, 2H), 4.87 (s, 1H), 4.68 (d, *J* = 5.2 Hz, 1H), 4.16 – 4.09 (m, 2H), 3.92 (dd, *J* = 10.6, 6.6 Hz, 1H), 3.82 (d, *J* = 7.2 Hz, 1H), 3.46 (d, *J* = 6.2 Hz, 2H), 3.41 (s, 3H), 3.31 (s, 3H), 3.27 – 3.06 (m, 8H), 2.90 (s, 16H), 2.61 – 2.32 (m, 9H), 2.32 – 2.22 (m, 3H), 2.18 – 2.13 (m, 1H), 2.06 (s, 3H), 2.00 (s, 3H), 1.98 (d, *J* = 1.4 Hz, 3H), 1.98 – 1.89 (m, 4H), 1.77 (td, *J* = 11.6, 10.8, 5.5 Hz, 4H), 1.62 (d, *J* = 23.5 Hz, 7H), 1.45 – 1.28 (m, 6H), 1.18 (s, 3H), 1.14 (s, 3H).

ESI-MS, positive mode: m/z = 1419.7 $[M+H]^+$.

HRMS (ESI) calcd for $C_{84}H_{99}N_4O_{16}$ $[M+H]^+$ 1419.7051, found 1419.7039.

**4-SiR-CTX** (**61**):

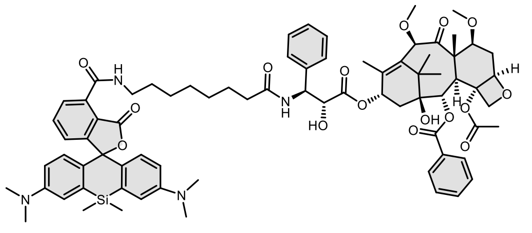

Compound was purified by preparative HPLC (solvent A: $H_2O$ +0.2% HCOOH, solvent B: MeOH; temperature 25 °C, gradient A:B - 3 min 50:50 isocratic, 4-20 min 50:50 to 0:100 gradient, 20-25 min 0:100 isocratic, 40 mL/min flow). Fractions containing the product were collected, evaporated and lyophilized from acetonitrile water mixture.to give product in 55% yield (3.7 mg) as a light blue solid.

$^1$H NMR (400 MHz, *d$_6$*-DMSO) δ 9.02 (t, *J* = 5.5 Hz, 1H), 8.37 (d, *J* = 9.0 Hz, 1H), 7.98 (d, *J* = 6.9 Hz, 2H), 7.77 – 7.74 (m, 2H), 7.69 – 7.64 (m, 1H), 7.59 (t, *J* = 7.5 Hz, 2H), 7.39 – 7.30 (m, 4H), 7.26 – 7.21 (m, 2H), 7.00 (d, *J* = 2.8 Hz, 2H), 6.70 (dd, *J* = 9.0, 1.2 Hz, 2H), 6.64 (ddd, *J* = 8.9, 2.8, 1.4 Hz, 2H), 5.95 (dd, *J* = 8.5, 6.4 Hz, 2H), 5.39 (d, *J* = 7.1 Hz, 1H), 5.29 (dd, *J* = 9.1, 5.8 Hz, 1H), 4.95 (d, *J* = 9.5 Hz, 1H), 4.70 (s, 1H), 4.65 (s, 1H), 4.43 (dd, *J* = 6.9, 5.8 Hz, 1H), 4.02 (s, 2H), 3.75 (dd, *J* = 10.6, 6.5 Hz, 1H), 3.63 (d, *J* = 7.1 Hz, 1H), 3.30 (s, 3H), 3.30 – 3.27 (m, 2H), 3.21 (s, 3H), 2.92 (s, 12H), 2.72 – 2.59 (m, 1H), 2.25 (s, 3H), 2.18 (t, *J* = 7.4 Hz, 2H), 2.01 – 1.93 (m, 1H), 1.90 – 1.84 (m, 1H), 1.83 (s, 3H), 1.57 – 1.46 (m, 8H), 1.37 – 1.20 (m, 6H), 1.03 (s, 3H), 0.97 (s, 3H), 0.62 (s, 3H), 0.52 (s, 3H).

ESI-MS, positive mode: m/z = 1353.6 $[M+Na]^+$.

HRMS (ESI) calcd for $C_{75}H_{90}N_4O_{16}SiNa$ $[M+H]^+$ 1353.6013, found 1353.6003.

**4-665SiR-CTX** (**62**):

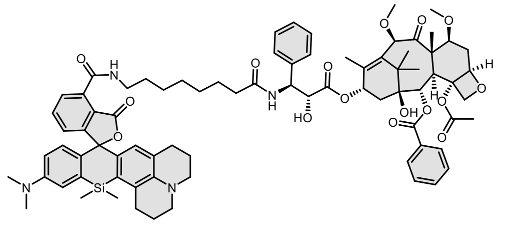

Compound was purified by preparative HPLC (solvent A: 10mM $NH_4COOH$ pH = 3.6, solvent B: MeOH; temperature 25 °C, gradient A:B - 3 min 50:50 isocratic, 4-20 min 50:50 to 0:100 gradient, 20-25 min 0:100 isocratic, 40 mL/min flow). Fractions containing the product were collected, evaporated and lyophilized from acetonitrile water mixture. The obtained solid was dissolved in 700 µL of $d_6$-DMSO. The samples from DMSO solution were diluted x100 in PBS +0.1% SDS and concentration was determined spectroscopically with Nanodrop. Determined stock concentration was 2.4 mM which constitutes to 34% yield (2.3 mg).

$^1$H NMR (400 MHz, $d_6$-DMSO) δ 9.06 (t, *J* = 5.5 Hz, 1H), 8.41 (d, *J* = 9.0 Hz, 1H), 8.01 – 7.93 (m, 2H), 7.75 – 7.63 (m, 3H), 7.59 (t, *J* = 7.4 Hz, 2H), 7.35 (ddd, *J* = 15.2, 8.3, 6.9 Hz, 4H), 7.21 (t, *J* = 7.1 Hz, 1H), 7.13 – 7.09 (m, 1H), 6.93 (d, *J* = 2.6 Hz, 1H), 6.70 – 6.62 (m, 2H), 6.33 (s, 1H), 6.00 (d, *J* = 6.7 Hz, 1H), 5.95 (t, *J* = 9.3 Hz, 2H), 5.39 (d, *J* = 7.1 Hz, 1H), 5.29 (dd, *J* = 9.0, 5.8 Hz, 1H), 4.95 (d, *J* = 9.5 Hz, 1H), 4.70 (s, 1H), 4.65 (s, 1H), 4.43 (t, *J* = 6.1 Hz, 1H), 4.02 (s, 2H), 3.75 (dd, *J* = 10.6, 6.6 Hz, 1H), 3.63 (d, *J* = 7.2 Hz, 1H), 3.31 – 3.24 (m, 5H), 3.21 (s, 3H), 3.16 (t, *J* = 5.8 Hz, 2H), 3.11 (t, *J* = 5.3 Hz, 2H), 3.02 – 2.78 (m, 8H), 2.68 – 2.62 (m, 1H), 2.44 (dd, *J* = 14.8, 8.3 Hz, 2H), 2.24 (s, 3H), 2.18 (t, *J* = 7.3 Hz, 2H), 1.97 – 1.73 (m, 8H), 1.58 – 1.44 (m, 7H), 1.38 – 1.24 (m, 6H), 1.03 (s, 3H), 0.97 (s, 3H), 0.66 (s, 3H), 0.55 (s, 3H).

ESI-MS, positive mode: m/z = 1383.7 $[M+H]^+$.

HRMS (ESI) calcd for $C_{79}H_{95}N_4O_{16}Si$ $[M+H]^+$ 1383.6507, found 1383.6513.

**4-670SiR-CTX** (**63**):

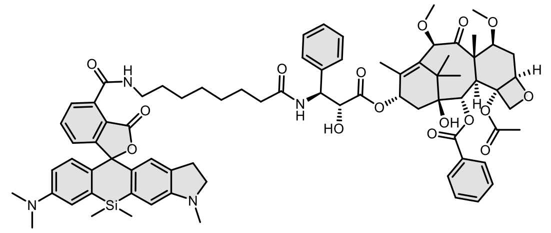

Compound was purified by preparative HPLC (solvent A: $H_2O$ +0.2% HCOOH, solvent B: MeOH; temperature 25 °C, gradient A:B - 3 min 50:50 isocratic, 4-20 min 50:50 to 0:100 gradient, 20-25 min 0:100 isocratic, 40 mL/min flow). Fractions containing the product were collected, evaporated and lyophilized from acetonitrile water mixture to give product in 34% yield (2.3 mg).

$^1$H NMR (400 MHz, $d_6$-DMSO) δ 9.03 (t, *J* = 5.5 Hz, 1H), 8.38 (d, *J* = 9.0 Hz, 1H), 8.01 – 7.94 (m, 2H), 7.77 – 7.71 (m, 2H), 7.67 (tt, *J* = 7.2, 1.4 Hz, 1H), 7.59 (t, *J* = 7.4 Hz, 2H), 7.40 – 7.30 (m, 4H), 7.24 – 7.17 (m,

2H), 6.98 (d, *J* = 2.8 Hz, 1H), 6.80 (s, 1H), 6.70 (dd, *J* = 9.0, 1.0 Hz, 1H), 6.64 (ddd, *J* = 9.1, 2.8, 1.6 Hz, 1H), 6.57 (s, 1H), 5.99 – 5.91 (m, 2H), 5.39 (d, *J* = 7.1 Hz, 1H), 5.29 (dd, *J* = 9.0, 5.8 Hz, 1H), 4.95 (d, *J* = 9.4 Hz, 1H), 4.70 (s, 1H), 4.65 (s, 1H), 4.43 (t, *J* = 6.3 Hz, 1H), 4.02 (s, 2H), 3.75 (dd, *J* = 10.6, 6.6 Hz, 1H), 3.63 (d, *J* = 7.1 Hz, 1H), 3.31 – 3.28 (m, 5H), 3.26 – 3.16 (m, 5H), 2.91 (s, 6H), 2.80 – 2.63 (m, 6H), 2.25 (s, 3H), 2.18 (t, *J* = 7.4 Hz, 2H), 1.97 (d, *J* = 6.3 Hz, 1H), 1.90 – 1.80 (m, 4H), 1.60 – 1.45 (m, 8H), 1.38 – 1.25 (m, 6H), 1.03 (s, 3H), 0.97 (s, 3H), 0.60 (s, 3H), 0.50 (s, 3H).

ESI-MS, positive mode: m/z = 1343.6 $[M+H]^+$.

HRMS (ESI) calcd for $C_{76}H_{91}N_4O_{16}Si$ $[M+H]^+$ 1343.6194, found 1343.6195.

**4-685SiR-CTX** (**64**):

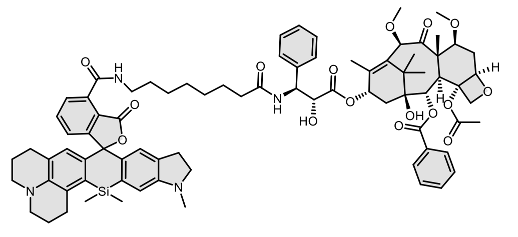

Compound was purified by preparative HPLC (solvent A: $H_2O$ +0.2% HCOOH, solvent B: MeOH; temperature 25 °C, gradient A:B - 3 min 40:60 isocratic, 4-20 min 40:60 to 0:100 gradient, 20-25 min 0:100 isocratic, 40 mL/min flow). Fractions containing the product were collected, evaporated and lyophilized from acetonitrile water mixture. The obtained solid was dissolved in 700 µL of $d_6$-DMSO. The samples from DMSO solution were diluted x100 in PBS +0.1% SDS and concentration was determined spectroscopically with Nanodrop. Determined stock concentration was 2.8 mM which constitutes to 39% yield (2.7 mg).

$^1$H NMR (400 MHz, $d_6$-DMSO) δ 9.06 (t, *J* = 5.5 Hz, 1H), 8.42 (d, *J* = 9.0 Hz, 1H), 8.00 – 7.93 (m, 2H), 7.73 – 7.62 (m, 3H), 7.58 (t, *J* = 7.5 Hz, 2H), 7.39 – 7.26 (m, 4H), 7.19 (tt, *J* = 7.1, 1.4 Hz, 1H), 7.06 (dd, *J* = 7.4, 1.3 Hz, 1H), 6.73 (s, 1H), 6.53 (s, 1H), 6.31 (s, 1H), 6.02 (d, *J* = 4.8 Hz, 1H), 5.96 – 5.90 (m, 1H), 5.37 (d, *J* = 7.1 Hz, 1H), 5.27 (dd, *J* = 8.9, 5.9 Hz, 1H), 4.94 (d, *J* = 9.6 Hz, 1H), 4.69 (s, 1H), 4.63 (s, 1H), 4.41 (t, *J* = 5.0 Hz, 1H), 4.01 (s, 2H), 3.74 (dd, *J* = 10.6, 6.6 Hz, 1H), 3.62 (d, *J* = 7.2 Hz, 1H), 3.29 – 3.28 (m, 5H), 3.22 – 3.08 (m, 9H), 2.89 (t, *J* = 6.3 Hz, 2H), 2.80 – 2.62 (m, 6H), 2.46 – 2.38 (m, 2H), 2.23 (s, 3H), 2.16 (t, *J* = 7.4 Hz, 2H), 1.99 – 1.79 (m, 7H), 1.77 – 1.70 (m, 2H), 1.61 – 1.41 (m, 8H), 1.35 – 1.23 (m, 6H), 1.02 (s, 3H), 0.96 (s, 3H), 0.63 (s, 3H), 0.52 (s, 3H).

ESI-MS, positive mode: m/z = 1395.6 $[M+H]^+$.

HRMS (ESI) calcd for $C_{80}H_{95}N_4O_{16}Si$ $[M+H]^+$ 1395.6507, found 1395.6494.

**4-690SiR-CTX** (**65**):

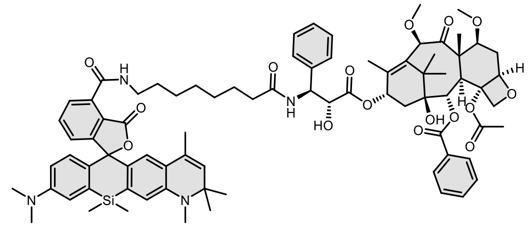

Compound was purified by preparative HPLC (solvent A: $H_2O$ +0.2% HCOOH, solvent B: MeOH; temperature 25 °C, gradient A:B - 3 min 30:70 isocratic, 4-20 min 30:70 to 0:100 gradient, 20-25 min 0:100 isocratic, 40 mL/min flow). Fractions containing the product were collected, evaporated and lyophilized from acetonitrile water mixture to give product in 52% yield (3.6 mg).

$^1$H NMR (400 MHz, $d_6$-DMSO) δ 9.01 (t, *J* = 5.1 Hz, 1H), 8.37 (d, *J* = 9.0 Hz, 1H), 8.01 – 7.95 (m, 2H), 7.83 – 7.76 (m, 2H), 7.69 – 7.64 (m, 1H), 7.59 (t, *J* = 7.4 Hz, 2H), 7.41 – 7.27 (m, 5H), 7.20 (tt, *J* = 7.1, 1.4 Hz, 1H), 7.01 (d, *J* = 2.8 Hz, 1H), 6.76 (s, 1H), 6.69 (dd, *J* = 8.9, 0.9 Hz, 1H), 6.62 (ddd, *J* = 9.1, 2.9, 1.3 Hz, 1H), 6.41 (s, 1H), 5.99 – 5.90 (m, 2H), 5.39 (d, *J* = 7.1 Hz, 1H), 5.34 (d, *J* = 1.5 Hz, 1H), 5.29 (dd, *J* = 9.0, 5.8 Hz, 1H), 4.95 (d, *J* = 9.6 Hz, 1H), 4.70 (s, 1H), 4.65 (s, 1H), 4.42 (t, *J* = 6.3 Hz, 1H), 4.02 (s, 2H), 3.75 (dd, *J* = 10.6, 6.6 Hz, 1H), 3.63 (d, *J* = 7.1 Hz, 1H), 3.31 – 3.24 (m, 5H), 3.21 (s, 3H), 2.91 (s, 6H), 2.81 (s, 3H), 2.69 – 2.62 (m, 1H), 2.25 (s, 3H), 2.17 (t, *J* = 7.4 Hz, 2H), 1.99 – 1.82 (m, 5H), 1.57 (d, *J* = 1.4 Hz, 3H), 1.56 – 1.40 (m, 8H), 1.35 – 1.21 (m, 12H), 1.03 (s, 3H), 0.97 (s, 3H), 0.63 (s, 3H), 0.51 (s, 3H).

ESI-MS, positive mode: m/z = 1397.7 $[M+H]^+$.

HRMS (ESI) calcd for $C_{80}H_{97}N_4O_{16}Si$ $[M+H]^+$ 1397.6663, found 1397.6662.

**4-700SiR-CTX** (**66**):

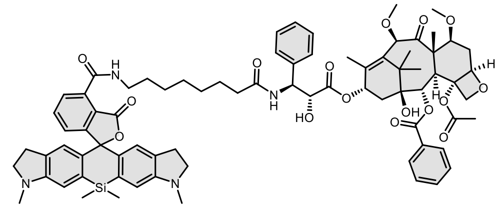

Compound was purified by preparative HPLC (solvent A: $H_2O$ +0.2% HCOOH, solvent B: MeOH; temperature 25 °C, gradient A:B - 3 min 30:70 isocratic, 4-20 min 30:70 to 0:100 gradient, 20-25 min 0:100 isocratic, 40 mL/min flow). Fractions containing the product were collected, evaporated and lyophilized from acetonitrile water mixture. The obtained solid was dissolved in 700 µL of $d_6$-DMSO. The samples from DMSO solution were diluted x100 in PBS +0.1% SDS and concentration was determined spectroscopically with Nanodrop. Determined stock concentration was 3.3 mM which constitutes to 46% yield (3.2 mg).

$^1$H NMR (400 MHz, $d_6$-DMSO) δ 9.03 (t, *J* = 5.5 Hz, 1H), 8.38 (d, *J* = 9.0 Hz, 1H), 7.99 – 7.94 (m, 2H), 7.73 – 7.64 (m, 3H), 7.58 (t, *J* = 7.5 Hz, 2H), 7.33 (ddd, *J* = 15.2, 8.3, 6.9 Hz, 4H), 7.19 (tt, *J* = 7.1, 1.5 Hz, 1H), 7.14 (dd, *J* = 6.8, 1.9 Hz, 1H), 6.76 (s, 2H), 6.56 (s, 2H), 5.98 – 5.90 (m, 2H), 5.37 (d, *J* = 7.1 Hz, 1H), 5.28 (dd, *J* = 9.0, 5.8 Hz, 1H), 4.94 (d, *J* = 9.5 Hz, 1H), 4.69 (s, 1H), 4.63 (s, 1H), 4.41 (t, *J* = 6.3 Hz, 1H), 4.01 (s, 2H),

3.74 (dd, *J* = 10.6, 6.6 Hz, 1H), 3.62 (d, *J* = 7.1 Hz, 1H), 3.30 – 3.25 (m, 5H), 3.24 – 3.14 (m, 7H), 2.83 – 2.63 (m, 11H), 2.23 (s, 3H), 2.16 (t, *J* = 7.3 Hz, 2H), 1.99 – 1.82 (m, 5H), 1.59 – 1.45 (m, 8H), 1.38 – 1.24 (m, 6H), 1.02 (s, 3H), 0.96 (s, 3H), 0.56 (s, 3H), 0.47 (s, 3H).

ESI-MS, positive mode: m/z = 1377.6 $[M+Na]^+$.

HRMS (ESI) calcd for $C_{77}H_{91}N_4O_{16}Si$ $[M+H]^+$ 1355.6194, found 1355.6190.

**4-720SiR-CTX** (**67**):

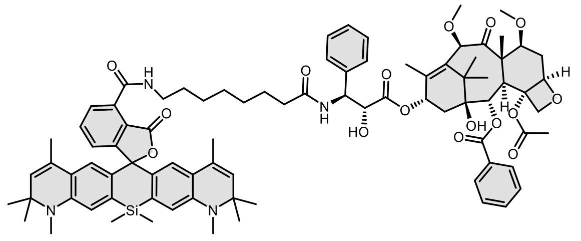

Compound was purified by preparative HPLC (solvent A: $H_2O$ +0.2% HCOOH, solvent B: MeOH; temperature 25 °C, gradient A:B - 3 min 30:70isocratic, 4-20 min 30:70 to 0:100 gradient, 20-25 min 0:100 isocratic, 40 mL/min flow). Fractions containing the product were collected, evaporated and lyophilized from acetonitrile water mixture to give product in 49% yield (3.6 mg).

$^1$H NMR (400 MHz, *d$_6$*-DMSO) δ 9.02 (t, *J* = 5.5 Hz, 1H), 8.39 (d, *J* = 9.0 Hz, 1H), 8.01 – 7.95 (m, 2H), 7.87 – 7.80 (m, 2H), 7.67 (tt, *J* = 7.3, 1.3 Hz, 1H), 7.59 (t, *J* = 7.4 Hz, 2H), 7.44 (t, *J* = 4.4 Hz, 1H), 7.40 – 7.28 (m, 4H), 7.20 (tt, *J* = 7.0, 1.6 Hz, 1H), 6.76 (s, 2H), 6.41 (s, 2H), 5.99 – 5.91 (m, 2H), 5.39 (d, *J* = 7.1 Hz, 1H), 5.34 (d, *J* = 1.5 Hz, 2H), 5.28 (dd, *J* = 9.0, 5.8 Hz, 1H), 4.95 (dd, *J* = 9.6, 1.8 Hz, 1H), 4.70 (s, 1H), 4.65 (s, 1H), 4.42 (t, *J* = 6.2 Hz, 1H), 4.02 (s, 2H), 3.75 (dd, *J* = 10.6, 6.6 Hz, 1H), 3.63 (d, *J* = 7.3 Hz, 1H), 3.31 – 3.24 (m, 5H), 3.21 (s, 3H), 2.81 (s, 6H), 2.69 – 2.64 (m, 1H), 2.24 (s, 3H), 2.19 – 2.13 (m, 2H), 1.98 – 1.82 (m, 5H), 1.57 (d, *J* = 1.1 Hz, 6H), 1.55 – 1.39 (m, 8H), 1.36 – 1.21 (m, 18H), 1.03 (s, 3H), 0.97 (s, 3H), 0.63 (s, 3H), 0.50 (s, 3H).

ESI-MS, positive mode: m/z = 1464.7 $[M+H]^+$.

HRMS (ESI) calcd for $C_{85}H_{103}N_4O_{16}Si$ $[M+H]^+$ 1463.7133, found 1463.7131.

<u>General procedure for the synthesis of compounds **68**-**79**:</u>

Into a solution of corresponding rhodamine dye (5 µmol, 1 eq; **2**, **4** and **7-16**) and DIPEA (5 µL) in DMSO (100 µL) a solution of HATU (6.5 µmol, 1.3 eq) in DMSO (100 µL) was added and the mixture was mixed for 1 min. Then a solution of **Cl-(CH$_2$)$_6$-PEG$_4$-NH$_2$**[22] (7.5 µmol ,1.5 eq) in DMSO (100 µL) was added at once. Reactions were usually over after 30 min and their course was monitored by LC/MS analysis. Once reaction was finished it was quenched with 20 µL of formic acid and diluted with water and acetonitrile to 2 mL volume and was further purified by the means of preparative HPLC (preparative column: Agilent 5 Prep-C18, 5 µm, 100 x 50 mm).

**4-610CP-Halo** (**68**):

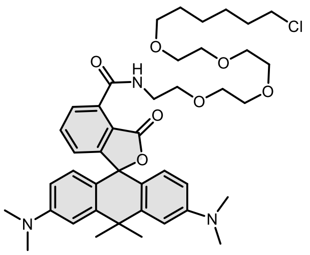

Compound was purified by preparative HPLC (solvent A: $H_2O$ +0.2% HCOOH, solvent B: MeOH; temperature 25 °C, gradient A:B - 3 min 70:30 isocratic, 4-20 min 70:30 to 0:100 gradient, 20-25 min 0:100 isocratic, 40 mL/min flow). Fractions containing the product were collected, evaporated and lyophilized from acetonitrile water mixture. The obtained solid was dissolved in 700 µL of $d_6$-DMSO. The samples from DMSO solution were diluted x100 in PBS +0.1% SDS and concentration was determined spectroscopically with Nanodrop. Determined stock concentration was 4.8 mM which constitutes to 68% yield (2.5 mg).

$^1$H NMR (400 MHz, $CD_3OD$) δ 8.10 (dd, *J* = 7.6, 0.9 Hz, 1H), 7.76 (t, *J* = 7.7 Hz, 1H), 7.14 (dd, *J* = 7.8, 1.0 Hz, 1H), 7.01 – 6.96 (m, 2H), 6.66 – 6.53 (m, 4H), 3.81 – 3.47 (m, 18H), 3.40 (t, *J* = 6.5 Hz, 2H), 2.99 (s, 12H), 1.88 (s, 3H), 1.80 – 1.66 (m, 5H), 1.57 – 1.49 (m, 2H), 1.45 – 1.32 (m, 4H).

ESI-MS, positive mode: m/z = 750.4 $[M+H]^+$.

HRMS (ESI) calcd for $C_{42}H_{57}ClN_3O_7$ $[M+H]^+$ 750.3880, found 750.3885.

**4-625CP-Halo (69)**:

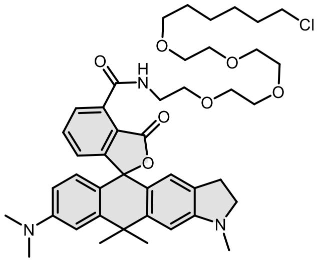

Compound was purified by preparative HPLC (solvent A: $H_2O$ +0.2% HCOOH, solvent B: MeOH; temperature 25 °C, gradient A:B - 3 min 70:30 isocratic, 4-20 min 70:30 to 0:100 gradient, 20-25 min 0:100 isocratic, 40 mL/min flow). Fractions containing the product were collected, evaporated and lyophilized from acetonitrile water mixture. The obtained solid was dissolved in 700 µL of $d_6$-DMSO. The samples from DMSO solution were diluted x100 in PBS +0.1% SDS and concentration was determined spectroscopically with Nanodrop. Determined stock concentration was 4.4 mM which constitutes to 62% yield (2.4 mg).

$^1$H NMR (400 MHz, , *$d_6$*-DMSO) δ 9.34 (t, *J* = 5.5 Hz, 1H), 7.81 (dd, *J* = 7.5, 0.9 Hz, 1H), 7.70 (t, *J* = 7.6 Hz, 1H), 7.05 (dd, *J* = 7.7, 1.0 Hz, 1H), 6.89 (d, *J* = 2.4 Hz, 1H), 6.76 (s, 1H), 6.57 (dd, *J* = 8.9, 2.4 Hz, 1H), 6.53 (d, *J* = 8.8 Hz, 1H), 6.37 (s, 1H), 3.71 – 3.40 (m, 18H), 3.35 (t, *J* = 6.5 Hz, 2H), 3.28 – 3.19 (m, 2H), 2.93 (s, 6H), 2.79 (s, 3H), 2.77 – 2.66 (m, 2H), 1.79 (s, 3H), 1.74 – 1.63 (m, 5H), 1.47 (p, *J* = 6.7 Hz, 2H), 1.39 – 1.26 (m, 4H).

ESI-MS, positive mode: m/z = 784.4 $[M+Na]^+$.

HRMS (ESI) calcd for $C_{43}H_{56}ClN_3O_7Na$ $[M+Na]^+$ 784.3699, found 784.3687.

**4-630CP-Halo** (**70**):

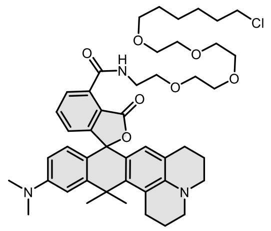

Compound was purified by preparative HPLC (solvent A: $H_2O$ + 10mM $NH_4COOH$ pH = 3.6, solvent B: MeOH; temperature 25 °C, gradient A:B - 3 min 70:30 isocratic, 4-20 min 70:30 to 0:100 gradient, 20-25 min 0:100 isocratic, 40 mL/min flow). Fractions containing the product were collected, evaporated and lyophilized from acetonitrile water mixture. The obtained solid was dissolved in 700 µL of $d_6$-DMSO. The samples from DMSO solution were diluted x100 in PBS +0.1% SDS and concentration was determined spectroscopically with Nanodrop. Determined stock concentration was 3.0 mM which constitutes to 42% yield (1.7 mg).

$^1$H NMR (400 MHz, *d*$_6$-DMSO) δ 9.38 (t, *J* = 5.5 Hz, 1H), 7.81 (dd, *J* = 7.5, 1.0 Hz, 1H), 7.68 (t, *J* = 7.6 Hz, 1H), 7.02 (dd, *J* = 7.7, 1.0 Hz, 1H), 6.80 (d, *J* = 2.5 Hz, 1H), 6.56 (dd, *J* = 9.0, 2.5 Hz, 1H), 6.47 (d, *J* = 8.8 Hz, 1H), 6.14 (s, 1H), 3.87 – 3.37 (m, 18H), 3.35 (t, *J* = 6.5 Hz, 2H), 3.24 – 3.07 (m, 4H), 2.92 (s, 8H), 2.49 – 2.35 (m, 2H), 2.03 – 1.85 (m, 5H), 1.82 (s, 3H), 1.79 – 1.63 (m, 4H), 1.47 (p, *J* = 6.8 Hz, 2H), 1.39 – 1.24 (m, 4H).

ESI-MS, positive mode: m/z = 824.4 $[M+Na]^+$.

HRMS (ESI) calcd for $C_{46}H_{60}ClN_3O_7Na$ $[M+Na]^+$ 824.4012, found 824.3995.

**4-640CP-Halo** (**71**):

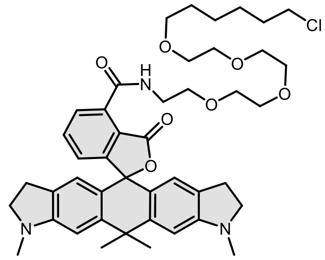

Compound was purified by preparative HPLC (solvent A: $H_2O$ + 10mM $NH_4COOH$ pH = 3.6, solvent B: MeOH; temperature 25 °C, gradient A:B - 3 min 60:40 isocratic, 4-20 min 60:40 to 0:100 gradient, 20-25 min 0:100 isocratic, 40 mL/min flow). Fractions containing the product were collected, evaporated and lyophilized from acetonitrile water mixture. The obtained solid was dissolved in 700 µL of $d_6$-DMSO. The samples from DMSO solution were diluted x100 in PBS +0.1% SDS and concentration was determined spectroscopically with Nanodrop. Determined stock concentration was 3.5 mM which constitutes to 47% yield (1.8 mg).

$^1$H NMR (400 MHz, *d*$_6$-DMSO) δ 9.36 (t, *J* = 5.5 Hz, 1H), 7.81 (dd, *J* = 7.5, 1.0 Hz, 1H), 7.69 (t, *J* = 7.6 Hz, 1H), 7.05 (dd, *J* = 7.7, 1.0 Hz, 1H), 6.74 (s, 2H), 6.34 (s, 2H), 3.73 – 3.38 (m, 18H), 3.35 (t, *J* = 6.5 Hz, 2H), 3.27 – 3.18 (m, 4H), 2.78 (s, 6H), 2.75 – 2.62 (m, 4H), 1.76 (s, 3H), 1.73 – 1.61 (m, 5H), 1.46 (p, *J* = 7.0 Hz, 2H), 1.38 – 1.28 (m, 4H).

ESI-MS, positive mode: m/z = 796.4 $[M+Na]^+$.

HRMS (ESI) calcd for $C_{44}H_{56}ClN_3O_7Na$ $[M+Na]^+$ 796.3699, found 796.3669.

**4-642CP-Halo** (**72**):

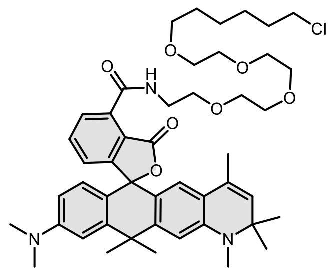

Compound was purified by preparative HPLC (solvent A: $H_2O$ + 10mM $NH_4COOH$ pH = 3.6, solvent B: MeOH; temperature 25 °C, gradient A:B - 3 min 30:70 isocratic, 4-20 min 30:70 to 0:100 gradient, 20-25 min 0:100 isocratic, 40 mL/min flow). Fractions containing the product were collected, evaporated and lyophilized from acetonitrile water mixture. The obtained solid was dissolved in 700 µL of $d_6$-DMSO. The samples from DMSO solution were diluted x100 in PBS +0.1% SDS and concentration was determined spectroscopically with Nanodrop. Determined stock concentration was 4.7 mM which constitutes to 66% yield (2.7 mg).

$^1$H NMR (400 MHz, *d*$_6$-DMSO) δ 9.38 (t, *J* = 5.5 Hz, 1H), 7.88 (dd, *J* = 7.6, 1.0 Hz, 1H), 7.74 (t, *J* = 7.7 Hz, 1H), 7.12 (dd, *J* = 7.7, 1.0 Hz, 1H), 6.91 (s, 1H), 6.67 (s, 1H), 6.56 (s, 2H), 6.16 (s, 1H), 5.33 (d, *J* = 1.6 Hz, 1H), 3.65 – 3.42 (m, 18H), 3.35 (t, *J* = 6.5 Hz, 2H), 2.94 (s, 6H), 2.84 (s, 3H), 1.81 (s, 3H), 1.73 – 1.64 (m, 5H), 1.55 (d, *J* = 1.4 Hz, 3H), 1.47 (p, *J* = 6.7 Hz, 2H), 1.38 – 1.25 (m, 10H).

ESI-MS, positive mode: m/z = 838.4 $[M+Na]^+$.

HRMS (ESI) calcd for $C_{47}H_{62}ClN_3O_7Na$ $[M+Na]^+$ 838.4169, found 838.4147.

**4-SiR-Halo** (**73**):

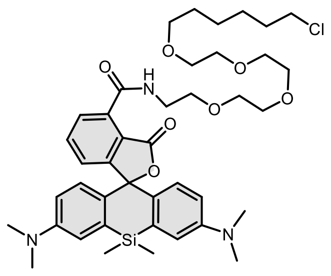

Compound was purified by preparative HPLC (solvent A: $H_2O$ + 10mM $NH_4COOH$ pH = 3.6, solvent B: MeOH; temperature 25 °C, gradient A:B - 3 min 30:70 isocratic, 4-20 min 30:70 to 0:100 gradient, 20-25 min 0:100 isocratic, 40 mL/min flow). Fractions containing the product were collected, evaporated and lyophilized from acetonitrile water mixture to give product in 70% yield (2.7 mg) as a light blue solid.

$^1$H NMR (400 MHz, *d*$_6$-DMSO) δ 9.22 (t, *J* = 5.5 Hz, 1H), 7.82 – 7.74 (m, 2H), 7.26 (dd, *J* = 7.2, 1.5 Hz, 1H), 7.00 (d, *J* = 2.7 Hz, 2H), 6.69 (d, *J* = 9.0 Hz, 2H), 6.64 (dd, *J* = 9.0, 2.8 Hz, 2H), 3.72 – 3.37 (m, 18H), 3.35 (t, *J* = 6.6 Hz, 2H), 2.92 (s, 12H), 1.68 (p, *J* = 6.7 Hz, 2H), 1.46 (p, *J* = 6.8 Hz, 2H), 1.35 – 1.21 (m, 4H), 0.62 (s, 3H), 0.52 (s, 3H).

ESI-MS, positive mode: m/z = 788.3 $[M+Na]^+$.

HRMS (ESI) calcd for $C_{41}H_{56}ClN_3O_7SiNa$ $[M+Na]^+$ 788.3468, found 788.3463.

**4-665SiR-Halo** (**74**):

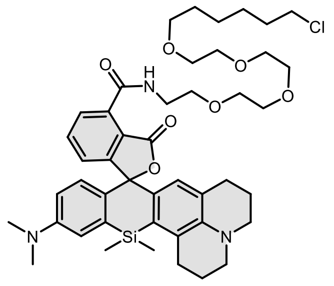

Compound was purified by preparative HPLC (solvent A: $H_2O$ + 10mM $NH_4COOH$ pH = 3.6, solvent B: MeOH; temperature 25 °C, gradient A:B - 3 min 30:70 isocratic, 4-20 min 30:70 to 0:100 gradient, 20-25 min 0:100 isocratic, 40 mL/min flow). Fractions containing the product were collected, evaporated and lyophilized from acetonitrile water mixture. The obtained solid was dissolved in 700 µL of $d_6$-DMSO. The samples from DMSO solution were diluted x100 in PBS +0.1% SDS and concentration was determined spectroscopically with Nanodrop. Determined stock concentration was 2.5 mM which constitutes to 35% yield (1.4 mg).

$^1$H NMR (400 MHz, *d₆*-DMSO) δ 9.26 (t, *J* = 5.5 Hz, 1H), 7.75 (d, *J* = 1.1 Hz, 1H), 7.71 (t, *J* = 7.6 Hz, 1H), 7.13 (dd, *J* = 7.6, 1.2 Hz, 1H), 6.93 (d, *J* = 2.5 Hz, 1H), 6.69 – 6.63 (m, 2H), 6.33 (s, 1H), 3.68 – 3.40 (m, 18H), 3.35 (t, *J* = 6.5 Hz, 2H), 3.16 (t, *J* = 5.8 Hz, 2H), 3.12 (t, *J* = 5.1 Hz, 2H), 2.96 – 2.85 (m, 8H), 2.47 – 2.40 (m, 2H), 1.98 – 1.93 (m, 2H), 1.79 – 1.74 (m, 2H), 1.68 (p, *J* = 6.7 Hz, 2H), 1.46 (p, *J* = 7.0 Hz, 2H), 1.36 – 1.27 (m, 4H), 0.66 (s, 3H), 0.55 (s, 3H).

ESI-MS, positive mode: m/z = 840.4 $[M+Na]^+$.

HRMS (ESI) calcd for $C_{45}H_{60}ClN_3O_7SiNa$ $[M+Na]^+$ 840.3781, found 840.3779.

**4-670SiR-Halo** (**75**):

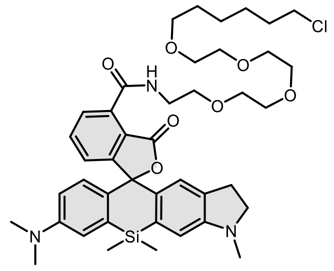

Compound was purified by preparative HPLC (solvent A: $H_2O$ + 0.2% HCOOH, solvent B: MeOH; temperature 25 °C, gradient A:B - 3 min 40:60 isocratic, 4-20 min 40:60 to 0:100 gradient, 20-25 min 0:100 isocratic, 40 mL/min flow). Fractions containing the product were collected, evaporated and lyophilized from acetonitrile water mixture to give product in 45% yield (1.75 mg) as a colorless solid.

$^1$H NMR (400 MHz, $d_6$-DMSO) δ 9.24 (t, *J* = 5.5 Hz, 1H), 7.79 (dd, *J* = 7.5, 1.3 Hz, 1H), 7.75 (t, *J* = 7.4 Hz, 1H), 7.21 (dd, *J* = 7.3, 1.4 Hz, 1H), 6.98 (d, *J* = 2.7 Hz, 1H), 6.80 (s, 1H), 6.69 (d, *J* = 9.0 Hz, 1H), 6.64 (dd, *J* = 9.1, 2.8 Hz, 1H), 6.57 (s, 1H), 3.62 – 3.42 (m, 18H), 3.35 (t, *J* = 6.5 Hz, 2H), 3.26 – 3.18 (m, 2H), 2.91 (s, 6H), 2.82 – 2.70 (m, 5H), 1.69 (p, *J* = 6.7 Hz, 2H), 1.46 (p, *J* = 6.8 Hz, 2H), 1.32 – 1.20 (m, 4H), 0.60 (s, 3H), 0.50 (s, 3H).

ESI-MS, positive mode: m/z = 800.3 $[M+Na]^+$.

HRMS (ESI) calcd for $C_{42}H_{56}ClN_3O_7SiNa$ $[M+Na]^+$ 800.3468, found 800.3464.

**4-685SiR-Halo** (**76**):

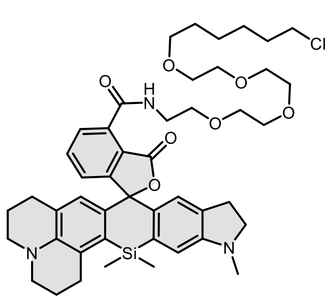

Compound was purified by preparative HPLC (solvent A: $H_2O$ + 0.2% HCOOH pH = 3.6, solvent B: MeOH; temperature 25 °C, gradient A:B - 3 min 40:60 isocratic, 4-20 min 40:60 to 0:100 gradient, 20-25 min 0:100 isocratic, 40 mL/min flow). Fractions containing the product were collected, evaporated and lyophilized from acetonitrile water mixture. The obtained solid was dissolved in 700 µL of $d_6$-DMSO. The samples from DMSO solution were diluted x100 in PBS +0.1% SDS and concentration was determined spectroscopically with Nanodrop. Determined stock concentration was 3.4 mM which constitutes to 48% yield (2.0 mg).

$^1$H NMR (400 MHz, $d_6$-DMSO) δ 9.29 (t, *J* = 5.5 Hz, 1H), 7.75 (dd, *J* = 7.4, 0.9 Hz, 1H), 7.69 (t, *J* = 7.6 Hz, 1H), 7.09 (dd, *J* = 7.6, 1.0 Hz, 1H), 6.74 (s, 1H), 6.54 (s, 1H), 6.31 (s, 1H), 3.63 – 3.42 (m, 18H), 3.34 (t, *J* = 6.8 Hz, 2H), 3.23 – 3.15 (m, 4H), 3.13 – 3.08 (m, 2H), 2.90 (t, *J* = 6.4 Hz, 2H), 2.76 (s, 5H), 2.48 – 2.38 (m, 2H), 1.94 (p, *J* = 6.3 Hz, 2H), 1.80 – 1.72 (m, 2H), 1.67 (p, *J* = 6.9 Hz, 2H), 1.46 (p, *J* = 6.8 Hz, 2H), 1.35 – 1.24 (m, 4H), 0.64 (s, 3H), 0.53 (s, 3H).

ESI-MS, positive mode: m/z = 852.4 $[M+Na]^+$.

HRMS (ESI) calcd for $C_{46}H_{60}ClN_3O_7SiNa$ $[M+Na]^+$ 852.3781, found 852.3770.

**4-690SiR-Halo** (**77**):

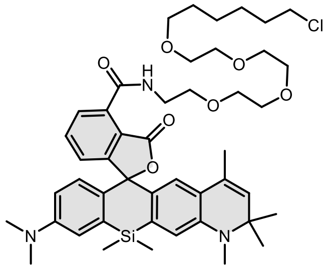

Compound was purified by preparative HPLC (solvent A: $H_2O$ + 0.2% HCOOH, solvent B: MeOH; temperature 25 °C, gradient A:B - 3 min 40:60 isocratic, 4-20 min 40:60 to 0:100 gradient, 20-25 min 0:100 isocratic, 40 mL/min flow). Fractions containing the product were collected, evaporated and lyophilized from acetonitrile water mixture to give product in 67% yield (2.8 mg) as a colorless solid.

$^1$H NMR (400 MHz, $d_6$-DMSO) δ 9.22 (t, *J* = 5.5 Hz, 1H), 7.86 – 7.79 (m, 2H), 7.37 (dd, *J* = 7.1, 1.6 Hz, 1H), 7.01 (d, *J* = 2.8 Hz, 1H), 6.76 (s, 1H), 6.68 (d, *J* = 8.9 Hz, 1H), 6.61 (dd, *J* = 9.0, 2.8 Hz, 1H), 6.40 (s, 1H), 5.35 (d, *J* = 1.5 Hz, 1H), 3.65 – 3.40 (m, 18H), 3.35 (t, *J* = 6.5 Hz, 2H), 2.92 (s, 6H), 2.81 (s, 3H), 1.68 (p, *J* = 7.0, 6.5 Hz, 2H), 1.57 (d, *J* = 1.4 Hz, 3H), 1.46 (p, *J* = 7.0 Hz, 2H), 1.35 – 1.26 (m, 4H), 1.25 (s, 3H), 1.24 (s, 3H), 0.63 (s, 3H), 0.51 (s, 3H).

ESI-MS, positive mode: m/z = 854.4 $[M+Na]^+$.

HRMS (ESI) calcd for $C_{46}H_{62}ClN_3O_7SiNa$ $[M+Na]^+$ 854.3938, found 854.3935.

**4-700SiR-Halo** (**78**):

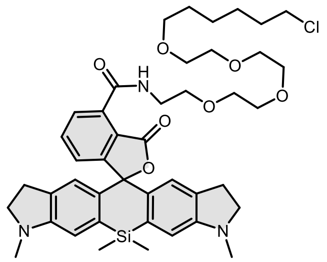

Compound was purified by preparative HPLC (solvent A: $H_2O$ + 0.2% HCOOH pH = 3.6, solvent B: MeOH; temperature 25 °C, gradient A:B - 3 min 40:60 isocratic, 4-20 min 40:60 to 0:100 gradient, 20-25 min 0:100 isocratic, 40 mL/min flow). Fractions containing the product were collected, evaporated and lyophilized from acetonitrile water mixture. The obtained solid was dissolved in 700 µL of $d_6$-DMSO. The samples from DMSO solution were diluted x100 in PBS +0.1% SDS and concentration was determined spectroscopically with Nanodrop. Determined stock concentration was 2.7 mM which constitutes to 38% yield (1.5 mg).

$^1$H NMR (400 MHz, *$d_6$*-DMSO) δ 9.26 (t, *J* = 5.5 Hz, 1H), 7.77 (dd, *J* = 7.5, 1.2 Hz, 1H), 7.72 (t, *J* = 7.5 Hz, 1H), 7.16 (dd, *J* = 7.5, 1.2 Hz, 1H), 6.78 (s, 2H), 6.57 (s, 2H), 3.69 – 3.38 (m, 18H), 3.35 (t, *J* = 6.5 Hz, 2H), 3.27 – 3.17 (m, 4H), 2.87 – 2.65 (m, 10H), 1.68 (p, *J* = 6.8 Hz, 2H), 1.46 (p, *J* = 6.8 Hz, 2H), 1.34 – 1.23 (m, 4H), 0.57 (s, 3H), 0.49 (s, 3H).

ESI-MS, positive mode: m/z = 812.3 $[M+Na]^+$.

HRMS (ESI) calcd for $C_{43}H_{56}ClN_3O_7SiNa$ $[M+Na]^+$ 812.3468, found 812.3462.

**4-720SiR-Halo** (**79**):

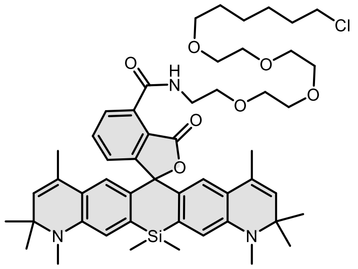

Compound was purified by preparative HPLC (solvent A: $H_2O$ + 0.2% HCOOH, solvent B: MeOH; temperature 25 °C, gradient A:B - 3 min 40:60 isocratic, 4-20 min 40:60 to 0:100 gradient, 20-25 min 0:100 isocratic, 40 mL/min flow). Fractions containing the product were collected, evaporated and lyophilized from acetonitrile water mixture to give product in 58% yield (2.6 mg) as a colorless solid.

$^1$H NMR (400 MHz, *$d_6$*-DMSO) δ 9.23 (t, *J* = 5.5 Hz, 1H), 7.88 (dd, *J* = 7.6, 1.7 Hz, 1H), 7.85 (t, *J* = 7.5, 7.1 Hz, 1H), 7.47 (dd, *J* = 7.1, 1.6 Hz, 1H), 6.76 (s, 2H), 6.39 (s, 2H), 5.34 (d, *J* = 1.5 Hz, 2H), 3.61 – 3.41 (m, 18H), 3.34 (t, *J* = 6.8 Hz, 2H), 2.81 (s, 6H), 1.68 (p, *J* = 6.8 Hz, 2H), 1.57 (d, *J* = 1.4 Hz, 6H), 1.46 (p, *J* = 6.8 Hz, 2H), 1.35 – 1.26 (m, 4H), 1.25 (s, 6H), 1.24 (s, 6H), 0.63 (s, 3H), 0.50 (s, 3H).

ESI-MS, positive mode: m/z = 920.4 $[M+Na]^+$.

HRMS (ESI) calcd for $C_{51}H_{68}ClN_3O_7SiNa$ $[M+Na]^+$ 898.4588, found 898.4570.

General procedure for the synthesis of compounds **80**-**86**:

Into a solution of corresponding rhodamine dye (5 µmol, 1 eq; **5**, **7, 8, 10, 13, 15, 23**) and DIPEA (5 µL) in DMSO (100 µL) a solution of HATU (6.5 µmol, 1.3 eq) in DMSO (100 µL) was added and a mixture was mixed for 1 min. Then a solution of (6-aminohexyl)triphenylphosphonium bromide hydrobromide (7.5 µmol ,1.5 eq, Sigma Aldrich) in DMSO (100 µL +5 µL DIPEA) was added at once. Reactions were usually over after 30 min and their course was monitored by LC/MS analysis. Once reaction was finished it was quenched with 20 µL of formic acid and diluted with water and acetonitrile to 2 mL volume and was further purified by the means of preparative HPLC (preparative column: Agilent 5 Prep-C18, 5 µm, 100 x 50 mm).

**4-525R-TPP (80):**

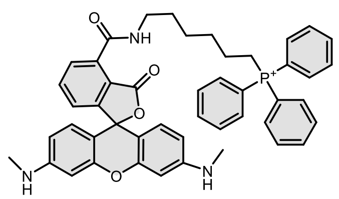

Compound was purified by preparative HPLC (solvent A: $H_2O$ + 10mM $NH_4COOH$ pH = 3.6, solvent B: MeCN; temperature 25 °C, gradient A:B - 3 min 70:30 isocratic, 4-20 min 70:30 to 0:100 gradient, 20-25 min 0:100 isocratic, 40 mL/min flow). Fractions containing the product were collected, evaporated and lyophilized from acetonitrile water mixture. The obtained solid was dissolved in 700 µL of $d_6$-DMSO. The samples from DMSO solution were diluted x100 in PBS +0.1% SDS and concentration was determined spectroscopically with Nanodrop. Determined stock concentration was 4.0 mM which constitutes to 56% yield (2.1 mg).

$^1$H NMR (400 MHz, $CD_3OD$) δ 7.87 – 7.74 (m, 11H), 7.69 – 7.61 (m, 6H), 7.37 (dd, *J* = 7.6, 1.2 Hz, 1H), 7.14 (d, *J* = 9.1 Hz, 2H), 6.76 (d, *J* = 2.2 Hz, 2H), 6.73 (dd, *J* = 9.2, 2.2 Hz, 2H), 3.45 – 3.36 (m, 4H), 3.01 (s, 6H), 1.67 – 1.51 (m, 8H).

ESI-MS, positive mode: m/z = 746.3 $[M]^+$.

HRMS (ESI) calcd for $C_{47}H_{45}N_3O_4P$ $[M]^+$ 746.3142, found 746.3141.

**4-580R-TPP (81):**

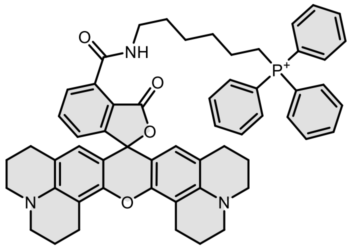

Compound was purified by preparative HPLC (solvent A: $H_2O$ + 10mM $NH_4COOH$ pH = 3.6, solvent B: MeCN; temperature 25°C, gradient A:B - 3 min 70:30 isocratic, 4-20 min 70:30 to 0:100 gradient, 20-25 min 0:100 isocratic, 40 mL/min flow). Fractions containing the product were collected, evaporated and lyophilized from acetonitrile water mixture. The obtained solid was dissolved in 700 µL of $d_6$-DMSO. The samples from DMSO solution were diluted x100 in PBS +0.1% SDS and concentration was determined spectroscopically with Nanodrop. Determined stock concentration was 1.9 mM which constitutes to 27% yield (1.2 mg).

$^1$H NMR (400 MHz, $CD_3OD$) δ 7.86 – 7.75 (m, 10H), 7.68 – 7.63 (m, 5H), 7.55 (t, *J* = 7.7 Hz, 1H), 7.30 (dd, *J* = 7.6, 1.3 Hz, 1H), 6.82 (s, 2H), 3.54 (t, *J* = 5.8 Hz, 4H), 3.50 – 3.44 (m, 4H), 3.43 – 3.33 (m, 4H), 3.06 (t, *J* = 6.2 Hz, 4H), 2.63 (q, *J* = 6.2 Hz, 4H), 2.09 (p, *J* = 6.3 Hz, 4H), 1.94 – 1.83 (m, 4H), 1.67 – 1.51 (m, 8H).

ESI-MS, positive mode: m/z = 878.4 $[M]^+$.

HRMS (ESI) calcd for $C_{57}H_{57}N_3O_4P$ $[M]^+$ 878.4081, found 878.4091.

**4-625-TPP (82):**

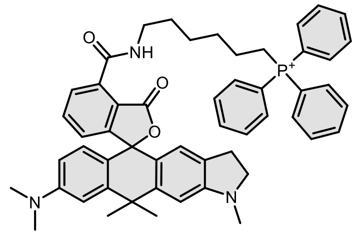

Compound was purified by preparative HPLC (solvent A: $H_2O$ + 10mM $NH_4COOH$ pH = 3.6, solvent B: MeCN; temperature 25°C, gradient A:B - 3 min 70:30 isocratic, 4-20 min 70:30 to 0:100 gradient, 20-25 min 0:100 isocratic, 40 mL/min flow). Fractions containing the product were collected, evaporated and lyophilized from acetonitrile water mixture. The obtained solid was dissolved in 700 µL of $d_6$-DMSO. The samples from DMSO solution were diluted x100 in PBS +0.1% SDS and concentration was determined spectroscopically with Nanodrop. Determined stock concentration was 2.4 mM which constitutes to 34% yield (1.4 mg).

$^1$H NMR (400 MHz, $CD_3OD$) δ 7.96 (dd, *J* = 7.6, 1.0 Hz, 1H), 7.80 – 7.71 (m, 10H), 7.65 – 7.58 (m, 6H), 7.15 (dd, *J* = 7.8, 1.0 Hz, 1H), 7.00 (d, *J* = 2.4 Hz, 1H), 6.81 (s, 1H), 6.59 (d, *J* = 8.8 Hz, 1H), 6.55 (dd, *J* = 8.9, 2.4 Hz, 1H), 6.41 (s, 1H), 3.52 – 3.41 (m, 4H), 3.36 – 3.32 (m, 2H), 3.01 (s, 6H), 2.88 (s, 3H), 2.72 (t, *J* = 8.2 Hz, 2H), 1.86 (s, 3H), 1.75 (s, 3H), 1.73 – 1.61 (m, 8H).

ESI-MS, positive mode: m/z = 812.4 $[M]^+$.

HRMS (ESI) calcd for $C_{53}H_{55}N_3O_3P$ $[M]^+$ 812.3976, found 812.3964.

**4-640-TPP (83):**

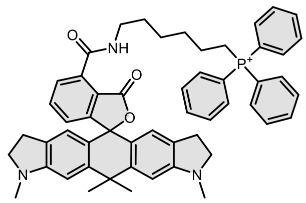

Compound was purified by preparative HPLC (solvent A: $H_2O$ + 10 mM $NH_4COOH$ pH = 3.6, solvent B: MeCN; temperature 25°C, gradient A:B - 3 min 40:60 isocratic, 4-20 min 40:60 to 0:100 gradient, 20-25 min 0:100 isocratic, 40 mL/min flow). Fractions containing the product were collected, evaporated and lyophilized from acetonitrile water mixture. The obtained solid was dissolved in 700 µL of $d_6$-DMSO. The samples from DMSO solution were diluted x100 in PBS +0.1% SDS and concentration was determined spectroscopically with Nanodrop. Determined stock concentration was 2.9 mM which constitutes to 41% yield (1.7 mg).

$^{1}H$ NMR (400 MHz, $CD_3OD$) δ 7.90 (dd, *J* = 7.6, 1.0 Hz, 1H), 7.86 – 7.73 (m, 10H), 7.68 (t, *J* = 7.7 Hz, 1H), 7.65 – 7.60 (m, 5H), 7.15 (dd, *J* = 7.7, 1.1 Hz, 1H), 6.82 (s, 2H), 6.48 (s, 2H), 3.51 – 3.43 (m, 4H), 3.40 (td, *J* = 8.2, 2.3 Hz, 4H), 2.93 (s, 6H), 2.77 – 2.72 (m, 4H), 1.82 (s, 3H), 1.73 (s, 3H), 1.65 (s, 8H).

ESI-MS, positive mode: m/z = 824.4 $[M]^+$.

HRMS (ESI) calcd for $C_{54}H_{55}N_3O_3P$ $[M]^+$ 824.3976, found 824.3950.

**4-665SiR-TPP (84):**

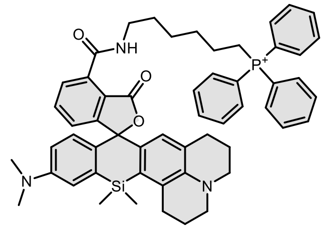

Compound was purified by preparative HPLC (solvent A: $H_2O$ + 10 mM $NH_4COOH$ pH = 3.6, solvent B: MeCN; temperature 25°C, gradient A:B - 3 min 40:60 isocratic, 4-20 min 40:60 to 0:100 gradient, 20-25 min 0:100 isocratic, 40 mL/min flow). Fractions containing the product were collected, evaporated and lyophilized from acetonitrile water mixture. The obtained solid was dissolved in 700 µL of $d_6$-DMSO. The samples from DMSO solution were diluted x100 in PBS +0.1% SDS and concentration was determined spectroscopically with Nanodrop. Determined stock concentration was 2.2 mM which constitutes to 31% yield (1.3 mg).

$^{1}H$ NMR (400 MHz $CD_3OD$) δ 7.90 (dd, *J* = 7.6, 1.0 Hz, 1H), 7.82 – 7.73 (m, 11H), 7.65 – 7.60 (m, 5H), 7.27 (dd, *J* = 7.8, 1.0 Hz, 1H), 7.02 (d, *J* = 2.8 Hz, 1H), 6.68 (d, *J* = 8.9 Hz, 1H), 6.62 – 6.59 (m, 1H), 6.32 (s, 1H), 3.50 – 3.41 (m, 4H), 3.23 (t, *J* = 5.9 Hz, 2H), 3.17 (t, *J* = 5.8 Hz, 2H), 3.01 (dt, *J* = 8.2, 4.0 Hz, 2H), 2.97 (s, 6H), 2.42 (t, *J* = 6.4 Hz, 2H), 2.07 – 2.02 (m, 2H), 1.84 – 1.79 (m, 2H), 1.76 – 1.65 (m, 8H), 0.69 (s, 3H), 0.54 (s, 3H).

ESI-MS, positive mode: m/z = 868.3 $[M]^+$.

HRMS (ESI) calcd for $C_{55}H_{59}N_3O_3PSi$ $[M]^+$ 868.4058, found 868.4038.

**4-685SiR-TPP (85):**

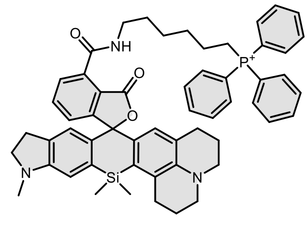

Compound was purified by preparative HPLC (solvent A: $H_2O$ + 10 mM $NH_4COOH$ pH = 3.6, solvent B: MeCN; temperature 25°C, gradient A:B - 3 min 40:60 isocratic, 4-20 min 40:60 to 0:100 gradient, 20-25 min 0:100 isocratic, 40 mL/min flow). Fractions containing the product were collected, evaporated and lyophilized from acetonitrile water mixture. The obtained solid was dissolved in 700 µL of $d_6$-DMSO. The samples from DMSO solution were diluted x100 in PBS +0.1% SDS and concentration was determined spectroscopically with Nanodrop. Determined stock concentration was 2.5 mM which constitutes to 35% yield (1.5 mg).

$^1$H NMR (400 MHz, $CD_3OD$) δ 7.85 (dd, *J* = 7.6, 0.9 Hz, 1H), 7.83 – 7.66 (m, 11H), 7.66 – 7.61 (m, 5H), 7.19 (dd, *J* = 7.8, 0.9 Hz, 1H), 6.77 (s, 1H), 6.55 (s, 1H), 6.33 (s, 1H), 3.50 – 3.43 (m, 4H), 3.28 – 3.21 (m, 4H), 3.18 – 3.14 (m, 2H), 3.00 (t, *J* = 6.3 Hz, 2H), 2.82 (s, 6H), 2.71 (t, *J* = 8.2 Hz, 2H), 2.40 (t, *J* = 6.3 Hz, 2H), 2.05 – 2.02 (m, 2H), 1.84 – 1.66 (m, 10H), 0.67 (s, 3H), 0.54 (s, 3H).

ESI-MS, positive mode: m/z = 880.4 $[M]^+$.

HRMS (ESI) calcd for $C_{56}H_{59}N_3O_3PSi$ $[M]^+$ 880.4058, found 880.4039.

**4-700SiR-TPP (86):**

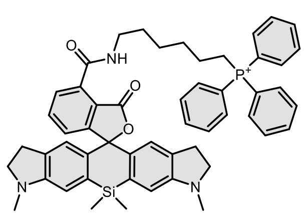

Compound was purified by preparative HPLC (solvent A: $H_2O$ + 10 mM $NH_4COOH$ pH = 3.6, solvent B: MeCN; temperature 25°C, gradient A:B - 3 min 40:60 isocratic, 4-20 min 40:60 to 0:100 gradient, 20-25 min 0:100 isocratic, 40 mL/min flow). Fractions containing the product were collected, evaporated and lyophilized from acetonitrile water mixture. The obtained solid was dissolved in 700 µL of $d_6$-DMSO. The samples from DMSO solution were diluted x100 in PBS +0.1% SDS and concentration was determined spectroscopically with Nanodrop. Determined stock concentration was 2.3 mM which constitutes to 32% yield (1.3 mg).

$^1$H NMR (400 MHz, $CD_3OD$) δ 7.88 (dd, *J* = 7.6, 0.9 Hz, 1H), 7.84 – 7.71 (m, 11H), 7.65 – 7.61 (m, 5H), 7.25 (dd, *J* = 7.8, 1.0 Hz, 1H), 6.79 (s, 2H), 6.57 (d, *J* = 0.5 Hz, 2H), 3.51 – 3.42 (m, 4H), 3.24 (h, *J* = 8.4 Hz, 4H), 2.82 (s, 6H), 2.72 (td, *J* = 8.2, 1.1 Hz, 4H), 1.67 (s, 8H), 0.60 (s, 3H), 0.47 (s, 3H).

ESI-MS, positive mode: m/z = 840.2 $[M]^+$.

HRMS (ESI) calcd for $C_{53}H_{55}N_3O_3PSi$ $[M]^+$ 840.3745, found 840.3745.

<u>General procedure for the synthesis of compounds **87**-**92**:</u>

Into a solution of corresponding rhodamine dye (5 µmol, 1 eq; **10**, **11, 15, 22, 23, 26**) and DIPEA (5 µL) in DMSO (100 µL) a solution of HATU (6.5 µmol, 1.3 eq) in DMSO (100 µL) was added and the mixture was mixed for 1 min. Then a solution of **Hoechst-$C_4$-$NH_2$**[23] (7.5 µmol ,1.5 eq) in DMSO (100 µL +5 µL DIPEA) was added at once. Reactions were usually over after 30 min and their course was monitored by LC/MS analysis. Once reaction was finished it was quenched with 20 µL of formic acid and diluted with water and acetonitrile to 2 mL volume and was further purified by the means of preparative HPLC (preparative column: Agilent 5 Prep-C18, 5 µm, 100 x 50 mm).

**4-CFL-Hoechst** (**87**):

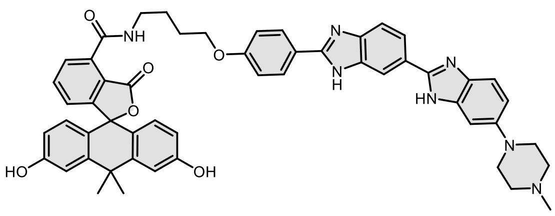

Compound was purified by preparative HPLC (solvent A: $H_2O$ + 0.2% HCOOH, solvent B: MeCN; temperature 25 °C, gradient A:B - 3 min 80:20 isocratic, 4-20 min 80:20 to 0:100 gradient, 20-25 min 0:100 isocratic, 40 mL/min flow). Fractions containing the product were collected, evaporated and lyophilized from acetonitrile water mixture to give product in 48% yield (2.1 mg) as a colorless solid.

$^1$H NMR (400 MHz, *d*$_6$-DMSO) δ 12.93 (br s, 1H), 12.55 (br s, 1H), 9.70 (br s, 2H), 9.06 (t, *J* = 5.7 Hz, 1H), 8.43 – 8.17 (br m, 2H), 8.17 – 8.06 (m, 3H), 7.99 (br s, 1H), 7.77 (dd, *J* = 7.5, 1.1 Hz, 1H), 7.71 (t, *J* = 7.5 Hz, 1H), 7.47 (br s, 1H), 7.16 (d, *J* = 9.0 Hz, 2H), 7.08 (dd, *J* = 7.6, 1.2 Hz, 2H), 6.94 (d, *J* = 8.6 Hz, 2H), 6.69 – 6.46 (m, 4H), 4.16 (t, *J* = 6.4 Hz, 2H), 3.46 (q, *J* = 6.7 Hz, 2H), 3.17-3.09 (m, 4H), 2.61-2.51 (m, 4H), 2.28 (s, 3H), 1.94 (p, *J* = 7.1 Hz, 2H), 1.80 (p, *J* = 7.2 Hz, 2H), 1.73 (s, 3H), 1.64 (s, 3H).

ESI-MS, positive mode: m/z = 880.4 $[M+H]^+$.

HRMS (ESI) calcd for $C_{53}H_{50}N_7O_6$ $[M+H]^+$ 880.3817, found 880.3807.

**4-505R-Hoechst** (**88**):

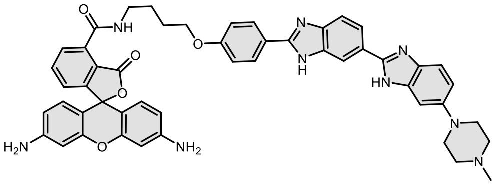

Compound was purified by preparative HPLC (solvent A: $H_2O$ + 10 mM $NH_4COOH$ pH = 3.6, solvent B: MeCN; temperature 25°C, gradient A:B - 3 min 80:20 isocratic, 4-20 min 80:20 to 0:100 gradient, 20-25 min 0:100 isocratic, 40 mL/min flow). Fractions containing the product were collected, evaporated and lyophilized from acetonitrile water mixture. The obtained solid was dissolved in 700 µL of d$_6$-DMSO. The samples from DMSO solution were diluted x100 in PBS +0.1% SDS and concentration was determined spectroscopically with Nanodrop. Determined stock concentration was 3.8 mM which constitutes to 53% yield (2.2 mg).

$^1$H NMR (400 MHz, *d*$_6$-DMSO) δ 9.10 (t, *J* = 5.6 Hz, 1H), 8.15 (d, *J* = 8.8 Hz, 2H), 7.99 (dd, *J* = 8.5, 1.6 Hz, 1H), 7.82 (d, *J* = 1.2 Hz, 1H), 7.77 (t, *J* = 7.5 Hz, 1H), 7.66 (d, *J* = 8.4 Hz, 1H), 7.44 (d, *J* = 8.7 Hz, 1H), 7.25 (dd, *J* = 7.5, 1.2 Hz, 1H), 7.15 (d, *J* = 8.9 Hz, 2H), 7.00 (s, 2H), 6.95 – 6.89 (m, 1H), 6.45 (d, *J* = 8.6 Hz, 2H), 6.38 (d, *J* = 2.2 Hz, 2H), 6.30 (dd, *J* = 8.5, 2.2 Hz, 2H), 5.59 (s, 4H), 4.15 (t, *J* = 6.3 Hz, 2H), 3.44 (q, *J* = 6.2 Hz, 2H), 3.13 (t, *J* = 4.9 Hz, 4H), 2.57 – 2.51 (m, 4H), 2.25 (s, 3H), 1.92 (p, *J* = 7.5 Hz, 2H), 1.78 (p, *J* = 7.3 Hz, 2H).

ESI-MS, positive mode: m/z = 852.4 $[M+H]^+$.

HRMS (ESI) calcd for $C_{50}H_{46}N_9O_5$ $[M+H]^+$ 852.3616, found 852.3612.

**4-525R-Hoechst** (**89**):

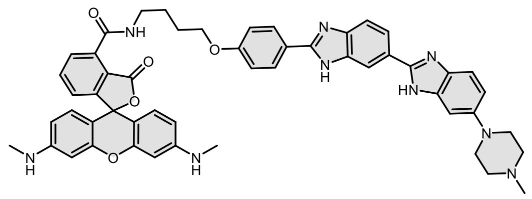

Compound was purified by preparative HPLC (solvent A: $H_2O$ + 10 mM $NH_4COOH$ pH = 3.6, solvent B: MeCN; temperature 25°C, gradient A:B - 3 min 70:30 isocratic, 4-20 min 70:30 to 0:100 gradient, 20-25 min 0:100 isocratic, 40 mL/min flow). Fractions containing the product were collected, evaporated and lyophilized from acetonitrile water mixture. The obtained solid was dissolved in 700 µL of $d_6$-DMSO. The samples from DMSO solution were diluted x100 in PBS +0.1% SDS and concentration was determined spectroscopically with Nanodrop. Determined stock concentration was 3.1 mM which constitutes to 44% yield (1.9 mg).

$^1$H NMR (400 MHz, $d_6$-DMSO + $CF_3COOD$) δ 9.11 (t, *J* = 5.6 Hz, 1H), 8.27 (s, 1H), 8.15 (d, *J* = 8.8 Hz, 2H), 7.99 (d, *J* = 8.5 Hz, 1H), 7.83 (dd, *J* = 7.5, 1.1 Hz, 1H), 7.77 (t, *J* = 7.6 Hz, 1H), 7.66 (d, *J* = 8.4 Hz, 1H), 7.44 (d, *J* = 8.0 Hz, 1H), 7.24 (dd, *J* = 7.6, 1.1 Hz, 1H), 7.15 (d, *J* = 9.0 Hz, 2H), 6.99 (s, 1H), 6.93 (dd, *J* = 8.8, 2.2 Hz, 1H), 6.50 (d, *J* = 8.5 Hz, 2H), 6.36 – 6.29 (m, 4H), 6.18 (q, *J* = 4.9 Hz, 2H), 4.15 (t, *J* = 6.3 Hz, 2H), 3.45 (p, *J* = 6.8 Hz, 2H), 3.13 (t, *J* = 5.0 Hz, 4H), 2.69 (d, *J* = 4.7 Hz, 6H), 2.54 – 2.51 (m, 4H), 2.25 (s, 3H), 1.96 – 1.89 (m, 2H), 1.83 – 1.76 (m, 2H).

ESI-MS, positive mode: m/z = 880.4 $[M+H]^+$.

HRMS (ESI) calcd for $C_{52}H_{50}N_9O_5$ $[M+H]^+$ 880.3929, found 880.3935.

**4-625CP-Hoechst** (**90**):

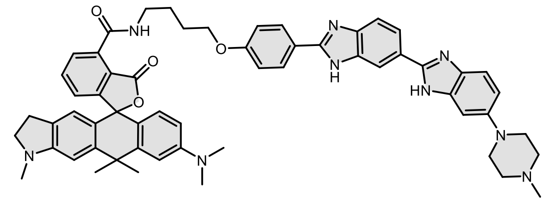

Compound was purified by preparative HPLC (solvent A: $H_2O$ + 10 mM $NH_4COOH$ pH = 3.6, solvent B: MeCN; temperature 25°C, gradient A:B - 3 min 70:30 isocratic, 4-20 min 70:30 to 0:100 gradient, 20-25 min 0:100 isocratic, 40 mL/min flow). Fractions containing the product were collected, evaporated and lyophilized from acetonitrile water mixture. The obtained solid was dissolved in 700 µL of $d_6$-DMSO. The samples from DMSO solution were diluted x100 in PBS +0.1% SDS and concentration was determined spectroscopically with Nanodrop. Determined stock concentration was 3.5 mM which constitutes to 49% yield (2.3 mg).

$^1$H NMR (400 MHz, $d_6$-DMSO + $CF_3COOD$) δ 9.17 (t, *J* = 5.6 Hz, 1H), 8.28 (s, 1H), 8.15 (d, *J* = 8.8 Hz, 2H), 7.99 (d, *J* = 8.5 Hz, 1H), 7.77 (dd, *J* = 7.5, 1.0 Hz, 1H), 7.72 – 7.63 (m, 2H), 7.44 (s, 1H), 7.16 (d, *J* = 9.0 Hz, 2H), 7.04 (dd, *J* = 7.7, 1.0 Hz, 1H), 6.99 – 6.87 (m, 3H), 6.76 (s, 1H), 6.59 – 6.53 (m, 2H), 6.39 (s, 1H), 4.16 (t, *J* = 6.4 Hz, 2H), 3.46 (q, *J* = 6.6 Hz, 2H), 3.25 – 3.22 (m, 2H), 3.13 (t, *J* = 4.7 Hz, 4H), 2.93 (s, 6H), 2.79

(s, 3H), 2.77 – 2.68 (m, 2H), 2.56 – 2.51 (m, 4H), 2.25 (s, 3H), 1.98 – 1.92 (m, 2H), 1.85 – 1.77 (m, 5H), 1.71 (s, 3H).

ESI-MS, positive mode: m/z = 946.5 $[M+H]^+$.

HRMS (ESI) calcd for $C_{58}H_{60}N_9O_4$ $[M+H]^+$ 946.4763, found 946.4743.

**4-630CP-Hoechst** (**91**):

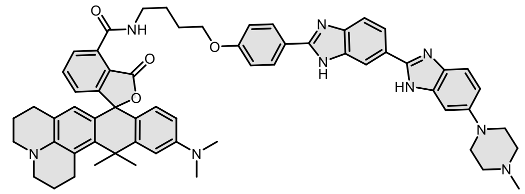

Compound was purified by preparative HPLC (solvent A: $H_2O$ + 10 mM $NH_4COOH$ pH = 3.6, solvent B: MeCN; temperature 25°C, gradient A:B - 3 min 70:30 isocratic, 4-20 min 70:30 to 0:100 gradient, 20-25 min 0:100 isocratic, 40 mL/min flow). Fractions containing the product were collected, evaporated and lyophilized from acetonitrile water mixture. The obtained solid was dissolved in 700 µL of $d_6$-DMSO. The samples from DMSO solution were diluted x100 in PBS +0.1% SDS and concentration was determined spectroscopically with Nanodrop. Determined stock concentration was 2.3 mM which constitutes to 32% yield (1.6 mg).

$^1$H NMR (400 MHz, $d_6$-DMSO) δ 12.93 (s, 1H), 12.57 (s, 1H), 9.20 (t, *J* = 5.6 Hz, 1H), 8.34 (s, 1H), 8.22 (s, 1H), 8.15 (d, *J* = 8.6 Hz, 3H), 8.03 – 7.96 (m, 1H), 7.76 (dd, *J* = 7.5, 1.0 Hz, 1H), 7.71 (d, *J* = 8.6 Hz, 1H), 7.67 (t, *J* = 7.6 Hz, 1H), 7.60 (d, *J* = 8.2 Hz, 1H), 7.16 (d, *J* = 8.8 Hz, 2H), 7.00 (dd, *J* = 7.7, 1.0 Hz, 1H), 6.94 (d, *J* = 7.2 Hz, 1H), 6.79 (d, *J* = 2.6 Hz, 1H), 6.57 (dd, *J* = 9.0, 2.5 Hz, 1H), 6.50 (d, *J* = 8.9 Hz, 1H), 6.16 (s, 1H), 4.17 (t, *J* = 6.4 Hz, 2H), 3.46 (q, *J* = 6.7 Hz, 2H), 3.19 – 3.12 (m, 8H), 2.97 – 2.89 (m, 8H), 2.62 – 2.56 (m, 4H), 2.47 – 2.36 (m, 2H), 2.31 (s, 3H), 1.98 – 1.90 (m, 4H), 1.89 (s, 3H), 1.83 (s, 3H), 1.82 – 1.71 (m, 4H).

ESI-MS, positive mode: m/z = 986.5 $[M+H]^+$.

HRMS (ESI) calcd for $C_{61}H_{64}N_9O_4Si$ $[M+H]^+$ 986.5076, found 986.5056.

**4-685SiR-Hoechst** (**92**):

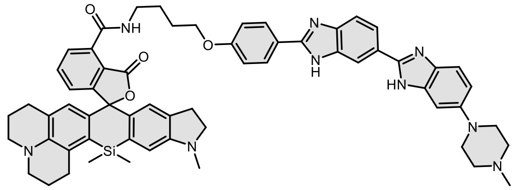

Compound was purified by preparative HPLC (solvent A: $H_2O$ + 10 mM $NH_4COOH$ pH = 3.6, solvent B: MeCN; temperature 25°C, gradient A:B - 3 min 70:30 isocratic, 4-20 min 70:30 to 0:100 gradient, 20-25 min 0:100 isocratic, 40 mL/min flow). Fractions containing the product were collected, evaporated and lyophilized from acetonitrile water mixture. The obtained solid was dissolved in 700 µL of $d_6$-DMSO. The samples from DMSO solution were diluted x100 in PBS +0.1% SDS and concentration was determined spectroscopically with Nanodrop. Determined stock concentration was 4.1 mM which constitutes to 58% yield (2.9 mg).

$^{1}H$ NMR (400 MHz, $d_6$-DMSO) δ 12.95 (s, 1H), 12.56 (s, 1H), 9.11 (t, $J$ = 5.7 Hz, 1H), 8.29 (s, 2H), 8.15 (d, $J$ = 8.8 Hz, 2H), 7.99 (d, $J$ = 8.4 Hz, 1H), 7.74 – 7.64 (m, 3H), 7.45 (s, 2H), 7.16 (d, $J$ = 9.0 Hz, 2H), 7.09 (dd, $J$ = 6.9, 1.8 Hz, 1H), 6.93 (d, $J$ = 8.4 Hz, 1H), 6.75 (s, 1H), 6.57 (s, 1H), 6.34 (s, 1H), 4.16 (t, $J$ = 6.3 Hz, 2H), 3.44 (q, $J$ = 6.2 Hz, 2H), 3.22 – 3.10 (m, 10H), 2.90 (t, $J$ = 6.3 Hz, 2H), 2.81 – 2.70 (m, 5H), 2.56 – 2.51 (m, 4H), 2.48 – 2.39 (m, 2H), 2.25 (s, 3H), 1.98 – 1.90 (m, 4H), 1.84 – 1.72 (m, 4H), 0.64 (s, 3H), 0.54 (s, 3H).

ESI-MS, positive mode: m/z = 1014.5 $[M+H]^+$.

HRMS (ESI) calcd for $C_{61}H_{64}N_9O_4Si$ $[M+H]^+$ 1014.4845, found 1014.4810.

**PepA-C6-NHBoc:**

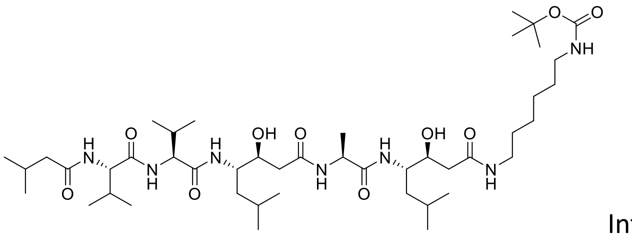

Into a solution of Pepstatin A (50 mg, 73 µmol, 1 eq;) and DIPEA (50 µL) in DMSO (1000 µL) a solution of TSTU (27 mg ,88 µmol, 1.2 eq) in DMSO (100 µL) was added and the mixture was mixed for 1-2 hours. Then a solution of N-(tert-Butoxycarbonyl)-1,6-diaminohexane (24 mg,110 µmol ,1.5 eq) in DMSO (100 µL +5 µL DIPEA) was added at once. Reaction was stirred for 1-2 hours the course of reaction was monitored by LC/MS analysis. Once reaction was finished it was quenched with 50 µL of formic acid and was further purified by the means of preparative HPLC (preparative column: Agilent 5 Prep-C18, 5 µm, 100 x 50 mm), collected fractions were collected and evaporated. The residue was dissolved in MeCN/$H_2O$ mixture and lyophilized to give 35 mg (54% yield) of colorless solid.

$^{1}H$ NMR (400 MHz, $d_6$-DMSO) δ 7.92 (d, $J$ = 7.3 Hz, 1H), 7.82 (d, $J$ = 8.8 Hz, 1H), 7.78 (d, $J$ = 8.8 Hz, 1H), 7.67 (t, $J$ = 5.6 Hz, 1H), 7.46 (d, $J$ = 8.8 Hz, 1H), 7.32 (d, $J$ = 9.1 Hz, 1H), 6.75 (t, $J$ = 5.0 Hz, 1H), 4.84 (t, $J$ = 5.7 Hz, 2H), 4.27 – 4.10 (m, 3H), 3.90 – 3.71 (m, 4H), 3.05 – 2.96 (m, 2H), 2.91 – 2.85 (m, 2H), 2.20 – 1.87 (m, 9H), 1.60 – 1.47 (m, 3H), 1.46 – 0.96 (m, 30H), 0.95 – 0.66 (m, 24H).

$^{13}C$ NMR (101 MHz, $d_6$-DMSO) δ 172.2, 171.6, 171.1, 170.8, 170.7, 170.6, 155.6, 77.3, 69.2, 69.0, 58.0, 57.8, 50.7, 50.4, 48.4, 44.4, 38.6, 38.4, 30.3, 30.1, 29.4, 29.1, 28.3, 26.1, 26.0, 25.7, 25.2, 24.2, 23.5, 23.3, 22.3, 22.3, 21.9, 21.6, 19.3, 19.3, 18.4, 18.3, 18.2.

ESI-MS, positive mode: m/z = 906.6 $[M+Na]^+$.

HRMS (ESI) calcd for $C_{45}H_{85}N_7O_{10}Na$ $[M+Na]^+$ 906.6250, found 906.6252.

General procedure for the synthesis of compounds **93**-**99**:

**PepA-C6-NH-Boc** (35 mg, 39 µmol) was dissolved in formic acid (1 mL) and was stirred for 3 hours at room temperature. The excess of formic acid was evaporated on rotary evaporator MeCN was added couple of times and evaporated to ensure complete removal of formic acid. Finally residue was dissolved in MeCN/$H_2O$ mixture and lyophilized to afford **PePA-C6-NH₂** quantitatively as a colorless solid. The obtained compound was further without any additional purifications.

Into a solution of corresponding rhodamine dye (5 µmol, 1 eq; **2**, **5**, **8**, **11**, **12**, **15**, **23**) and DIPEA (5 µL) in DMSO (100 µL) a solution of HATU (6.5 µmol, 1.3 eq) in DMSO (100 µL) was added and the mixture was mixed for 1 min. Then a solution of **PepA-C6-NH$_2$** (7.5 µmol ,1.5 eq) in DMSO (100 µL +5 µL DIPEA) was added at once. Reactions were usually over after 30 min and their course was monitored by LC/MS analysis. Once reaction was finished it was quenched with 20 µL of formic acid and diluted with water and acetonitrile to 2 mL volume and was further purified by the means of preparative HPLC (preparative column: Agilent 5 Prep-C18, 5 µm, 100 x 50 mm).

**4-525R-PepA (93):**

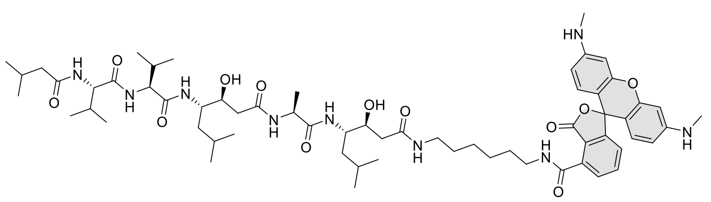

Compound was purified by preparative HPLC (solvent A: $H_2O$ + 10 mM $NH_4COOH$ pH = 3.6, solvent B: MeCN; temperature 25°C, gradient A:B - 3 min 70:30 isocratic, 4-20 min 70:30 to 0:100 gradient, 20-25 min 0:100 isocratic, 40 mL/min flow). Fractions containing the product were collected, evaporated and lyophilized from acetonitrile water mixture. The obtained solid was dissolved in 700 µL of $d_6$-DMSO. The samples from DMSO solution were diluted x100 in PBS +0.1% SDS and concentration was determined spectroscopically with Nanodrop. Determined stock concentration was 2.2 mM which constitutes to 31% yield (1.8 mg).

$^1$H NMR (400 MHz, $d_6$-DMSO) δ 9.05 (t, *J* = 5.6 Hz, 1H), 7.96 (s, 1H), 7.90 (d, *J* = 7.4 Hz, 1H), 7.82 – 7.74 (m, 3H), 7.70 – 7.65 (m, 1H), 7.45 (d, *J* = 8.9 Hz, 1H), 7.30 (s, 1H), 7.22 – 7.18 (m, 1H), 6.53 – 6.43 (m, 2H), 6.44 – 6.21 (m, 4H), 6.16 (d, *J* = 5.1 Hz, 2H), 4.85 – 4.79 (m, 2H), 4.24 – 4.10 (m, 3H), 3.80 (s, 4H), 3.06 – 2.93 (m, 4H), 2.68 (d, *J* = 4.8 Hz, 6H), 2.11 – 1.89 (m, 9H), 1.68 – 1.07 (m, 23H), 0.84 – 0.76 (m, 24H).

ESI-MS, positive mode: m/z = 1168.7 $[M+H]^+$.

HRMS (ESI) calcd for $C_{63}H_{94}N_9O_{12}$ $[M+H]^+$ 1168.7016, found 1168.6960.

**4-580R-PepA** (**94**):

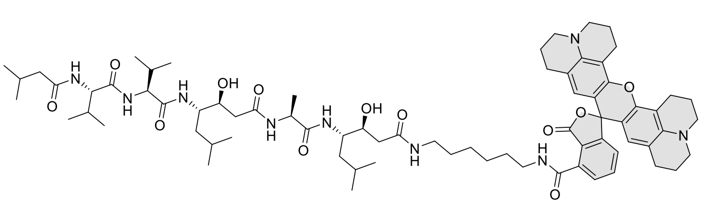

Compound was purified by preparative HPLC (solvent A: $H_2O$ + 10 mM $NH_4COOH$ pH = 3.6, solvent B: MeCN; temperature 25°C, gradient A:B - 3 min 70:30 isocratic, 4-20 min 70:30 to 0:100 gradient, 20-25 min 0:100 isocratic, 40 mL/min flow). Fractions containing the product were collected, evaporated and lyophilized from acetonitrile water mixture. The obtained solid was dissolved in 700 µL of $d_6$-DMSO. The samples from DMSO solution were diluted x100 in PBS +0.1% SDS and concentration was determined spectroscopically with Nanodrop. Determined stock concentration was 2.7 mM which constitutes to 38% yield (2.5 mg).

$^1$H NMR (400 MHz, $d_6$-DMSO) δ 9.19 (t, *J* = 5.6 Hz, 1H), 8.33 (s, 1H), 7.92 (d, *J* = 7.3 Hz, 1H), 7.87 – 7.71 (m, 4H), 7.47 (d, *J* = 8.8 Hz, 1H), 7.33 (d, *J* = 9.1 Hz, 1H), 7.25 (d, *J* = 7.6 Hz, 1H), 6.11 (s, 2H), 4.85 (dd, *J* = 10.0, 4.9 Hz, 2H), 4.26 – 4.10 (m, 3H), 3.87 – 3.72 (m, 4H), 3.17 (t, *J* = 5.5 Hz, 4H), 3.13 (t, *J* = 5.6 Hz, 4H), 3.03 (dt, *J* = 14.7, 7.0 Hz, 4H), 2.85 (t, *J* = 6.6 Hz, 4H), 2.49 – 2.37 (m, 4H), 2.20 – 1.70 (m, 17H), 1.68 – 1.06 (m, 23H), 0.92 – 0.57 (m, 24H).

ESI-MS, positive mode: m/z = 1300.8 $[M+H]^+$.

HRMS (ESI) calcd for $C_{73}H_{106}N_9O_{12}$ $[M+H]^+$ 1300.7955, found 1300.7953.

**4-630CP-PepA** (**95**):

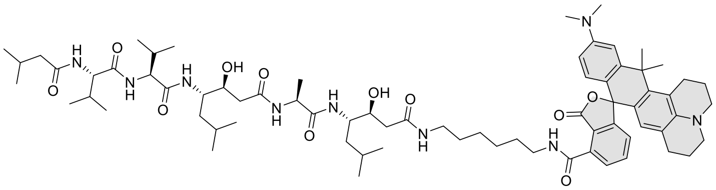

Compound was purified by preparative HPLC (solvent A: $H_2O$ + 10 mM $NH_4COOH$ pH = 3.6, solvent B: MeCN; temperature 25°C, gradient A:B - 3 min 70:30 isocratic, 4-20 min 70:30 to 0:100 gradient, 20-25 min 0:100 isocratic, 40 mL/min flow). Fractions containing the product were collected, evaporated and lyophilized from acetonitrile water mixture. The obtained solid was dissolved in 700 µL of $d_6$-DMSO. The samples from DMSO solution were diluted x100 in PBS +0.1% SDS and concentration was determined spectroscopically with Nanodrop. Determined stock concentration was 2.0 mM which constitutes to 28% yield (1.8 mg).

$^1$H NMR (400 MHz, $d_6$-DMSO) δ 9.16 (t, *J* = 5.6 Hz, 1H), 7.92 (d, *J* = 7.4 Hz, 1H), 7.83 – 7.63 (m, 5H), 7.46 (d, *J* = 8.8 Hz, 1H), 7.33 (d, *J* = 9.2 Hz, 1H), 7.00 (dd, *J* = 7.7, 0.9 Hz, 1H), 6.79 (d, *J* = 2.6 Hz, 1H), 6.57 (dd, *J* = 9.0, 2.5 Hz, 1H), 6.48 (d, *J* = 8.8 Hz, 1H), 6.14 (s, 1H), 4.85 (dd, *J* = 9.7, 5.0 Hz, 2H), 4.26 – 4.10 (m, 3H), 3.87 – 3.73 (m, 4H), 3.40 – 3.35 (m, 2H), 3.22 – 3.11 (m, 4H), 3.04 (p, *J* = 7.1 Hz, 2H), 3.01 – 2.78 (m,

8H), 2.48 – 2.36 (m, 2H), 2.31 – 1.82 (m, 16H), 1.82 (s, 3H), 1.77 – 1.67 (m, 2H), 1.65 – 1.17 (m, 21H), 0.89 – 0.70 (m, 24H).

ESI-MS, positive mode: m/z = 1274.9 $[M+H]^+$.

HRMS (ESI) calcd for $C_{72}H_{108}N_9O_{11}$ $[M+H]^+$ 1274.8163, found 1274.8162.

**4-642CP-PepA** (**96**):

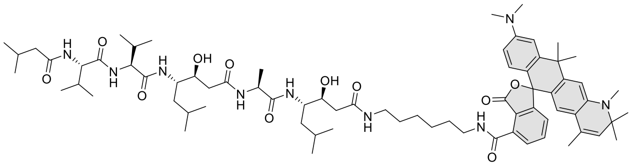

Compound was purified by preparative HPLC (solvent A: $H_2O$ + 10 mM $NH_4COOH$ pH = 3.6, solvent B: MeOH; temperature 25°C, gradient A:B - 3 min 70:30 isocratic, 4-20 min 70:30 to 0:100 gradient, 20-25 min 0:100 isocratic, 40 mL/min flow). Fractions containing the product were collected, evaporated and lyophilized from acetonitrile water mixture. The obtained solid was dissolved in 700 µL of $d_6$-DMSO. The samples from DMSO solution were diluted x100 in PBS +0.1% SDS and concentration was determined spectroscopically with Nanodrop. Determined stock concentration was 3.0 mM which constitutes to 42% yield (3.1 mg).

$^1$H NMR (400 MHz, $d_6$-DMSO) δ 9.17 (t, *J* = 5.6 Hz, 1H), 7.92 (d, *J* = 7.4 Hz, 1H), 7.84 – 7.67 (m, 5H), 7.47 (d, *J* = 8.9 Hz, 1H), 7.33 (d, *J* = 9.1 Hz, 1H), 7.10 (d, *J* = 7.6 Hz, 1H), 6.91 (s, 1H), 6.67 (s, 1H), 6.57 (s, 2H), 6.16 (s, 1H), 5.33 (d, *J* = 1.6 Hz, 1H), 4.85 (dd, *J* = 8.8, 5.1 Hz, 2H), 4.25 (t, *J* = 7.2 Hz, 1H), 4.18 (dd, *J* = 8.8, 7.3 Hz, 1H), 4.12 (dd, *J* = 8.9, 7.2 Hz, 1H), 3.85 – 3.72 (m, 4H), 3.38 – 3.35 (m, 2H), 3.04 (p, *J* = 6.7 Hz, 2H), 2.94 (s, 6H), 2.84 (s, 3H), 2.12 – 1.92 (m, 9H), 1.81 (s, 3H), 1.70 (s, 3H), 1.62 – 1.18 (m, 32H), 0.86 – 0.77 (m, 24H).

ESI-MS, positive mode: m/z = 1288.9 $[M+H]^+$.

HRMS (ESI) calcd for $C_{73}H_{110}N_9O_{11}$ $[M+H]^+$ 1288.8319, found 1288.8329.

**4-SiR-PepA (97):**

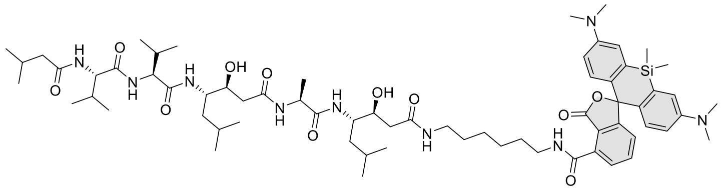

Compound was purified by preparative HPLC (solvent A: $H_2O$ + 0.2% HCOOH, solvent B: MeCN; temperature 25 °C, gradient A:B - 3 min 40:60 isocratic, 4-20 min 40:60 to 0:100 gradient, 20-25 min 0:100 isocratic, 40 mL/min flow). Fractions containing the product were collected, evaporated and lyophilized from acetonitrile water mixture to give product in 45% yield (2.8 mg) as a light blue solid.

$^1$H NMR (400 MHz, $d_6$-DMSO) δ 9.01 (t, *J* = 5.5 Hz, 1H), 7.91 (d, *J* = 7.4 Hz, 1H), 7.86 – 7.73 (m, 4H), 7.70 (t, *J* = 5.6 Hz, 1H), 7.46 (d, *J* = 8.8 Hz, 1H), 7.33 (d, *J* = 9.1 Hz, 1H), 7.24 (t, *J* = 4.4 Hz, 1H), 7.00 (d, *J* =

2.8 Hz, 2H), 6.70 (d, *J* = 9.0 Hz, 2H), 6.64 (dd, *J* = 9.0, 2.8 Hz, 2H), 4.84 (dd, *J* = 8.4, 5.0 Hz, 2H), 4.25 (t, *J* = 7.2 Hz, 1H), 4.18 (dd, *J* = 8.8, 7.3 Hz, 1H), 4.12 (dd, *J* = 8.9, 7.2 Hz, 1H), 3.85 – 3.73 (m, 4H), 3.30 – 3.20 (m, 2H), 3.10 – 3.01 (m, 2H), 2.92 (s, 12H), 2.13 – 1.90 (m, 9H), 1.72 – 1.03 (m, 23H), 0.88 – 0.77 (m, 24H), 0.62 (s, 3H), 0.52 (s, 3H).

ESI-MS, positive mode: m/z = 1260.7 $[M+Na]^+$.

HRMS (ESI) calcd for $C_{67}H_{103}N_9O_{11}SiNa$ $[M+Na]^+$ 1260.7439, found 1260.7410.

**4-685SiR-PepA (98):**

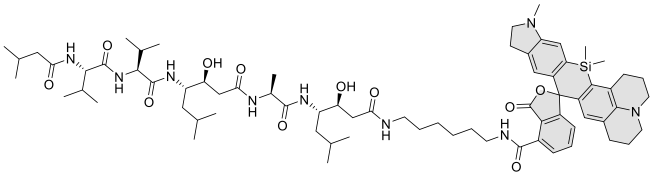

Compound was purified by preparative HPLC (solvent A: $H_2O$ +0.2% HCOOH, solvent B: MeCN; temperature 25°C, gradient A:B - 3 min 70:30 isocratic, 4-20 min 70:30 to 0:100 gradient, 20-25 min 0:100 isocratic, 40 mL/min flow). Fractions containing the product were collected, evaporated and lyophilized from acetonitrile water mixture. The obtained solid was dissolved in 700 µL of $d_6$-DMSO. The samples from DMSO solution were diluted x100 in PBS +0.1% SDS and concentration was determined spectroscopically with Nanodrop. Determined stock concentration was 3.2 mM which constitutes to 45% yield (2.9 mg).

$^1$H NMR (400 MHz, $d_6$-DMSO) δ 9.07 (t, *J* = 5.6 Hz, 1H), 7.92 (d, *J* = 7.2 Hz, 1H), 7.82 (d, *J* = 8.8 Hz, 1H), 7.78 (d, *J* = 9.0 Hz, 1H), 7.75 – 7.63 (m, 3H), 7.47 (d, *J* = 8.8 Hz, 1H), 7.34 (d, *J* = 9.1 Hz, 1H), 7.08 (dd, *J* = 7.3, 1.5 Hz, 1H), 6.74 (s, 1H), 6.55 (s, 1H), 6.32 (s, 1H), 4.89 – 4.81 (m, 2H), 4.25 (t, *J* = 7.2 Hz, 1H), 4.18 (dd, *J* = 8.8, 7.3 Hz, 1H), 4.12 (dd, *J* = 9.0, 7.2 Hz, 1H), 3.87 – 3.73 (m, 4H), 3.45 – 3.38 (m, 2H), 3.24 – 3.09 (m, 6H), 3.08 – 3.00 (m, 2H), 2.90 (t, *J* = 6.1 Hz, 2H), 2.82 – 2.66 (m, 5H), 2.43 (dd, *J* = 16.1, 6.4 Hz, 2H), 2.35 – 1.87 (m, 13H), 1.80 – 1.72 (m, 2H), 1.59 – 1.18 (m, 21H), 1.04 – 0.69 (m, 24H), 0.64 (s, 3H), 0.53 (s, 3H).

ESI-MS, positive mode: m/z = 1302.7 $[M+H]^+$.

HRMS (ESI) calcd for $C_{72}H_{108}N_9O_{11}Si$ $[M+H]^+$ 1302.7932, found 1302.7932.

**4-700SiR-PepA** (**99**):

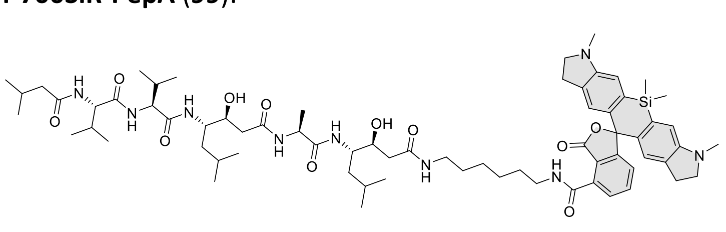

Compound was purified by preparative HPLC (solvent A: $H_2O$ +0.2% HCOOH, solvent B: MeCN; temperature 25°C, gradient A:B - 3 min 70:30 isocratic, 4-20 min 70:30 to 0:100 gradient, 20-25 min 0:100 isocratic, 40 mL/min flow). Fractions containing the product were collected, evaporated and lyophilized from acetonitrile water mixture. The obtained solid was dissolved in 700 µL of $d_6$-DMSO. The samples from DMSO solution were diluted x100 in PBS +0.1% SDS and concentration was determined spectroscopically with Nanodrop. Determined stock concentration was 2.8 mM which constitutes to 39% yield (2.5 mg).

$^1$H NMR (400 MHz, *d*$_6$-DMSO) δ 9.04 (t, *J* = 5.6 Hz, 1H), 7.92 (d, *J* = 7.4 Hz, 1H), 7.82 (d, *J* = 8.8 Hz, 1H), 7.78 (d, *J* = 9.0 Hz, 1H), 7.75 – 7.67 (m, 3H), 7.46 (d, *J* = 8.8 Hz, 1H), 7.33 (d, *J* = 9.2 Hz, 1H), 7.15 (dd, *J* = 6.5, 2.2 Hz, 1H), 6.78 (s, 2H), 6.58 (s, 2H), 4.84 (dd, *J* = 8.8, 5.0 Hz, 2H), 4.25 (t, *J* = 7.2 Hz, 1H), 4.18 (t, *J* = 9.4, 7.3 Hz, 1H), 4.12 (dd, *J* = 9.4, 7.1 Hz, 1H), 3.86 – 3.74 (m, 4H), 3.29 – 3.14 (m, 6H), 3.07 – 2.99 (m, 2H), 2.86 – 2.62 (m, 10H), 2.13 – 1.90 (m, 9H), 1.64 – 1.08 (m, 23H), 0.88 – 0.74 (m, 24H), 0.57 (s, 3H), 0.49 (s, 3H).

ESI-MS, positive mode: m/z = 1284.7 $[M+Na]^+$.

HRMS (ESI) calcd for $C_{69}H_{103}N_9O_{11}SiNa$ $[M+Na]^+$ 1284.7439, found 1284.7433.

<u>General procedure for the synthesis of compounds **100**-**104**:</u>

Into a solution of corresponding rhodamine dye (1 µmol, 2.7 eq; **3**, **4**, **11**, **15**, **22**), **des-bromo-des-methyl-Lys-jasplakinolide**[24] (250 µg, 0.37 µmol ,1.0 eq, unless stated otherwise) and DIPEA (5 µL) in DMSO (50 µL) a solution of HATU (1.3 µmol, 1.3 eq) in DMSO (20 µL) was added and the mixture was mixed for 30 min. Reactions were usually over after 30 min and their course was monitored by LC/MS analysis. Once reaction was finished it was quenched with 10 µL of formic acid and diluted with water and acetonitrile to 1 mL volume and was further purified by the means of preparative HPLC (preparative column: Agilent 5 Prep-C18, 5 µm, 100 x 50 mm).

**4-505R**-**JAS** (**100**):

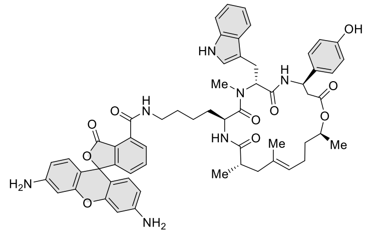

Compound was purified by preparative HPLC (solvent A: $H_2O$ 10 mM $NH_4COOH$ pH = 3.6, solvent B: MeCN; temperature 25°C, gradient A:B - 3 min 70:30 isocratic, 4-20 min 70:30 to 0:100 gradient, 20-25 min 0:100 isocratic, 40 mL/min flow). Fractions containing the product were collected, evaporated and lyophilized from acetonitrile water mixture. The obtained solid was dissolved in 600 µL of *d*$_6$-DMSO. The samples from DMSO solution were diluted x100 in PBS +0.1% SDS and concentration was determined spectroscopically with Nanodrop. Determined stock concentration was 0.31 mM which constitutes to 50% yield (192 µg).

$^1$H NMR (600 MHz, *d*$_6$-DMSO) δ 10.81 (s, 1H), 9.30 (s, 1H), 9.03 (t, *J* = 5.7 Hz, 1H), 8.65 (d, *J* = 8.8 Hz, 1H), 7.85 (dd, *J* = 7.5, 1.0 Hz, 1H), 7.76 (t, *J* = 7.6 Hz, 1H), 7.69 – 7.67 (m, 1H), 7.33 (d, *J* = 8.0 Hz, 1H),

7.25 (dd, *J* = 7.7, 1.0 Hz, 1H), 7.13 (d, *J* = 8.7 Hz, 2H), 7.09 (d, *J* = 2.3 Hz, 1H), 7.04 (t, *J* = 7.2 Hz, 1H), 6.95 (t, *J* = 7.5 Hz, 1H), 6.69 (d, *J* = 8.6 Hz, 2H), 6.53 (s, 1H), 6.43 (dd, *J* = 8.5, 6.9 Hz, 2H), 6.38 (d, *J* = 2.2 Hz, 2H), 6.29 (dd, *J* = 8.4, 2.2 Hz, 2H), 5.59 (s, 4H), 5.53 (dd, *J* = 11.5, 5.0 Hz, 1H), 5.46 (d, *J* = 9.5 Hz, 1H), 5.18 (t, *J* = 11.3 Hz, 1H), 4.92 (t, *J* = 7.0 Hz, 1H), 4.67 (h, *J* = 6.5 Hz, 1H), 4.58 (td, *J* = 8.8, 3.8 Hz, 1H), 3.19 – 3.01 (m, 6H), 2.93 (dd, *J* = 15.0, 4.9 Hz, 1H), 2.68 (dd, *J* = 14.8, 11.4 Hz, 1H), 2.62 – 2.57 (m, 2H), 2.40 – 2.37 (m, 1H), 2.18 – 2.13 (m, 1H), 1.84 (q, *J* = 16.0, 12.4 Hz, 2H), 1.70 (d, *J* = 15.0 Hz, 1H), 1.48 (s, 3H), 1.38 – 1.31 (m, 2H), 1.16 (d, *J* = 6.3 Hz, 3H), 0.98 – 0.92 (m, 2H), 0.88 – 0.79 (m, 5H), 0.76 – 0.67 (m, 2H).

ESI-MS, positive mode: m/z = 1030.4 $[M+H]^+$.

HRMS (ESI) calcd for $C_{59}H_{64}N_7O_{10}$ $[M+H]^+$ 1030.4709, found 1030.4714.

**4-TMR-JAS** (**101**):

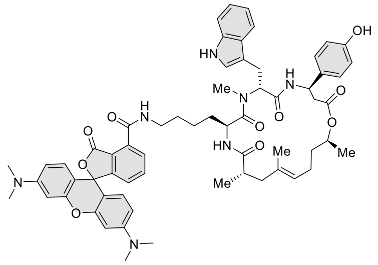

Compound was purified by preparative HPLC (solvent A: $H_2O$ + 10 mM $NH_4COOH$ pH = 3.6 B: MeCN; temperature 25°C, gradient A:B - 3 min 70:30 isocratic, 4-20 min 70:30 to 0:100 gradient, 20-25 min 0:100 isocratic, 40 mL/min flow). Fractions containing the product were collected, evaporated and lyophilized from acetonitrile water mixture. The obtained solid was dissolved in 600 µL of *d*6-DMSO. The samples from DMSO solution were diluted x100 in PBS +0.1% SDS and concentration was determined spectroscopically with Nanodrop. Determined stock concentration was 0.26 mM which constitutes to 42% yield (169 µg).

$^1$H NMR (600 MHz, $d_6$-DMSO) δ 10.82 (d, *J* = 2.3 Hz, 1H), 9.30 (s, 1H), 8.97 (t, *J* = 5.6 Hz, 1H), 8.66 (d, *J* = 8.9 Hz, 1H), 7.85 (dd, *J* = 7.5, 1.0 Hz, 1H), 7.76 (t, *J* = 7.6 Hz, 1H), 7.71 (d, *J* = 8.7 Hz, 1H), 7.69 (d, *J* = 8.0 Hz, 1H), 7.34 (d, *J* = 8.3 Hz, 1H), 7.24 (dd, *J* = 7.7, 1.0 Hz, 1H), 7.13 (d, *J* = 8.7 Hz, 2H), 7.10 (d, *J* = 2.4 Hz, 1H), 7.05 (t, *J* = 7.3 Hz, 1H), 6.96 (t, *J* = 7.7 Hz, 1H), 6.70 (d, *J* = 8.6 Hz, 2H), 6.62 (dd, *J* = 8.8, 3.0 Hz, 2H), 6.52 – 6.44 (m, 4H), 5.54 (dd, *J* = 11.6, 4.9 Hz, 1H), 5.21 – 5.16 (m, 1H), 4.92 (t, *J* = 7.0 Hz, 1H), 4.69 – 4.65 (m, 1H), 4.57 (td, *J* = 9.0, 3.8 Hz, 1H), 3.28 – 3.23 (m, 2H), 3.08 – 3.00 (m, 5H), 2.94 (d, *J* = 3.3 Hz, 12H), 2.68 (dd, *J* = 14.7, 11.6 Hz, 1H), 2.63 – 2.55 (m, 2H), 2.45 (d, *J* = 21.7 Hz, 2H), 2.39 (p, *J* = 1.8 Hz, 1H), 2.18 – 2.13 (m, 1H), 1.89 – 1.79 (m, 2H), 1.66 (d, *J* = 15.2 Hz, 2H), 1.48 (s, 3H), 1.39 – 1.34 (m, 2H), 1.16 (d, *J* = 6.3 Hz, 3H), 0.99 – 0.92 (m, 2H), 0.80 (d, *J* = 6.8 Hz, 3H).

ESI-MS, positive mode: m/z = 1086.5 $[M+H]^+$.

HRMS (ESI) calcd for $C_{63}H_{72}N_7O_{10}$ $[M+H]^+$ 1086.5335, found 1086.5297.

**4-610CP-JAS** (**102**):

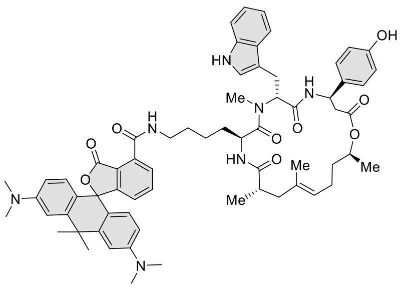

The reaction was carried from 100 µg of **des-bromo-des-methyl-Lys-jasplakinolide**[24] (0.15 µmol). Compound was purified by preparative HPLC (solvent A: $H_2O$ + 10 mM $NH_4COOH$ pH = 3.6, solvent B: MeCN; temperature 25°C, gradient A:B - 3 min 70:30 isocratic, 4-20 min 60:400 to 0:100 gradient, 20-25 min 0:100 isocratic, 40 mL/min flow). Fractions containing the product were collected, evaporated and lyophilized from acetonitrile water mixture. The obtained solid was dissolved in 100 µL of $d_6$-DMSO. The samples from DMSO solution were diluted x100 in PBS +0.1% SDS and concentration was determined spectroscopically with Nanodrop. Determined stock concentration was 0.42 mM which constitutes to 28% yield (46 µg).

The amount of product was not sufficient to obtain reasonable $^1$H NMR spectra. Therefore copies of LC/MS and HRMS are provided to support molecular structure and purity of the product.

ESI-MS, positive mode: m/z = 1112.5 $[M+H]^+$.

HRMS (ESI) calcd for $C_{66}H_{78}N_7O_9$ $[M+H]^+$ 1112.5856, found 1112.5848.

LC-MS analysis data:

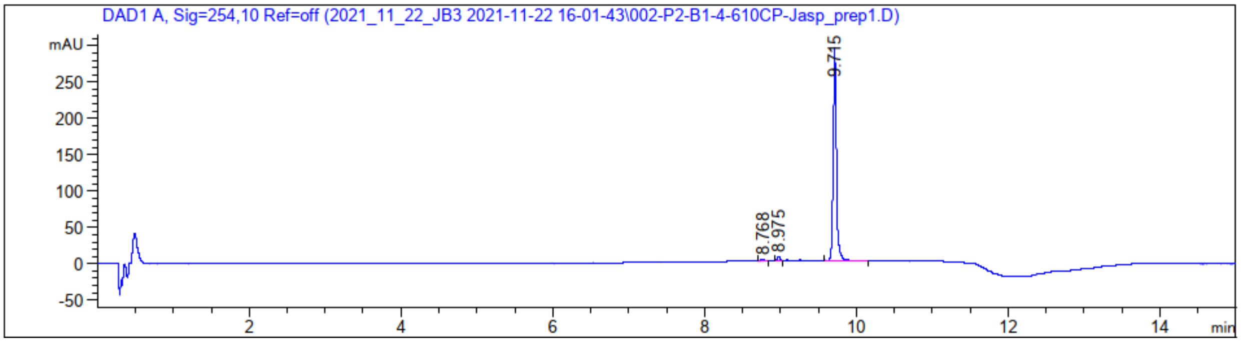

Signal 1: DAD1 A, Sig=254,10 Ref=off

| Peak # | RetTime [min] | Type | Width [min] | Area [mAU*s] | Height [mAU] | Area % |
|---|---|---|---|---|---|---|
| 1 | 8.768 | BB | 0.0452 | 8.57062 | 2.76319 | 0.9672 |
| 2 | 8.975 | BB | 0.0392 | 14.35857 | 5.74317 | 1.6203 |
| 3 | 9.715 | BB | 0.0444 | 863.21344 | 292.86401 | 97.4125 |

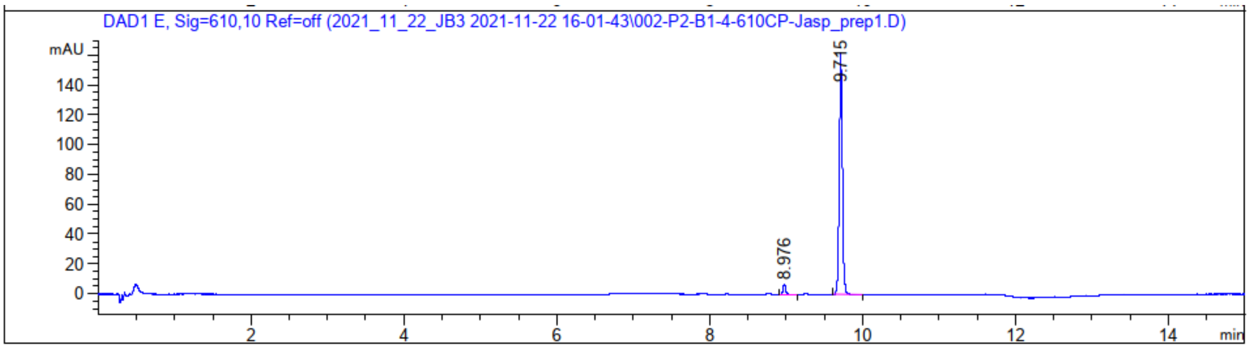

Signal 2: DAD1 E, Sig=610,10 Ref=off

| Peak # | RetTime [min] | Type | Width [min] | Area [mAU*s] | Height [mAU] | Area % |
|---|---|---|---|---|---|---|
| 1 | 8.976 | BB | 0.0414 | 17.91189 | 6.65488 | 3.7760 |
| 2 | 9.715 | BB | 0.0429 | 456.45078 | 162.02919 | 96.2240 |

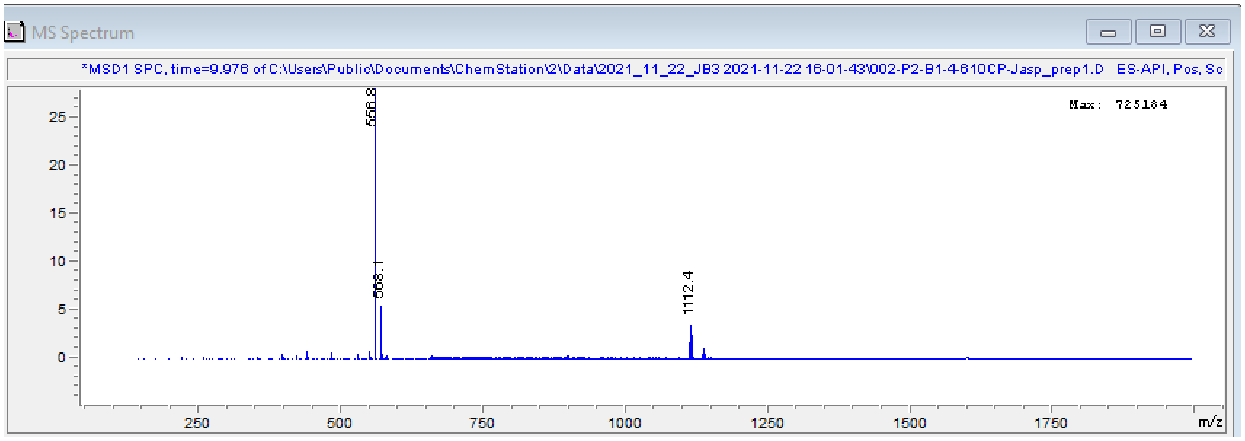

**4-630CP-JAS** (**103**):

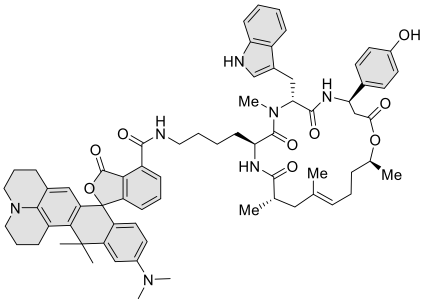

Compound was purified by preparative HPLC (solvent A: $H_2O$ + 10 mM $NH_4COOH$ pH = 3.6 B: MeCN; temperature 25°C, gradient A:B - 3 min 70:30 isocratic, 4-20 min 70:30 to 0:100 gradient, 20-25 min 0:100 isocratic, 40 mL/min flow). Fractions containing the product were collected, evaporated and lyophilized from acetonitrile water mixture. The obtained solid was dissolved in 600 µL of $d_6$-DMSO. The samples from DMSO solution were diluted x100 in PBS +0.1% SDS and concentration was determined spectroscopically with Nanodrop. Determined stock concentration was 0.28 mM which constitutes to 45% yield (194 µg).

$^1$H NMR (600 MHz, $d_6$-DMSO) δ 10.82 (s, 1H), 9.30 (s, 1H), 9.16 – 9.08 (m, 1H), 8.66 (dd, *J* = 8.8, 4.5 Hz, 1H), 7.79 (dd, *J* = 9.9, 7.4 Hz, 1H), 7.72 (dd, *J* = 8.7, 4.2 Hz, 1H), 7.70 – 7.63 (m, 2H), 7.34 (d, *J* = 8.1 Hz, 1H), 7.13 (d, *J* = 8.5 Hz, 2H), 7.10 (d, *J* = 2.3 Hz, 1H), 7.05 (t, *J* = 7.5 Hz, 1H), 7.00 (d, *J* = 7.8 Hz, 1H), 6.96 (t, *J* = 7.5 Hz, 1H), 6.80 (d, *J* = 2.5 Hz, 1H), 6.69 (d, *J* = 8.5 Hz, 2H), 6.55 – 6.49 (m, 2H), 6.14 (d, *J* = 2.8 Hz, 1H), 5.54 (dd, *J* = 11.6, 5.0 Hz, 1H), 5.18 (t, *J* = 11.3 Hz, 1H), 4.92 (t, *J* = 7.4 Hz, 1H), 4.67 (h, *J* = 6.8, 6.1, 6.0 Hz, 1H), 4.61 – 4.56 (m, 1H), 3.26 – 3.23 (m, 2H), 3.15 (t, *J* = 17.9, 6.2 Hz, 4H), 3.10 – 3.03 (m, 4H), 2.96 – 2.87 (m, 7H), 2.68 (dd, *J* = 14.6, 11.5 Hz, 1H), 2.62 – 2.58 (m, 2H), 2.54 – 2.52 (m, 4H), 2.47 (q, *J* = 1.9 Hz, 2H), 2.39 (q, *J* = 1.9 Hz, 1H), 2.18 – 2.13 (m, 1H), 1.96 – 1.86 (m, 5H), 1.83 (s, 3H), 1.54 – 1.43 (m, 5H), 1.40 – 1.34 (m, 2H), 1.21 (s, 3H), 1.16 (d, *J* = 6.3 Hz, 3H), 0.99 – 0.95 (m, 2H), 0.85 – 0.80 (m, 4H).

ESI-MS, positive mode: m/z = 1164.6 $[M+H]^+$.

HRMS (ESI) calcd for $C_{70}H_{82}N_7O_9$ $[M+H]^+$ 1164.6169, found 1164.6155.

**4-685SiR-JAS** (**104**):

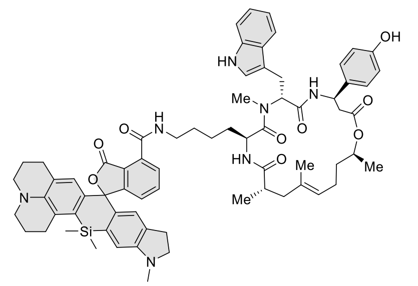

Compound was purified by preparative HPLC (solvent A: $H_2O$ + 10 mM $NH_4COOH$ pH = 3.6 B: MeCN; temperature 25°C, gradient A:B - 3 min 70:30 isocratic, 4-20 min 70:30 to 0:100 gradient, 20-25 min 0:100 isocratic, 40 mL/min flow). Fractions containing the product were collected, evaporated and lyophilized from acetonitrile water mixture. The obtained solid was dissolved in 600 µL of $d_6$-DMSO. The samples from DMSO solution were diluted x100 in PBS +0.1% SDS and concentration was determined spectroscopically with Nanodrop. Determined stock concentration was 0.24 mM which constitutes to 39% yield (172 µg).

$^1$H NMR (600 MHz, $d_6$-DMSO) δ 10.81 (s, 1H), 9.30 (s, 1H), 9.07 (t, *J* = 5.7 Hz, 1H), 8.65 (d, *J* = 8.8 Hz, 1H), 7.75 (dd, *J* = 7.5, 1.0 Hz, 1H), 7.70 – 7.64 (m, 3H), 7.34 (d, *J* = 8.1 Hz, 1H), 7.13 (d, *J* = 8.6 Hz, 2H), 7.11 – 7.08 (m, 2H), 7.04 (t, *J* = 7.8 Hz, 1H), 6.95 (t, *J* = 7.6 Hz, 1H), 6.75 (s, 1H), 6.69 (d, *J* = 8.6 Hz, 2H), 6.55 (d, *J* = 5.5 Hz, 1H), 6.33 (d, *J* = 9.6 Hz, 1H), 5.53 (dd, *J* = 11.5, 5.1 Hz, 1H), 5.18 (t, *J* = 12.0, 11.6 Hz, 1H), 4.92 (t, *J* = 7.2 Hz, 1H), 4.67 (h, *J* = 6.4 Hz, 1H), 4.59 (td, *J* = 8.8, 4.1 Hz, 1H), 3.24 – 3.08 (m, 11H), 3.06 (s, 3H), 2.95 (d, *J* = 5.0 Hz, 1H), 2.92 – 2.89 (m, 2H), 2.76 (d, *J* = 1.0 Hz, 3H), 2.73 – 2.65 (m, 3H), 2.62 – 2.58 (m, 2H), 2.47 – 2.42 (m, 2H), 2.38 (p, *J* = 1.9 Hz, 1H), 2.18 – 2.14 (m, 1H), 1.96 – 1.92 (m, 2H), 1.87 – 1.80 (m, 2H), 1.75 (q, *J* = 6.1 Hz, 2H), 1.69 (d, *J* = 15.8 Hz, 1H), 1.48 (s, 3H), 1.40 – 1.34 (m, 2H), 1.28 – 1.26 (m, 2H), 1.21 (s, 3H), 1.16 (d, *J* = 6.4 Hz, 4H), 0.95 (q, *J* = 6.9 Hz, 2H), 0.75 – 0.68 (m, 2H), 0.64 (s, 3H), 0.53 (s, 3H).

ESI-MS, positive mode: m/z = 1192.5 $[M+H]^+$.

HRMS (ESI) calcd for $C_{70}H_{82}N_7O_9Si$ $[M+H]^+$ 1192.5901, found 1192.5901.

General procedure for the synthesis of compounds **105**-**109**:

Into a solution of corresponding rhodamine dye (5 µmol, 1 eq;**4, 7, 10, 11, 24**) and DIPEA (5 µL) in DMSO (100 µL) a solution of HATU (6.5 µmol, 1.3 eq) in DMSO (100 µL) was added and the mixture was mixed for 1 min. Then a solution of O6-[4-(aminomethyl)benzyl]guanine (7.5 µmol ,1.5 eq, Sigma Aldrich) in DMSO (100 µL +5 µL DIPEA) was added at once. Reactions were usually over after 30 min and their course was monitored by LC/MS analysis. Once reaction was finished it was quenched with 20 µL of formic acid and diluted with water and acetonitrile to 2 mL volume and was further purified by the means of preparative HPLC (preparative column: Agilent 5 Prep-C18, 5 µm, 100 x 50 mm).

**4-580CP-BG** (**105**):

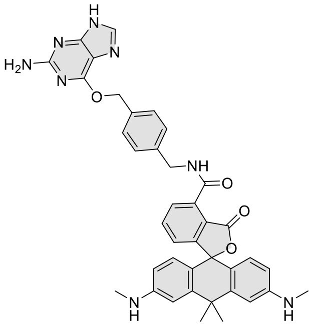

Compound was purified by preparative HPLC (solvent A: $H_2O$ + 10 mM $NH_4COOH$ pH = 3.6 B: MeCN; temperature 25°C, gradient A:B - 3 min 70:30 isocratic, 4-20 min 70:30 to 0:100 gradient, 20-25 min 0:100 isocratic, 40 mL/min flow). Fractions containing the product were collected, evaporated and lyophilized from acetonitrile water mixture. The obtained solid was dissolved in 700 µL of $d_6$-DMSO. The samples from DMSO solution were diluted x100 in PBS +0.1% SDS and concentration was determined spectroscopically with Nanodrop. Determined stock concentration was 4.2 mM which constitutes to 59% yield (2.0 mg).

$^1$H NMR (400 MHz, $d_6$-DMSO) δ 12.42 (s, 1H), 9.58 (t, $J$ = 5.7 Hz, 1H), 8.48 (s, 1H), 7.81 – 7.79 (m, 1H), 7.70 (t, $J$ = 7.6 Hz, 1H), 7.50 (s, 4H), 7.06 (dd, $J$ = 7.7, 1.0 Hz, 1H), 6.75 (d, $J$ = 2.3 Hz, 2H), 6.48 (d, $J$ = 8.6 Hz, 2H), 6.37 (dd, $J$ = 8.7, 2.3 Hz, 2H), 6.28 (s, 2H), 5.85 (q, $J$ = 5.1 Hz, 2H), 5.48 (s, 2H), 4.60 (d, $J$ = 5.8 Hz, 2H), 2.70 (d, $J$ = 5.0 Hz, 6H), 1.75 (s, 3H), 1.66 (s, 3H).

ESI-MS, positive mode: m/z = 703.3 $[M+Na]^+$.

HRMS (ESI) calcd for $C_{39}H_{36}N_8O_4Na$ $[M+Na]^+$ 703.2752, found 703.2726.

**4-610CP-BG** (**106**):

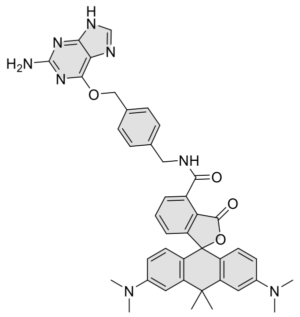

Compound was purified by preparative HPLC (solvent A: $H_2O$ + 10 mM $NH_4COOH$ pH = 3.6 B: MeCN; temperature 25°C, gradient A:B - 3 min 70:30 isocratic, 4-20 min 70:30 to 0:100 gradient, 20-25 min 0:100 isocratic, 40 mL/min flow). Fractions containing the product were collected, evaporated and lyophilized from acetonitrile water mixture. The obtained solid was dissolved in 700 µL of *d*₆-DMSO. The samples from DMSO solution were diluted x100 in PBS +0.1% SDS and concentration was determined spectroscopically with Nanodrop. Determined stock concentration was 3.2 mM which constitutes to 45% yield (1.6 mg).

$^1$H NMR (400 MHz, $d_6$-DMSO) δ 12.43 (s, 1H), 9.57 (t, *J* = 5.8 Hz, 1H), 8.53 (s, 1H), 7.80 (dd, *J* = 7.5, 1.0 Hz, 1H), 7.70 (t, *J* = 7.6 Hz, 1H), 7.50 (s, 4H), 7.05 (dd, *J* = 7.6, 1.0 Hz, 1H), 6.92 – 6.89 (m, 2H), 6.61 – 6.55 (m, 4H), 6.28 (s, 2H), 5.49 (s, 2H), 4.60 (d, *J* = 5.8 Hz, 2H), 2.94 (s, 12H), 1.82 (s, 3H), 1.72 (s, 3H).

ESI-MS, positive mode: m/z = 731.3 $[M+Na]^+$.

HRMS (ESI) calcd for $C_{41}H_{40}N_8O_4Na$ $[M+Na]^+$ 731.3065, found 731.3059.

**4-625CP-BG** (**107**):

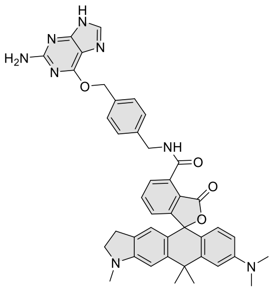

Compound was purified by preparative HPLC (solvent A: $H_2O$ + 10 mM $NH_4COOH$ pH = 3.6 B: MeCN; temperature 25°C, gradient A:B - 3 min 70:30 isocratic, 4-20 min 70:30 to 0:100 gradient, 20-25 min 0:100 isocratic, 40 mL/min flow). Fractions containing the product were collected, evaporated and lyophilized from acetonitrile water mixture. The obtained solid was dissolved in 700 µL of $d_6$-DMSO. The samples from DMSO solution were diluted x100 in PBS +0.1% SDS and concentration was determined spectroscopically with Nanodrop. Determined stock concentration was 3.8 mM which constitutes to 53% yield (1.9 mg).

$^1$H NMR (400 MHz, $d_6$-DMSO) δ 9.59 (t, *J* = 5.8 Hz, 1H), 7.83 – 7.75 (m, 2H), 7.69 (t, *J* = 7.6 Hz, 1H), 7.51 (s, 4H), 7.04 (dd, *J* = 7.7, 1.0 Hz, 1H), 6.89 (s, 1H), 6.76 (s, 1H), 6.60 – 6.52 (m, 2H), 6.40 (s, 1H), 6.27 (s, 2H), 5.49 (s, 2H), 4.61 (d, *J* = 5.8 Hz, 2H), 3.27 – 3.23 (m, 2H), 2.93 (s, 6H), 2.79 (s, 3H), 2.76 – 2.67 (m, 2H), 1.79 (s, 3H), 1.71 (s, 3H).

ESI-MS, positive mode: m/z = 721.3 $[M+H]^+$.

HRMS (ESI) calcd for $C_{42}H_{41}N_8O_4$ $[M+H]^+$ 721.3245, found 721.3238.

**4-630CP-BG** (**108**):

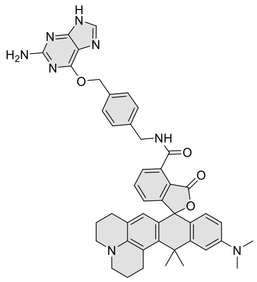

Compound was purified by preparative HPLC (solvent A: $H_2O$ + 10 mM $NH_4COOH$ pH = 3.6 B: MeCN; temperature 25°C, gradient A:B - 3 min 70:30 isocratic, 4-20 min 70:30 to 0:100 gradient, 20-25 min 0:100 isocratic, 40 mL/min flow). Fractions containing the product were collected, evaporated and lyophilized from acetonitrile water mixture. The obtained solid was dissolved in 700 µL of *d*$_6$-DMSO. The samples from DMSO solution were diluted x100 in PBS +0.1% SDS and concentration was determined spectroscopically with Nanodrop. Determined stock concentration was 2.3 mM which constitutes to 32% yield (1.2 mg).

$^1$H NMR (400 MHz, *d*$_6$-DMSO) δ 12.39 (s, 1H), 9.61 (t, *J* = 5.7 Hz, 1H), 7.78 (dd, *J* = 7.5, 1.0 Hz, 2H), 7.67 (t, *J* = 7.6 Hz, 1H), 7.51 (s, 4H), 7.01 (dd, *J* = 7.7, 1.0 Hz, 1H), 6.79 (d, *J* = 2.5 Hz, 1H), 6.56 (dd, *J* = 8.9, 2.5 Hz, 1H), 6.50 (d, *J* = 8.9 Hz, 1H), 6.28 (s, 2H), 6.17 (s, 1H), 5.49 (s, 2H), 4.61 (d, *J* = 5.8 Hz, 2H), 3.20 – 3.11 (m, 4H), 2.99 – 2.86 (m, 8H), 2.47 – 2.36 (m, 2H), 1.94 – 1.89 (m, 2H), 1.89 (s, 3H), 1.83 (s, 3H), 1.79 – 1.72 (m, 2H).

ESI-MS, positive mode: m/z = 761.3 [M+H]+.

HRMS (ESI) calcd for $C_{45}H_{45}N_8O_4$ $[M+H]^+$ 761.3558, found 761.3545.

**4-640CP-BG** (**109**):

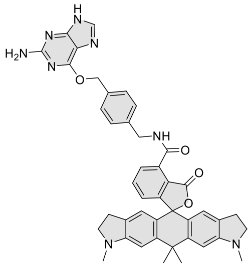

Compound was purified by preparative HPLC (solvent A: $H_2O$ + 10 mM $NH_4COOH$ pH = 3.6 B: MeCN; temperature 25°C, gradient A:B - 3 min 70:30 isocratic, 4-20 min 70:30 to 0:100 gradient, 20-25 min 0:100 isocratic, 40 mL/min flow). Fractions containing the product were collected, evaporated and lyophilized from acetonitrile water mixture. The obtained solid was dissolved in 700 µL of *d*$_6$-DMSO. The samples from DMSO solution were diluted x100 in PBS +0.1% SDS and concentration was determined spectroscopically with Nanodrop. Determined stock concentration was 4.0 mM which constitutes to 56% yield (2.0 mg).

$^1$H NMR (400 MHz, *d*$_6$-DMSO) δ 9.60 (t, *J* = 5.8 Hz, 1H), 7.82 (s, 1H), 7.79 (dd, *J* = 7.5, 1.0 Hz, 1H), 7.69 (t, *J* = 7.6 Hz, 1H), 7.51 (s, 4H), 7.04 (dd, *J* = 7.7, 1.0 Hz, 1H), 6.74 (s, 2H), 6.37 (s, 2H), 6.26 (s, 2H), 5.49 (s, 2H), 4.61 (d, *J* = 5.8 Hz, 2H), 3.28 – 3.18 (m, 4H), 2.78 (s, 6H), 2.76 – 2.64 (m, 4H), 1.76 (s, 3H), 1.69 (s, 3H).

ESI-MS, positive mode: m/z = 733.3 [M+H]+.

HRMS (ESI) calcd for $C_{43}H_{41}N_8O_4$ $[M+H]^+$ 733.3245, found 733.3236.

### References

1. Åkerlöf, G.; Short, A. O., The Dielectric Constant of Dioxane—Water Mixtures between 0 and 80°. *J. Am. Chem. Soc.* **1936,** *58* (7), 1241–1243.
2. Grimm, J. B.; Tkachuk, A. N.; Xie, L.; Choi, H.; Mohar, B.; Falco, N.; Schaefer, K.; Patel, R.; Zheng, Q.; Liu, Z.; Lippincott-Schwartz, J.; Brown, T. A.; Lavis, L. D., A general method to optimize and functionalize red-shifted rhodamine dyes. *Nat Methods* **2020,** *17* (8), 815-821.
3. Frisch, M. J.; Trucks, G. W.; Schlegel, H. B.; Scuseria, G. E.; Robb, M. A.; Cheeseman, J. R.; Scalmani, G.; Barone, V.; Petersson, G. A.; Nakatsuji, H.; Li, X.; Caricato, M.; Marenich, A. V.; Bloino, J.; Janesko, B. G.; Gomperts, R.; Mennucci, B.; Hratchian, H. P.; Ortiz, J. V.; Izmaylov, A. F.; Sonnenberg, J. L.; Williams; Ding, F.; Lipparini, F.; Egidi, F.; Goings, J.; Peng, B.; Petrone, A.; Henderson, T.; Ranasinghe, D.; Zakrzewski, V. G.; Gao, J.; Rega, N.; Zheng, G.; Liang, W.; Hada, M.; Ehara, M.; Toyota, K.; Fukuda, R.; Hasegawa, J.; Ishida, M.; Nakajima, T.; Honda, Y.; Kitao, O.; Nakai, H.; Vreven, T.; Throssell, K.; Montgomery Jr., J. A.; Peralta, J. E.; Ogliaro, F.; Bearpark, M. J.; Heyd, J. J.; Brothers, E. N.; Kudin, K. N.; Staroverov, V. N.; Keith, T. A.; Kobayashi, R.; Normand, J.; Raghavachari, K.; Rendell, A. P.; Burant, J. C.; Iyengar, S. S.; Tomasi, J.; Cossi, M.; Millam, J. M.; Klene, M.; Adamo, C.; Cammi, R.; Ochterski, J. W.; Martin, R. L.; Morokuma, K.; Farkas, O.; Foresman, J. B.; Fox, D. J. *Gaussian 09 Rev. D.01*, Wallingford, CT, 2009.
4. Becke, A. D., Density-functional thermochemistry. III. The role of exact exchange. *The Journal of Chemical Physics* **1993,** *98* (7), 5648-5652.
5. Lee, C.; Yang, W.; Parr, R. G., Development of the Colle-Salvetti correlation-energy formula into a functional of the electron density. *Phys Rev B Condens Matter* **1988,** *37* (2), 785-789.
6. Davidson, E. R.; Feller, D., Basis set selection for molecular calculations. *Chemical Reviews* **1986,** *86* (4), 681-696.
7. Tomasi, J.; Mennucci, B.; Cammi, R., Quantum Mechanical Continuum Solvation Models. *Chemical Reviews* **2005,** *105* (8), 2999-3094.
8. Lukinavicius, G.; Lavogina, D.; Orpinell, M.; Umezawa, K.; Reymond, L.; Garin, N.; Gonczy, P.; Johnsson, K., Selective chemical crosslinking reveals a Cep57-Cep63-Cep152 centrosomal complex. *Curr Biol* **2013,** *23* (3), 265-70.
9. Bach, M.; Grigat, S.; Pawlik, B.; Fork, C.; Utermohlen, O.; Pal, S.; Banczyk, D.; Lazar, A.; Schomig, E.; Grundemann, D., Fast set-up of doxycycline-inducible protein expression in human cell lines with a single plasmid based on Epstein-Barr virus replication and the simple tetracycline repressor. *FEBS J* **2007,** *274* (3), 783-90.
10. D'Este, E.; Kamin, D.; Gottfert, F.; El-Hady, A.; Hell, S. W., STED nanoscopy reveals the ubiquity of subcortical cytoskeleton periodicity in living neurons. *Cell Rep* **2015,** *10* (8), 1246-51.
11. Molyneux, P. C.; Pyle, L. A.; Dillon, M.; Harrington, M. E., A Mouse Primary Hepatocyte Culture Model for Studies of Circadian Oscillation. *Curr Protoc Mouse Biol* **2015,** *5* (4), 311-329.
12. Schindelin, J.; Arganda-Carreras, I.; Frise, E.; Kaynig, V.; Longair, M.; Pietzsch, T.; Preibisch, S.; Rueden, C.; Saalfeld, S.; Schmid, B.; Tinevez, J. Y.; White, D. J.; Hartenstein, V.; Eliceiri, K.; Tomancak, P.; Cardona, A., Fiji: an open-source platform for biological-image analysis. *Nat Methods* **2012,** *9* (7), 676-82.
13. Gottlieb, H. E.; Kotlyar, V.; Nudelman, A., NMR Chemical Shifts of Common Laboratory Solvents as Trace Impurities. *J Org Chem* **1997,** *62* (21), 7512-7515.
14. Gribble, G. W.; Nutaitis, C. F., Reactions of Sodium Borohydride in Acidic Media; XVI.1 N-Methylation of Amines with Paraformaldehyde/Trifluoroacetic Acid. *Synthesis* **1987,** *1987* (08), 709-711.
15. Mayer, R. J.; Hampel, N.; Mayer, P.; Ofial, A. R.; Mayr, H., Synthesis, Structure, and Properties of Amino-Substituted Benzhydrylium Ions – A Link between Ordinary Carbocations and Neutral Electrophiles. *European Journal of Organic Chemistry* **2019,** *2019* (2-3), 412-421.
16. Hanaoka, K.; Kagami, Y.; Piao, W.; Myochin, T.; Numasawa, K.; Kuriki, Y.; Ikeno, T.; Ueno, T.; Komatsu, T.; Terai, T.; Nagano, T.; Urano, Y., Synthesis of unsymmetrical Si-rhodamine fluorophores and

application to a far-red to near-infrared fluorescence probe for hypoxia. *Chem Commun (Camb)* **2018,** *54* (50), 6939-6942.
17. Grimm, J. B.; Sung, A. J.; Legant, W. R.; Hulamm, P.; Matlosz, S. M.; Betzig, E.; Lavis, L. D., Carbofluoresceins and carborhodamines as scaffolds for high-contrast fluorogenic probes. *ACS Chem Biol* **2013,** *8* (6), 1303-10.
18. Pastierik, T.; Sebej, P.; Medalova, J.; Stacko, P.; Klan, P., Near-infrared fluorescent 9-phenylethynylpyronin analogues for bioimaging. *J Org Chem* **2014,** *79* (8), 3374-82.
19. Bachman, J. L.; Escamilla, P. R.; Boley, A. J.; Pavlich, C. I.; Anslyn, E. V., Improved Xanthone Synthesis, Stepwise Chemical Redox Cycling. *Org Lett* **2019,** *21* (1), 206-209.
20. Bucevicius, J.; Kostiuk, G.; Gerasimaite, R.; Gilat, T.; Lukinavicius, G., Enhancing the biocompatibility of rhodamine fluorescent probes by a neighbouring group effect. *Chem Sci* **2020,** *11* (28), 7313-7323.
21. Butkevich, A. N., Modular Synthetic Approach to Silicon-Rhodamine Homologues and Analogues via Bis-aryllanthanum Reagents. *Org Lett* **2021,** *23* (7), 2604-2609.
22. Benink, H.; McDougall, M.; Klaubert, D.; Los, G., Direct pH measurements by using subcellular targeting of 5(and 6-) carboxyseminaphthorhodafluor in mammalian cells. *Biotechniques* **2009,** *47* (3), 769-74.
23. Bucevicius, J.; Keller-Findeisen, J.; Gilat, T.; Hell, S. W.; Lukinavicius, G., Rhodamine-Hoechst positional isomers for highly efficient staining of heterochromatin. *Chem Sci* **2019,** *10* (7), 1962-1970.
24. Belov, V. N.; Stoldt, S.; Ruttger, F.; John, M.; Seikowski, J.; Schimpfhauser, J.; Hell, S. W., Synthesis of Fluorescent Jasplakinolide Analogues for Live-Cell STED Microscopy of Actin. *J Org Chem* **2020,** *85* (11), 7267-7275.
